# BRCA1-A and LIG4 complexes mediate ecDNA biogenesis and cancer drug resistance

**DOI:** 10.1101/2025.02.18.638901

**Authors:** Oliver W. Chung, Shun Yao, Fu Yang, Ling Wang, Christian Cerda-Smith, Haley M. Hutchinson, Kris C. Wood, Weijia Su, Mustafa Khasraw, Lee Zou, Dale A. Ramsden, ZZ Zhao Zhang

**Author notes:** These authors contributed equally (Oliver W. Chung, Shun Yao, Fu Yang).

## Abstract

Extrachromosomal circular DNA (ecDNA) are commonly produced within the nucleus to drive genome dynamics and heterogeneity, enabling cancer cell evolution and adaptation. However, the mechanisms underlying ecDNA biogenesis remain poorly understood. Here using genome-wide CRISPR screening in human cells, we identified the BRCA1-A and the LIG4 complexes mediate ecDNA production. Following DNA fragmentation, the upstream BRCA1-A complex protects DNA ends from excessive resection, promoting end-joining for circularization. Conversely, the MRN complex, which mediates end resection and thus antagonizes the BRCA1-A complex, suppresses ecDNA formation. Downstream, LIG4 conservatively catalyzes ecDNA production in *Drosophila* and mammals, with patient tumor ecDNA harboring junctions marked by LIG4 activity. Notably, disrupting LIG4 or BRCA1-A in cancer cells impairs ecDNA-mediated adaptation, hindering resistance to both chemotherapy and targeted therapies. Together, our study reveals the roles of the LIG4 and BRCA1-A complexes in ecDNA biogenesis, and uncovers new therapeutic targets to block ecDNA-mediated adaptation for cancer treatment.

## INTRODUCTION

Although originally reported six decades ago, extrachromosomal circular DNA (henceforth, abbreviated as “ecDNA” for all circles regardless of size) have recently emerged as a common DNA form broadly produced in eukaryotes^1–4^. With the size ranging from hundreds to millions of bases, ecDNA often harbor genes or genetic regulatory elements^3–10^. Together with the feature of unrestricted copy numbers––often up to hundreds of copies per cell, ecDNA bring one important layer of genome dynamics that allows cells to fully exploit their genetic information for adaptation and evolution^11,12^. Reflecting this fundamental function, ecDNA have been documented to play pivotal roles in diverse processes: magnifying histone genes in yeast to achieve dose compensation ^13^, amplifying rDNA to drive frog egg development^1,14^, and re-writing the cattle genome for coat color sidedness patterning^12^.

Notably, forming ecDNA is a common mechanism for the amplification of oncogenes and genes that render cancer cells drug resistant^3,4,15^. Strikingly, nearly one-in-three cancer patients harbor ecDNA in their cancer cells^3,7,8,11,16–19^. This percentage increases even further in certain aggressive cancer types: to 60% in glioblastoma and ∼40-50% in sarcoma and esophageal cancers^8,19^. Given their high copy numbers and decompacted chromatin structure, ecDNA can massively amplify and express oncogenes residing on them^3,7,8,11,16–18^. Meanwhile, ecDNA have shown strong structural and spatial dynamics, in which they can hop on and off the chromosomes during cancer evolution^16,20^. As such, charactering the molecular mechanisms underlying ecDNA biology and correspondingly drugging these processes could provide new therapeutic strategies for cancer treatment.

Despite their essential functions, our understanding of the ecDNA formation process is limited. Our previous work elucidated the mechanism by which the retrotransposons residing in the host genome form ecDNA during their life cycle^21^. These DNA circles represent replication products and typically only harbor retrotransposon sequences^21^. Given that retrotransposons are often silenced in the host genome^22–24^, retrotransposon-derived ecDNA are not frequently produced. A more common process of ecDNA biogenesis is directly circularizing one or more DNA fragments generated from chromosomal breaks^3,4,15^. This process appears to be more prevalent in cancer cells, especially when these cells undergo genome rupture, such as chromothripsis^16,17^. However, the molecular mechanisms that drive this process remain largely unexplored.

In this study we employed the genome-wide CRISPR screening to identify factors mediating ecDNA biogenesis. We found ecDNA formation is orchestrated by BRCA1-A and LIG4 complexes. BRCA1-A complex acts upstream by protecting DNA free ends from resection and then hands off these DNA molecules to LIG4 for end-to-end joining, resulting in ecDNA formation. Importantly, mutations in LIG4 or components of the BRCA1-A complex abrogate ecDNA-mediated adaptation to cancer therapeutic drugs. Our findings provide a crucial step in decoding the mechanism of ecDNA formation and offer promising drug targets for addressing therapeutic resistance in cancer.

## RESULTS

### CRISPR screens identify ecDNA biogenesis regulators

Current tools for studying and observing ecDNA are limited. To address this limitation, the CRISPR-C approach, using CRISPR/Cas9 cleavage to generate DNA fragments with the promoter located downstream of eGFP at the opposite end, was invented^25^. Under this design, circularization of the fragments brings the promoter upstream of eGFP, thus giving rise to fluorescence that reflects ecDNA formation (Figure 1A)^25^. Given that eGFP expression can be used to reflect ecDNA biogenesis, we sought to perform genome-wide CRISPR screening to identify factors regulating this process. For the screen, we used HEK293T cells harboring the eGFP reporter to mutate one gene/cell for 18,761 genes, by transducing the genome-wide MinLib CRISPR lentiviral library^26^. Upon inducing ecDNA production and eGFP expression, eGFP-positive and eGFP-negative cells were sorted to achieve 1,000-fold coverage of the sgRNA library. Under this design, sgRNAs targeting genes required for ecDNA biogenesis were enriched in the eGFP-negative population and excluded from the eGFP-positive population (Figure 1B). Conversely, sgRNAs targeting genes encoding factors that suppress ecDNA formation would show the reverse pattern (Figure 1B).

**Figure 1.**
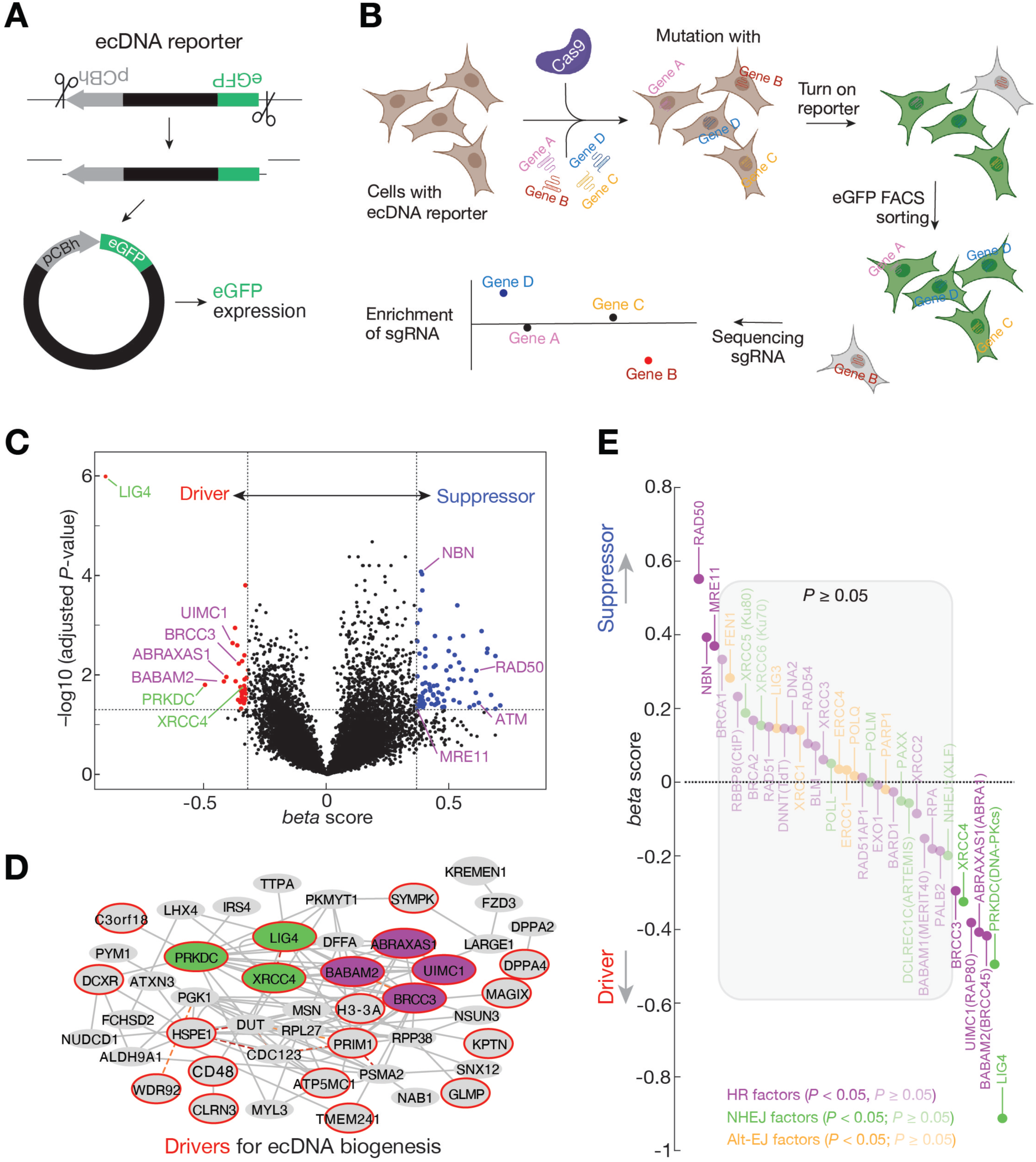
CRISPR screening to identify factors regulating ecDNA biogenesis. (A) Schematic design of the reporter used for monitoring ecDNA production. (B) Schematic design of the genome-wide CRISPR screens, which were performed in 3 biological replicates. (C) Volcano plot to show the regulators of ecDNA biogenesis. The factors further investigated in this study are labelled. (D) Interactome analysis of the factors that drive ecDNA biogenesis. LIG4 and BRCA1-A complexes are highlighted by green and magenta respectively. Factors identified as drivers in panel C are circled with red. Factors that show co-essentiality are connected with dashed red lines. (E) Snake plot to show the potential function of factors from distinct DNA break repair pathways in ecDNA biogenesis. See also Figure S1.

Given that ecDNA biogenesis is coupled with the DNA breakage, factors from DNA repair pathways were the most significantly enriched screen hits (Figure 1C and S1A). Meanwhile, we also detected factors from other processes as potential regulators for ecDNA biogenesis (Figure S1). For example, histone variant and factors related with rRNA processing or ribosome biogenesis may regulate ecDNA production under our screen design (Figure 1D and S1). While these findings point to directions for future investigations, here we primarily focus on the function of DNA repair factors in ecDNA biogenesis.

Among our top hits for driving ecDNA production are LIG4 and its physical partner XRCC4, and DNA-PKcs (encoded by the *PRKDC* gene) (Figure 1C-1E). Notably, DNA-PKcs has previously been implicated in mediating ecDNA production when cancer cells undergo chromothripsis^16,27,28^, giving credence to our screening approach. Despite the LIG4/XRCC4 complex and DNA-PKcs being classified into the Non-homologous end joining (NHEJ) DNA repair process^29,30^, the other factors from this pathway were not uncovered as significant hits from our screen (Figure 1E, see discussion). Meanwhile, although previous research has suggested the potential function of alternative end joining (alt-EJ) in circular DNA formation under specific settings^28,31,32^, our screening results showed that all factors from this DNA repair process are dispensable for ecDNA biogenesis from the designed reporter (Figure 1E).

Meanwhile, the other top hits for driving ecDNA biogenesis are factors can form a complex: BRCC36 encoded by gene *BRCC3*, BRCC45/BRE encoded by *BABAM2*, ABRA1/ABRAXAS encoded by *ABRAXAS1*, and RAP80 encoded by *UIMC1* (Figure 1C-1E). Together with MERIT40, these four factors physically interact with each other to build a complex that can sequester BRCA1 protein^33^. This complex thus is named as BRCA1-A complex. It is worth noting that BRCA1 itself was not screened as a regulator, suggesting that its canonical function in promoting homologous recombination–– executed by forming BRCA1-C complex––is unessential for ecDNA formation. Screening four different components from the same complex as strong hits indicates its pivotal role in mediating the circularization process. The BRCA1-A complex has been characterized for its function in preventing end-resection from the DNA break sites^34–37^. This suggests that preventing end-resection upon DNA segmentation promotes ecDNA production. Satisfyingly, our screen identified factors that drive end-resection as suppressors of ecDNA biogenesis (Figure 1C-1E and S1B). These factors are MRE11, NBN1, and RAD50 (Figure 1C-1E and S1B), which form the MRN complex that resects the DNA break ends and recruits ATM^38^. Consistently, ATM is screened as a negative regulator of ecDNA production (Figure 1C and S1B). In summary, our screen results suggest that upon DNA fragmentation, BRCA1-A complex prevents end resection to drive the ecDNA production process that is potentially catalyzed by LIG4.

### Validating BRCA1-A, MRN, and LIG4 complexes

To validate the screen hits that serve as drivers for ecDNA production, we used CRISPR/Cas9 on the eGFP reporter cells to generate mutant cell lines for the following seven genes: *LIG4*, *XRCC4*, *PRKDC*, *UIMC1*, *BRCC3*, *BABAM2*, and *ABRAXAS1* (Figure S2A and S2B). For each factor, we employed at least two different sgRNAs to generate distinct mutant clones (Figure S2A and S2B), minimizing the possibility of obtaining false-positive findings from the off-targeting mutation by an individual sgRNA. Using eGFP expression as the proxy for ecDNA production from the reporter, we obtained findings consistent with our screen results. Mutating any of the component from either LIG4 or BRCA1-A complex showed no eGFP expression (Figure S3), suggesting that the cells without either complex lost the capability of ecDNA formation. Besides using eGFP to indicate ecDNA production, we also designed qPCR or digital droplet PCR (ddPCR) assay to precisely quantify the number of ecDNA produced from the reporter (Figure 2A-2C). Consistent with eGFP expression, mutating *LIG4*, *XRCC4*, *PRKDC*, *BRCC3*, *BABAM2*, *ABRAXAS1*, *UIMC1* abrogated ecDNA production, as measured by either qPCR or ddPCR assay (Figure 2A-2C).

**Figure 2.**
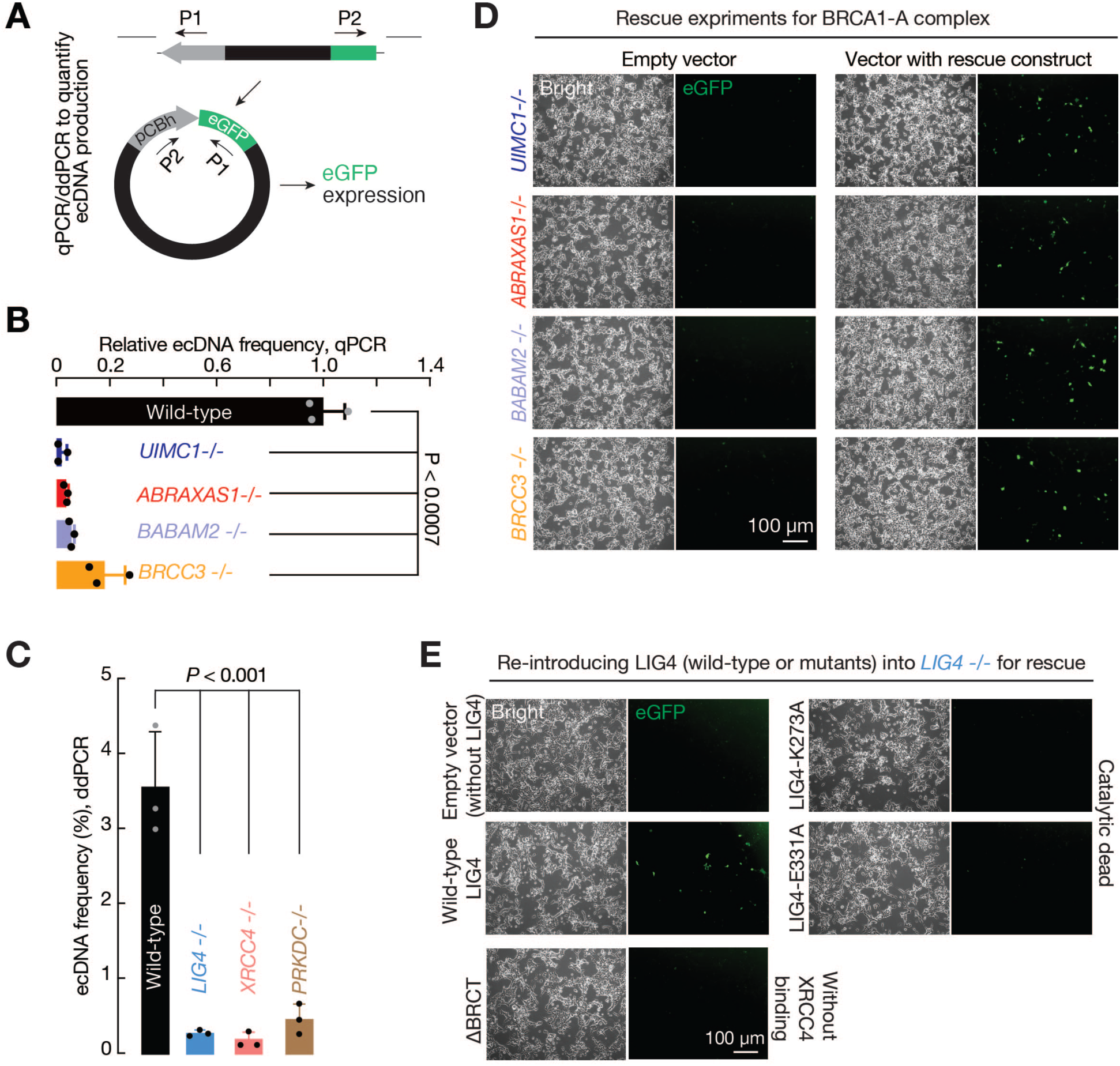
BRCA1-A and LIG4 complexes drives ecDNA biogenesis from the reporter. (A) Schematic design of the qPCR and Droplet digital PCR (ddPCR) assays to quantify ecDNA production from the pre-integrated reporter. (B) TaqMan qPCR to quantify ecDNA production when BRCA1-A function is disrupted. (C) ddPCR to quantify ecDNA production when LIG4 complex is mutated. For both panel B and C, the bars report mean ± standard deviation from three biological replicates (n=3). *P-*values were calculated with a two-tailed, two-sample unequal variance *t* test. For each gene, the mutant cell line generated by sgRNA-1 was picked for qPCR. (D) Re-introducing wild-type genes into corresponding BRCA1-A mutants rescues ecDNA production from the pre-integrated reporter, as evidenced by the restoration of eGFP expression. (E) Re-introducing wild-type *LIG4* rescues ecDNA production from the pre-integrated reporter. Mutant versions of *LIG4*––either catalytically dead (E331A or K273A) or with the XRCC4-binding BRCT domains deleted––fail to rescue. Each rescue construct achieves a similar level of protein expression (Figure S2D). See also Figure S2A-2D, S3, and S4.

To finally validate the screen outcome, we performed the rescue experiment by re-introducing either wild-type gene of *LIG4*, *BRCC3*, *BABAM2*, *ABRAXAS1*, and *UIMC1* via a plasmid vector into corresponding mutant cells (Figure 2D, 2E, S2C, and S2D). For LIG4, we also generated catalytically-dead LIG4 transgene for the rescue experiment to examine its ligase function in ecDNA formation (Figure S2D). Meanwhile, since XRCC4 stabilizes LIG4 by directly interacting with its two BRCT domains ^39,40^, we additionally included a mutant version of LIG4 by removing the BRCT domains––termed as LIG4-ΔBRCT––for the rescue experiment (Figure S2D). As expected, re-introducing wild-type *LIG4, BRCC3*, *BABAM2*, *ABRAXAS1*, and *UIMC1* restored ecDNA production in the mutant cells, as evidenced by eGFP expression from the reporter (Figure 2D, 2E, S2C, and S2D). However, either mutating LIG4 ligase activity or suppressing its interaction with XRCC4 abolished the rescue outcome (Figure 2E and S2D). These findings indicate a pivotal role of LIG4 and BRCA1-A complex in circularizing the reporter.

On the other end, we sought to validate the MRN complex as a suppressor of ecDNA production via two independent approaches. First, we generated two *RAD50* mutant cell lines using two distinct sgRNAs (Figure S4A). Upon mutating *RAD50*, the ecDNA-producing cells significantly increased from 21.2% to 37.4% (*P*-value = 0.030, sgRNA-1) and 36% (*P*-value = 0.046, sgRNA-2, Figure S4A and S4B). Next, we further validated the antagonizing function of the MRN complex by using a small molecule drug, Mirin, which serves as an MRN inhibitor to block the end-resection function of MRE11^41,42^. Applying this inhibitor significantly increased the eGFP-positive population by 2.7–fold (*P*-value = 0.004, Figure S4C). Altogether, our results robustly validated the screen findings and support a model that suppressing end-resection by BRCA1-A complex promotes LIG4-mediated ecDNA biogenesis.

### BRCA1-A and LIG4 drive ecDNA formation in cancer cell lines

All data described above were collected based on the CRISPR-C reporter, which relies on CRISPR/Cas9 to generate a DNA segment for ecDNA formation. Although unlikely, it is possible that LIG4 or BRCA1-A complex is essential for DNA fragmentation by Cas9, thus leading to no ecDNA production upon its mutation. To exclude this possibility, we adapted the original CRIPSR-C system to directly introduce linear DNA molecules that mimic the fragments generated by CRISPR/Cas9 cleavage (Figure 3A). Under this design, the sequences of linear DNA can be easily modified. Moreover, another advantage is that these linear molecules can be introduced into any cells without a pre-integration event, allowing us to monitor ecDNA production in different cell types. As such, we named our design as “Versatile biosensor”. As a proof-of-principal, we initially designed the biosensor with just a promoter and eGFP (Figure S5A). Transfecting these DNA molecules into HEK293T cells gave eGFP positive cells, suggesting the formation of ecDNA (Figure S5B).

**Figure 3.**
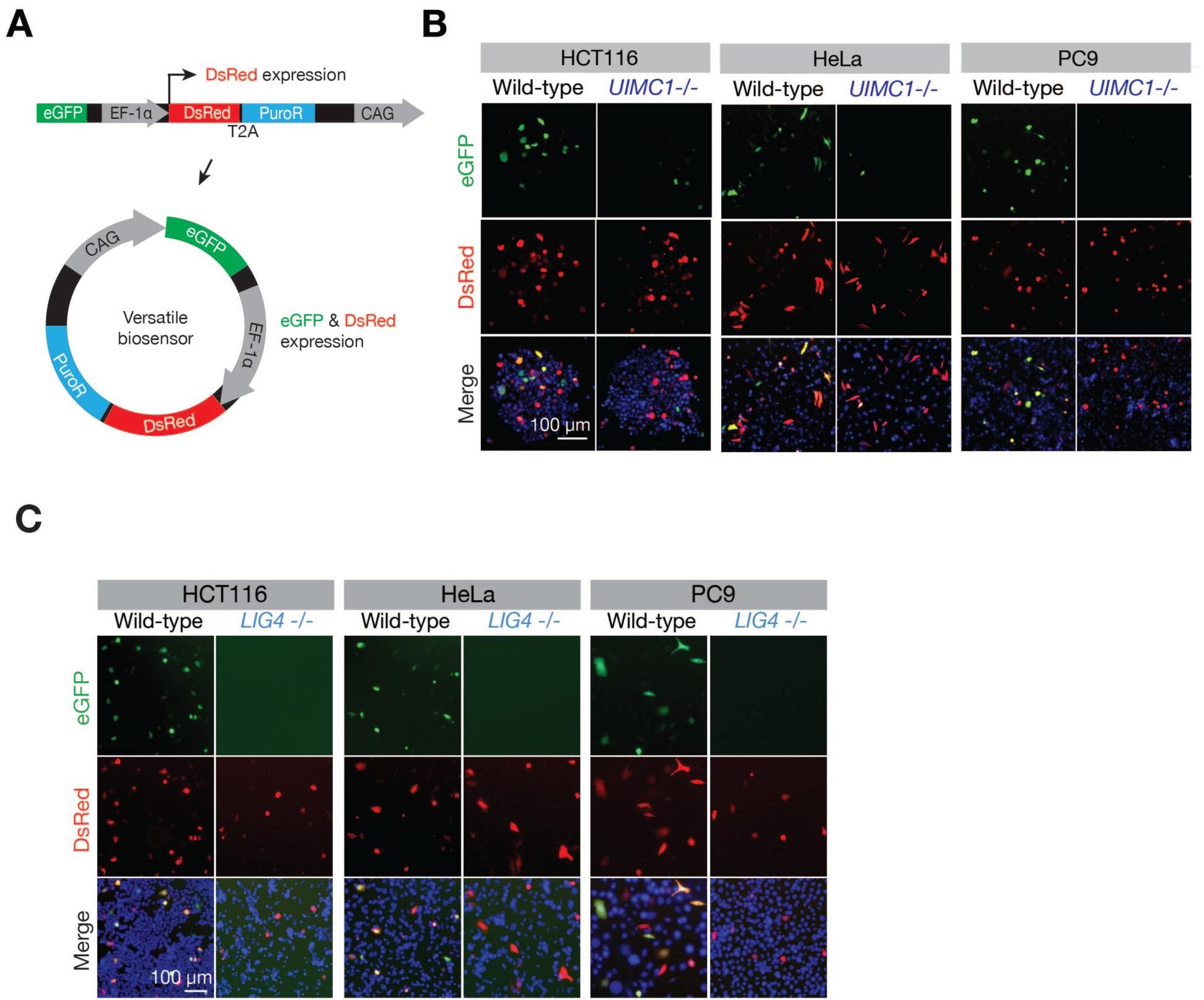
BRCA1-A and LIG4 complexes drives ecDNA biogenesis in cancer cells. (A) Schematic design of the versatile biosensor. Circularization brings the CAG promoter upstream of the eGFP sequences to initiate eGFP expression. EF-1α promoter drives the expression of DsRed and the puromycin resistance gene, enabling selection of the cells harboring the biosensors. (B) BRCA1-A complex drives ecDNA production in cancer cells: HCT116, HeLa, and PC9. (C) LIG4 drives ecDNA production in cancer cells: HCT116, HeLa, and PC9. See also Figure S2E, S2F, and S5.

To validate that eGFP expression from our versatile biosensor reflects ecDNA production, and is not due to the introduced eGFP sequences integrating into the host cell genome with a local promoter driving eGFP expression, we designed two other control constructs (Figure S5B). One is just the eGFP sequence without a promoter (Figure S5B). The other is a circle harboring the designed biosensor (Figure S5B), thus preventing the joining of the eGFP and promoter. Introducing either construct into cultured cells could not give eGFP expression (Figure S5B), suggesting eGFP signals from our biosensor reflect circle formation. Meanwhile, given that cells can take up multiple biosensor molecules, their concatenation can form linear DNA and bring a promoter upstream of eGFP. To test this possibility, we treated DNA from the eGFP-positive cells with exonuclease, which removes linear DNA but leaves ecDNA intact. By using a pair of primers that spans the junction site of eGFP and promoter (Figure S5A), we observed that the PCR amplicon signals were preserved upon exonuclease treatment (Figure S5C), indicating that our biosensor mainly forms ecDNA.

Since eGFP expression reflects the circularization process from our versatile biosensor, we sought to test the function of LIG4 and BRCA1-A complex for ecDNA biogenesis in different cancer lines when Cas9-mediated DNA fragmentation is bypassed. Given that these cancer lines have different efficiency on uptake our engineered biosensor, we introduced a DsRed cassette that is directly driven by its own promoter into the middle of the biosensor (Figure 3A). This allows us to measure the percentage of cells harboring biosensors upon delivery. With this design, we next tested the function of LIG4 and BRCA1-A complex in three cancer cell lines: cervical cancer HeLa cells, non-small cell lung cancer PC9 cells, and colon cancer HCT116 cells. In all cell lines, mutating *LIG4* or the BRCA1-A complex key component *UIMC1* led to the abolishment of ecDNA production (Figure 3B, 3C, S2E, and S2F), suggesting a generic function of LIG4 and BRCA1-A complex in mediating ecDNA biogenesis across different cell types.

Lastly, while our findings suggest LIG4 and BRCA1-A complex as the key factors driving ecDNA formation, these findings were based on the reporter system that produced DNA fragments with fixed DNA sequences at the two ends. Are LIG4 and BRCA1-A still essential if the DNA sequences at the two ends are altered? To address this question, we harnessed the versatility of our ecDNA biosensor design by including 6 random nucleotides at the two ends (Figure S5D). In all cells tested (HCT116, PC9, and HeLa cells), loss the function of BRCA1-A by mutating *UIMC1* or mutating *LIG4* led to abrogation of ecDNA formation from the biosensor with random end sequences (Figure S5E). Altogether, our findings suggest that when cells experience DNA fragmentation, LIG4 and BRCA1-A complex drives ecDNA biogenesis regardless of the nucleotide composition at the two ends.

### LIG4 mediates natural ecDNA formation in vivo

Given that LIG4 is conserved in *Drosophila*, we sought to test the function of LIG4 in driving natural ecDNA formation in vivo. During *Drosophila* egg development, two chorion genomic clusters undergo massive amplification in the oocyte-surrounding follicle cells to immensely produce chorion protein for egg encapsulation^43^. Previous studies showed that the uneven amplification of chorion clusters often leads to DNA breaks with variable length (Figure 4A) ^44^. We previously sequenced all circular DNA in *Drosophila* ovaries to examine retrotransposon-derived ecDNA^21^. Mining our dataset, we found sequencing reads with head-to-tail junctions that support ecDNA generated from the two chorion clusters (Figure S6A). To investigate whether LIG4 is required for the formation of chorion ecDNA, we sequenced ovarian ecDNA from *LIG4* homozygotes (*Lig4^-/-^*) and heterozygotes (*Lig4^-/+^*). While the *LIG4* heterozygotes still produced ecDNA from both chorion regions, loss of LIG4 resulted in a nearly 100% abolishment of ecDNA production (Figure 4B and S6B). To further validate our findings, we designed a pair of primers that can only yield PCR products when ecDNA is generated (Figure 4A and 4C). Similarly, while ecDNA with variable length can be detected from *Lig4^-/+^* ovaries, *Lig4^-/-^*flies lost chorion-ecDNA production (Figure 4C).

**Figure 4.**
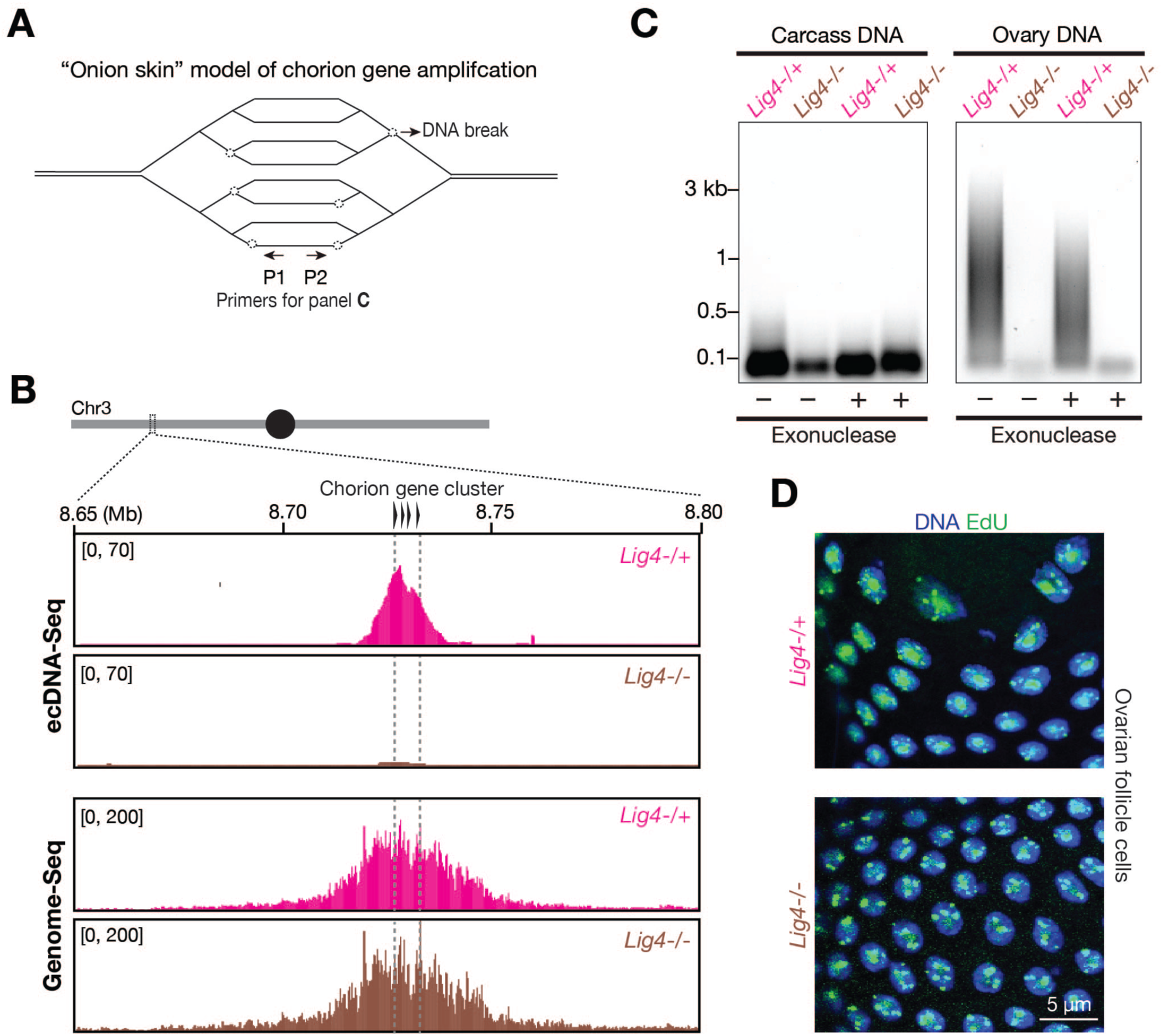
LIG4 drives natural ecDNA biogenesis in vivo. (A) Cartoon depicting the “Onion skin” model of chorion gene amplification occurring in *Drosophila* ovarian follicle cells. Dashed circles stand for DNA breaks. (B) ecDNA-Seq and Genome-Seq to measure the amplification of the chorion locus on the 3^rd^ chromosome and ecDNA production from this region. (C) PCR-based assay to measure the production of chorion ecDNA from *Drosophila* ovaries. Fly bodies without ovaries were used as the carcass for DNA extraction, serving as a negative control. (D) EdU incorporating into the replicated DNA in *Drosophila* ovarian follicle cells. *Lig4* mutation has no impact on DNA amplification, as indicated by EdU immunostaining. See also Figure S6A and S6B.

While our findings suggest LIG4 directly drives chorion ecDNA production in vivo, we are cognizant of the possibility that LIG4 is essential for chorion region amplification, thus its depletion would abolish ecDNA biogenesis. To test this possibility, we sequenced the ovarian genomic DNA from both *Lig4^-/-^* and *Lig4^-/+^* flies. Regardless of the genotype, the two chorion clusters amplified to the same magnitude (Figure 4B and S6B), suggesting that LIG4 is dispensable for the genomic amplification of these two regions. The amplification process of these two regions can incorporate a large amount of EdU into the replicated DNA and be visualized by EdU staining. Consistently, we observed comparable EdU foci in the follicle cells from both *Lig4^-/-^* and *Lig4^-/+^* flies (Figure 4D), further indicating that mutating *LIG4* does not alter chorion cluster amplification but does abolish ecDNA production. In summary, our data suggest that LIG4 plays a conserved function in driving natural ecDNA production during *Drosophila* egg development. Given that *Drosophila* does not possess BRCA1-A complex, we were unable to test its function under this oogenesis setting.

### Glioblastoma ecDNA harbor end-end junctions with a LIG4 signature

ecDNA are frequently produced in cancer cells and have been extensively shown to contribute to tumor heterogeneity and cancer cell adaptation^3–10^. Since 0-3 bp microhomology is the strong signature of LIG4-mediated ligation process ^45,46^, this can be used to test whether LIG4 likely mediates ecDNA formation in patient tumors by examining junction site sequences. Previous research assessed the possible contribution of distinct DNA repair pathways in mediating ecDNA formation by using datasets from The Cancer Genome Atlas, which often lack paired germline genome from the same patients to serve as controls^8^. Given glioblastoma shows the highest frequency for ecDNA production^8^, we sequenced tumor DNA from 36 patients with 100X genome coverage to reconstruct ecDNA and to identify the junction sites (Figure 5A). Meanwhile, we sequenced blood DNA from the same patients to build the personalized germline genome (Figure 5A), which serves as the reference for ecDNA and mutation detection. Among these patients, 21 (58%) harbor cancer-related ecDNA within their tumors (Figure 5B). Similar as previously reported^8^, EGFR is the gene that most frequently amplified via ecDNA formation: EGFR ecDNA are abundantly produced in the tumors of 14 patients (39%, Figure 5B). From all ecDNA, we in total detected 190 junction sites for microhomology analysis (Figure 5C-5E). Analyzing the sequences of junction sites revealed a strong LIG4 signature: 43 sites without homology sequences from the two ends (supporting direct ligation events); 51 sites with 1 bp homology; 41 sites with 2 bp homology; and 26 sites with 3 bp homology (Figure 5D and 5E). In total, 85% of the junction sites are highly likely to be catalyzed by LIG4 (Figure 5E). Meanwhile, we also detected 15 sites with 4 bp homology; 9 sites with 5 bp; and 5 sites with homology sequences ≥ 6 bp (Figure 5E). These 15% events were potentially driven by either LIG4 or Polθ, as both can mediate the repair process that leaves a footprint with > 3 bp homology ^47,48^. In summary, our data support LIG4 to be the major driver for ecDNA formation within glioblastoma tumor cells.

**Figure 5.**
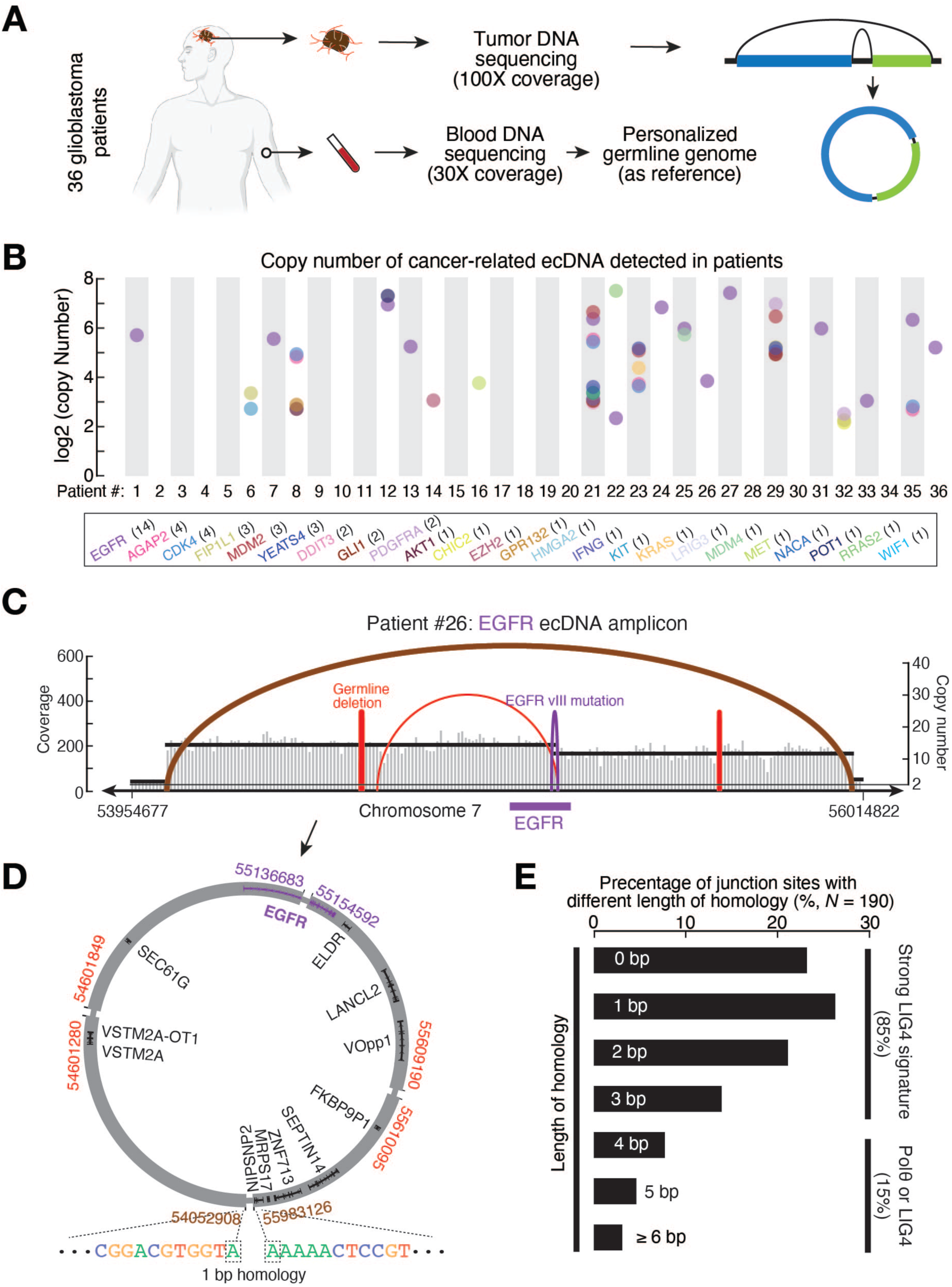
LIG4 likely mediates ecDNA formation in glioblastoma. (A) Cartoon depicting the experiment design to sequence ecDNA from tumor tissues from 36 glioblastoma patients. (B) Copy number of cancer-related ecDNA being detected in glioblastoma from each patient. Color indicates the oncogene detected on the circle. Bottom black box serves as the color key, and the total number of patients harboring the labelled oncogene on ecDNA was shown in the parenthesis. For example, 14 patients having EGFR on their ecDNA. (C) Copy number and breakpoints of the EGFR amplicon from patient #26, which serves as a representative example. The brown arc labels the junction site for circularization. The purple arc illustrates EGFR vIII mutation occurs in tumor tissues, but not in blood DNA. The left red arc labels a germline deletion, which can be detected in blood DNA. The right red arc indicates a 905 bp deletion occurs in tumor tissues. (D) Reconstructed ecDNA from the EGFR amplicon showing in panel C. The junction sequences from the two ends share 1-bp homology. (E) The bar graph to show the percentage of junction sites harboring distinct length of homologies.

### BRCA1-A complex protects the DNA ends from resection

We next aimed to dissect the function of BRCA1-A complex in ecDNA formation. The aforementioned data demonstrated the function of BRCA1-A complex in circularizing DNA less than 10 kb. Given that the ecDNA driving cancer cell evolution often reach megabases, we next examined the function of BRCA1-A complex in large circle formation. Previous research established the CRISPR-C approach to generate megabase-sized ecDNA species^11^. We employed similar approach to generate a 1.8 mega-base segment in PC9 cells harboring *EGFR* (Figure 6A), an oncogene frequently forming massive amount of ecDNA in cancer patients, such as glioblastoma^8,49^. Wild-type PC9 cells generated 1.8 mega-bases *EGFR* ecDNA (Figure 6B). Treating these cells with Osimertinib, the 3^rd^-generation EGFR inhibitor that has been widely used in clinics, led to accumulation of a massive amount of *EGFR* ecDNA (Figure 6C and 6D), suggesting that these circles confer drug resistance. However, in *UIMC1* mutant cells with BRCA1-A function disrupted, the amount of ecDNA produced significantly dropped to 15% of the wild-type level (Figure 6B; P < 0.0001, measured by ddPCR). Correspondingly, loss of BRCA1-A complex led to complete cell-death upon Osimertinib exposure (Figure 6C).

**Figure 6.**
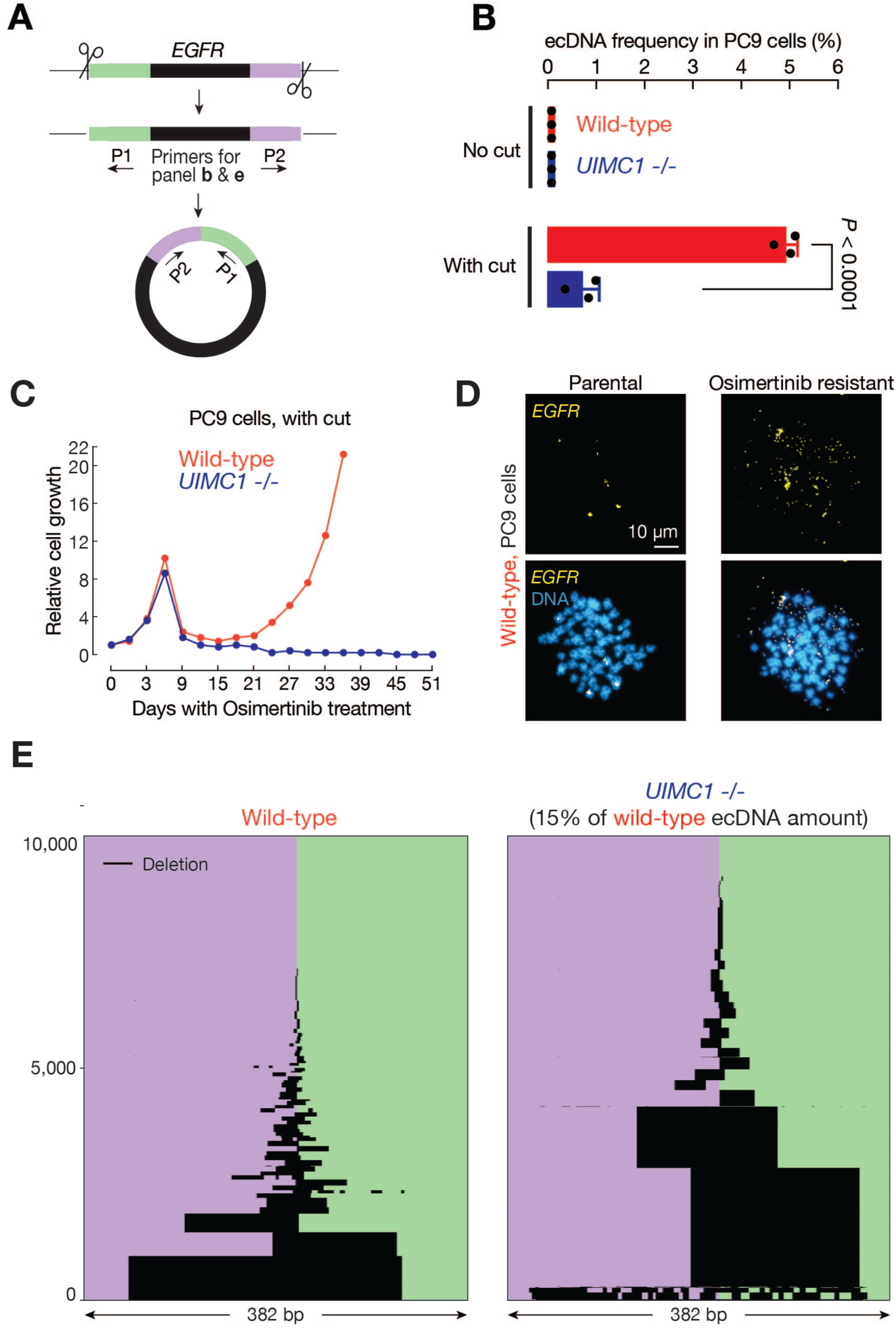
BRCA1-A complex prevents end-resection to drive ecDNA formation. (A) Schematic design of the CRISPR-C approach to generate mega-bases *EGFR* ecDNA in PC9 lung cancer cells. (B) ddPCR to quantify ecDNA production from the *EGFR* region in PC9 cells. The bars report mean ± standard deviation from three biological replicates (n = 3). *P*-values were calculated with a two-tailed, two-sample unequal variance *t* test. (C) Cell growth curve upon Osimertinib treatment. CRISPR-C approach was applied to generate *EGFR* ecDNA in PC9 cells before Osimertinib exposure. (D) DNA-FISH to measure *EGFR* ecDNA in parental and Osimertinib-resistant cells. (E) High-throughput sequencing to survey the end-resection events at the end-end junction site. Ten thousands reads were randomly selected for figure display. Black lines reflect the deletion regions from each read. See also Figure S6C.

Does BRCA1-A complex protect the two ends of DNA segments from resection, thus promoting ecDNA formation? To seek an answer, we sought to examine the sequences of the end-end junction site of ecDNA to search the “footprint” of the resection events––nucleotide deletion. For this purpose, we amplified the junction region and employed high-throughput sequencing to survey the junction sequences of *EGFR* ecDNA generated in PC9 cells from precisely defined DNA segment. In wild-type cells, direct ligation of the two free ends is the dominant form: taking 25.2% of total events (Figure 6E). Meanwhile, we also observed ecDNA with deletions at variable lengths (Figure 6E), suggesting that end-resection is an active process even in wild-type cells.

Given the collapse of ecDNA biogenesis upon loss of BRCA1-A complex (Figure 6B), examining the end-end junction sites from trace amount of ecDNA produced under this condition is technically challenging. By using a 5-times more *UIMC1* mutant cells to perform the experiment, we managed to obtain enough ecDNA for sequencing-library preparation. As expected, the ecDNA species from this condition dropped to only 49% of wild-type level, indicating sever deficiency in ecDNA formation. From these ecDNA, the percentage of having direct ligation dropped from 25.2% in wild-type to 3.7% (Figure 6E and S6C). Meanwhile, we also observed dramatic increase of ecDNA with large deletions at the junction site upon loss of BRCA1-A complex (Figure 6E and S6C). The junction sites with > 50 bp deletions increased from 27.4% in wild-type to 42% upon loss of RAP80 by mutating *UIMC1* (Figure S6C). For the sites with > 100 bp deletions, the frequency increased from 20.6% to 42% (Figure S6C). This suggests the occurrence of excessive resection without BRCA1-A complex. In summary, our data suggest without the protection of BRCA1-A complex, DNA segments are more prone for resection at the ends and followed by degradation, as reflected by the abolishment of ecDNA production.

### BRCA1-A and LIG4 drive ecDNA-mediated cancer cell adaptation

The CRISPR-C approach relies on CRISPR/Cas9 to generate pre-defined DNA fragments for ecDNA biogenesis. Without CRISPR/Cas9, is BRCA1-A complex still required for natural ecDNA biogenesis in cancer cells and enabling drug resistance? To answer this question, we tested the function of BRCA1-A complex in two systems: (1) methotrexate (MTX) induced spontaneous *DHFR* ecDNA production in HeLa cells; and (2) EGFR inhibitor induced natural *EGFR* ecDNA biogenesis in PC9 cells for drug resistance (Figure 7A). Pioneering research in 1970s established the system that upon methotrexate treatment, HeLa cells rely on ecDNA––at that time termed “double minutes”––harboring the *DHFR* gene to evolve resistance for this chemotherapy drug^50–53^. We adapted this system and tested the function of BRCA1-A complex during this natural ecDNA-mediated evolution process (Figure 7B-7D). Wild-type cells evolved resistance to methotrexate by producing *DHFR* ecDNA, which enable the expression of *DHFR* increased from 53 fpkm in parental cells to 1748 fpkm in methotrexate resistant cells (Figure 7B, 7C, S7A, and S7B). Mutating *UIMC1* to disrupt BRCA1-A function led to the abolishment of resistance development (Figure 7D).

**Figure 7.**
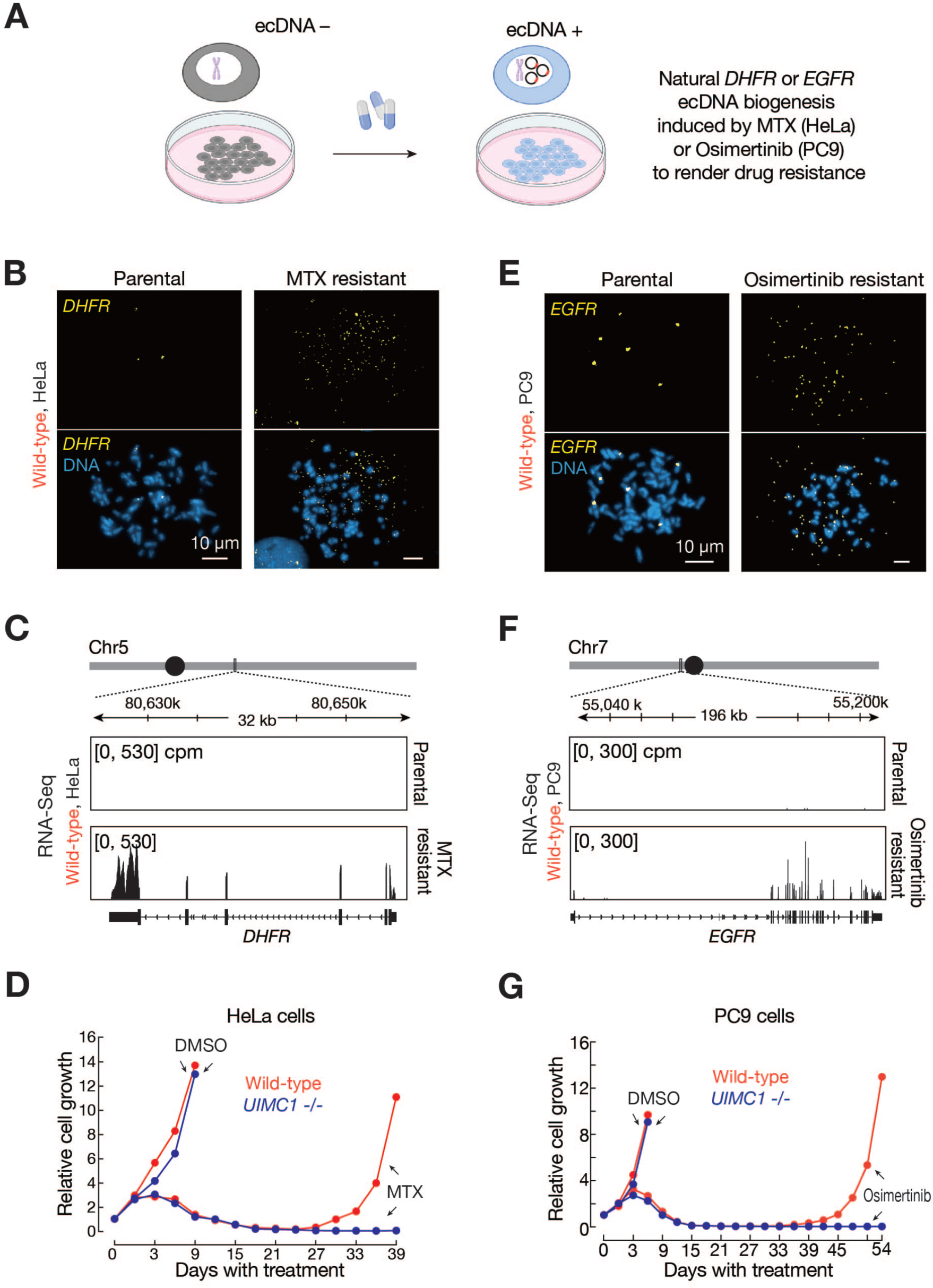
BRCA1-A complex drives ecDNA-mediated cancer cell evolution to acquire drug resistance. (A) Schematic design of using cancer treatment drugs to induce natural ecDNA production in cell culture. Methotrexate =MTX. (B) DNA-FISH to measure *DHFR* ecDNA in HeLa cells. Upon drug treatment, wild-type cells formed and accumulated ecDNA to acquire drug resistance. (C) RNA-Seq to show the expression of *DHFR* in parental and MTX resistant cells from the biological replicate #1. (replicate #2 and #3 were shown as Figure S7A and S7B). cpm = counts per million. (D) Cell growth curve upon MTX treatment. Mutating *UIMC1* abolishes HeLa cells evolving to MTX resistance. (E) DNA-FISH to measure *EGFR* ecDNA in PC9 cells. Upon Osimertinib treatment, wild-type cells formed and accumulated *EGFR* ecDNA to acquire drug resistance. (F) RNA-Seq to show the expression of *EGFR* in parental and Osimertinib resistant cells from the biological replicate #1. (replicate #2 was shown as Figure S7C and S7D). cpm = counts per million. (G) Cell growth curve upon Osimertinib treatment. Without *UIMC1*, PC9 cells failed to evolution resistance to the third generation EGFR inhibitor. Note, without CRISPR cut, it took one extra month for acquiring resistance, comparing with Figure 6C. See also Figure S7.

To examine whether BRCA1-A complex is broadly required for natural ecDNA production, we established PC9 lung cancer cells as a new system to study spontaneous *EGFR* ecDNA formation. After treating PC9 cells with EGFR inhibitor Osimertinib, we observed the production of natural *EGFR* ecDNA (Figure 7E), which drives a 12.9-fold increase of the expression of *EGFR* (33 fpkm in parental cells *vs* 424 fpkm in Osimertinib resistant cells, Figure 7F, S7C and S7D). Similar to HeLa cells, while wild-type cells evolved resistance to Osimertinib, mutating *UIMC1* resulted in the abrogation of adaptation (Figure 7G). Notably, we used similar approach to examine the function of *LIG4* and found that mutating it also led to the abolishment of ecDNA production and the incapability of cancer cells to evolve drug resistance in both HeLa-methotrexate and PC9-Osimertinib models (Figure S7E). In summary, our data provide strong evidence to suggest the essentiality of BRCA1-A complex and LIG4 for the process of ecDNA-mediated adaptation that allows cancer cells to acquire drug resistance.

## DISCUSSION

### DNA repair factors mediate ecDNA biogenesis

The biogenesis of ecDNA is a crucial biological process that occurs across various species. Previous studies have explored the involvement of specific DNA repair pathways in this process, with suggestive data leading to models proposing distinct roles for different repair mechanisms^3,4,25,31,54^. In this study, we performed a genome-wide CRISPR screening and identified LIG4 and BRCA1-A complexes play key roles in mediating ecDNA formation. While LIG4 complex serves as core components from NHEJ DNA repair, BRCA1-A acts at the upstream of DNA break signaling and is linked to determine the Homologous Recombination (HR) pathway choice^34–37^. Thus, our findings indicate that ecDNA biogenesis is not driven by a single DNA repair pathway. Instead, it is orchestrated by selective factors from various steps of the DNA repair process. Notably, in an ATM-deficient background, BRCA1-A and LIG4 have been reported to work synergistically to repair DNA breaks^55^. Our findings highlight that their cooperative relationship is also pivotal underlying the mechanism of ecDNA formation.

Upon DNA fragmentation, the upstream BRCA1-A complex protects the DNA ends from resection, likely funneling the un-resected ends towards downstream ligation to form a circle. Meanwhile, from the screen we uncovered MRN complex, which mediates end-resection, as a suppressor of ecDNA formation. Our findings thus highlight upon DNA fragmentation, the end-processing step is a highly dynamic process, which may ultimately determine the fate of ecDNA formation. At downstream, a few NHEJ core factors––LIG4, XRCC4, and DNA-PKcs––mediate ecDNA formation with LIG4 as the key enzyme to catalyze the process. However, the other NHEJ factors were not identified as essential for ecDNA biogenesis. Ku80 and Ku70 are essential for the viability of the human cells ^56,57^, confounding the efforts to discern them as ecDNA biogenesis factors. Additional NHEJ factors (ARTEMIS, PAXX, POLL, and POLM) are required when the ends possess non-complementary overhangs ^58^, and this could explain why they are dispensable for ecDNA biogenesis when blunt ends are generated during DNA fragmentation.

### Targeting ecDNA formation for cancer therapy

Given that ecDNA is frequently generated in cancer cells to drive tumor evolution and adaptation^3,7,8,11,16–18^, targeting ecDNA biogenesis presents a promising strategy for novel cancer therapies. Here, we report that both the BRCA1-A and LIG4 complexes play critical roles in ecDNA formation and contribute to drug resistance, highlighting new avenues for therapeutic intervention.

We showed that the LIG4 complex directly catalyzes ecDNA formation and thus represents ideal target. Humans appear to be able to tolerate the loss of LIG4 at a high level ^59^. Mutations in *LIG4* leads to LIG4 syndrome, a disease from which patients show developmental microcephaly and growth retardation but only have manageable immune deficiency in adult life ^59^. Additionally, our findings show that LIG4 is generally not essential for cell viability but becomes critical for cancer cells undergoing drug treatment, suggesting minimal and controllable toxicity from LIG4-targeted therapies. In fact, with the potential usage to sensitize cancer cells for radiotherapy, efforts were spent in the past to develop LIG4 drugs ^60–63^, but failed to reach a fruition. Uncovering its essential function in driving ecDNA biogenesis and cancer cell adaptation, our work calls for innovative approaches to develop LIG4 inhibitors for cancer therapy.

In parallel, the BRCA1-A complex, a key player in safeguarding ecDNA formation and mediating drug resistance, represents another therapeutic target. Notably, BRCA1-A shares certain components with the BRISC complex^33^, a deubiquitination complex involved in interferon responses, spindle assembly during mitosis, and hematopoiesis^64–66^. Therefore, targeting proteins that are exclusive to the BRCA1-A complex, such as ABRAXAS and RAP80, may help avoid broader side effects. Encouragingly, loss of either ABRAXAS or RAP80 in mice does not lead to developmental defects^67,68^, and our data indicate that these proteins are not normally required for cell viability but are essential when cancer cells adapt to drug treatment. Since these proteins lack enzymatic activities, developing compounds that block their interactions with partner proteins or induce their degradation is a promising path forward. In summary, by elucidating the essential roles of BRCA1-A and LIG4 complexes in ecDNA formation and drug resistance, our work underscores the potential for innovative, selective inhibitors of these complexes as cancer therapies.

### Limitations of the study

In this study, we identified the critical roles of BRCA1-A and LIG4 complexes in ecDNA formation and drug resistance. Although we provide compelling in vivo data for LIG4 in the context of ecDNA formation during *Drosophila* oogenesis, we are limited by the absence of murine models that would enable testing the function of these two complexes in vivo in a cancer setting. Additionally, alternative explanations exist for our data showing that LIG4 mutations led to HeLa cell death under methotrexate treatment. Methotrexate is a genotoxin capable of inducing chromothripsis^16,17^. Beyond its role in promoting DHFR ecDNA formation, LIG4 may also be required to repair genomic damage to support cancer cell survival under methotrexate stress. Lastly, while our findings advance understanding of ecDNA biogenesis in the context of genome fragmentation, other DNA repair factors and pathways can potentially contribute to ecDNA formation during development, disease, or in cases where the factors we identified are dysfunctional. Future studies are necessary to explore these additional players in ecDNA formation and their roles across various biological contexts.

## RESOURCE AVAILABILITY

### Lead contact

Further information and requests for resources and reagents should be directed to and will be fulfilled by the lead contact, Zhao Zhang.

### Materials availability

This study did not generate new unique reagents.

### Data and code availability

The sequencing data were deposited to the National Center for Biotechnology Information (NCBI) with accession number PRJNA1068592.

## ACKNOWLEDGMENTS

We thank Roberto Zoncu and Claire Goul for sharing scripts on CRISPR network analysis. We thank Charlie Gersbach and his lab for assisting with ddPCR. We thank David MacAlpine for suggestions on detecting chorion ecDNA. We thank members from ZZ lab for constructive suggestions during the project development and for reading the manuscript. This work was supported by the grants to Z.Z. from the Pew Biomedical Scholars Program, the Sontag Distinguished Scientist Award program, and the National Institutes of Health (R01 GM141018 and R01 GM152423), and to K.C.W. from the National Institutes of Health (R01CA263593), to Christian Cerda-Smith from the National Institutes of Health (F30CA247323), to Lee Zou from National Cancer Institute (R01 CA263394), and to D.A.R from the National Cancer Institute (R01 CA097096), and to M.K. from The Cooperative Trials Group for Neuro-Oncology.

## AUTHOR CONTRIBUTIONS

Z.Z., O.C., S.Y., and F.Y. conceived the project. L.Z. and D.A.R. provided critical comments on CRISPR screen data interpretation. All authors designed the experiments. M.K. sequenced glioblastoma DNA. F.Y. performed ecDNA versatile biosensor-related experiments. L.W. performed computational analysis. S.Y. performed cancer cell-related experiments. C.C.S., H.M.H., and K.C.W. provided guidance on CRISPR screen and performed related data analysis. O.C. performed all the rest of experiments. Z.Z. and O.C. wrote the manuscript. All authors read and approved the manuscript.

## DECLARATION OF INTERESTS

Z.Z, O.C., F.Y., and S.Y. are co-inventors on two US provisional patent applications filed by Duke University related to this work. Z.Z. is the co-founder of Muye Therapeutics.

---

**Figure S1.**
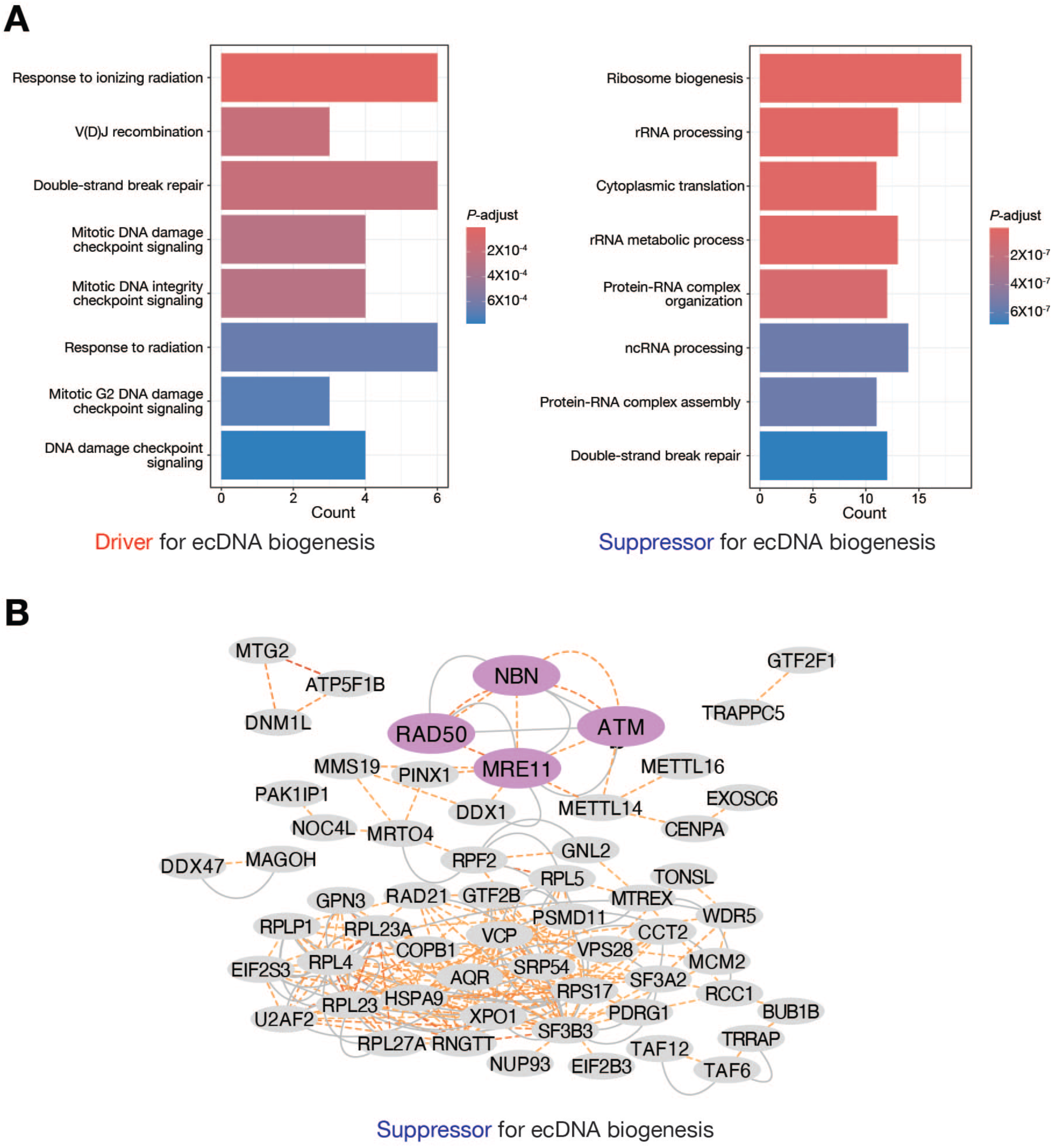
Bioinformatic analysis of CRISPR screen hits, related to Figure 1. (A) GO analysis to characterize the pathways that potentially regulate ecDNA biogenesis. (B) Interactome analysis of the factors that suppress ecDNA biogenesis. MRN complex components and ATM are highlighted by magenta. Factors that show co-essentiality are connected with dashed orange lines.

**Figure S2.**
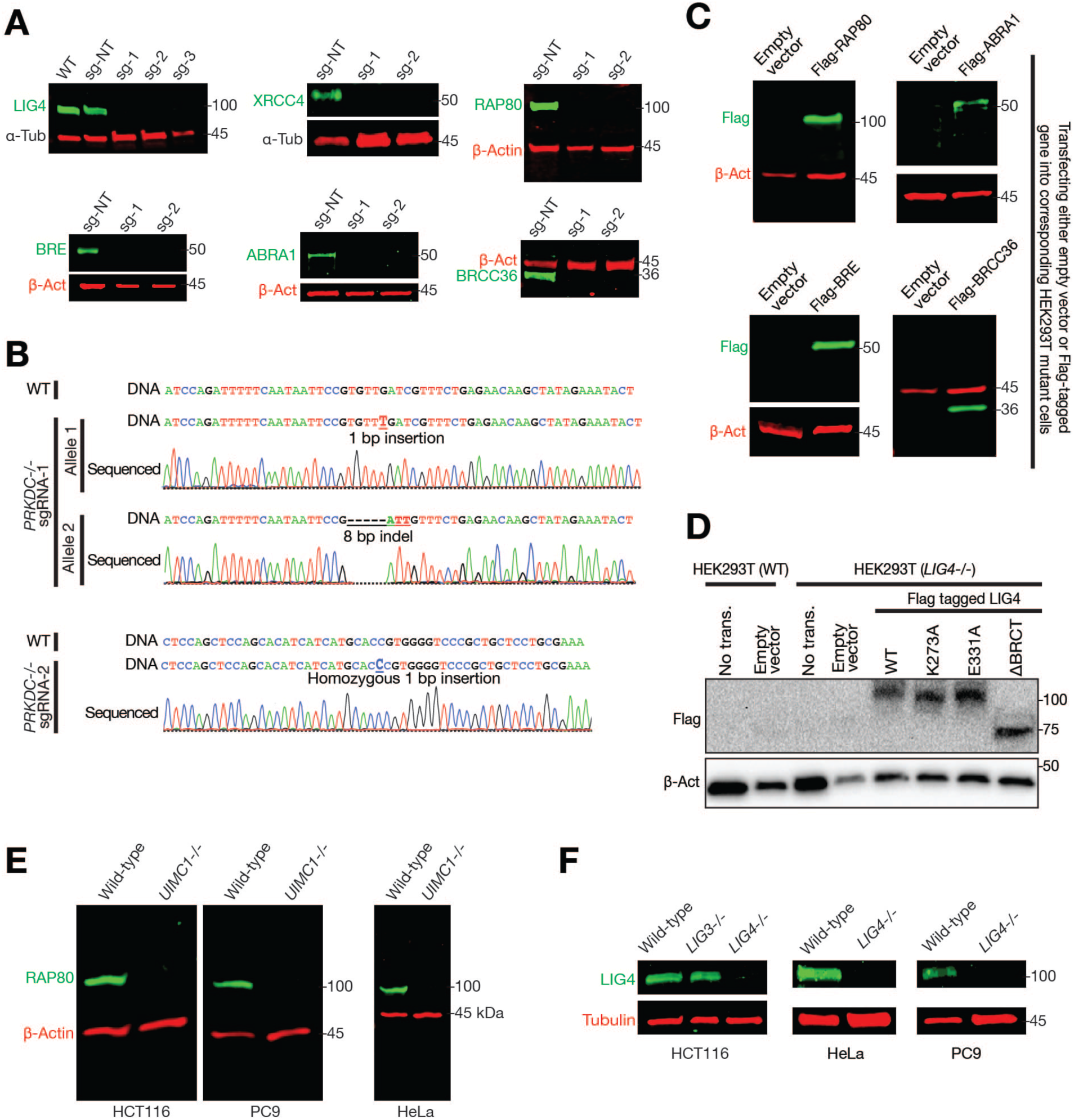
Validate mutation and rescue expression of genes from LIG4 and BRCA1-A complexes, related to Figure 2 and 3. (A) Western blotting to validate the mutation of *LIG4*, *XRCC4*, *UIMC1* (encodes RAP80 protein), *BABAM2* (encodes BRE protein), *ABRAXAS1* (encode ABRA1 protein), and *BRCC3* (encode BRCC36 protein) in HEK293T cells. (B) Sanger sequencing to validate *PRKDC* mutation in the HEK293T cells. The mutant allele generated by sgRNA-1 is trans-heterozygotes. The mutant allele generated by sgRNA-2 is a homozygote allele. The sequencing traces were manually divided to represent deletions. (C) Western blotting to examine the expression of BRCA1-A complex rescue constructs. Empty vector = transfecting the vector that does not harbor the rescue gene. (D) Western blotting to examine the expression of LIG4 rescue constructs. No trans. = no plasmid transfection. Empty vector = transfecting the vector that does not harbor LIG4 sequences. (E) Western blotting to validate the mutation of *UIMC1* in different cancer cell lines. (F) Western blotting to validate the mutation of *LIG4* in different cancer cell lines.

**Figure S3.**
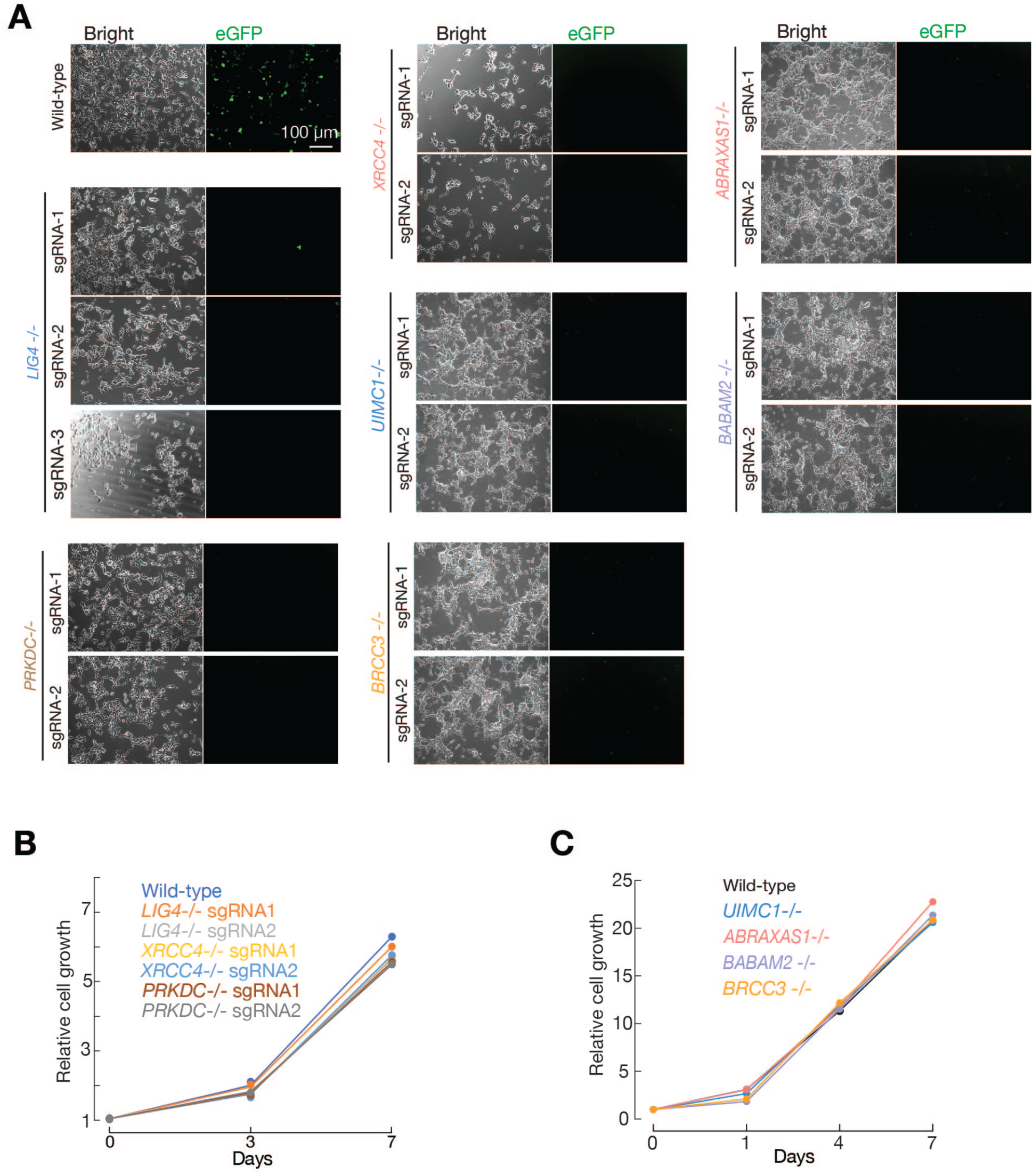
Mutating genes in LIG4 and BRCA1-A complexes abolishes ecDNA production from the pre-integrated reporter, related to Figure 2. (A) Individually mutating *LIG4*, *XRCC4*, *PRKDC*, *UIMC1*, *ABRAXAS1*, *BABAM2*, and *BRCC3* abolishes ecDNA production, as indicated by the loss of eGFP expression from the pre-integrated reporter. (B) Cell growth curve upon mutating *LIG4*, *XRCC4*, and *PRKDC* individually. (C) Cell growth curve upon mutating *UIMC1*, *ABRAXAS1*, *BABAM2*, and *BRCC3* individually. Mutants generated by sgRNA-1 for each gene were used for data display.

**Figure S4.**
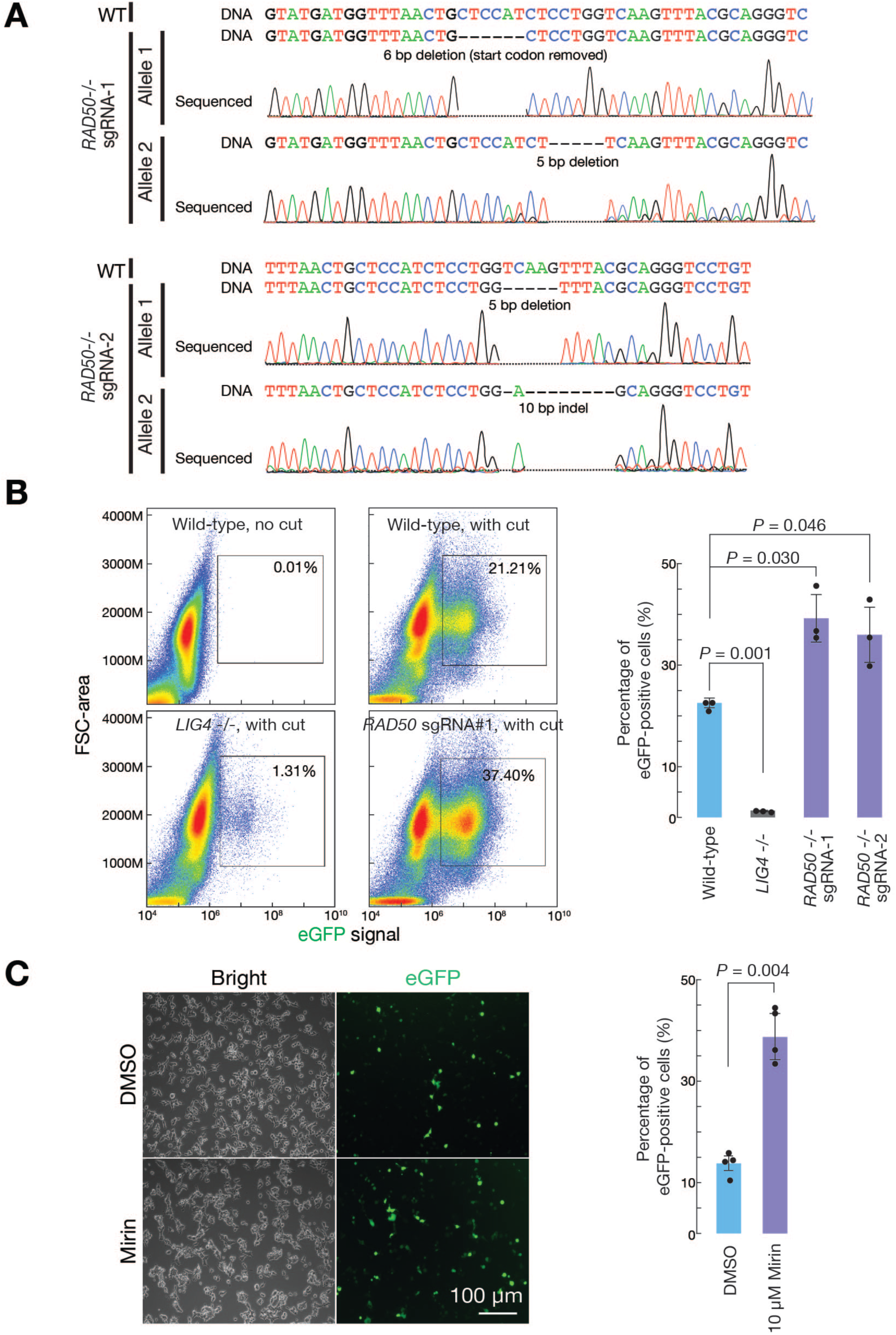
Suppressing the function of MRN complex enhances ecDNA production, related to Figure 2. (A) Sanger sequencing to validate *RAD50* mutation of the HEK293T cells. The sequencing traces were manually divided to represent deletions. (B) FACS to quantify ecDNA positive cells. The bar graph shows the quantification of three biological replicates. The bars report mean ± standard deviation from three biological replicates (n=3). *P*-values were calculated with a two-tailed, two-sample unequal variance *t* test. (C) Blocking the MRN complex function by using Mirin enhances ecDNA production. The bar graph shows the quantification of three biological replicates. The bars report mean ± standard deviation from three biological replicates (n=3). *P*-values were calculated with a two-tailed, two-sample unequal variance *t* test.

**Figure S5.**
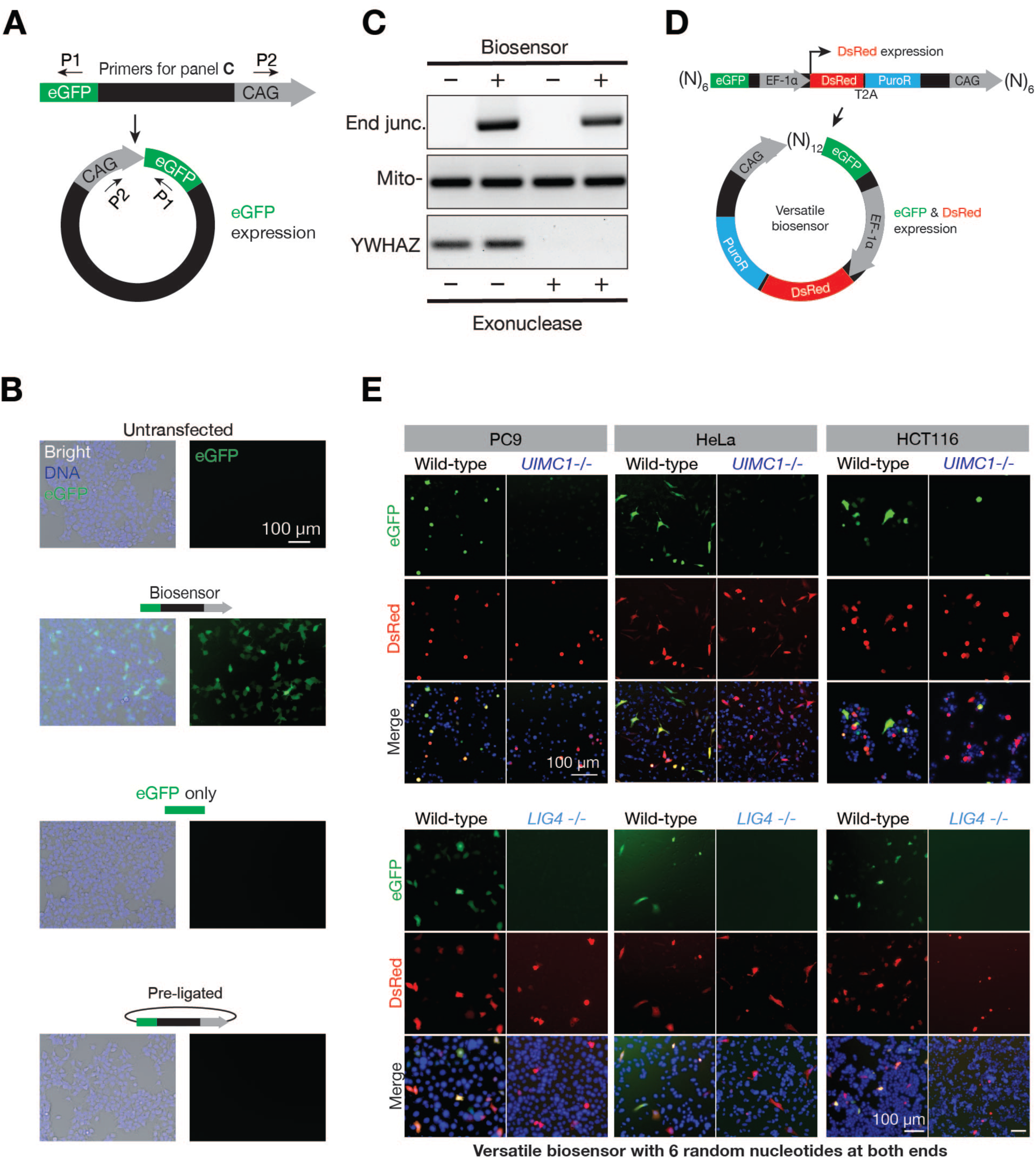
Versatile and robust biosensors to monitor ecDNA biogenesis, related to Figure 3. (A) Schematic design of the biosensor. Circularization brings the CAG promoter upstream of the eGFP sequences to initiate eGFP expression. (B) Testing the biosensor in HEK293T cells. Introducing the eGFP sequence only or pre-ligating the biosensor into a circle prevents eGFP expression, suggesting the eGFP expression from the biosensor relies on the circularization process, not random integration of the biosensor into the genome. (C) PCR assay to measure the production of ecDNA from the biosensor. Exonuclease treatment removes linear DNA, as evidenced by the signals from the YWHAZ gene. (D) Schematic design of the biosensor flanked by six random nucleotides. (E) Mutating *UIMC1* or *LIG4* abolishes ecDNA formation from the biosensor flanked by six random nucleotides.

**Figure S6.**
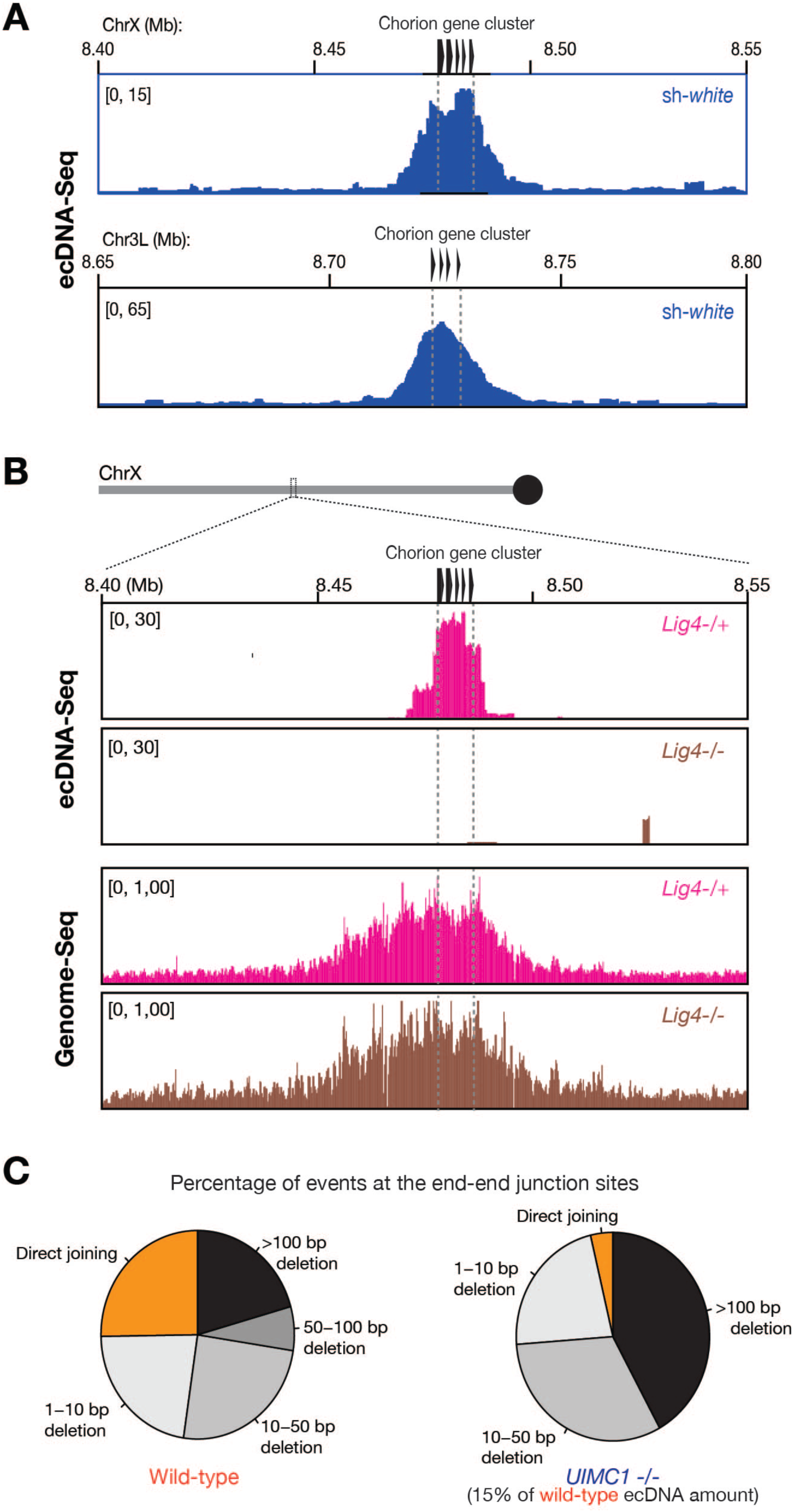
LIG4 drives natural ecDNA production during *Drosophila* oogenesis, related to Figure 4 and 6. (A) Mining the published ecDNA-Seq data from the *Drosophila* ovary shows ecDNA production from the two chorion loci. (B) ecDNA-Seq and Genome-Seq to measure the amplification of the chorion locus on the X chromosome and ecDNA production from this region. (C) Pie chart to show the percentage of sequencing reads harboring each event category (Related to Figure 6).

**Figure S7.**
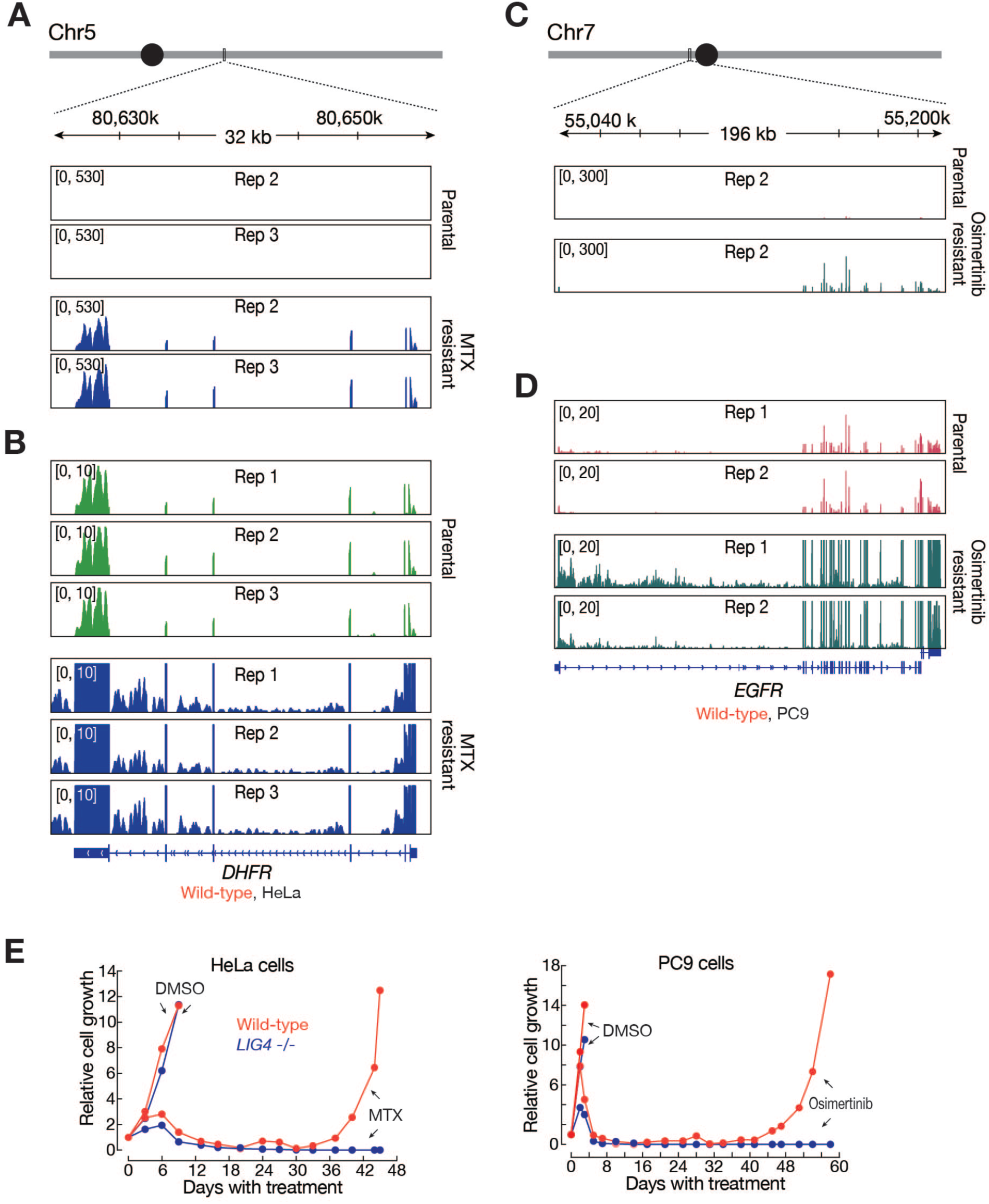
LIG4 drives ecDNA-mediated cancer cell evolution to acquire drug resistance, related to Figure 7. (A) Genome browser to show the expression of *DHFR* in parental and methotrexate (MTX) resistant cells from the biological replicate #2 and #3 (replicate #1 was shown as Figure 7C). (B) Genome browser to show the expression of *DHFR* in parental and methotrexate (MTX) resistant cells from three biological replicates. The y-axis was adjusted to display signals from parental cells. (C) Genome browser to show the expression of *EGFR* in parental and Osimertinib resistant cells from the biological replicate #2 (replicate #1 was shown as Figure 7F). (D) Genome browser to show the expression of *EGFR* in parental and Osimertinib resistant cells from two biological replicates. The y-axis was adjusted to display signals from parental cells. In all plots, the unit of y-axis is counts per million (cpm). (E) Cell growth curve upon Methotrexate or Osimertinib treatment. Without *LIG4*, HeLa cells failed to adapt to Methotrexate and PC9 cells failed to evolution resistance to the third generation EGFR inhibitor.

