## Supplementary material for "BRCA1-A and LIG4 complexes mediate ecDNA biogenesis and cancer drug resistance": Materials and Methods

#### STAR METHODS

#### KEY RESOURCE TABLE

| REAGENT or RESOURCE | SOURCE | IDENTIFIER |
|---|---|---|
| **Antibodies** | | |
| Rabbit polyclonal anti-XRCC4 | Proteintech | Cat# 21468-1-AP; RRID: AB_2878865_ |
| Mouse monoclonal anti-LIG4 | Proteintech | Cat# 66705-1-Ig RRID:AB_2882057 |
| Rabbit polyclonal anti-RAP80 | Fortis Life Sciences | Cat# A300-764A RRID:AB_2304339 |
| Rabbit polyclonal anti-ABRA1 | Fortis Life Sciences | Cat# A302-180A; RRID:AB_1659775 |
| Rabbit polyclonal anti-BRE | Cell Signaling Technologies | Cat# 12457S RRID:AB_2797924 |
| Rabbit polyclonal anti-BRCC36 | Cell Signaling Technologies | Cat# 18215S RRID:AB_2798795 |
| Mouse monoclonal anti-β-Actin | Proteintech | Cat# 66009-1-Ig; RRID:AB_2687938 |
| Rabbit monoclonal anti-tubulin-$\alpha$ | MilliporeSigma | Cat# SAB4500087 RRID:AB_10743646 |
| Mouse monoclonal anti-FLAG | MilliporeSigma | Cat# F3165 RRID:AB_259529 |
| Goat polyclonal anti-mouse IgG-HRP | Thermo Scientific | Cat# G-21040; RRID:AB_2536527 |
| Goat polyclonal anti-rabbit IgG-HRP | Thermo Scientific | Cat# G-21234; RRID:AB_1500696 |
| Goat anti-mouse IRDye680RD | Licor | Cat# 926-68070 RRID:AB_10956588 |
| Goat anti-rabbit IRDye800CW | Licor | Cat# 926-32211 RRID:AB_621843 |
| **Chemicals, peptides, and recombinant proteins** | | |
| Plasmid-safe DNase | Lucigen | Cat# E3110K |
| AMPure XP beads | Beckman Coulter | Cat# A63881 |
| Phi29 DNA Polymerase | New England Biolabs | Cat# M0269L |
| SpCas9 | Integrated DNA Technologies | Cat #1081059 |
| Electroporation Enhancer | Integrated DNA Technologies | Cat #1075916 |
| Exo Resistant Random Primer | Thermo Scientific | Cat# SO181 |
| Turbo DNase | Thermo Scientific | Cat# AM2239 |
| RNA FISH Wash Buffer A | LGC Biosearch Tech | Cat# SMF-WA1-60 |
| RNA FISH Hybridization Buffer | LGC Biosearch Tech | Cat# SMF- HB1-10 |
| RNA FISH Wash Buffer B | LGC Biosearch Tech | Cat# SMF- WB1-20 |
| Lipofectamine™ 3000 Transfection Reagent | Thermo Scientific | Cat# L3000001 |
| Methotrexate | MilliporeSigma | Cat# 454126-100MG |
| Osimertinib | Selleck Chemicals | Cat# S7297 |
| Mirin | Medchem Express | Cat# HY-117693 |
| 2X SSC buffer | Thermo Fisher Scientific | Cat #AM9763 |

| Critical commercial assays | | |
|---|---|---|
| Quick-gDNA MicroPrep kit | Zymo Research | Cat# D3021 |
| RNA Clean & Concentrator-5 | Zymo Research | Cat# R1016 |
| Ligation Sequencing Kit | Oxford Nanopore | Cat# SQK-LSK109 |
| Rapid Sequencing Kit | Oxford Nanopore | Cat# SQK-RAD004 |
| Magnetic mRNA Isolation Kit | New England Biolabs | Cat# S1550S |
| mirVana miRNA Isolation Kit | Thermo Fisher Scientific | Cat# AM1560 |
| SuperSignal West Pico PLUS Chemiluminescent Substrate Kit | Thermo Fisher Scientific | Cat# 34577 |
| Qubit dsDNA HS Assay kit | Thermo Fisher Scientific | Cat# Q33231 |
| Qubit dsDNA BR Assay kit | Thermo Fisher Scientific | Cat# Q32850 |
| Iscript cDNA Synthesis Kit | Bio-Rad | Cat# 1708890 |
| Ssofast EvaGreen | Bio-Rad | Cat# 1725204 |
| Vectashield Mounting Medium | VECTOR LABORATORIES INC MS | Cat# 101098-042 |
| CloneAMP HiFi PCR Premix | Takara Bio | Cat# 639398 |
| NEBNext Ultra II Q5 Master Mix | New England Biolabs | Cat# M0544L |
| Gotaq Green Master Mix | Promega | Cat# M7123 |
| 2x Phanta Max Master Mix | Vazyme | Cat# P515 |
| Short Read Eliminator XL kit | Circulomics | Cat# SS-100-111-01 |
| Gibco LV-MAX Lentiviral Production system | Thermo Fisher Scientific | Cat# A35684 |
| Quick-DNA Miniprep Kit | Zymo Research | Cat# D3025 |
| PureYield Plasmid Miniprep System | Promega | Cat# A1222 |
| CloneExpress Ultra One Step Cloning Kit | Cellagen Technology | Cat# C115-02 |
| Mix & Go Competent Cells-Zymo strain 10B | Zymo Research | Cat# T3020 |
| E.Z.N.A. Gel Extraction Kit | Omega Bio-Tek | Cat# D2500 |
| SE solution | Lonza | Cat# V4XC-1024 |
| DNeasy Blood & Tissue Kit | QIAGEN | Cat# 69594 |
| 2X TaqMan™ Universal PCR Master Mix | Thermo Fisher Scientific | Cat# 4304437 |
| iBlot™ 2 Transfer Stacks, PVDF | Thermo Fisher Scientific | Cat# IB24001 |
| Q5® Hot Start High-Fidelity polymerase reaction | New England Biolabs | Cat# M0494S |
| TruSeq Stranded mRNA and Total RNA kit | illumina | Cat# 20020594 |
| SuperScript IV | Invitrogen | Cat# 18090010 |
| Click-iT EdU Cell Proliferation Kit | Thermo Fisher Scientific | Cat# C10337 |
| Nextera XT DNA Library Preparation Kit | illumina | Cat# FC-131-1024 |
| **Deposited data** | | |
| Sequencing raw data for this paper | This paper | GEO: PRJNA794176 |
| **Experimental models: Cell lines** | | |
| HeLa cells | Duke Cell Culture Facility | Cat# CL-3272-FV |

| | | |
|---|---|---|
| HCT116 cells | Duke Cell Culture Facility | Cat# CL-2780-FV |
| PC9 cells | Duke Cell Culture Facility | Cat# CL-3238-FV |
| HEK293T cells | Duke Cell Culture Facility | Cat# CL-3395-FV |
| **Experimental models: Organisms/strains** | | |
| *D.melanogaster*: *DNA-lig4*[57] | This paper | Dr. Jefferey Sekelsky lab |
| *D.melanogaster*: $w^{1118}$ | Bloomington Drosophila Stock Center | Cat# 5905; RRID:BDSC_5905 |
| *H.sapien: HEK-293T biosensor cells* | This paper | Dr. Alun Luo lab |
| *H.sapien : patient derived glioblatoma and blood* | This paper | Dr. Mustafa Khasraw lab |
| **Oligonucleotides** | | |
| Custom CRISPR screen sequencing Forward primer: 5′-AATGATACGGCGACCACCGAGATCTACACTCCAAGGTCGGGCAGGAAGAGG | This paper | N/A |
| Custom CRISPR screen sequencing Reverse primer: 5′- CAAGCAGAAGACGGCATACGAGAT-(6 bp index sequence)-CCTCCCCTACCCGGTAGAATTGG | This paper | N/A |
| CRISPR/Cas9 sgRNA sequences (listed in Table S1) | Integrated DNA Technologies | N/A |
| LIG4-Wild-type primer: TACCGAGCTCGGATCCatggctgcctcacaaact | This paper | |
| LIG4-K273A primer: aaaccGCgctagatggtgaacgtatgc | This paper | |
| LIG4-ΔE331A primer: ccatcatcGcaccatcaagaatacagatt | This paper | |
| LIG4- BRCT primer: GATATCTGCAGAATTCTTATTTATCGTCATCATCTTTGTAGTCCTTGTCATCATCGTCCTTATAGTCCTTATCGTCGTCATCCTTGTAATCGttaacgttagtaaggttaggtgct | This paper | |
| MP-P2 indexing primers (listed in Table S1) | This paper | N/A |
| Primers for PCR and ddPCR (listed in Table S1) | Integrated DNA Technologies | N/A |
| Oligopain FISH probes (listed in Table S2) | This paper | N/A |
| **Recombinant DNA** | | |
| lentiCas9-Blast | addgene | Cat# 52962 |
| MinLibCas9 Library | addgene | Cat# 164896 |
| pU6-(BbsI)-CBh-Cas9-T2A-BFP plasmid | addgene | Cat# 64323 |
| pDR119 | addgene | Cat# 13332 |
| pcDNA3.1 | Thermo Fisher Scientific | Cat# V79020 |
| pOZ-N-FH RAP80 | addgene | Cat#27501 |
| pOZ-N-FH Abraxas | addgene | Cat# 27495 |
| pOZ-N-FH BRCC36 | addgene | Cat# 27496 |
| GenEZ$^{tm}$ORF clone BABAM2 | Genscript | Cat# OHU14947D |
| pCAG-GFP | addgene | Cat# 11150 |
| **Software and algorithms** | | |

| | | |
|---|---|---|
| GraphPad Prism 8 | N/A | https://www.graphpad.com/scientific-software/prism/ |
| MinKNOW version 21.05.25 | N/A | Oxford Nanopore Technologies |
| Porechop 0.2.4 | N/A | https://github.com/rrwick/Porechop |
| Minimap2 | N/A | https://github.com/lh3/minimap2 |
| Samtools 1.12 | N/A | http://www.htslib.org/ |
| IGV | N/A | https://igv.org/doc/desktop/ |
| R studio | N/A | https://posit.co/download/rstudio-desktop/ |
| MAGeCK | N/A | https://sourceforge.net/p/mageck/ |
| FIREWORKS | N/A | https://github.com/mendillolab/fireworks/ |
| Cytoscape 3.10.1 | N/A | https://cytoscape.org/ |
| CRISPResso2 | N/A | https://github.com/lucapinello/CRISPResso |
| StringTie v2.2.3 | N/A | https://github.com/gpertea/stringtie |
| AmpliconArchitect 1.3.r2 | N/A | https://github.com/virajbdeshpande/AmpliconArchitect |
| BEDTools (v2.26.0) | N/A | https://github.com/arq5x/bedtools2 |
| **Other** | | |
| DMEM | Thermo Fisher Scientific | Cat# 10569044 |
| RPMI 1640 | Millipore Sigma | Cat# R8758-500ML |
| Fetal Bovine Serum | Cytivia | Cat# SH30396.03 |
| 100X Penicillan-Strptomycin | Thermo Fisher Scientific | Cat# 15140122 |
| Opti-MEM reduced serum medium | Thermo Fisher Scientific | Cat# 31985062 |
| SPRIselect beads | Beckman Coulter | Cat# B23317 |
| RIPA buffer | Thermo Fisher Scientific | Cat# 89900 |
| 1x Complete protease inhibitor | Roche | Cat #4693159001 |
| KaryoMAX™ Colcemid™ Solution | Thermo Fisher Scientific | Cat# 15212012 |
| Vectashield Antifade Mounting Media | Vector Laboratories | Cat# H-1200-10 |
| AMPure XP beads | Beckman Coulter | Cat# A63881 |

### EXPERIMENTAL MODEL AND SUBJECT DETAILS

#### Mammalian cells and related drug treatment

HEK293T, HeLa, and HCT116 cells were cultured under the following conditions: Gibco DMEM (High Glucose, GlutaMAX Supplement, Pyruvate; Thermo Fisher Scientific, Cat #10569044), supplemented with 10% FBS (Cytvia, Cat #SH30396.03) and 1% penicillin–streptomycin (Thermo Fisher Scientific, Cat #15140122). PC9 cells were cultured in RPMI 1640 Medium (Sterile, pH 7.0 to 7.6, with L-glutamine and sodium bicarbonate, suitable for cell culture; Sigma-Aldrich, Cat # R8758-500ML), supplemented similarly with 10% FBS (Cytvia, Cat #SH30396.03) and 1% penicillin–streptomycin (Thermo Fisher Scientific, Cat #15140122). HEK293T biosensor cells were a generous gift from Dr. Alun Luo[1] and were re-engineered with lentivirus containing vector lentiCas9-Blast (Addgene #52962) to express multiple copies of Cas9 protein. These cells were cultured in similar conditions as normal HEK293T cells. The incubation conditions were maintained at 37°C and 5% $CO_2$.

To induce natural ecDNA formation in HeLa or PC9 cells: $8x10^5$ of HeLa cells or $5x10^5$ of PC9 cells were seeded into 10 cm dishes. After 24 hours of recovering, HeLa cells were subjected to treatment with 100 nM Methotrexate (MilliporeSigma, Cat#454126-100MG) for up to one and half months. PC9 cells were treated with 20 nM Osimertinib (Selleck Chemicals, Cat #S7297) for up to two months. Culture medium supplemented with Methotrexate or Osimertinib were replaced twice per week. Cell numbers were monitored with a microscope when replacing the medium.

PC9 cells (either wild-type or *UIMC1-/-)* were also subjected to CRISPR-C to induce EGFR ecDNA formation and incubated for seventy-two hours after. Cells were then counted and $5x10^5$ cells were seeded into 10 cm dishes. Cells were allowed to recover for 24 hours, then treated with 20 nM of Osimertinib (Selleck Chemicals, Cat #S7297) for one-to-two months. Medium (with 20 nM Osimertinib) was replaced every three days, meanwhile the live cells were counted under microscope.

To validate MRN function by small molecule inhibition, wild-type HEK293T cells were plated at 300,000 cells in 6-well with 2 ml of media containing 10 μM Mirin (Medchem Express, Cat #HY-117693) and incubated overnight. The next day, CRISPR-C left and right sgRNA lentivirus was added with 8 μg/ml polybrene to the cells and allowed to grow for 72 hours. Cells were observed for eGFP with Zeiss Axio Observer microscope.

**Fly strains, housing, and husbandry conditions**

All flies were grown on standard agar-corn medium and maintained at 25°C. Flies carrying the *Lig4[57]* mutation were generously gifted by Dr. Jefferey Sekelsky. Female flies aged 3-6 days were selected for experiments. Ovaries were carefully dissected and washed in 1x Phosphate-buffered saline (PBS) prior to experiments.

#### METHOD DETAILS

**Lentivirus production and genome-wide CRISPR screen**

Transfection reagents were prepared in Opti-MEM reduced serum medium (Thermo Fisher Scientific, Cat#31985062) with appropriate scaling for culture surface area, according to manufacturer instructions. For the lentiviral preparation of the genome-wide sgRNA lentiviral library, 80% confluent HEK293T cells were transfected in a T-225 Flask with 42.4 $\mu$g of the MinLib plasmid library (MinLibCas9 Library was a gift from Dr. Mathew Garnett, Addgene #164896), 32.4 $\mu$g of psPAX2, and 21.2 $\mu$g of pMD2.G, using 374 $\mu$l of Lipofectamine 2000 supplemented with 17.54 $\mu$l of PLUS Reagent. After 6 hours following lipofection, the media was replaced with fresh pre-warmed harvest media (HEK293T media supplemented with 25% FBS). After 48 hours, the viral supernatant was collected, filtered using a 0.45 $\mu$m PES filter, and either used directly or stored at -80°C for future use. The titer of the lentiviral sgRNA library was determined by flow cytometry. Large batch of lentivirus containing biosensor CRISPR-C cutters—left and right sgRNA––vectors were prepared using the Gibco LV-MAX Lentiviral Production system (Thermo Fisher Scientific Cat # A35684) as per manufacturer instructions. Equal portions of left cut and right cut virus were pooled together to make CRISPR-C biosensor cutting virus.

For genome-wide CRISPR screen, ~$1.0x10^8$ HEK293T biosensor cells were seeded into 15 cm tissue culture dishes with 8 μg/ml polybrene and transduced with the titered lentiviral sgRNA library at a low multiplicity of infection ~0.4 (statistically ensuring that most cells harbor no more than 1 sgRNA) to achieve greater than 1,000x coverage of the sgRNA library, in biologic triplicate. Twenty-four hours post-transduction, media was replaced with fresh media and cells were incubated for 72 hours. Cells were then selected for BFP expression by Fluorescent Activated Cell sorting (FACS). Sorted cells containing sgRNA were allowed to recover for 24 hours in fresh media with 30% FBS. Cells were then infected with optimal CRISPR-C biosensor cutting virus with 8 μg/ml polybrene for 24 hours to allow for excision of the DNA biosensor. Virus media was replaced with fresh media and cells were passaged and maintained above 1000x coverage for 72 hours post-transduction. Replicates were harvested and cells were then sorted by FACS based on GFP expression; the cells were collected in FBS-coated tubes, maintaining approximately 1000× coverage per population. An ungated control sample was also collected for each replicate.

Immediately following collection, replicate samples were individually subjected to genomic DNA extraction using the Quick-DNA Miniprep Kit (Zymo Research, Cat #D3025). sgRNA libraries were recovered from gDNA via PCR amplification using NEBNext Ultra II Q5 Master Mix (NEB, Cat #M0544L) according to manufacturer instructions and using customized primers:

FWD 5′-
AATGATACGGCGACCACCGAGATCTACACTCCAAGGTCGGCAGGAAGAGG
REV 5′-
CAAGCAGAAGACGGCATACGAGAT-(6 bp index sequence)-
CCTCCCCTACCCGGTAGAATTGG

Amplified libraries were purified using SPRIselect beads (Beckman Coulter, Cat #B23317) employing right-sided selection of 0.8x then to 1.2x the original volume. Each DNA sample was quantified using the Qubit dsDNA Broad Range Assay Kit (Thermo Fisher Scientific, Cat #Q32850) and quality-checked using an Agilent 4150 TapeStation System. DNA was analyzed using D5000 ScreenTape (Agilent, Cat #5067-5588). Samples were pooled and sequenced on a NextSeq 500 (Illumina) with 20 bp single-end sequencing using customized read and index primers.

Custom Read 5′-
CTTGGCTTTATATATCTTGTGGAAAGGACGAAACACCG
Custom Index 5′-
GGATCCAATTCTACCGGGTAGGGGAGG

**Gene mutation by CRISPR-Cas9**

The sgRNAs were derived from the MinLib CRISPR guide RNA library. To ensure efficiency, an additional guanine was appended to the sgRNAs that do not start with a guanine. Each sgRNA was cloned into pU6-(BbsI)-CBh-Cas9-T2A-BFP plasmid (Addgene, Cat #64323) and verified by Sanger sequencing. To generate gene mutations, cells were seeded at 300,000 cells per well in 6-well plates with complete media and allowed to grow overnight. The next day, 2 ml of culture media was replaced prior to transfection. Transfection was carried out by delivering 3 μg of plasmid to cells using

Lipofectamine™ 3000 Transfection Reagent (Thermo Fisher Scientific, Cat #L3000001) according to manufacturer's recommended protocol. Cells were incubated with transfection mixture for 24 hours and then incubated with fresh media for an additional 48 hours. Then cells were sorted for BFP signal using a Beckman Coulter Astrios EQ High-Speed Cell Sorter. The collected cells were plated into 6-cm dishes and allowed to recover for 48 hours post-sorting. Next, the cells were dissociated and diluted to 30 cells/ml. One hundred μl of the cell suspension was distributed into 96-well plates per well. Single clones were allowed to grow into stable colonies over a period of approximately 10-14 days for HEK293T cells and 30-40 days for HeLa, HCT116, and PC9 cells. Finally, the colonies were validated for mutation by Sanger sequencing the site of sgRNA directed mutation or protein depletion by western blot. sgRNA sequences are listed in Table S1.

**Cloning and expression of LIG4 wild-type and mutants for rescue**

The cloning of the *LIG4* wild-type and mutant alleles were amplified by using CloneAMP HiFi PCR Premix (Takara, Cat # 639298) from the pDR119 plasmid (Addgene, Cat# 13332). The specific primers used for this purpose are listed in Table S1. Following amplification, the PCR products were assembled into a pcDNA3.1 plasmid (Thermo Fisher Scientific, Cat # V79020), which was pre-digested with EcoRI (NEB, Cat # R3101S) and BamHI (NEB, Cat # R3136S). This assembly was facilitated by the CloneExpress Ultra One Step Cloning Kit (Cellagen Technology, Cat# C115-02). For plasmid propagation, Mix & Go Competent Cells-Zymo strain 10B (Zymo Research, Cat# T3020), a chemically competent E. coli strain, was utilized. The propagated plasmids were then extracted using the PureYield Plasmid Miniprep System (Promega, Cat# A1222). To confirm the accuracy of the constructed vectors, Sanger sequencing was employed for verification.

HEK293T biosensor cells (WT and *LIG4*-/-) were seeded at 300,000 cells/well in 6-well plates with 2 ml of culture media and incubated overnight. The following day, plasmids were transfected into cells using Lipofectamine 3000 Transfection Reagent (Thermo Scientific, Cat # L3000008) as per manufacturer's instruction. The culture medium was refreshed 16 hours post-transfection. Images were acquired at 48 hours post-transfection using a Zeiss AxioObserver microscope. For western blotting, cells were harvested at 48 hours after transfection.

**Transfection of wild-type BRCA1-A factors in mutants for rescue**

pOZ-N-FH RAP80 (Addgene, Cat#27501), pOZ-N-FH Abraxas (Addgene, Cat# 27495), and pOZ-N-FH BRCC36 (Addgene, Cat# 27496) expression plasmids were purchased from Addgene for transient expression of human wild-type RAP80, ABRAXAS, and BRCC36 protein, respectively. GenEZ$^{tm}$ORF clone BABAM2 (Genscript, Cat# OHU14947D) expression plasmid was purchased from Genscript for transient expression of human wild-type BRE protein. For plasmid propagation, Mix & Go Competent Cells-Zymo strain 10B (Zymo Research, Cat #T3020), a chemically competent E. coli strain, was utilized. The propagated plasmids were then extracted using the PureYield Plasmid Miniprep System (Promega, Cat #A1222). Sanger sequencing was employed to confirm the accuracy of the vectors.

HEK293T biosensor cells with mutations on BRCA1-A complex genes were seeded at 300,000 cells/well in 6-well plates with 2 ml of culture media and incubated overnight. The following day, plasmids were transfected into cells using Lipofectamine 3000 Transfection Reagent (Thermo Fisher Scientific, Cat #L3000008) according to manufacturer's instruction. The culture media was refreshed 16 hours post-transfection and replaced with CRISPR-C left and right sgRNA lentivirus infused media. Images were acquired at 48 hours post-transfection using a Zeiss Axio Observer microscope. For western blotting, cells were harvested 48 hours after transfection.

**Design, cloning, and transfection of versatile biosensors**

To construct the versatile biosensor, the pCAG-GFP plasmid (Addgene, Cat #11150) was first digested with EcoRI (NEB, Cat #R3101S) and HindIII (NEB, Cat #R3104S), followed by re-ligation. GFP-polyA was then inserted into the NotI (NEB, Cat #R3189S) site, this construct (Version 1) was used to test the concept of biosensor (Extended Data Fig. 7). To build the final biosensor, DsRed-T2A-PuroR fragment was amplified from a U6-sgRNA-DsRed-2A-PuroR vector using CloneAMP HiFi PCR Premix (Takara, Cat #639298), and subsequently integrated into Version 1 biosensor that is pre-digested with SpeI (NEB, Cat #R3133S). The biosensor plasmid was validated via Sanger sequencing. For linearization to get the biosensor, 10 $\mu$g of plasmids (50 $\mu$l volume) was treated with 1 $\mu$l EcoRV (NEB, Cat # R3195T) and 5 $\mu$l CutSmart buffer, incubated at 37 ºC for 4 hours. The products were then purified by using 0.8% agarose gel. Bands of expected size were extracted using E.Z.N.A. Gel Extraction Kit (Omega Bio-Tek, Cat #D2500). HEK293T, HeLa, PC9, and HCT116 cells ($1x10^6$ cells each) were transfected with 500 ng (HEK293T) or 2000 ng (HeLa, HCT116, and PC9) of linearized biosensor using Lipofectamine 3000 Transfection Reagent (Thermo Fisher Scientific, Cat #L3000008). The culture media was refreshed 4 hours post-transfection. Images were acquired at 24 hours post-transfection using a Zeiss Axio Observer microscope. Image assembly and processing were conducted using Adobe Photoshop and Illustrator. To prepare Versatile biosensors with random nucleotides at the two ends, 1 ng of the biosensor was amplified with primers harboring six random nucleotides by using CloneAMP HiFi PCR Premix (Takara, Cat #639298). The amplification conditions were: initial denaturation at 95 ºC for 3 min; 30 cycles of 95 ºC for 15 s, 64 ºC for 15 s, and 72 ºC for 2 min; final extension at 72 ºC for 5 min (Primers listed in Table S1). The PCR products underwent electrophoresis and subsequent gel purification as described above.

**CRISPR-C mediated ecDNA induction by nucleofection**

sgRNAs to induce *EGFR*-containing ecDNA were designed using CHOPCHOP (Version 3) webtool, which generated a 1.8 Mb size ecDNA. All the sgRNAs were purchased from Synthego (https://www.synthego.com/). The sequences of the sgRNAs were listed in Table S1.

To generate ecDNA, $5x10^5$ PC9 cells were collected and centrifuged at 150 g for 10 minutes. The Cas9-sgRNA Ribonucleoprotein (RNP) complexes were introduced into cells by electroporation with Lonza 4D-Nucleofector™ system (X Unit). Briefly, the RNP complexes were assembled by mixing 2 µl of SpCas9 (IDT, Cat #1081059, 61 µM), 1 µl of left-sgRNA and 1 µl of right-sgRNA (100 µM, in TE buffer, pH 8.0). The RNP mixtures were incubated at room temperature for 15 minutes. Then, 4 µl of RNP, 5 µl of electroporation enhancer (IDT, Cat #1075916, 100 µM), 91 µl of cells (in SE solution, Lonza, Cat #V4XC-1024) were mixed, transferred to the Nucleocuvette (Lonza, Cat #V4XC-1024) and electroporated with Lonza 4D-Nucleofector™ system with the preset code: CN114. After electroporation, cells were transferred to 24-well plates with pre-warmed culture medium. For ecDNA junction detection by ddPCR, cells were collected at 12 hours post nucleofection.

**TaqMan Real-Time PCR**

Cells were washed with 2 ml PBS, followed by adding 100 µl 0.25% Trypsin+EDTA and incubated in 37 °C for 5-10 minutes. Trypsin was quenched by adding 1 ml of culture media (10% FBS). Cells were pelleted at 300 g for 5 minutes. The supernatant was aspirated, and the cell pellets were stored at -80 °C.

Genomic DNA was isolated using DNeasy Blood & Tissue Kit (QIAGEN, Cat #69594) according to the manufacturer's instructions. Briefly, the cell pellets were resuspended in 200 µl PBS, and then mixed with 200 µl lysis buffer with 20 µl of 20mg/mL Proteinase K solution. The protein was digested at 56 °C for 10 min. The cell lysate was transferred to the binding columns and washed 2 times. Genomic DNA (gDNA) was eluted with 80 µl RNase free water.

ecDNA abundance was analyzed by Taqman Real-Time PCR assay according to the manufacturer's supplied protocols (Thermo Fisher Scientific). Briefly, approximately 100 ng of gDNA was used in a 20 µl reaction with 10 µl 2X TaqMan™ Universal PCR Master Mix (Thermo Fisher Scientific, Cat #4304437), 900 nM of primers (225 nM for each primer), 125 nM of FAM probe (ecDNA), and 125 nM of HEX probe (GAPDH). TaqMan PCR was conducted with the CFX96 Real-Time PCR System (Bio-Rad Laboratories), with the following setting: 50 °C for 2 minutes, 95 °C for 10 minutes, 45 cycles (95 °C for 15 seconds, 60 °C for 60 seconds). The amount of ecDNA and GAPDH (serving as reference) were quantified by the comparative CT method[2]. All results were normalized to GAPDH; each sample was repeated three times, and the average value was used. Primers and probes were the same ones used in ddPCR assay listed in Table S1.

**Quantitative droplet digital PCR (ddPCR) of CRISPR-C junction sites**

Amplicons for the circular DNA junctions and glyceraldehyde 3-phosphate dehydrogenase (GAPDH) were designed using the IDT PrimerQuest™ Tool (https://www.idtdna.com/pages/tools/primerquest). Dual-quenched probes (IDT) were used. Probes for ecDNA junction amplicons were labeled with FAM, and probe for

GAPDH amplicon was labeled with HEX to facilitate the multiplexing. The sequences of probes and primers are listed in Table S1.

ddPCR was performed on samples using the QX200TM ddPCR system (Bio-Rad Laboratories). The amplification reactions were set up according to the manufacturer's specifications (Bio-Rad Laboratories). Briefly, approximately 50 ng of gDNA was used in a 20 μl reaction with 10 μl ddPCR Supermix for probes (no dUTP) (Bio-Rad Laboratories), 900 nM for primers (225 nM for each primer), 125 nM of FAM probe, and 125 nM of HEX probe. Then, the droplets were created using droplet-generating oil for probes, DG8 cartridges, DG8 gaskets and the QX200 Droplet generator (Bio-Rad Laboratories). The 96-well plate with droplets was sealed with a PX1 PCR plate sealer (Bio-Rad Laboratories). PCR of droplets was performed using the following setting: 95 °C for 10 minutes, 40 cycles (95 °C for 30 seconds, 55.8 °C for 60 seconds), 98 °C for 10 minutes. After PCR amplification, the plate was analyzed with a QX200 Droplet Reader (Bio-Rad Laboratories). The positive and negative droplets were quantified by QuantaSoft Software. The ecDNA frequency was calculated as:

$$ecDNA\ frequency\ (\%) = \frac{ecDNA\ concentration}{GAPDH\ concentration}$$

**Western blotting**

Cells were lysed in RIPA buffer (Thermo Fisher Scientific, Cat #89900) with 1x complete protease inhibitor cocktail (Roche, Cat #4693159001). The lysate was resolved by SDS-PAGE gels, transferred to PVDF membranes (Thermo Fisher Scientific, Cat #IB24001) with an iBlot2, and analyzed by immunoblotting with indicated primary antibodies. The following primary antibodies were used in overnight incubation at 4 °C: anti-ABRA1 (Fortis Life Sciences, Cat #A300-764A; 1:1000), anti-RAP80 (Fortis Life Sciences, Cat

#A300-764A; 1:500), anti-BRE (Cell Signalling Technology, Cat #12457S; 1:1000), anti-BRCC36 (Cell Signalling Technology, Cat #18215S; 1:1000), anti-$\alpha$-Tubulin (MilliporeSigma, Cat # SAB4500087; 1:10000), anti-β-Actin (Proteintech, Cat #66009-1-Ig; 1:10000), and anti-FLAG (MilliporeSigma, Cat#F3165; 1:1000). Secondary antibodies include: anti-mouse and anti-rabbit IgG-HRP (Thermo Fisher Scientific, Cat #G-21040 and #G-21234; 1:5000) or anti-mouse IRDye680RD (Licor, Cat #926-68070; 1:10000) and anti-rabbit IRDye800CW (Licor, Cat #926-32211; 1:10000). Protein bound membranes were washed with 1x TBST (Tris-Buffered Saline, 0.1% Tween 20) and incubated with IRDye secondary antibodies for 1 hour at room temperature, followed by 1x TBST washes. Membranes were imaged with a Licor Odyssey CLx.

**Metaphase DNA-FISH**

Cells were arrested in metaphase with KaryoMAX™ Colcemid™ Solution (Thermo Fisher Scientific, Cat# 15212012) treatment at 0.2 μg/ml for 6 hours (HeLa), or 16 hours (PC9). Cells were washed with 1X PBS, trypsinized, and resuspended in 75 mM KCl for 15 min. Then the cells were fixed with fresh ice-cold Carnoy's fixative (3:1 methanol: glacial acetic acid, v/v) for 10 minutes, followed by three additional washes with the fixative. Finally, the cells suspended in the fixative were dropped onto humidified glass slides. The slides were air-dried and aged at room temperature overnight. The slides with fixed cells were briefly equilibrated in 2X SSC buffer (Thermo Fisher Scientific, Cat #AM9763), followed by dehydration in ascending ethanol series (70%, 85%, 100%) for 2 minutes each. Then the slides were air dried completely. Oligopaint FISH probes with hybridization buffer were mixed well and applied onto the slides, covered by coverslips, and sealed with rubber cement. Then the probes and samples were co-denatured at 75°C for 5 minutes, and hybridization was carried out at 37°C overnight. The coverslips were removed, and the slides were washed in 0.4× SSC with 0.3% IGEPAL (72°C), and 2× SSC with 0.1% IGEPAL, for 5 minutes each. DNA was counter-stained with DAPI, and the slides were mounted with VECTASHIELD® Antifade Mounting Medium with DAPI (Vector Laboratories, Cat #H-1200-10). FISH images were acquired by Zeiss Axio Observer microscope using a 40X objective.

The custom Oligopaint probes used in this study were prepared based on the method developed from the laboratory of C.T. Wu[3]. Briefly, the oligo library was designed from the algorithm developed from the Wu lab. Each individual oligo consisted of a unique set of primer pairs for PCR amplification, and a T7 promoter sequence was attached in the forward primer to enable in vitro transcription. The Oligopaint-covered genomic regions used in this study were as follows: DHFR (chr5: 80,626,226-80,654,983), EGFR (chr7:55,016,118-55,213,816). The oligo pools for each ecDNA were synthesized by Twist Bioscience. All the sequences of the oligos are listed in Table S2. The oligos were amplified by PCR, followed by in vitro transcription, and then the RNA was converted to ssDNA by reverse transcription, in which the fluorophores were introduced to the probes.

**Library preparation for targeted deep sequencing of ecDNA junctions**

Genomic DNA from the cells subjected to CRISPR-C experiments was isolated using DNeasy Blood & Tissue Kit (QIAGEN, Cat #69594) according to the manufacturer's instructions. Then two-step PCR was performed to amplify the ecDNA junctions and barcode the libraries. For WT samples, 35 cycles of primary PCR were performed using ~100 ng of genomic DNA in a 25 µL Q5® Hot Start High-Fidelity polymerase reaction (NEB, Cat #M0494S). For UIMC1 mutants, multiple PCR reactions using ~500 ng of genomic DNA in total were set-up in order to acquire enough amplicon quantities comparable to the wild-type. Then 20 μl AMPure XP beads (Beckman Coulter, Cat #A63881) were used to purify DNA. The concentrations of the purified DNA were measured by Nanodrop. Illumina adapters and indexing sequences were added via 10 cycles of secondary PCR with 30 ng of primary PCR product in a 25 μL Q5® Hot Start High-Fidelity polymerase reaction. The final PCR products were purified by 20 μl AMPure XP beads. The concentrations of the purified DNA were measured by Qubit dsDNA HS Assay kit (Thermo Fisher Scientific, Cat #Q33231). Then, amplicons were sequenced by Next Generation Sequencing (Amplicon-EZ, 150-500 bp, Azenta). The sequences of primers are listed in Table S1.

**RNA Extraction and RNA sequencing**

Three million HeLa and PC9 cells were plated in 10 cm dishes with complete media and allowed to incubate at 37°C overnight. ecDNA positive HeLa and PC9 cells were plated with 1 μM Methotrexate and 20 nM Osimertinib infused media, respectively. Cells were washed with 1X PBS and detached with 0.25% Trypsin+EDTA and incubated in 37°C for 5-10 minutes. Trypsin was quenched by adding 1 ml of culture media (10% FBS). Cells were pelleted at 150 x g for 5 mins and then placed on ice. 300 μl of Lysis/Binding buffer from mirVana miRNA Isolation Kit (Invitrogen, Cat# AM1560) was used to resuspend pellet, and RNA was extracted according to manufacturer recommended protocol. Total RNA concentration and quality was measured by Nanodrop. RNA was stored at -80˚C.

Ribosomal RNA depletion was performed using the TruSeq Stranded mRNA and Total RNA kit (illumina, Cat# 20020594) and followed by TurboDNase (Ambion, Cat# AM2239) to remove DNA. To prepare RNA-seq libraries, 5 μl of purified RNA was fragmented by mixing 1 μl dNTP mixture (Qiagen, Cat#1005631), 1 μl of 3 μg/ml Random primers (Invitrogen, Cat# 58875), and 6 μl of water. Mixture was incubated at

65°C for 5 mins then placed on ice for 1 minute. RNA was then reverse-transcribed by adding 1 μl of SuperScript IV (200 U/μl) (Invitrogen, Cat# 18090010), 4 μl of 5X First strand buffer (Invitrogen SuperScript IV), and 1 μl of 100 mM Dithiothreitol (DTT). Sample was incubated in a in the following condition: 23 ˚C for 10 minutes, 50 ˚C for 10 minutes, 80˚C for 10 minutes. cDNA was purified using 36 μl Ampure XP beads (Beckman-Coulter, Cat# A63880) and eluted with 23 μl of 10 mM Tris-HCl, pH 8.5. Then 3 μl of 10X NEB buffer 2 (New England Biolabs, Cat# B7002S), 2 μl of dUTP mixure (dUTP: 20mM, dATP, dCTP, dGTP: 10mM), 1 μl of RNase H (Invitrogen, Cat# 100004927), 2 μl of 10 U/μl DNA polymerase I (New England Biolabs, Cat# M0209L), and 0.5 μl of 100 mM DTT was added. Sample was incubated at 16˚C for 2.5 hours. Sample was then purified using 45 μl of Ampure XP beads and eluted with 33 μl of water.

DNA end-prep was performed by mixing 32 μl of DNA, 5 μl of T4 DNA ligase buffer with 10 mM ATP (New England Biolabs, Cat# B0202S), 2 μl of dNTP mixture (Qiagen, Cat# 201901), 2 μl of 3U/μl T4 DNA polymerase (New England Biolabs, Cat# M0203S), 1 μl of 5U/μl Klenow DNA polymerase (New England Biolabs, Cat# M0210S), 2 μl of 10U/μl T4 PNK (New England Biolabs, Cat# M0201L), and 6 μl of water followed by incubation at 20˚C for 30 minutes. DNA was purified with 70 μl Ampure XP beads and eluted with 33 μl of water. A-tailing was performed by adding 5 μl of NEB buffer 2, 1 μl of 10 mM dATP (Promega, Cat# U120B), 2 μl of 5U/μl Klenow 3' to 5' exo (New England Biolabs, Cat# M0212L), and 9 μl water to 32 μl of end-prepped DNA followed by incubation at 37˚C for 30 minutes. DNA was purified with 60 μl of Ampure XP beads and eluted with 25 μl of water. Adapter ligation was performed by mixing 23 μl of DNA, 2X Rapid ligation buffer (Enzymatics Inc, Cat# B1010L), 1 μl of 10 μM Adapter, and 1.5 μl of T4 DNA ligase (Enzymatics Inc, Cat# L603-HC-L) followed by incubation at room temperature for 15 minutes. DNA was purified with 50 μl of Ampure XP beads and eluted with 31 μl of water. The eluted DNA was then treated with 2 μl Uracil-DNA Glycosylase (New England Biolab, Cat# M0280S) and incubated at 37˚C for 30 minutes. The RNA-Seq libraries were amplified by mixing 32 μl of DNA sample with 0.5 μl of Phusion Polymerase (New England Biolabs, Cat# M0530S), 10 μl of 5XHF buffer, 1 μl of 10 μM MP-P2-IdX primer, 1.25 μl of 10 mM dNTP, and 5.25 μl of water. The mixture was then incubated in the following: 98 ˚C for 30 seconds, 98 ˚C for 10 seconds, 65 ˚C for 30 seconds, and 72 ˚C for 30 seconds. Then 1 μl of 10 μM MP-primer-1 was added. The DNA was then amplified by 12 cycles of: 98˚C for 10 seconds, 65˚C for 30 seconds, and 72˚C for 30 seconds. RNA-seq library was purified with 35 μl Ampure XP beads and eluted with 21 μl of water. Twenty μl of library was collected and analyzed for quality with a 4150 Tapestation System (Agilent, Cat# G2992AA). The sequences of Adapter and barcoding primers are listed in Table S1.

**Divergent PCR of chorion-ecDNA**

DNA from *Drosophila* ovary and carcass was purified using the Quick-DNA Microprep Kit (ZymoResearch, Cat# D3020). One hundred ng total DNA (in 10 µl volume) was mixed with 1 µl 10X Plasmid-safe DNase buffer, 0.5 µl Plasmid-safe DNase (Biosearch Tech., Cat# E3101K), and 0.5 µl 100 mM ATP. Controls did not contain Plasmid-safe DNase. The mixture was incubated at 37 °C for 16 hours on thermocycler and followed by 70 °C for 30 minutes. One µl of the digested DNA was used for divergent PCR using CloneAMP HiFi PCR Premix (Takara Bio, Cat # 639398). The primer sequences are listed in Table S1. The amplified product was assayed by electrophoresis through a 0.8% agarose gel.

**5-ethynyl-2'-deoxyuridine (EdU) staining of *Drosophila* follicle cells**

Dissected fly ovaries were stained with the Click-iT EdU Cell Proliferation Kit (Thermo Fisher Scientific, Cat# C10337). The tissue was first washed twice in 1x Phosphate-buffered saline (PBS) and incubated with 20 μM EdU for 60 minutes at room temperature. The tissue was then washed twice with 1x PBS followed by fixation in 4% Paraformaldehyde for 15 minutes. Fixed tissues were washed twice with 1x PBS and detected with Click-iT reaction as per manufacturer's instructions. Tissues were washed twice with 1x PBS and whole-mounted on microscope slides in Vectashield Antifade Mounting Medium with DAPI (Vector labs, Cat# H-1200-10). Stained tissues were imaged using a Leica SP5 Inverted Confocal Microscope.

**ecDNA-sequencing and genome-sequencing of *Drosophila* ovaries**

For ecDNA-sequencing, total DNA from ovaries was extracted by using Quick-gDNA MicroPrep Kit (Zymo Research, Cat # D3021). After removing linear DNA, rolling circle amplification, and debranching as described previously[1], the library was prepared with the Ligation Sequencing Kit (Oxford Nanopore, Cat #SQK-LSK109). Briefly, 2 μg of total DNA was mixed with 2 μl Plasmid-safe DNase (Biosearch Tech., Cat # E3110K), 5 μl 10x Plasmid-safe buffer, 2 μl 100 mM ATP (Thermo Fisher Scientific, Cat # R0441), and ultrapure water (Thermo Fisher Scientific, Cat #10977023) to 50 μl. On a thermocycler machine, the mixture was incubated at 37 °C for 3 hours. Then 2 μl Plasmid-safe DNase and 1 μl ATP were added to the mixture. The mixture was further incubated at 37°C for 16 hours and 70°C for 30 minutes on a thermocycler machine. Then 50 μl AMPure XP beads (Beckman Coulter, Cat # A63881) was used to purify DNA. The concentration of the purified circular DNA was measured by Qubit dsDNA HS Assay kit (Thermo Fisher Scientific, Cat # Q33231). The RCA reaction was conducted as follows: 2 ng circular DNA, 5 μl 10x Phi29 DNA Polymerase buffer, 1 μl Phi29 DNA Polymerase (NEB, Cat # M0269L),

2.5 μl 10 mM dNTP (Qiagen, Cat # 201901), 2.5 μl Exo Resistant Random Primer (Thermo Scientific, Cat # SO181), and ultrapure water to 50 μl. The mixture was incubated at 30 °C for 16 hours and 65 °C for 10 minutes on a thermocycler machine. The RCA product was purified by isopropanol precipitation and debranched by T7 Endonuclease I (NEB, Cat # 0302L). The short fragments were eliminated by Short-Read Eliminator XL kit (Circulomics, Cat # SS-100-111-01). Then the library was constructed following the Oxford Nanopore SQK- LSK109 protocol. All libraries were sequenced in R9.4 flow cells on a GridION instrument according to the manufacturer's instructions.

For genome sequencing, fly ovarian gDNA libraries were generated using the Nextera XT DNA Library Preparation Kit (illumina, Cat# FC-131-1024). Briefly, fifty ng of gDNA was mixed with 10 μL of 2x Tagmentation buffer, 2.5 μl of Tagmentase, and up to 20 μl of nuclease-free water. The mixture was then incubated at 55 °C for 10 minutes. Thirty μl of nuclease-free water was then added to the mix, and bead purified with 90 μl (1.8x) of AMPure XP beads (Beckman Coulter, Cat # A63881). Tagmented DNA was eluted with 10 μl of water. PCR amplification reaction was composed of the following: 10 μl of tagmented DNA was mixed with 5 μl of i7 index primer, 5 μl of i5 index primer, 5 μl of Universal primer mix/PPC, and 25 μl of 2X Ultra II Q5 Hotstart Master Mix (NEB, Cat# M0544L). On a thermocycler, the mixture was incubated at 72°C for 3 minutes, 98°C for 30 seconds, 10 cycles of the following steps: 98°C for 15 seconds, 63°C for 30 seconds, 72°C for 3 minutes, and 72°C for 5 minutes. Libraries were size selected using AMPure XP beads, first at 0.6x to remove large fragments, then 1.2X beads to remove small fragments. Beads were washed with fresh 80% ethanol. DNA was eluted from beads with 28 μl of water. Library concentrations and quality were quantified using Qubit dsDNA HS Assay kit and an Agilent 4150 TapeStation System, respectively. Final libraries were sequenced at 150 bp paired-ends on an Illumina NextSeq 550 Instrument. Table S2 summaries all sequencing libraries prepared for this study.

**Glioblastoma tumor DNA sequencing**

High quality DNA was extracted from both the blood and tumor samples. The tumor DNA was extracted by The Maxwell® 16 FFPE Tissue LEV DNA Purification Kit, the whole blood DNA was extracted by Azenta. DNA Libraries were prepared using the NEBNext Ultrall DNA Library Kit for Illumina following the manufacturer's protocol (New England Biolabs) and were sequenced on a NovaSeq S4 platform with 2 × 150 bp paired-end reads.

**Computational analysis for CRISPR screen and other sequencing experiments**

Raw sequencing read counts from CRISPR screen were processed, and analysis of enrichment and depletion metrics comparing the eGFP+ and eGFP- populations was performed using the MAGeCK software analysis pipeline under default settings [4]. Volcano plot and snake plot was generated using RStudio [5]. To conduct Co-essentiality and Gene Ontology analysis, Project Achilles gene essentiality data was first obtained from the Broad Institute's DepMap portal (20q1 release). The locus-adjusted gene coessentiality was then determined with FIREWORKS (https://github.com/mendillolab/fireworks/) as described [6]. Essentiality scores for each gene were adjusted by subtracting half of the median value of its 40 nearest neighbor genes' (20 upstream, 20 downstream) while excluding those located within duplicate gene clusters, followed by pairwise Pearson correlations conducted at a genome scale. Protein-Protein Interactions (PPIs) were combined to enhance the gene network. PPIs of genes were extracted from Metascape (3.5) [7] which utilizes physical PPIs captured in

BioGrid [8] and STRING database (version 12.0) database (version 12.0) [9] as the main data source. A network was constructed in Cytoscape 3.10.1 [10] for the top 65 drivers for ecDNA biogenesis with beta scores below -0.350, based on protein-protein interactions (PPIs) with coessentiality relations. Additionally, for suppressors with beta scores above 0.375 and a p-value less than 0.05, the network was constructed based on their coessentiality with added PPI information. Gene ontology analysis was conducted using enrichGO in the clusterProfiler R package (https://rdrr.io/bioc/clusterProfiler/) [11]. Top 30 ecDNA drivers with beta scores below -0.350 and top 80 ecDNA suppressors with beta scores above 0.375 were analyzed for enrichment of biological processes (BP). Bar plots for GO analysis were generated using RStudio [5].

All junction sequences were analyzed using CRISPResso2 [12] in batch mode. To detect large deletions, two rounds of CRISPResso2 analyses were performed. In the first round, paired-end reads were processed using the parameters: -q 30 -s 20 --trim_sequences --fastp_options_string '--umi --umi_loc read1 --umi_len 4' --default_min_aln_score 13 --keep_intermediate. The minimum homology score was set to 13, ensuring retention of the PCR primer regions. After this round, an additional filtering step was applied, keeping only reads that contained both the left and right primers. The second round of CRISPResso2 was performed on the filtered reads using the parameters: --default_min_aln_score 13 --keep_intermediate --bam_output. Throughout the analysis, a "pseudo-guide" sequence was used, with its last 3 bases corresponding to the right end of the junction. All downstream analyses and visualizations were based on the alignment outputs from CRISPResso2.

For RNA-seq data analysis, raw reads were trimmed using Trimmomatic v0.39 to remove adapter sequences and low-quality bases with the following parameters: LEADING:15, TRAILING:15, SLIDINGWINDOW:4:15 MINLEN:36. The trimmed reads were aligned to the human reference genome GRCh38.p14 using STAR v2.7.11a with default settings. Gene expression levels were quantified as FPKM (Fragments Per Kilobase of transcript per Million mapped reads) using StringTie v2.2.3, with gencode.v47.basic.annotation.gtf from Gencode serving as the reference annotation. For visualization, coverage tracks were generated using bamCoverage from deepTools v3.5.5, with reads normalized to CPM (Counts Per Million) and a bin size of 10 base pairs. The summary of high-throughput sequencing data was listed in Table S3.

For the detection of ecDNA in glioblastoma patients, the wrapper tool AmpliconSuite-pipeline was used, which facilitated the execution of AmpliconArchitect (1.3.r2) and AmpliconClassifier (0.4.13) from the fastq files. BWA (0.7.17-r1188) MEM was used for alignment to the reference, and CNVkit (0.9.9) was used for copy-number variant calling. GRCh38 (hg38) was used for reference genome mapping and annotation. For oncogene-related ecDNA detection, the parameters were applied as -s *$name* -t 32 --fastqs *$name"_R1_001.fastq.gz" $name"_R2_001.fastq.gz"* --ref hg38 --downsample *cov* --run_AA --run_AC, where *cov* indicated the optimized coverage. A downsampled coverage depth of 30x was utilized for the precise detection of junction sites. The segments from AmpliconClassifier cycle file which indicates "ecDNA-like" cyclic path were used for junction analyses. To analyze the sequences of junction sites, the sequencing reads around ecDNA cycle segment 1000 bp were reassembled using SvABA (1.1.0). From consensus sequence of each junction, the positions and the level of homology was taken from the output. For all detected junction events, they were also checked in the corresponding paired blood samples. The presence of a junction event in the blood sample was considered a germline-inherited events.

**Nanopore sequencing reads pre-processing and mapping**

All sequencing data and analytical pipelines will be released upon paper acceptance. The fast5 files generated by the Nanopore GridION machine were used as input in MinKNOW version 21.05.25 (MinKNOW core 4.3.12). Guppy 5.0.16 is integrated into MinKNOW. The basic data preprocessing parameters are the following: Basecall model = High-accuracy base-calling; Read filtering = 9; The passed fastq files produced by MinKNOW were used for further quality control. Adapter sequences were detected and trimmed by porechop (0.2.4) with parameters: --extra_end_trim 0 --discard_middle. This setting only removes the adapter sequencing detected at the beginning and the end of the reads, and if the adapter sequence is detected in the middle of the reads, the reads were filtered out. Output files of porechop were used for further analysis. Reads were mapped to the reference genome of *Drosophila melanogaster* version dm6 (GCA_000001215.4). Read mapping was performed using the minimap2 (2.17-r941)[24] software with parameter settings -ax map-ont -Y -t 16 to keep the soft clipping sequences for all supplementary alignments in the SAM output. Mapped reads were converted to bam format, sorted by reference coordinates, and indexed by samtools (1.12). Data visualization was achieved by R studioand Python (3.9.12). IGV (2.12.0) was used to visualize mapping results.
