## Supplemental Table 1 for "BRCA1-A and LIG4 complexes mediate ecDNA biogenesis and cancer drug resistance"

**Table S1: Summary of oligos**

| Name | Sequence (5' to 3') | Description |
|---|---|---|
| biosensor-6N-F | NNNNNNATCGCGGCCGCCAGCTGCTC | Primers for introducing 6 random nucleotide ends |
| biosensor-6N-R | NNNNNNATCGAGACGgaattctttgcca | Primers for introducing 6 random nucleotide ends |
| Version 1 junction-F | GGTTCGGCTTCTGGCGTGTGACC | PCR primer for biosensor Version 1 junction (Extended Data Fig 7c) |
| Version 1 junction-R | GAACTTCAGGGTCAGCTTGC | PCR primer for biosensor Version 1 junction (Extended Data Fig 7c) |
| MT-F | GGCTATATACAACTACGCAAAGGC | PCR primer for mitochondrial DNA |
| MT-R | GGTAGATGTGGCGGGTTTTAGG | PCR primer for mitochondrial DNA |
| YWHAZ-F | ACTTTTGGTACATTGTGGCTTCAA | PCR primer for YWHAZ linear DNA control |
| YWHAZ-R | CCGCCAGGACAAACCAGTAT | PCR primer for YWHAZ linear DNA control |
| chorion-ecDNA-F | TGGGCGAGATCACGTTTCGAG | Divergent PCR primers for chorion ecDNA detection |
| chorion-ecDNA-R | ACTCCTTCACGGGGACTACC | Divergent PCR primers for chorion ecDNA detection |
| LIG4-WT-F | TACCGAGCTCGGATCCatggctgcctcacaaact | Wild-type LIG4 primer |
| LIG4-WT-R | GATATCTGCAGAATTCttaTTTATCGTCATCATCTTTGTAGTCCTTGTCATCATCGTCCTTATAGTCCTTATCGTCGTCATCCTTGTAATCaatcaaatactggttttcttcttgtaat | Wild-type LIG4 primer |
| LIG4-K273A-F | aaaccGCgctagatggtgaacgtatgc | Primer for introducing K274A point mutation into LIG4 |
| LIG4-K273A-R | catctagcGCggtttctatgtagaaactct | Primer for introducing K274A point mutation into LIG4 |
| LIG4-E331A-F | ccatcatcGcaccatcaagaatacagatt | Primer for introducing E331A point mutation into LIG4 |
| LIG4-E331A-R | atggtgCgatgatggcctataatccta | Primer for introducing E331A point mutation into LIG4 |
| LIG4-ΔBRCT-R | GATATCTGCAGAATTCttaTTTATCGTCATCATCTTTGTAGTCCTTGTCATCATCGTCCTTATAGTCCTTATCGTCGTCATCCTTGTAATCgttaacgttagtaaggttaggtgct | Primer for introducing BRCT domain deletion into LIG4 |
| LIG4_sgRNA1-F | CACCGGCACTATTTCTATTCTACC | sgRNA to LIG4 gene locus #1, sense strand |
| LIG4_sgRNA1-R | AAACGGTAGAATAGAAATAGTGCC | sgRNA to LIG4 gene locus #1, anti-sense strand |
| LIG4_sgRNA2-F | CACCGCATAATGTCACTACAGATC | sgRNA to LIG4 gene locus #2, sense strand |
| LIG4_sgRNA2-R | AAACGATCTGTAGTGACATTATGC | sgRNA to LIG4 gene locus #2, anti-sense strand |
| LIG4_sgRNA3-F | CACCGCGTCGAAACATACTGAGAGG | sgRNA to LIG4 gene locus #3, sense strand |
| LIG4_sgRNA3-R | AAACCCTCTCAGTATGTTTCGACGC | sgRNA to LIG4 gene locus #3, anti-sense strand |
| PRKDC_sgRNA1-F | CACCGTTCTCAGAAACGATCAACA | sgRNA to PRKDC gene locus #1, sense strand |
| PRKDC_sgRNA1-R | AAACTGTTGATCGTTTCTGAGAAC | sgRNA to PRKDC gene locus #1, anti-sense strand |
| PRKDC_sgRNA2-F | CACCGCACATCATCATGCACCGTG | sgRNA to PRKDC gene locus #2, sense strand |
| PRKDC_sgRNA2-R | AAACCACGGTGCATGATGATGTGC | sgRNA to PRKDC gene locus #2, anti-sense strand |
| XRCC4_sgRNA1-F | CACCGGGAGACTGATCTTTATAAG | sgRNA to XRCC4 gene locus #1, sense strand |
| XRCC4_sgRNA1-R | AAACCTTATAAAGATCAGTCTCCC | sgRNA to XRCC4 gene locus #1, anti-sense strand |
| XRCC4_sgRNA2-F | CACCGGTTCTCTAATGACTTCAGC | sgRNA to XRCC4 gene locus #2, sense strand |
| XRCC4_sgRNA2-R | AAACGCTGAAGTCATTAGAGAACC | sgRNA to XRCC4 gene locus #2, anti-sense strand |
| NT_sgRNA1-F | CACCGCGGGCAGAACGACCCTGACTGG | Non-targeting sgRNA control, sense strand |
| NT_sgRNA1-R | AAACCCAGTCAGGGTCGTTCTGCCCGC | Non-targeting sgRNA control, anti-sense strand |
| RAD50_sgRNA1-F | CACCGCGTAAACTTGACCAGGAGA | sgRNA to RAD50 gene locus #1, sense strand |
| RAD50_sgRNA1-R | AAACTCTCCTGGTCAAGTTTACGC | sgRNA to RAD50 gene locus #1, anti-sense strand |
| RAD50_sgRNA2-F | CACCGACCCTGCGTAAACTTGACC | sgRNA to RAD50 gene locus #2, sense strand |
| RAD50_sgRNA2-R | AAACGGTCAAGTTTACGCAGGGTC | sgRNA to RAD50 gene locus #2, anti-sense strand |
| UIMC1_sgRNA1-F | CACCGATGAGTGAGCAGGAAGCTA | sgRNA to UIMC1 gene locus #1, sense strand |
| UIMC1_sgRNA1_R | AAACTAGCTTCCTGCTCACTCATC | sgRNA to UIMC1 gene locus #1, anti-sense strand |
| UIMC1_sgRNA2-F | CACCGGGTTAGGGTCTACTCCATC | sgRNA to UIMC1 gene locus #2, sense strand |
| UIMC1_sgRNA2-R | AAACGATGGAGTAGACCCTAACCC | sgRNA to UIMC1 gene locus #2, anti-sense strand |
| BABAM2_sgRNA1-F | CACCGCTCCAAAGATAAAATCGGG | sgRNA to BABAM2 gene locus #1, sense strand |
| BABAM2_sgRNA1-R | AAACCCCGATTTTATCTTTGGAGC | sgRNA to BABAM2 gene locus #1, anti-sense strand |
| BABAM2_sgRNA2-F | CACCGAGTTTAAATCGGTCACAGT | sgRNA to BABAM2 gene locus #2, sense strand |
| BABAM2_sgRNA2-R | AAACACTGTGACCGATTTAAACTC | sgRNA to BABAM2 gene locus #2, anti-sense strand |
| ABRAXAS1_sgRNA1-F | CACCGGAGGGGGAGAGTACGTCGG | sgRNA to ABRAXAS1 gene locus #1, sense strand |
| ABRAXAS1_sgRNA1-R | AAACCCGACGTACTCTCCCCCTCC | sgRNA to ABRAXAS1 gene locus #1, anti-sense strand |
| ABRAXAS1_sgRNA2-F | CACCGTGTTAATAGCAGAAAAACA | sgRNA to ABRAXAS1 gene locus #2, sense strand |
| ABRAXAS1_sgRNA2-R | AAACTGTTTTTCTGCTATTAACAC | sgRNA to ABRAXAS1 gene locus #2, anti-sense strand |
| BRCC3_sgRNA1-F | CACCGCTGTTTGGCCTTCACATGT | sgRNA to BRCC3 gene locus #1, sense strand |
| BRCC3_sgRNA1-R | AAACACATGTGAAGGCCAAACAGC | sgRNA to BRCC3 gene locus #1, anti-sense strand |
| BRCC3_sgRNA2-F | CACCGCTTACGACGTTCTGATAAG | sgRNA to BRCC3 gene locus #2, sense strand |
| BRCC3_sgRNA2-R | AAACCTTATCAGAACGTCGTAAGC | sgRNA to BRCC3 gene locus #2, anti-sense strand |
| EGFR-ecDNA sgRNA-L | GCAGCCTAGCCACATGACAA | sgRNA for CRISPR-C of EGFR-ecDNA |
| EGFR-ecDNA sgRNA-R | GGATGGCAAGGAACAAAGAA | sgRNA for CRISPR-C of EGFR-ecDNA |
| Biosensor junction ddPCR_F (5'-3') | CACCTGCCTGAAATCACTTT | Junction primer for CRISPR-C biosesensor for ddPCR experiments |
| Biosensor junction ddPCR_Probe (5'-3') | 5' 6-FAM-ATGGTGAGCAAGGGCGAGGA-3' Iowa Black® FQ | Probe for internal control CRISPR-C biosesensor for ddPCR experiments |
| Biosensor junction ddPCR_R (5'-3') | CTGAACTTGTGGCCGTTTAC | Junction primer for CRISPR-C biosesensor for ddPCR experiments |
| EGFR-ecDNA ddPCR_F (5'-3') | GGCTCACGTGTCCACTTAAA | Junction primer for EGFR for ddPCR experiments |
| EGFR-ecDNA ddPCR_Probe (5'-3') | 5' 6-FAM-AACCTGCGGCAACACCACAAAG-3' Iowa Black® FQ | Probe for internal control EGFR for ddPCR experiments |
| EGFR-ecDNA ddPCR_R (5'-3') | TTCTCAGATGCAAGAGCAAAGA | Junction primer for EGFR for ddPCR experiments |
| GAPDH ddPCR_F (5'-3') | CCCTTCATACCCTCACGTATTC | Junction primer for GAPDH for ddPCR experiments |
| GAPDH ddPCR_Probe (5'-3') | 5' HEX-AGCTTCCCGTTCTCAGCCTTGAC-3' Iowa Black® FQ | Probe for internal control GAPDH for ddPCR experiments |
| GAPDH ddPCR_R (5'-3') | TCTTCCATTCTGTCTTCCACTC | Junction primer for GAPDH for ddPCR experiments |
| EGFR-ecDNA junction_F (5'-3') | ACACTCTTTCCCTACACGACGCTCTTCCGATCTNNNNCTGCACATCCTGCGCATGTACTT | ecDNA junction primer for CRISPR-C deep sequencing experiments |
| EGFR-ecDNA junction_R (5'-3') | TGGAGTTCAGACGTGTGCTCTTCCGATCTCGTATCTGGCAGAACAGCTTTCCA | ecDNA junction primer for CRISPR-C deep sequencing experiments |
| Adapter | ACACTCTTTCCCTACACGACGCTCTTCCGATCT | Adapter for RNA seq library preparation |
| MP-primer-1 | AATGATACGGCGACCACCGAGATCTACACTCTTTCCCTACACGACGCTCTTCCGATCT | Forward primer for RNA seq library preparation |
| MP-P2-Id1 | CAAGCAGAAGACGGCATACGAGAT**CGTGAT**GTGACTGGAGTTCAGACGTGTGCTCTTCCGATCT | Reverse primer for RNA seq library preparation with barcode |
| MP-P2-Id2 | CAAGCAGAAGACGGCATACGAGAT**ACATCG**GTGACTGGAGTTCAGACGTGTGCTCTTCCGATCT | Reverse primer for RNA seq library preparation with barcode |
| MP-P2-Id3 | CAAGCAGAAGACGGCATACGAGAT**GCCTAA**GTGACTGGAGTTCAGACGTGTGCTCTTCCGATCT | Reverse primer for RNA seq library preparation with barcode |
| MP-P2-Id4 | CAAGCAGAAGACGGCATACGAGAT**TGGTCA**GTGACTGGAGTTCAGACGTGTGCTCTTCCGATCT | Reverse primer for RNA seq library preparation with barcode |
| MP-P2-Id5 | CAAGCAGAAGACGGCATACGAGAT**CACTGT**GTGACTGGAGTTCAGACGTGTGCTCTTCCGATCT | Reverse primer for RNA seq library preparation with barcode |
| MP-P2-Id6 | CAAGCAGAAGACGGCATACGAGAT**ATTGGC**GTGACTGGAGTTCAGACGTGTGCTCTTCCGATCT | Reverse primer for RNA seq library preparation with barcode |
| MP-P2-Id7 | CAAGCAGAAGACGGCATACGAGAT**GATCTG**GTGACTGGAGTTCAGACGTGTGCTCTTCCGATCT | Reverse primer for RNA seq library preparation with barcode |
| MP-P2-Id8 | CAAGCAGAAGACGGCATACGAGAT**TCAAGT**GTGACTGGAGTTCAGACGTGTGCTCTTCCGATCT | Reverse primer for RNA seq library preparation with barcode |
| MP-P2-Id9 | CAAGCAGAAGACGGCATACGAGAT**CTGATC**GTGACTGGAGTTCAGACGTGTGCTCTTCCGATCT | Reverse primer for RNA seq library preparation with barcode |
| MP-P2-Id10 | CAAGCAGAAGACGGCATACGAGAT**AAGCTA**GTGACTGGAGTTCAGACGTGTGCTCTTCCGATCT | Reverse primer for RNA seq library preparation with barcode |
