## Supplemental Table 2 for "BRCA1-A and LIG4 complexes mediate ecDNA biogenesis and cancer drug resistance"

**Table S2: Oligopaint probe sequences**

DHFR Oligopaint probes

ACCTGCATGCGGCCCAATCACTGCTAGTAAGCAAAGGCAAAGTTATCTGGCTGACGATCACGG
ACCTGCATGCGGGTTCCCTTCTACCTACCTTTTCACCAAGTGGTTCTTCATGACGATCACGG
ACCTGCATGCGGCTGTCATTCTCTGCACAGGCTCCTCTTCTGGGCTTTGACGATCACGG
ACCTGCATGCGGAAGCTATTGTCCTTTGAGACCCATCTGAACACCATAGTTTCTGACGATCACGG
ACCTGCATGCGGGGGTTATTGTGTCACAAAGTTTTGGAAGGACAGGATGCTGACGATCACGG
ACCTGCATGCGGAACTGGCCTTCTGGGTTATCTATAAGCAGCAACTCATGACGATCACGG
ACCTGCATGCGGGCCTCCTCTCTCCATTTCTCATCCCTGAAACCCTGTGACGATCACGG
ACCTGCATGCGGAAAATGAAAGCGGCATGCAGAGAAGAGGGTCATCTTGACGATCACGG
ACCTGCATGCGGTCAGATTTTGGAGTCCAGACTCAGCCTAAAGCTCATTTTGACGATCACGG
ACCTGCATGCGGTGGGGTTCCTGAGAATCCACAGGCTCATAATACCCTGACGATCACGG
ACCTGCATGCGGATTTGATTAAGCTACCATGAGAAGGCTTCTGTTACACGTGGTGACGATCACGG
ACCTGCATGCGGCAACTCCTCCTCAGATGATCTTTGATGCTATCACTGATTGGTGACGATCACGG
ACCTGCATGCGGAGCAGGAAGACTTGGGAAGCCACTGGGATGGATTATGACGATCACGG
ACCTGCATGCGGACAATTCGGAAAACTCACTGAGAGTCTTCTGAGTGTAGAGCTGACGATCACGG
ACCTGCATGCGGAGGATTCTAGGTATGTGAGCCTCGCCGATGCTTTTTGACGATCACGG
ACCTGCATGCGGGGTGCCTTACATCACATCCCCACCTCTGTTTTCACTGACGATCACGG
ACCTGCATGCGGAACACATGAGTGGTACAGATTCCAGTCCATATAGTTGCTGTTGACGATCACGG
ACCTGCATGCGGATCTCATGGAAGCACTCTGTTCTCAGCTAAGTATGAAGAGGTGACGATCACGG
ACCTGCATGCGGTGGGATTGGGTACACTTTGTACCATATCCTTAGTGATCTGTTGACGATCACGG
ACCTGCATGCGGACAGAACACTTAGGTCTTCACAAAATCATCCTGGAACATGTTGACGATCACGG
ACCTGCATGCGGTAGCCCTTAATCTGTTAGAACTGTATCCTGAACAGCCTTGATGACGATCACGG
ACCTGCATGCGGAAAAGGCATTGTATCAGTTTAGGTTAAATGCATGGCTGGCTGACGATCACGG
ACCTGCATGCGGTTCGCTCTTCTACTTTGTTGATATCAGCTTTCACAGCATCTTGACGATCACGG
ACCTGCATGCGGTCCTTGGCAGTGTCTTTCAGCTCATTCCAGCATCCTGACGATCACGG
ACCTGCATGCGGTGCTTTGCCTTGCCTTTGAGGTCTAACATCCTGGGTGACGATCACGG
ACCTGCATGCGGCCATTCTGTATTTAAGTCATGCACAGTCGCTTGTTTGCATGACGATCACGG
ACCTGCATGCGGGCGGCTGTAGATGAACATCCTCTCATCATCTGCTGTGACGATCACGG
ACCTGCATGCGGGCTTGGTCTCAAGGTGCTTAATATCCTTGGCAGCTTGACGATCACGG
ACCTGCATGCGGTTCTCAAACAGCCTGAGACATACTGGCAGTTGTTGCTGACGATCACGG
ACCTGCATGCGGTGGAGTGTTAGGAAGTCATCCTGAAAGATTTGCTAGTTCCATGACGATCACGG
ACCTGCATGCGGGTTGGCCTTCACCATCCACTATGCAGAAATATAAAGGAGGTGACGATCACGG
ACCTGCATGCGGTGTTTTCCCAATTCCCTTTCTCTACTAGGCCCATTTAAACCTGACGATCACGG
ACCTGCATGCGGAGGAATTCTGGTTGCCATATTCATGAAAGAAGGCACTAGATGACGATCACGG
ACCTGCATGCGGTACATGCTTAATATAGCACCTTGTCCAGGGGTTTTCTCTGCTGACGATCACGG
ACCTGCATGCGGTATACCCCTGAGATATGTGAAGGCCATTCTCAAGTTCTCATGACGATCACGG
ACCTGCATGCGGTTCTAAGTTTAGGCATTTTGGTTTAGCATGAGCGATCCAGATGACGATCACGG
ACCTGCATGCGGTACTAAGCAGTAGGCCTCCTTCAAAATACAATGTCAATGGGTGACGATCACGG
ACCTGCATGCGGTTTTCAGAAAATGAATTTGAGGCATGTCTCCATCGTTTGCCTGACGATCACGG
ACCTGCATGCGGAGGATTATTCATCAAGTCACATCAGAGAAAGCAACCTGGGTGACGATCACGG
ACCTGCATGCGGGCAAGCCAGCTCTTTAGCCATGGATAATTATCAAGGTTTCTGACGATCACGG
ACCTGCATGCGGAACTGACCTCATGCAGAATCAAGAGTAGCATTATGTTCTGGTGACGATCACGG
ACCTGCATGCGGAGAGGCCATCTATGAAGCTGCTGAGTTTTATAGAAAAGCCTGACGATCACGG
ACCTGCATGCGGAGATTCACTGAAGTAGGAAGGGAGACAAGTTGAGAATCAGGTGACGATCACGG
ACCTGCATGCGGTTCTTTAAGAGCCACTTTTGAACTGTCTTCTAGCACTTGCTTGACGATCACGG
ACCTGCATGCGGTCCCATTTACAACTCTACGAATTTTCGCAAATTGAACAGCCTGACGATCACGG
ACCTGCATGCGGAAAGCAGGAATATGTGAGATGTTGTCCTTCACTTTCATCCCTGACGATCACGG
ACCTGCATGCGGATACACTTTATGGTCTAAGCATGTGTTGCCCTGACCTTGACGATCACGG

ACCTGCATGCGGTTACATATCTATAGTCCTTGGCCAGCTCCATCTCACATTGGTGACGATCACGG
ACCTGCATGCGGCTGCCAGTTCTGGGGCTCACATTCTTCAAGCATGTTGACGATCACGG
ACCTGCATGCGGGCTCCTCCTCCTCAGAGAGCCTTGCTCCATCTCTCTGACGATCACGG
ACCTGCATGCGGCTGGGCCATGGGGCCTTCACATTCAGGTTTTGGTTTGACGATCACGG
ACCTGCATGCGGTGTGTGTGTATCTTGACACTCAACACCCTGTCCCGTGACGATCACGG
ACCTGCATGCGGCCCAATCACATGAGGTGATTCTAGTGTTTGTTCCTTGCTGACGATCACGG
ACCTGCATGCGGCATTTCAGGGCAAGGGGCATTGTCTTTGTGCAGATTGACGATCACGG
ACCTGCATGCGGAAAACATGTAATTGTAAATGTGAGTGGGCAGAGACAGCCATGACGATCACGG
ACCTGCATGCGGGCAAGGGAAACCAAATCCCCTCTCAATTCAATGGATGACGATCACGG
ACCTGCATGCGGGACCAGTAGGAGGGTCTCAAGAAACTGAAAAGAGGCTGACGATCACGG
ACCTGCATGCGGCCTGATTACAGTGCCTGTGTGTACAAGGCATCCGCTGACGATCACGG
ACCTGCATGCGGCTTCTCTGCTCTCTCTCTACCCAAATCCAGGGGAGTGACGATCACGG
ACCTGCATGCGGCACTGCTAGCCTCAAGGCTGCCAGAGTGCAAAGAATGACGATCACGG
ACCTGCATGCGGTGCCAGGATAATGGCTTTATCCACGTGTGGCCTTTTGACGATCACGG
ACCTGCATGCGGTATTCCAGGTACAAATAGTTCTTTCCCACGAGCTGTTTCTGTGACGATCACGG
ACCTGCATGCGGCAGGTCTCCAAACAGACAATTACCCGGTTCCACTTTGACGATCACGG
ACCTGCATGCGGGGGCTGTTGTCCTGCCCTCTCCCAGCTCTAACATTTGACGATCACGG
ACCTGCATGCGGCTGCGCAGCCTTTCATGCATTGAATGGGCAGCTTCTGACGATCACGG
ACCTGCATGCGGATGATGCTGCTGCGACAATAGCGACAATAGCGGGGTGACGATCACGG
ACCTGCATGCGGGCAGAGGAGTTTGCTCACCCTCTTGCTTCTCTGAATGACGATCACGG
ACCTGCATGCGGAAGATGCCTCTGCCTGACAGTTACCTACCCTGGGGTGACGATCACGG
ACCTGCATGCGGTATAATCAACATTTCCCAGAAATCACTCACAAGCCACAGGCTGACGATCACGG
ACCTGCATGCGGAGAAACTGCATTCAATGGACATCATTGTTCAAAGTTTGCCGTGACGATCACGG
ACCTGCATGCGGAGCTGAAGGCCGGCTGTTGGAATATTTTCAGATTTTGACGATCACGG
ACCTGCATGCGGAAAAGCAAGAGTGGGATGTCCTATTTGAGCTGGAATTTCATGACGATCACGG
ACCTGCATGCGGAAAATCACTGAACTATGGGGAAGGAGGTGACAATTAAGACCTGACGATCACGG
ACCTGCATGCGGCAACCACAAATCAACAACCTGCGGAATGTGGTTGTTGACGATCACGG
ACCTGCATGCGGACTTAAAGAGAATGTTGCCTTCTTCCCAAGACACTTTCTGATGACGATCACGG
ACCTGCATGCGGTAACTCCAAGGAGGCATATAGCACAATTGCATTTTGTCACTTGACGATCACGG
ACCTGCATGCGGTACGCTGGAAATGCAGAACTGTTCAATCTGAATAAGCATGTTGACGATCACGG
ACCTGCATGCGGAACATTCTTAATTTCAACCAGAATGCCACAAAGTCCTGCATGACGATCACGG
ACCTGCATGCGGAAAATGTTTGAGTTGTCACACAGCCCGAGGGTGAGTGACGATCACGG
ACCTGCATGCGGAGAACACTGGGTGGTGAAGCCCAGCAGGGAACATTTGACGATCACGG
ACCTGCATGCGGCACCAAAGAAGGCCACATGGTTCATATTTCCCTCTCCTGACGATCACGG
ACCTGCATGCGGCTTACATCCGGGAACAAAGATCTTCCAGCGTTGAGTTGACGATCACGG
ACCTGCATGCGGTTGAGGACAGAATGTGGAGAAGTCTATGTTACTAATGGCCTTGACGATCACGG
ACCTGCATGCGGGTTAGGCAAGGACAGGCTTCCTAGTCTAGATGGAAAGGTGACGATCACGG
ACCTGCATGCGGAGAACTAAAACCCAAGCCCAATGTGTAATGTAGGAAAAGGGTGACGATCACGG
ACCTGCATGCGGAAAATTATCTCCACATGCCAGCCAAGCATAGTACTCATTCCTGACGATCACGG
ACCTGCATGCGGAATGCAACATCTGTCTCTGCTCATAAATGAGTCACTTTCCTTGACGATCACGG
ACCTGCATGCGGAACTCCTTTGAGTCTTCTGGTTTTAAACACCTCAAGGACCTGACGATCACGG
ACCTGCATGCGGCTCACTACAAGCAAGATGCATTTCCAAACCCTCCACTGACGATCACGG
ACCTGCATGCGGAAAATGTGGGGAAGCGAGAGTCAAGTTGTAAGCCTTGACGATCACGG
ACCTGCATGCGGGGCTGCCTGTGGATCAGAAACACCTGTGGTAAGCTTGACGATCACGG
ACCTGCATGCGGGTGGGTGAACAGCATTGCTTTTCACTTTTAAGGTAAGACATGACGATCACGG
ACCTGCATGCGGTATTACAGGAAGAGACTGGAATCTGTAGTAACCTGGGAGGATGACGATCACGG
ACCTGCATGCGGAAGCAGATAGCAGGATGGCTGGGGAGGAAGCTGAATGACGATCACGG
ACCTGCATGCGGATTTCCTTTCTAGGTATTTTGCCCATACTTTGGGCACATGGTGACGATCACGG
ACCTGCATGCGGTTTTCTTGCCAAGAGACCCTCATTTTGTTTGTTGGGGTGACGATCACGG
ACCTGCATGCGGAGCAATATGCCAGACCCAGATGATGAATCAAGGTTGGTGACGATCACGG
ACCTGCATGCGGAAAACAGTATTTAAGGGCTGAAATAGGTAGTGCTGCATGGTTGACGATCACGG

ACCTGCATGCGGTTTGGTTAATGCCACTGTAGAGGGCCTTAAAGACCATGACGATCACGG
ACCTGCATGCGGGATTGTCTTTGTGGCCATGTTTATGAGAGACTGTTGGAGATGACGATCACGG
ACCTGCATGCGGTGAACTAGATGCGGTTATGAAAGAGAACAGATTGAAAGGCATGACGATCACGG
ACCTGCATGCGGGTCCTGGGTTTTAGAATCTGGAGATGAAGTTTCTTCATGGCTGACGATCACGG
ACCTGCATGCGGCCTTAGTCATTGCTTTTCTAGCCTCTTCAAGGCCAGTTGACGATCACGG
ACCTGCATGCGGGGGGGAGGAATCCCTAGTCTAAGTTTTGCCAATTCACATGACGATCACGG
ACCTGCATGCGGATCAAAAGTAAAGACAACTGGCAGTGAGAAGTCAACCTGTTGACGATCACGG
ACCTGCATGCGGGGACAGTGAATCAGGTCAACTAGAAGCCTCCTAACTCCTGACGATCACGG
ACCTGCATGCGGTTGTCTCTGGCCTTTTGGAGGGGAACATGACTAGGTGACGATCACGG
ACCTGCATGCGGATTCAGGATCCCTTTCCCTAATTACTTTAGCTGGTAGGTGCTGACGATCACGG
ACCTGCATGCGGCCCCAGGTTTTCCTGACCTCCAGCACCATGAAGTCTGACGATCACGG
ACCTGCATGCGGTTTTCCTCCTTGCAGACAAAACTCTATCCCTTTAGGCTTGACGATCACGG
ACCTGCATGCGGTTGTCTTGTGTTTGTCTTTGTTAGGGCTCCTAGGCATGACGATCACGG
ACCTGCATGCGGTGTATTTACAGAGTAACTTCCTGGTACACATGAGGTCAGCTTGACGATCACGG
ACCTGCATGCGGGGGGGAACAGAAAGGACAGGAGCAGATGATTAGAAACGTGACGATCACGG
ACCTGCATGCGGCCCCGGGTAACAGGAACAGCACTTCATGTGGTTTATGACGATCACGG
ACCTGCATGCGGTAGGCCTTCTTTCTTCTGTTTGGCCAGGGGCTTTATGACGATCACGG
ACCTGCATGCGGTAGTCATTTAATCATGCACTTCTTTTCTTTCTCCCCGCTGGTGACGATCACGG
ACCTGCATGCGGTTCATAGATAACTGTTTGTCTGAATCCTTTCAACGGCTGGCTGACGATCACGG
ACCTGCATGCGGCAGCCAATCTTTTCCTCACAGTCATGTTGCCTAATTAGGCTGACGATCACGG
ACCTGCATGCGGTTTGCTTTCCTGTTTTGCTTCACTGAGATGTTCTAGAGACGTGACGATCACGG
ACCTGCATGCGGAGCAGGTACAACATTTTCAGAAACATTGCCTTTAAGGAAGCTGACGATCACGG
ACCTGCATGCGGTATTTACATGTTTGCCCCAAGTTATCACCTTACAGTGCTGTTGACGATCACGG
ACCTGCATGCGGAAGAACAACAGGAGGCCAGCACTTATTAACAAGTGTAACTTGACGATCACGG
ACCTGCATGCGGACAGCTTCCCTGATTCTCAGTTTAAATGCATCAATAGTGCATGACGATCACGG
ACCTGCATGCGGAGCTGTTTAGAACATAACTGAAATTATGTGCCCTGGACAGTTGACGATCACGG
ACCTGCATGCGGCAGGTTCTCTGCTTCTGTTTGATTGATGGAGACTATACTGCTGACGATCACGG
ACCTGCATGCGGTTAGGATCTGCTATTCCTGTTCCCTAACTCTCACTTAAGCCTGACGATCACGG
ACCTGCATGCGGTGTGTTCTCTCCAGCTCCCCTTCCAAACCCAGCTTTGACGATCACGG
ACCTGCATGCGGATGAACCCCAATCTTTGGTTACTTCTTCAGTACTATGCAGGTGACGATCACGG
ACCTGCATGCGGCAGTATGCTCCAGGTGGTCAGGAACGCCTGCTTGATGACGATCACGG
ACCTGCATGCGGATCAGACTGTGCTTCACTCTGGTTCTGCTGTGTGATGACGATCACGG
ACCTGCATGCGGCATGCCTCCGAGCCTGGCTGTCAATAAAGTCTTCATGACGATCACGG
ACCTGCATGCGGACAATTAAAATCAAGACTCTACAGGAAGCACAGGCTCAGGTGACGATCACGG
ACCTGCATGCGGAGCAAAGGGATATCCGAGTCATGTGGACGAGTATGTGACGATCACGG
ACCTGCATGCGGCCAGCATCCTGACAGGAAGTTTTAGGGTCAGAAGATGACGATCACGG
ACCTGCATGCGGAGAAACATCAGCCCATCCAAAACCTACTTCAAGGTGTTGACGATCACGG
ACCTGCATGCGGCCACTGATAAGGGGCATCTGTCCCAGCTCTGGAGCTGACGATCACGG
ACCTGCATGCGGCAACTACTGCCTCCACAGGCCTCTTTTAATCTGTCTTGACGATCACGG
ACCTGCATGCGGACAGAAGATTGTTAACTAGAAGTAAAAGCCTGTTTGGGGCTTGACGATCACGG
ACCTGCATGCGGAAGTAGAAAACTGCCAATATAAAATATCTGGGCCTGCTGGTTGACGATCACGG
ACCTGCATGCGGAGTCTGAGAAAGATTTCTTCCAGGCAAGATGAATGAGAGCTGACGATCACGG
ACCTGCATGCGGAGAGCCCTAGAGGATAGGCTGGCACAGTGCTGTTGTGACGATCACGG
ACCTGCATGCGGAAAATCGACTGCACAATGACGTACAGAGCTGACGGTGACGATCACGG
ACCTGCATGCGGAACCACCTTACTTTATTAATATGACCGTCTGCTCTGGCTCCTGACGATCACGG
ACCTGCATGCGGACCACAAGAGGTTAACTGATAGAGACAACGGGACTTAAACTTGACGATCACGG
ACCTGCATGCGGAAGTGAGTAGCTGTGGGGAAAGGGATAGGGGAGGTTGACGATCACGG
ACCTGCATGCGGTATATTTGGACAGTAGAAAGCAAATCTAAGTTCCCCAGCGCTGACGATCACGG
ACCTGCATGCGGACAGAGATCTGGAGAGAAAGCCGCTCCAGAACTGATGACGATCACGG
ACCTGCATGCGGTTCTCCATAGAATGAAGGACTGACTTACTCTAGGTCTCCCGTGACGATCACGG
ACCTGCATGCGGACATGCCTGAAAGTACAAATGACCATCAAATGTTGCAATGGTGACGATCACGG

ACCTGCATGCGGTTTGCTTTCTCCCATAGCCTTGTTTAGAATCTCCACTTTGGTGACGATCACGG
ACCTGCATGCGGATACTATTCTTTCATCTGGGGTGCTGCAAAGTATGCCTTGACGATCACGG
ACCTGCATGCGGTGAGAGACTGAGTTTAAATATAGTCTGGCCTCCAGTTTGCTTGACGATCACGG
ACCTGCATGCGGTTTATTGTTTTCCCTTTGGCTGTTTAGAGTTAAGAGCCCCATGACGATCACGG
ACCTGCATGCGGACAATTTAATGAGACCTTGACTATGGAAAACCCCAAGAGCATGACGATCACGG
ACCTGCATGCGGATAAGCCTGAATGATATCTACAAGCTGTTGTGATGTGAGGCTGACGATCACGG
ACCTGCATGCGGTTACAAATGGGGAACATAAGAGGTTCTAGAAACAGGAGCCCTGACGATCACGG
ACCTGCATGCGGTTTCACTGTATATATCTTACCACAAGACCACCCACATGGCATGACGATCACGG
ACCTGCATGCGGACATCTAGTTAGTGTATATCAACTGAGAGCCCCAACCCTTGACGATCACGG
ACCTGCATGCGGTCATCCTGAAAGACAAAGGCATAACTCAGTGTATCTAGGGTTGACGATCACGG
ACCTGCATGCGGACATTCAGGGAGTAGAAACCAGTGCAATGCAAGTACATGACGATCACGG
ACCTGCATGCGGATTTACAGTACAGATAATGTGCTGCTTCCTATACGCATGGCTGACGATCACGG
ACCTGCATGCGGTATACCTGTTTCTTCCACTTCCTTTAAATGGCAAGGAAGCTTGACGATCACGG
ACCTGCATGCGGTGTTTGAATGACTGCAGAAAAGCTTGGAATGACATTCAGACTGACGATCACGG
ACCTGCATGCGGACATTGACTAGCTTAGTCCCTCCCAATCTCTGAAACTACTTGACGATCACGG
ACCTGCATGCGGTCAAGAATAGGGACTATTCATTCTCTTGATTGCCTCCTCCCTGACGATCACGG
ACCTGCATGCGGAAAAGACATGGTGGCCAGAGAGAAGTTAAATAAGCCCATGACGATCACGG
ACCTGCATGCGGGGTATCTCCTAAGGTATCAGGGCAGTTACAGGGACCTGACGATCACGG
ACCTGCATGCGGAGGTTGGATTAGGCTCTTCAGGGAGAGAACTCAATCTTGACGATCACGG
ACCTGCATGCGGTTAGTTAATAAACAGATGATGGCCAAAACAGAGCCGTGACGTGACGATCACGG
ACCTGCATGCGGGTGTTAGTAAGTGCAGCTGTGAAGACTTCCTTGCCTGACGATCACGG
ACCTGCATGCGGAGAGCAAAGGTATGAAGAATATGCTTCCGTAAAATGTGGCTTGACGATCACGG
ACCTGCATGCGGTTAGCAGTTTAGAGGTAAGTATCACAGGAAACACTGGACCTTGACGATCACGG
ACCTGCATGCGGAAAAGCTGAGCTACGGTTTGAGTTCATCAGCCCTTTGACGATCACGG
ACCTGCATGCGGGGCACAAGCGCTAGAAATACAGTAAATGACAAATAGCAAGCTGACGATCACGG
ACCTGCATGCGGAAACCAATCTAGATTGGATTCCCTGACAAAGGAGAGTTGGTGACGATCACGG
ACCTGCATGCGGGCAATTTGTTACATGGACAAAAGACTTCAGTTGGTGCCTTGACGATCACGG
ACCTGCATGCGGGCAGGGTCCTCCGATTCAGGGGCCCTTAAACATCATGACGATCACGG
ACCTGCATGCGGGGATGGTGGGGCTTAGATTAGGCAGAACCCTCTGATGACGATCACGG
ACCTGCATGCGGCTCTTGTCTTCTGGCCTTATCTGATTATCTCGGGTGCTGACGATCACGG
ACCTGCATGCGGTTTTCACTGTTCCTGATGTGAGGACTGCCGATGGATGACGATCACGG
ACCTGCATGCGGATTCTTGCTCAATTTCCCTGAGGAGTGTTGGGAAACCTGACGATCACGG
ACCTGCATGCGGAAAAGAGGTGGGTTACAAAGTGGTACTATGTGGGATGGTGACGATCACGG
ACCTGCATGCGGATACTAAAAGGTTGGGACAGCAGAGCAAGGTGAAGATGACGATCACGG
ACCTGCATGCGGAAAAGGAATCCAGAATCAAGTCAGGGGAGATGAGAGAGTTGACGATCACGG
ACCTGCATGCGGGGGGGAGAGAGAACTGCACCTTCAAGAAGTTTTAATCACTTGACGATCACGG
ACCTGCATGCGGAATGGAGGCAATTTTGGATTTTCTAAAGATGGTGTCCAGGCTGACGATCACGG
ACCTGCATGCGGTAAAGCCACAACTTGACTGTCTAGACAACTTTGGAAAAGCATGACGATCACGG
ACCTGCATGCGGTTGGCATAAGCAAGTGATTTCCATGTATAAATCTCTGCCCTTGACGATCACGG
ACCTGCATGCGGTGTGTGGGACTGACCAACAAGTAAGGAAGTGGAAGTGACGATCACGG
ACCTGCATGCGGAACTGGTATCACTATTTCCCTGGCATGCTTGATAACAAAGATGACGATCACGG
ACCTGCATGCGGATACAGAAGAAGCAACCAAACCCAATTACATGCATCACAGGTGACGATCACGG
ACCTGCATGCGGATTACACTCATCATATTCGAGGAGAAATGCTGGCTCAAGTGTGACGATCACGG
ACCTGCATGCGGATCTACCAAGGAAAGAGATAATCAGAGCATCCTAGCAACCATGACGATCACGG
ACCTGCATGCGGGAAGAAAGAACAGCATGAAGATGGAAGCAGGCAGCTGACGATCACGG
ACCTGCATGCGGAAGATGCACATCAACAAATCTGCCCCGTCGCCCAGTGACGATCACGG
ACCTGCATGCGGAGGAAGAACTTGGGAGAAAATCTATGTAAGCCTGAATCCCCTGACGATCACGG
ACCTGCATGCGGATGACTACTGTTCCCAAGAAAGTAGTGCCCTAGGATACTGTGACGATCACGG
ACCTGCATGCGGTAGACTAAGAATGATTAACATGCCAGATCCCAGACCTTCCCTGACGATCACGG
ACCTGCATGCGGCAATTCTGGCCACCCCTCAACCTCCCAAGCCATTGTGACGATCACGG
ACCTGCATGCGGGAGTTGACTTCCTGGGCAGTCTTAATATGATTACCCCTCCTGACGATCACGG

ACCTGCATGCGGCACTCTCCAGTCACCAGGGCATATCCAGGCTTCATTGACGATCACGG
ACCTGCATGCGGAAAATTCTTAGCAGAGGCTAACTGGAGATACCCGCTTTGACGATCACGG
ACCTGCATGCGGAATTCACATTTTATTGCATTTAGGGTTGGGTCCAGAAAGGGTGACGATCACGG
ACCTGCATGCGGTTGGGCCTACTGAATGATGGTTCAAGATAACTTGCAAATCTTGACGATCACGG
ACCTGCATGCGGTTTGGGGCTAGGTTTAGCTCAGAGTGGTCAAGTTGTGACGATCACGG
ACCTGCATGCGGGCTGGCACGAACTTGACGAACCCTAGGGTTAGTATTGACGATCACGG
ACCTGCATGCGGGGTATGAGCCTGTCGAAACATTTTCAGTGTATTGCTTTGAGTGACGATCACGG
ACCTGCATGCGGTGTTGAGTAAGGTGGGCAGGGAGTTGCATTGATGGTGACGATCACGG
ACCTGCATGCGGAGTAGAATGTGAGTTGAGGGAGGATGATGTGTTAGTTGAGGTGACGATCACGG
ACCTGCATGCGGATGCATACCGCCAAAAGATGAAATTTGAAATCTGGTTAGGCTGACGATCACGG
ACCTGCATGCGGTTTTGAGACTGGCTTTTATCAATTCTTTAGCTACGGGCTCTTGACGATCACGG
ACCTGCATGCGGTGTCATCATCTCTACCAGTGACCTAAGTGTCAAACCCATGACGATCACGG
ACCTGCATGCGGAATGCCTTGTATCTGTCCCATTAAAGAGATGCAGCATCTTGACGATCACGG
ACCTGCATGCGGGCTCCTTTCTTACTGTTTCCATTTCCTCTGCCATGCTGACGATCACGG
ACCTGCATGCGGACAACCATAAATATCCAGGTCTCTTAGGTTTTAAACGGGGCTGACGATCACGG
ACCTGCATGCGGCCCCACATTCTTTTCCTTGGTTATTCCCTTCCCTCCTGACGATCACGG
ACCTGCATGCGGAACAGTTCAATTCACCTAGATCCCCACGCCTGAAATGACGATCACGG
ACCTGCATGCGGTTATCCTAGATGTCCTAGAGGCGCCTCATCATTACAATGGTGACGATCACGG
ACCTGCATGCGGTACATTATTCTCCACTCCTTTACATGTCACGCCAGCTTTGACGATCACGG
ACCTGCATGCGGTCAAACTGAAAATCTGAGCGTTCATCCCTGGTGCATGACGATCACGG
ACCTGCATGCGGTTTTACCCTCTCTTCTCCACTGCCCTTGTTCAGGCTGACGATCACGG
ACCTGCATGCGGCTTATCTCTTCCAGCAGCTGTTCCAAAGGCCTACTTGACGATCACGG
ACCTGCATGCGGCTGTTTTCCTTTCGGAGTGCTCACCTCCACCGAAGTGACGATCACGG
ACCTGCATGCGGCACCCAGCTGCCAATTCTGCCCCATGCCTGATAATTGACGATCACGG
ACCTGCATGCGGAACTAACATTTGCACTGGTAAACTTCATCAAGCAAGACCCTTGACGATCACGG
ACCTGCATGCGGAAGTTAGAGAAACAGCGTTACTCGAAACATTATCCCTTGGGTGACGATCACGG
ACCTGCATGCGGCTCTAAGGCACCTGACAAACGGAGCGCTGTGGGTATGACGATCACGG
ACCTGCATGCGGCTGCGAGGCAAGCGGTTTTGAGCCGATTCTTCCAGTGACGATCACGG
ACCTGCATGCGGTCTACGGGAAGCCTGAAATCCACCTCCTCCTCCACTGACGATCACGG
ACCTGCATGCGGCCGGGGCTACAAATTGGGTGAAGCGCTGAGGTTTTTGACGATCACGG
ACCTGCATGCGGGCTTGTGGGCAAGGGGTGCGTCTTTTAACCTCCATTGACGATCACGG
ACCTGCATGCGGAAAAGTGCTGGATTGGGTGACTAGAAGAAAGCTGCTGACGATCACGG
ACCTGCATGCGGAAGACTGCAACAGCATGGCTCAATATTTGAAATCACTACGGTGACGATCACGG
ACCTGCATGCGGTTAAAAGCAGTCATATTGTGCAGTTCCCCAGTGAACTCTTGACGATCACGG
ACCTGCATGCGGATGGTATAGATGTGACTAATCATGGAGTCCTTCCTGTTGCCTGACGATCACGG
ACCTGCATGCGGTTTTAAACCAGAGATACTGCCACAGGAAAAGCCCAAGGTGACGATCACGG
ACCTGCATGCGGGTGACCCCTCACTTGGGTCACCTGCACAGTATTTTTGACGATCACGG
ACCTGCATGCGGTCAGATTGAGGCTGTGGAAGGGCATTGAAGACTGGTGACGATCACGG
ACCTGCATGCGGTGTTTAGCCACCTACCGTGTTTTAATGTAGGTGGCATGACGATCACGG
ACCTGCATGCGGTGAGGACGAGCTACGTTTTCATGAATTCACCAAAATTTTGCTGACGATCACGG
ACCTGCATGCGGACCCACCATTTAAAGGGTAGCTGGTGAAGGACTTATGACGATCACGG
ACCTGCATGCGGTGTTCTTTTATTACCTATCTTTTCCCCTCTGTGCCTTTGCTTGACGATCACGG
ACCTGCATGCGGAATTAGAGAAAGCTGAAGGCTCATTGAATCAAGACCGGTTGACGATCACGG
ACCTGCATGCGGAGCAGCCTTGGTTTCAATTAATTCTAAACACGGCTTGATGACGATCACGG
ACCTGCATGCGGGTTGCTGAGGCTCCTGACTATCAGACTCTACCTTGTGACGATCACGG
ACCTGCATGCGGAACAGTTCACCCATAGGGTTTTCCAGAGCTAAAAGTAGAGGTGACGATCACGG
ACCTGCATGCGGATTCTTTGAAGAGTGGATGGCAAGGAACCTTGTCATTCATGACGATCACGG
ACCTGCATGCGGTTGTAAGGTTTAAATTGCTTCACATACGTCAGGCCTTCTCATGACGATCACGG
ACCTGCATGCGGATTGACAGAAGAAAGAAGAGACCATTGGAAAATGATGGGCCTGACGATCACGG
ACCTGCATGCGGAGTAAAGAAAGTCCAACAAAAGGAAGGAGGAAGTGATCTGGTGACGATCACGG
ACCTGCATGCGGTTTTCTCCTCAGTCATGGCTCTGGTATTCTGGAATTCTCCTGACGATCACGG

ACCTGCATGCGGTTTTCTTTGTGGAGAAAGGTCCCTTTTGTTCCTGAGCTGACGATCACGG
ACCTGCATGCGGCCCCTATAGGTGATGGTGATGTGTATAGGAGGTTAAGGGTGACGATCACGG
ACCTGCATGCGGTGGTAGAGCTACAATTCTGCAATTTAGCTGTAGTTCTCCCATGACGATCACGG
ACCTGCATGCGGTGAACAATTTTCAGTATCAGGACCTGCAGATCTACTCTCCCTGACGATCACGG
ACCTGCATGCGGCCAGCCTGCCTGAGTGCCTGAGTTCCTGGATTTCTTGACGATCACGG
ACCTGCATGCGGTGGTACCAGAGAATTTGTGATAAGGGAAGACAGGAAATCCATGACGATCACGG
ACCTGCATGCGGAAAGTAAATCTAAAACTTCACCTTTCTGGAGGGGTCATGCATGACGATCACGG
ACCTGCATGCGGAGGAGGCGTGACTTGAGAGAATTTGCCACGGTTAATGACGATCACGG
ACCTGCATGCGGTGTTACAATTCTGGGGCAGCCTACCTTTCATTGCTTGACGATCACGG
ACCTGCATGCGGAAGTAACTTAACTAAGACCACATGGGGACAGTTGAGATGGATGACGATCACGG
ACCTGCATGCGGGGGGAGGATCATTCACTAGTCAGAAGCTGGGTGACTGACGATCACGG
ACCTGCATGCGGTCTGCCCCATATAATCTGCAGGAACCCATATTTGATTTTGGTGACGATCACGG
ACCTGCATGCGGTTCATCTAGTAGCATCTGCCAGACATAAGAGTCCCTGATGACGATCACGG
ACCTGCATGCGGTTTTCCCAGCAAATACAACTCCAAGATCTTCATCCCACTTGACGATCACGG
ACCTGCATGCGGTGTGGCTAGAAAATTCACAGGAGATCTAATGAAAGCCTGTGTGACGATCACGG
ACCTGCATGCGGTTAATAGGGAAGGTAAGAGGTGATAAATGTGTGCAGTCCCATGACGATCACGG
ACCTGCATGCGGAAACGGTGAAATACTGTGTTCTAGTACATGGTAGGGATCCATGACGATCACGG
ACCTGCATGCGGTCTTCTTATTTGCTGCCTAAGAGCCAAAGAAATGTCTGAGGTGACGATCACGG
ACCTGCATGCGGAAAATTGAAAGAATTCTGCTGCGATTCTGCCCTTCCTTGACGATCACGG
ACCTGCATGCGGCAAAGTAGAGTCCAGACAGAATCTCTGCAGGAGAGATTTGCTGACGATCACGG
ACCTGCATGCGGATCAGTCTTCTACACGCAAAGAATGCAGTTTCTTCTGAAGATGACGATCACGG
ACCTGCATGCGGTCCAGTTACCTAGAATAGTGGGTTCTGAAGTACTGTCAGGTTGACGATCACGG
ACCTGCATGCGGTCCTTAGAGTTTCCATGATTACCGTGGAAAATCATGATGGGTGACGATCACGG
ACCTGCATGCGGAACTGCAGAATCAGGGAATTTAACTCATGCTCCATCATTGATGACGATCACGG
ACCTGCATGCGGTGCCATTATTACAGGTGATAAGCTGTTTGATGGCATGGATGACGATCACGG
ACCTGCATGCGGAAAAGCTACTGAGTGAACATAAATGCCCCATGGTCCATGACGATCACGG
ACCTGCATGCGGGGACCATTAGGGTGGGGTCAGCAATGCCATAACTCTGACGATCACGG
ACCTGCATGCGGTAATGACCTGGATTTCTCTTGCCAACCTCTCCATGCTGACGATCACGG
ACCTGCATGCGGCTCACCGTCTTCTTATTGCTGATCTTTGGAGGTTTGCTGACGATCACGG
ACCTGCATGCGGTTGAAAAGTTAGTATCTTTACCTGATGTCCCGGTGAAGCTGTGACGATCACGG
ACCTGCATGCGGTTCCCATTTCTGGGCCATTTTGCTTCTCTCTCACTTGACGATCACGG
ACCTGCATGCGGAGAAGTTGGCAGTGACAGAGTGGCAATTAATTAGAAGTTGGTGACGATCACGG
ACCTGCATGCGGAGTAGGAGTACTAATGTTGAAAATGTGGCCTTGTGCTGTTGACGATCACGG
ACCTGCATGCGGTAAAACATTAAGTGCAGCTTTATCTGTTGGCATGCTTTGCTTGACGATCACGG
ACCTGCATGCGGCAAAGTGGTTTTGCCCTACTCCTGAGAGAGAGCCATGACGATCACGG
ACCTGCATGCGGGATTTGTCCCCATTGGCCTTTTGTGCATTACACACTGACGATCACGG
ACCTGCATGCGGATTTACAGAAAACTGCTTCCAAATCAGCTAACAAACGGTCCTGACGATCACGG
ACCTGCATGCGGTACGCCGCTAGAATTACAATACATAGAAATGAAGCAGCAGCTGACGATCACGG
ACCTGCATGCGGAAGTATAGATTCTTTGGGGAAGATGCAGAGGTAAGTCGTCTTGACGATCACGG
ACCTGCATGCGGAAAATGAACACAAGAAACTTGGTGGGAACCAGACTTGGTGACGATCACGG
ACCTGCATGCGGAAAGTGAGGGGTCTCTATTGGAATGTTAGCGCTTATTCTCCTGACGATCACGG
ACCTGCATGCGGTCATAGCCAGGAACAAAGTTAAGTCTCAATGGAAGAACACGTGACGATCACGG
ACCTGCATGCGGACCAGCTACCACCAGAAGTTGTTTGAAGTTTTACAGATCTTGACGATCACGG
ACCTGCATGCGGCTTTGGGTGTCAGTCAACTTGGTGTCTTAGTCTAAATTGGGTGACGATCACGG
ACCTGCATGCGGTTTCATAGGCAAACTTAGGGAAAGTATTGCAGATGACACCTTGACGATCACGG
ACCTGCATGCGGAGTCATGAATGCTTGTGATGATTCTATAGTTTGGGACCTGGTGACGATCACGG
ACCTGCATGCGGTTGGATTGTACACTAGAGAGTGATGAGGTAGCATGTCTGTTGACGATCACGG
ACCTGCATGCGGGCTTCTTGGATGGGAAGTAAAGACTTGGCCAACTTTTGACGATCACGG
ACCTGCATGCGGTTTTCTTCAAGTTTCCTAAATCCTCTGTGGCAGGCAAGCTGACGATCACGG
ACCTGCATGCGGAGCGTTCTCAAAGTCAGATGCATTATGCAATGTTCTGTGTGACGATCACGG
ACCTGCATGCGGAAATTAGGCTCTGAGATAATGTCACCATGTTTTCTGAGGGCTGACGATCACGG

ACCTGCATGCGGAGGAACAAATATGTAGTTGTCATCCCCTAGGCGAAGGTGACGATCACGG
ACCTGCATGCGGGAAACTGTCTAGGTTAAACGGTAAATTCAGACCTGACCACGTGACGATCACGG
ACCTGCATGCGGTTTTCTGCCCTTCTGTGAGGGACTAATAGAGTTCAGAGATGACGATCACGG
ACCTGCATGCGGAGGGGTGAGGTTGGGGAGCGAGAGTGTTAGAATTATGACGATCACGG
ACCTGCATGCGGGAGTTCCATGGCTTCAGTGGGCAATATCTAAGCCATGACGATCACGG
ACCTGCATGCGGTGGTGCTTTCTAGAGGAAATCAGGGAATCACATGGTTGACGATCACGG
ACCTGCATGCGGTTTCTATCAGTCGTGCATCATTAATTAGTTAGCCAGCCTGGTGACGATCACGG
ACCTGCATGCGGTTTCAATGATTTCTCTGTCCGGATGGGTTCATCATTTCTGCTGACGATCACGG
ACCTGCATGCGGACTCTGTAGTGTCTTCTAAATGTCAGTCATTTGCACATGCCTGACGATCACGG
ACCTGCATGCGGAGTCTTTGTGTAGGGAAAGCTCTAAGGATGTAGAAGACCTGTGACGATCACGG
ACCTGCATGCGGATACAGATGTTGATACTGAAGTTAGGTCACTGGAACTGCCCTGACGATCACGG
ACCTGCATGCGGAGATTGCAGCCCGAGAGCTCAATATTTATTGCCATTTAGATGACGATCACGG
ACCTGCATGCGGCAGCAAGTATACCTACTCACAGACTGTTTGTTCATGTACGCTGACGATCACGG
ACCTGCATGCGGCGCCTGGTGGCAAAAGGATATAAGGTCAGCTTTGGTGACGATCACGG
ACCTGCATGCGGCTTTAACTTGTGGGGAAAGGAAATTGGGATTCTCCTCCATGACGATCACGG
ACCTGCATGCGGGAGAGTGCAAACGTGTTTTGTAAGGTGGTAGGTAGGCTGACGATCACGG
ACCTGCATGCGGAGCAAATTATCCACCCTTGTCAACTCCTTTAAACCCTACATGACGATCACGG
ACCTGCATGCGGTTTTGTTGAAGGTGGGAGTTGTGAAGCAAACTGAAACTTGACGATCACGG
ACCTGCATGCGGGCAGCATTAAAGGCCATTGGAGACAACAGAAGTTCATGACGATCACGG
ACCTGCATGCGGTTAGATGAGCTGAGAGGCATTAAGGTGAACCCTTGATTAGGTGACGATCACGG
ACCTGCATGCGGTGTGCTATTCAAGAGGGAAGTAGTGGCTGGAAACATGACGATCACGG
ACCTGCATGCGGAAATAGGGAACAGCTATCACTCAAAATGGGCTCCAAACTTGACGATCACGG
ACCTGCATGCGGATTAACAGCCTTGTGTTGTTCTAAGCTGATCAGGCTGTTGACGATCACGG
ACCTGCATGCGGGCTTGAGTCTGTTATTTGAAGCCCATTTCTAGTACAGTCCCTGACGATCACGG
ACCTGCATGCGGTATGTCCTGATGTCCAAAGATATGTCCAGACTTTGCCCTGACGATCACGG
ACCTGCATGCGGATGGGATTGTTAAGGATTCAAGTCTCAGCATCATCCAAGTTGACGATCACGG
ACCTGCATGCGGGATGAAGTGGGTTGATATTTGGGCCTCCATGGGGTTGACGATCACGG
ACCTGCATGCGGCTGGTGTACTCCACTAAGCCCTTTGGAAGTAAGATACTGGTGACGATCACGG
ACCTGCATGCGGAATGTGAATCCCCTAATCAAGCTGGATGATGCTGTAAATGTTGACGATCACGG
ACCTGCATGCGGATACTCTATGCTATGGAGAGATCCAGATGGTCTTGGTCCATGACGATCACGG
ACCTGCATGCGGTGTGCCCTGAGATTTTAAGGTTGAAGATAGACACAATGGCTGACGATCACGG
ACCTGCATGCGGACAGATAGCAGAATAAAACACTAAGCCAACACAAGCATGGTTGACGATCACGG
ACCTGCATGCGGATTTGCTCATTTACAGACTTACGTGGACTCTGCTAAAGGTCTGACGATCACGG
ACCTGCATGCGGTTTTAAGGCCCTTCAGAACAATCAGACACCCCTTTTCTTGACGATCACGG
ACCTGCATGCGGTGCTTAACCAATAAGCCCTTCTTTCAGAGAGTTAGCATTCCTGACGATCACGG
ACCTGCATGCGGTATCCTTGAGAAGCACGTTTTATCTAGCTACAGGCATTTGGTGACGATCACGG
ACCTGCATGCGGTCCTACTTAGTGCATTTAGAAATACCATCTGGCTTACCGCCTGACGATCACGG
ACCTGCATGCGGAGAGTTAACCATGTGTGCGAACTGGGACTCGGAAATGACGATCACGG
ACCTGCATGCGGAGTCTGAAATTCAAAGGGCTTTGCTGTTGTAGTATCACTGCTGACGATCACGG
ACCTGCATGCGGGTCAGGCTCTGTGCCTTCAGTCTGGGCAGTGCAGTTTTATATGACGATCACGG
ACCTGCATGCGGATGTACTGATAGTCCATCTCTCTAGGCTCTAGAGAAAGCCTTGACGATCACGG
ACCTGCATGCGGTTAATTACAGTTAGAGGAGCTATAACACTGCCTCCAACCCATGACGATCACGG
ACCTGCATGCGGTCTTTGCTATTATTGGCTTAGGATGTTAGGGAGCAGGTTCTGACGATCACGG
ACCTGCATGCGGTTGGCATGATTATTCTACTGGATAGACTGTCTACTTGGGCTTGACGATCACGG
ACCTGCATGCGGATTTACTACAGTACTGGTGGTCTGGTTACTAGGACCATGTCTGACGATCACGG
ACCTGCATGCGGTATCTGTGAGTCTGACTCTTCGTGTAAGAGTCCTGAAAAGCTGACGATCACGG
ACCTGCATGCGGTTTCCAGGACTCTGCTTCTCGTTCAGAGCTAGAAACTGACGATCACGG
ACCTGCATGCGGGATGTCAAGCCTGCAGCCAGTAGAGCTGCTGCTTCTGACGATCACGG
ACCTGCATGCGGCCTTGTCCGAGCAAACAGAGGCGCTCATCCACAGATGACGATCACGG
ACCTGCATGCGGGCCACATCTGTTAGGTAAGTTGGCACATCACTGGATGACGATCACGG
ACCTGCATGCGGACAAAGTTTATGTAACTTTGGAGATGCTTTGTCTGTGCTGCTGACGATCACGG

ACCTGCATGCGGAAGTTTCAATTCATGGCTGTGTCCAGTGGTTTTGTTTACTGTGACGATCACGG
ACCTGCATGCGGATCTAAGATCAGACCACTTACGTATGAAGCAGTGTGCTAGGTGACGATCACGG
ACCTGCATGCGGCGCTGTGCGTACAGAGGAAGAAGATGTGGTTATGATGACGATCACGG
ACCTGCATGCGGTATATGTCTGGCAGAGTTTACAACCTCACATGAGAGTCAGATGACGATCACGG
ACCTGCATGCGGTTAAAGGCATGATGTATTAAGTGCTAAACGTGAAGGTGCCTTGACGATCACGG
ACCTGCATGCGGTGTTGGAAAGACTCCGTACTGCCACCTGGAGTGTGTGACGATCACGG
ACCTGCATGCGGGTGGGAAAGAGACCAGAAAGGTGTGCTGGGGCTTCTGACGATCACGG

EGFR Oligopaint probes

ACCTGCATGCGGCTCAGATCTTCACCCTGCCAAATAACGTGTTTCTCCTTGACGATCACGG
ACCTGCATGCGGCCCCTTTCACAGAGCATTTGGTTTTAGGAAATTCAGAGCTGACGATCACGG
ACCTGCATGCGGAAAAGTTTCTCATGTACAGCTGGCAAAGGGATGGAACTGACGATCACGG
ACCTGCATGCGGTTTTAGGTCCTGACATCACCATTTGTCTCAGGTTTTGAAGCTGACGATCACGG
ACCTGCATGCGGGGATTCAAAGTTGGCCTCTCATACTAGCAAGTATACCTTGGTGACGATCACGG
ACCTGCATGCGGTATCCTGGTCACTTCTCCCGGCCACAGCATCACATTGACGATCACGG
ACCTGCATGCGGTTAACCAGCTCACAGGTTATCAGATAAGCTTAGTCTGACCATGACGATCACGG
ACCTGCATGCGGATGCTTAACACAGCAACTGGGCCACTATTGTCATTTGACGATCACGG
ACCTGCATGCGGCATGGTCTAGGACCACCCTCCACAGAGGCTGTGAGTGACGATCACGG
ACCTGCATGCGGCTAGAGCCCTAACTGTGCAGGGCCCTAACTATGCCTGACGATCACGG
ACCTGCATGCGGACAAGCCGTACTATGAAGGCTGTTGTCTCATATAGTCCTTGACGATCACGG
ACCTGCATGCGGATTGATTTGTGTTGTGACTTGTACACACAGGTCACATTCCCTGACGATCACGG
ACCTGCATGCGGGCCTCTGCATTCAGGGCATGACTTCAACGCACAGTTGACGATCACGG
ACCTGCATGCGGAGGAACAGATTACTTCTCCTAATTCACAGGGAAGTTCCAGGTGACGATCACGG
ACCTGCATGCGGGGGTGCGAGACCCAGGCACAAACATTTTGCTGGATTGACGATCACGG
ACCTGCATGCGGGAGGAGGAAAGATGTAAGGTTGCTCCCCTTCAGAGTGACGATCACGG
ACCTGCATGCGGTTTTGACAGAGCCGAGAGATCAGGGTTGTTGAACCTGACGATCACGG
ACCTGCATGCGGGAGGGGTCTGGCTGTAACATGGACCTAGAGGACATTGACGATCACGG
ACCTGCATGCGGTTTTACTGCAGGAGAAGGAACAGTGGGGATGGGGTTGACGATCACGG
ACCTGCATGCGGGGACTTGCCAAAGGAATATAGCTCAAGTTCCTGCAGCTGACGATCACGG
ACCTGCATGCGGAAAAGCTCAGTTTCTTTTGGCCAAAGCTTCCGCGATGACGATCACGG
ACCTGCATGCGGGGAGCTACAGGGGCAGTGGGACACTTAGCCTCTCTATGACGATCACGG
ACCTGCATGCGGAAAGCACCTCCACGGCTGTTTGTGTCAAGCCTTTATGACGATCACGG
ACCTGCATGCGGTTCCAAGAGCTTCACTTTTGCGAAGTAATGTGCTTCACTGACGATCACGG
ACCTGCATGCGGAAACATTAAGGAGGCCTGTCTCTGCACCCGGAGTTTGACGATCACGG
ACCTGCATGCGGGGGTGCCCTCATTTCAGATGATTTCGAGGGTGCTTTGACGATCACGG
ACCTGCATGCGGGACAAGATCTGAAGGACCCTCGGACTTTAGAGCACTGACGATCACGG
ACCTGCATGCGGGTCGGTGCCATTATCCGACGCTGGCTCTAAGGCTCTGACGATCACGG
ACCTGCATGCGGGGCCAGTCTGTCTAAAGCTGGTACAAGTTTGCTTTTGACGATCACGG
ACCTGCATGCGGGGACCCTGGCACAGATTTGGCTCGACCTGGACATATGACGATCACGG
ACCTGCATGCGGGCAGTGCTGGGAACGCCCCTCTCGGAAATTAACTCTGACGATCACGG
ACCTGCATGCGGCCCGCCCGTCCTTTCCTGTTTCCTTGAGATCAGCTTGACGATCACGG
ACCTGCATGCGGCCCCGAACCGCTCCCAACTTTCTTCCCTCACTTTCTGACGATCACGG
ACCTGCATGCGGCCGCCGAGACTGCACTGTTTAGGGAAGCTGAGGAATGACGATCACGG
ACCTGCATGCGGCGGCGGTTAGTTTCCTCGTTGGCAAAAGGCAGGTGTGACGATCACGG
ACCTGCATGCGGTCAAAATGGGGCTCACAGCAAACTTCTCCTCAAAATGACGATCACGG
ACCTGCATGCGGCGGATTGCTGTCCCTGGTTCAAGTGTGCCAAGTGTTGACGATCACGG
ACCTGCATGCGGGCAGACAGAACATGAGCGAGTCTGGCTTCGTGACTTGACGATCACGG
ACCTGCATGCGGCTTGGAAGGTTTAATTGCACAATTCCAACCTTGAGCTGCTGACGATCACGG
ACCTGCATGCGGGGTTCCAAGAGCCAGGCCCGTACTTGCTGTTGATGTGACGATCACGG

ACCTGCATGCGGAGACGGAAGAAGGATTGTTAAAGTCTCCGGAGATGTTACTTGACGATCACGG
ACCTGCATGCGGTGCCAATGCTAAGAGCTCTTTGAGGACATCTGGAATGACGATCACGG
ACCTGCATGCGGATAAAGGATTCCGCTTTTCATTGAAGGAACTGGTGGAAAGGTGACGATCACGG
ACCTGCATGCGGGGAGACTGCTGACTGCCGGTCCTGTTAGTCAGGTGTGACGATCACGG
ACCTGCATGCGGTCAGCCCTGTCTCTGCCGAAGAGACTCTTCTCTTTTGACGATCACGG
ACCTGCATGCGGTAAATACCACTTAATTTGCCTGTTTGTCCCGCATTCACTCATGACGATCACGG
ACCTGCATGCGGAAACAGAATGCTCCTGAAGACAAGAGAGAGAGTAGGAGAATGACGATCACGG
ACCTGCATGCGGTCAAGATGCACTTTGTTTAGTGGGAGTTTTGTTTGGTTCTGTGACGATCACGG
ACCTGCATGCGGTGCTAACTTCTAGACTTGCTGCTCTTAGAGGTAATGACTGCTGACGATCACGG
ACCTGCATGCGGTCCGAGTTCTGGTTTCAGGAGATCCAAATCAGGTGTGACGATCACGG
ACCTGCATGCGGTGTGCAAATGTCTAATGTCAGAGCTGGCAAGGGGATGACGATCACGG
ACCTGCATGCGGCAGTCTGCATTCCTGCCGAGTTCCTCAGCCCTCTGTGACGATCACGG
ACCTGCATGCGGTTGGGTCACCTTCCATAGAGGCAGCTTAGTCCTCATGACGATCACGG
ACCTGCATGCGGGTTCAGTGAGCATGGAGTGGAGACTGCTTGAGGGGTGACGATCACGG
ACCTGCATGCGGTGCTGAGCAAAGCCCTGCCTCTTACAGGATGAAGGTGACGATCACGG
ACCTGCATGCGGTGCTCTCCAGAAGGGACACTGGAAAGTATTCCAAGTGACGATCACGG
ACCTGCATGCGGGCGAGTCGAATTCCCAACTGAGGGAGCTTTGTGGATGACGATCACGG
ACCTGCATGCGGCCCAGCCCCACTTCTGGAGACGTTCCCATTCAGTATGACGATCACGG
ACCTGCATGCGGATATTAATGGTATCCAAAGTGAGATCTACCCCACCCTCCCTGACGATCACGG
ACCTGCATGCGGTCCTCAAAGGAGGTCAGATCAAGAAAGCCCAAGCCTGACGATCACGG
ACCTGCATGCGGCGGCCTGGCAATTGGGACCTTTCTTCTCACTCCAGTGACGATCACGG
ACCTGCATGCGGCCCAGGGTGAAGGTGGACAAGTCACTTTGACCCTTTGACGATCACGG
ACCTGCATGCGGGGGGTGAGTTTTAAGGGTTTCTGCTCTTGCTTCCATGACGATCACGG
ACCTGCATGCGGGATTGCTTTCAAATCTGGAAAGGACACCAGTGGTTTGTTGACGATCACGG
ACCTGCATGCGGGTGTTAGACCCACACTGCCGTAGCACAGAATACAATGACGATCACGG
ACCTGCATGCGGAAAACTTCTGGTTTGAAGACACCGATTGCCGAAAGTTGACGATCACGG
ACCTGCATGCGGCCATTGTGCTGCATAATTACACTTGGTCCACGTGATGACGATCACGG
ACCTGCATGCGGTTTAGAAGTAGTCTCAGCAAAGATGAAGGATTCCTCCCTGTTGACGATCACGG
ACCTGCATGCGGTATTAAAGGCAATTGATGATTATGGGCACTGAAGGGAAGGTTGACGATCACGG
ACCTGCATGCGGCTGGTGCCCCGGAATAGGGATGGGTCACAATGTTGTGACGATCACGG
ACCTGCATGCGGAGGACATTTCGCCTGTTGCAGAACCCACCTGCAACTGACGATCACGG
ACCTGCATGCGGTCAAAGTTTTAGAATATGGTTCACCCACATGTTGCTTCCCTTGACGATCACGG
ACCTGCATGCGGTGAATTTTCTAAGAAAATTCCAGCAGTTGGCTCTTTGGACGTGACGATCACGG
ACCTGCATGCGGACATCTCTAGATTGTCCTCCATTGGGCCCATAGGCTGACGATCACGG
ACCTGCATGCGGACAAGCTGGCCAGTTTGAATTTGGGCAAGAATCCATGACGATCACGG
ACCTGCATGCGGTAGATACACTCAATCCAGTTGTCTAGAAAGTTCCCTGAGCCTGACGATCACGG
ACCTGCATGCGGGGGAGCAGGAGGGGTAGTTGGGGCCAGGAATATTGTGACGATCACGG
ACCTGCATGCGGGGGGTGTGTTTACTGAGCCCCTAGAAAGTAAGTGCTGACGATCACGG
ACCTGCATGCGGGTATCAAATGACTGGTCCTGTGGACTGAGCATCTGTTGACGATCACGG
ACCTGCATGCGGAATTTTACTCCTACCAGTTTCAGCAGCTTGCTTTAGCAAGCTGACGATCACGG
ACCTGCATGCGGTCCAAGTTTTATCTCATTCTGGTGGGTTTTCAAGGAGAGACTGACGATCACGG
ACCTGCATGCGGTCGGAGTCCAAGTGTCCTTTCTGAATATATCTGGAACTTCTTGACGATCACGG
ACCTGCATGCGGAACAAAAGACTCAAGTTATAATTTAGGGGACAAGGCACCCATGACGATCACGG
ACCTGCATGCGGATGAGAATGCCTTGCAGGCAGCCCTAAGTACACCTTGACGATCACGG
ACCTGCATGCGGGCAATTACACCATTACTAGCGCGGCAGCACACATGTGACGATCACGG
ACCTGCATGCGGTATGTTTGAGCCATGATGAGTGATTTACACTGTTGCATCCATGACGATCACGG
ACCTGCATGCGGTTAGACTTGGTAATTGCCCTTAAGTGTCTGGAAGTCAACTGTGACGATCACGG
ACCTGCATGCGGAGTGTGAAACAATGTCACAGAATCAATGACGGAACTCTTCCTGACGATCACGG
ACCTGCATGCGGTTTGAGTTCAGTCTTTGCCTTTAATTCTACATGGGGAGGAGTGACGATCACGG
ACCTGCATGCGGAGCACGTTTAGCCACAAATGGAAGGGATTACTCATTTGTGACGATCACGG
ACCTGCATGCGGGGTTATATGATTATTTCCCCAGAGAATAGGATGTGCAGGGCTGACGATCACGG

ACCTGCATGCGGATTACACAAGCAGTGCCAATAGCAGCAAAGTTCTTGATGACGATCACGG
ACCTGCATGCGGTGTCTTAGATCACAGAACAAGTTCTCCTCTCACAGTTTCCTTGACGATCACGG
ACCTGCATGCGGGCTCCACCTGTCTCATGCTCACCGTCAGCATCGAATGACGATCACGG
ACCTGCATGCGGATTGAGCCACACCAGGGGTTCTGGATACCAGCTTCTGACGATCACGG
ACCTGCATGCGGTCTCTAGGTGAGGCTGCTATAGTCAGCAGCTGATTTGACGATCACGG
ACCTGCATGCGGATATATTGCTGCAAATACTCGGAATGGGGATCTCTTGTTCCTGACGATCACGG
ACCTGCATGCGGTGCTTAAGACCACATCACATTACTTGGTTTTGTACGCTAGTTGACGATCACGG
ACCTGCATGCGGTGTAGCTCGTAACGAAAGAAATCTTGTCTTTGCTCTCAGACTGACGATCACGG
ACCTGCATGCGGATATACTCCAGGCACATCGCATGCTGGGATCTAGATGACGATCACGG
ACCTGCATGCGGTACACCAAGGGAACAAAATAACTGCACTTGTCCTCTGATGACGATCACGG
ACCTGCATGCGGGGACCGACTTACCTTTTGGAAGGGCTGAGAAAGAGTGACGATCACGG
ACCTGCATGCGGTGTTTTATAACAGATGTGATTTGGGATTTCAGTGGGAGCCCTGACGATCACGG
ACCTGCATGCGGAAAAGAGGGACTGACTAATTCAGCCTCTGTGACAAGGTGACGATCACGG
ACCTGCATGCGGGGAGTTTCTCAGAAACAGAATGCTTAGCTGGGCCTTGACGATCACGG
ACCTGCATGCGGGGCCCTGTGCTCCTTCAGCAAAACCCTCAGTTTCTTGACGATCACGG
ACCTGCATGCGGTGGAGTCCAGGTTATAGTTGATAGCTTTAACTTTCTCCCCATGACGATCACGG
ACCTGCATGCGGAGGTAGATTTGCAACAAATAAAGAGTGGAGTACAGCTGCTTGACGATCACGG
ACCTGCATGCGGAATGTGATATCTGACGATTTTACCACTACATGCTTGCAGGCTGACGATCACGG
ACCTGCATGCGGCAGTGCACAGCAGATGACGTCATGATTGTTTTAGCATGACGATCACGG
ACCTGCATGCGGTTGATGCATTGCACATAGTCTCTCTGATAAGACAAACTGGCTGACGATCACGG
ACCTGCATGCGGAAACAAACCACTTAGTGAATTTGTGCAGGTCCTTAAGGACATGACGATCACGG
ACCTGCATGCGGAAGTTTGTTCTAGTGCTAACTTGCAAGATCTAATGGCTCCATGACGATCACGG
ACCTGCATGCGGAAAACCCTGGAACCCCAATGTCAGATGTCATCCACTGACGATCACGG
ACCTGCATGCGGACAAGTATGCATACACATGTATTGCTCCCTGAAAAGTGGTTGACGATCACGG
ACCTGCATGCGGAGTTATAATGGGGAGAGAAAACAGGTTAGAGTCTCCCCTCTGACGATCACGG
ACCTGCATGCGGTCACCTTCCATAAAACAGCTAAACTAGACGATCGTCAGACTTGACGATCACGG
ACCTGCATGCGGCTGCTGTTGTCCGTAATCACCAGGCTGGTGATCAGTGACGATCACGG
ACCTGCATGCGGGCACTGGGTGCTCTCCTGCTACACAGCACTGTCTCTGACGATCACGG
ACCTGCATGCGGGGATGCCCCATCACTTTTAGTTCTATTTGTGAATCAAAGGCTGACGATCACGG
ACCTGCATGCGGATGTGTTATAACTGCCCATTCCATAAGATACAGGGCAGTGTGACGATCACGG
ACCTGCATGCGGAAAATGAAAGGAGAAGGCAACTTGATTTAGCAGTTGGGTCTGACGATCACGG
ACCTGCATGCGGAGTTAGCAATGCCTATGGCAAGCTGTAGTAATTCCCTTGACGATCACGG
ACCTGCATGCGGAATGAAGGCATACACTTAACCTCTTTAGGGTGTGAAACAGCTGACGATCACGG
ACCTGCATGCGGAAAAGAGACAAACTTAAGAAACAGTGTGGCCCTCCAAGATGACGATCACGG
ACCTGCATGCGGTAATAAGCTATTCTGGCTGGGATTTACTTGCAAGCATTGGCTGACGATCACGG
ACCTGCATGCGGTTCTGGGTTGAAGGAATGGAATGGAGAGTGCGGCATGACGATCACGG
ACCTGCATGCGGGTGAGTAGATCTCTCAGTGACGGTGATGTGCCTCTTGACGATCACGG
ACCTGCATGCGGAATGCAGTGTTCATTTTCCTCCACAAGAAAGGAAGAAACTGTGACGATCACGG
ACCTGCATGCGGACATAGTGGAAACTTTTCAGTACTCTGGCAGAAATTTCCCATGACGATCACGG
ACCTGCATGCGGAGTTGTTATCTAGGTGCTGCAGGTTTAGAGAGGATTGCTGACGATCACGG
ACCTGCATGCGGCAGGCCTGTGATGACGTATCTCTTGTTGAATAAGTAAACCCTGACGATCACGG
ACCTGCATGCGGAGTTAGTATATTGATTTCAGGGTGGCTTTAGCCACTGAACCTGACGATCACGG
ACCTGCATGCGGAAAATCTACCTGGCCAAGGCAGAACAGAAAGTTCAGCTGACGATCACGG
ACCTGCATGCGGATTTGATGAAGTGGGACAACATGAAGAATCAGGTGAGTTGCTGACGATCACGG
ACCTGCATGCGGCCTTTAGAGATTCTTGTTTAGATGCAGAGTAGTGACGTGCCTGACGATCACGG
ACCTGCATGCGGTGGTGTCAGGGAGAGAGTTGAATGAGAAAAGTCCCTGACGATCACGG
ACCTGCATGCGGAGAAGGGCAGAAGACTTGGGTGATTATCTGAGTCCATGACGATCACGG
ACCTGCATGCGGTTCCTTATCACATGACAGAGTTCTTGAAGTCTTGGCTAGGATGACGATCACGG
ACCTGCATGCGGTCAAACTATAGCCAGGGAGATAAGTAGTCACGAACTCAAGGTGACGATCACGG
ACCTGCATGCGGCCTAAATTCTGCTGATGGAGCCGATGAGAATTGGGTTGACGATCACGG
ACCTGCATGCGGGCTAAGGCAAAGAGAGTTGCCAATATTATATTCTTCGGGGTTGACGATCACGG

ACCTGCATGCGGAGCATTCCTCTGACTTAATATTGAGAAGACATTGGGCACTCTGACGATCACGG
ACCTGCATGCGGTTTTCCTCCCACACTTGTCTTCTTTCACTAGGTGACAATGACGATCACGG
ACCTGCATGCGGGGGAAGAGGTAGCATGAGGTGGTGGTCACAGGTGATGACGATCACGG
ACCTGCATGCGGGGGCTGTTGTGAGCACAGGCATGTTGACTGCACATTGACGATCACGG
ACCTGCATGCGGATTAACAGTAGTACTGATCTCACAATCGCCCTATGTCCCATGACGATCACGG
ACCTGCATGCGGTTCACAAGATGTTCTGCCAAGCCATAAAAGGCCCATGACGATCACGG
ACCTGCATGCGGAACATACCAATTCAGTGGTATCATCCTTAGAGGCAGACAGATGACGATCACGG
ACCTGCATGCGGGGATGATTAAATCATTCAGCCCATCTCTGTCTGAGGACGTGACGATCACGG
ACCTGCATGCGGAAGAGTTTGCTCCCGTGAATTTTCACTCTCTATGTAGAACGTGACGATCACGG
ACCTGCATGCGGGACCCGGACGTGTTCACAGTTTGACACATCTGACTTGACGATCACGG
ACCTGCATGCGGCCCAGATCAGGGACAGCTAGCTTTGCTGGCTGGTTTGACGATCACGG
ACCTGCATGCGGATGCATCCTGGATGGCATAGAGTTGATTCTCCTAACAAATCTGACGATCACGG
ACCTGCATGCGGAAATGGAGAAAATGTTCAGGAGTTTCTTGACTCAGAGAGGGTGACGATCACGG
ACCTGCATGCGGTTTTCAGCCAGGAATTTACTACCTTTGCAGGGTAAAGGGTGACGATCACGG
ACCTGCATGCGGGACTCACCACGCTGGAAGTCAAAATAAGCCACCAGTGACGATCACGG
ACCTGCATGCGGTGCCAAGTGTTCAAAGCCCTTAGAATCACAATGCTTGACGATCACGG
ACCTGCATGCGGAAAGCAAAGTCTTCAACAATGCTTGAAAACTTCCACTGGTTGACGATCACGG
ACCTGCATGCGGTTGTTTTCATTAAAGTGACTTGTTAGCCTCTTGCTCCCTGTTGACGATCACGG
ACCTGCATGCGGCCCTGGGAAGTCGCCAAGAGCCAGACAGGAGAAAGTGACGATCACGG
ACCTGCATGCGGGCTCCACTTGGCTCTCCTCTTTGGTGACCATCCCTTGACGATCACGG
ACCTGCATGCGGGGGACTCTCAGGTGACATCCCACCAACCCTCACTTTGACGATCACGG
ACCTGCATGCGGTGCTTCCCTGGTGGGTCTCACTTTCCCTCAAGAGTTGACGATCACGG
ACCTGCATGCGGTTTTGTTTCCTGCATAGTCCTGGGCCAGTTTTGATTGACGATCACGG
ACCTGCATGCGGACCCTCTTCATTTCACTTCAGAAACCCTGATGATTTCTTCCTGACGATCACGG
ACCTGCATGCGGTTTTACCTTAGGACTTTTACTATGACGACTGTGACTGGCCCTGACGATCACGG
ACCTGCATGCGGCCATGCACACTTTCCAAGGGTCCAGCTGTAGATCTTGACGATCACGG
ACCTGCATGCGGTCATGGTTCCCCAGGTGCCTGGACCATCTTGTGAGTGACGATCACGG
ACCTGCATGCGGCTGGAGCACTTGGCCCTTTCGGCAATGTTTTGGCTTTGACGATCACGG
ACCTGCATGCGGTTTGATCTGGAATTAACTGAGCTACTGCAGGAATTGCTTGATGACGATCACGG
ACCTGCATGCGGTTCACTGATGACTGGTGTTGAGCCAGTACACACCCTGACGATCACGG
ACCTGCATGCGGACACCCAAGGACTGTGACTGTCTTCTGAGGTCCATTGACGATCACGG
ACCTGCATGCGGAAAATGCGCCAACCATCTGTCCTTCCTCTTTATCTGATGACGATCACGG
ACCTGCATGCGGTAAATTAATGCATGCTGGAGACTTGCTGTTTCCTACTAGCATGACGATCACGG
ACCTGCATGCGGATAAATTGCTTTGCTGCAAGAAACACCAAACATGGAAAAGCTGACGATCACGG
ACCTGCATGCGGATGGACATCACTAGTGGCATAGTTGCTTTAAACAGTGAGAGTGACGATCACGG
ACCTGCATGCGGTAGATATTTGATTTGCAAGTGGGATGAAGGGTGGTCTAACCTGACGATCACGG
ACCTGCATGCGGTTTGTCCTGTGTTTACCTTCCATGAGATCCTAGAGGTTGTTGACGATCACGG
ACCTGCATGCGGACAGCACAGTAGTGGCATGTGACACACTTGAGAGTTGACGATCACGG
ACCTGCATGCGGAAGTGGCCTTCGAGCTTAACACATAAGACTTGGGATGACGATCACGG
ACCTGCATGCGGTTTTAGGTGTTAGGAACAGGGAAAACTGTGTGGTTAGGAAGTGACGATCACGG
ACCTGCATGCGGTTTTGGTCTTTAAGGTTTGGAGGCCATGGAGGCATTGACGATCACGG
ACCTGCATGCGGTGAGGGAGAGAGAGCTAAGATAGATAAAGACAGAGATGGGGTGACGATCACGG
ACCTGCATGCGGTTTGGTCTGTTCTGCCTTCTTTATTTTCAGGATAGGTGTCCTGACGATCACGG
ACCTGCATGCGGTATATATGGTTCACCTGAATCCCCAGCCATTTGGTAGAGATGACGATCACGG
ACCTGCATGCGGAGAAAATTCAGCTTCTCGACGGAGGTATTGGCTTTAAAGTTGACGATCACGG
ACCTGCATGCGGAAACCTCTAAGGAACACCTCCTCTCTTAACAGGCATTAACCTGACGATCACGG
ACCTGCATGCGGAACTGCAGAAACTGCAGAAGGACAGGGCTATTTGGTGACGATCACGG
ACCTGCATGCGGGAATAACACAGCTCCCTTCCTTGTCTGTTCCCTCCTGACGATCACGG
ACCTGCATGCGGCATTGTCAGGCTTCTGTGGAGCCATATTCAGAGCATGACGATCACGG
ACCTGCATGCGGGGGGAAGAGAAAATCAACCCCTTGGTGAAGGAAAGTGACGATCACGG
ACCTGCATGCGGCTCCCAATTCACAGAGCAAACATGGGTACTCTTGTTTGTGACGATCACGG

ACCTGCATGCGGCTCCCAGCTCACCGAGCATTCTGAGCCCTGATCCTTATGACGATCACGG
ACCTGCATGCGGGCTCAATTCAACTCAATAGGCATGTGTCAAATGTCTACTGCTGACGATCACGG
ACCTGCATGCGGAGACTGAGCACTGAAAAGCTGCTGGGTACAGGGTTTGACGATCACGG
ACCTGCATGCGGACATGGATAGAAAACGTAGCCTCTGACCCCTAAGGATGACGATCACGG
ACCTGCATGCGGGCCTGTAATCCAGATCCCCATTCTTTCCATCCCATTGACGATCACGG
ACCTGCATGCGGCCTCAGAGCCTTCCGCACGCGGTGTCTCATTTGAATATGACGATCACGG
ACCTGCATGCGGCTTGTGTGAGGATAGCCTCATACCCCTCAGTGAGCTGACGATCACGG
ACCTGCATGCGGTAACACCTTCCTCAGGAGATCTATCTCAGTTCTGAATGCTGTGACGATCACGG
ACCTGCATGCGGTCTAACTAAGAAGGATATTTGGCTACATGCTGGGAAGAGGGTGACGATCACGG
ACCTGCATGCGGGCACGCCGCGATTCCACTCCAGCATTTCCAGTTAGTGACGATCACGG
ACCTGCATGCGGGGCGCCCAGTTAGTTGTGTACTCTGGGCTGTCCCTATATGACGATCACGG
ACCTGCATGCGGCTGGAGTCCTAAAACACTTACGACTGCAGATAGGGGTGACGATCACGG
ACCTGCATGCGGAACTTGATTCTGAGGACACCAGTGAGGTGGTAGTGTGACGATCACGG
ACCTGCATGCGGTGCAAATGATGTGTGAGGAAACTTTGGAGGAGTCTCATGACGATCACGG
ACCTGCATGCGGCTGGATGAAGCTTGTCCCACCAGAGGCTCGCTTGTTGACGATCACGG
ACCTGCATGCGGTTAATAGGCAGGGGTGGAACAGTTATGAAACTGCTCTTGACGATCACGG
ACCTGCATGCGGTCATTTTCTCAAACTGGGAGTGACAAATGTATCCACTTGGATGACGATCACGG
ACCTGCATGCGGTCCATCACTTTAAGTGAATGTAGGCCCTGGATACAGTGGTGACGATCACGG
ACCTGCATGCGGGAGCTCAGAAGAGCAAATCAGTTGGTCACCTTGCTTGACGATCACGG
ACCTGCATGCGGTTTTACTAAGGGCATCAGTAAGGCTTTCTATGACCTGCTCCTGACGATCACGG
ACCTGCATGCGGTTCAATGCTTGGTTGACATTTGGGGAGCAAAGATAAACTTGACGATCACGG
ACCTGCATGCGGATCTCAAACCTCTAGGTGGAGGAGTGCAGAGATGATGACGATCACGG
ACCTGCATGCGGGATGGTGGAAGCCTGCAGGAGAGCTGAACACCTGATGACGATCACGG
ACCTGCATGCGGAGACACCCAGTGGGAAGACCAGGACCTTTAACGCCTGACGATCACGG
ACCTGCATGCGGCATATCTGCTGCTCAAGACTGGCAGAGAGAAGAGGTGACGATCACGG
ACCTGCATGCGGGTTTGTGATGAGAAAAGGTGGTGAAAGGCACAAGGATGACGATCACGG
ACCTGCATGCGGGGCACAGAGCATGTCAGGTCCCATATCCCAAAAGGTGACGATCACGG
ACCTGCATGCGGGGAGAGCTCCTCCATGGCTGGAGGCATTCAGAGACTGACGATCACGG
ACCTGCATGCGGATAAAGGGAGTGAGAAGAGAAGGTCTGTGGATACTTGAGTGTGACGATCACGG
ACCTGCATGCGGAAATTGTGTACATCCCATTTCTTTCCACATTTTCCTGGGCTGACGATCACGG
ACCTGCATGCGGTGTCACAGTGGCTGCAAAGAAAGCAGTCCGTGAACTGACGATCACGG
ACCTGCATGCGGTGAACTGTGATCCCAGACAGGCAAGCACACCAGGATGACGATCACGG
ACCTGCATGCGGATCTCTTCTCAGCTGTTGATAATGAGGGAGCGCTGTGACGATCACGG
ACCTGCATGCGGGGGAGAGAAATGGGGTCCTCTTTGAGTTTCCTCTGTGACGATCACGG
ACCTGCATGCGGTCTTCACTGACAGTTCTCTTACTATACACTGATGGTGACGCTGACGATCACGG
ACCTGCATGCGGAAAGCCACTGTGCTTTGCTGTGGATTAGGATCAGATGACGATCACGG
ACCTGCATGCGGTGTTCCAACAAGGAAAGTTGCTTATTGGAAAGTTTTGCTGCTGACGATCACGG
ACCTGCATGCGGAGGGAGCCTTGAGTTCTGCATCAGGCTTGGAAGTGTGACGATCACGG
ACCTGCATGCGGTCAAAGCGATATTAATCCTGAGACCTGCAGCTAAAGTCAAGTGACGATCACGG
ACCTGCATGCGGTAGGGATAATTAATAGGAGGAAGGTGGGGTTGGAAGATCTGTGACGATCACGG
ACCTGCATGCGGTCATTGGGTTATGAAATGACTGGGGAAAGACTTTTCTTGCTTGACGATCACGG
ACCTGCATGCGGCTGTGTGTCTCCTGTAAATAAGTGCATTAGCAGTACACAGGTGACGATCACGG
ACCTGCATGCGGAAAACGTGGAGTGAGCCCAAGGTGCCATTTATCCATGACGATCACGG
ACCTGCATGCGGGCCCATACACATCGTACCATGTACAGAGTGGACACTGACGATCACGG
ACCTGCATGCGGATACACATCTGCAGCCCTAACACATCTTTATTTGCTAACGATGACGATCACGG
ACCTGCATGCGGTATGTCCAAAGGAGTGAATAGGCCTTATCTGAAACAGGTGATGACGATCACGG
ACCTGCATGCGGATAGAAGGATCTAAGAATTTTCTGCAGCATTCACCAGCACCTGACGATCACGG
ACCTGCATGCGGACCTTGGAAACAGCTATTAACTGGTAACCTCAAGAAACACCTGACGATCACGG
ACCTGCATGCGGTCCAGTAATGACAAGAGGTAGAAGTTACATCCTTGCTGTCTTGACGATCACGG
ACCTGCATGCGGAAATGGAATTTCACGGACAAGAAAATCCATGTCCATTTGGTTGACGATCACGG
ACCTGCATGCGGTTTTACACTTGGTACATGGTTGCAAGATTGCCCCTTGACGATCACGG

ACCTGCATGCGGGGGGTTAATGCAGGTCACTCTACGTATGTGCACGATGACGATCACGG
ACCTGCATGCGGGGTCGTTATAAAGCTCGAAAATATGGGCTCACCAACCATGACGATCACGG
ACCTGCATGCGGAGTAAAATAATGATAACAGGTCAAACTTTGCGGTCCCACGTTGACGATCACGG
ACCTGCATGCGGGAGGCTGGAGATGCGTATTGTCTTGACTTTGCATCTGACGATCACGG
ACCTGCATGCGGTGAATTTCCTTTCGCTGTGTTTACGATAGACTCTCACGTCTGACGATCACGG
ACCTGCATGCGGATTCCACCTCTTTCAAATCGGATGGTACCTATCCTTCAAACTGACGATCACGG
ACCTGCATGCGGTGCTATTTAATGACTGTCTGCTATGTTCAAGGCACTGCTTGACGATCACGG
ACCTGCATGCGGATCTGAAATATGGCCTACGAGCTGAGTTGTGAATCAAGTTTGACGATCACGG
ACCTGCATGCGGAAATTGCATAAGAATGCAGTCCTGTTCTTCATTCCTCTTGCTGACGATCACGG
ACCTGCATGCGGTTTGCAGTTAATGATCCTTTGCCTGGACTTTCTAAGTGCCTGACGATCACGG
ACCTGCATGCGGCAGAAGAGCAACAGCCAGCATGCAGGATGGCATTCTGACGATCACGG
ACCTGCATGCGGAGTGAAAGCAAGAGCCACGTGAAGACGCCTTAGTTTGACGATCACGG
ACCTGCATGCGGATATGCACCCACCCAGACACTTGCTCAGAAAGGAATGACGATCACGG
ACCTGCATGCGGGCCCTGGCCTTAGAAACTGGCTCCTTCACTGCTGTATGACGATCACGG
ACCTGCATGCGGTTTTCCTGTCTTGGCAATGCAAGGATGCCATTAGGTGACGATCACGG
ACCTGCATGCGGTAAAGAAATCCTTCACCACACTAATCCTGCAGAGCCATGACGATCACGG
ACCTGCATGCGGGAAGAGAAACCAGCTTGTTCTAACCCAGCTTTGTCATGTGACGATCACGG
ACCTGCATGCGGTCTTTTGGTAGCAGCCTGCCCAAGGGTGAAAGTGTTGACGATCACGG
ACCTGCATGCGGAGTGAACAGTAAATGTATGCCTGTTTCTGCTTTATGGGACTTGACGATCACGG
ACCTGCATGCGGCATTTGTGGTGCTTCGAGCGAGTTACCAGAGTCCTTGACGATCACGG
ACCTGCATGCGGCATTTGGGCAGGGCAGGTGACCAAGGGTCTTGTTGTGACGATCACGG
ACCTGCATGCGGGAACTGAAGTCCAGCTTGAAAAGCAAATCTGGTTGTGATGACGATCACGG
ACCTGCATGCGGATGGGATCTTAGTCCAACTCTCTTATTTAACGAGGTCCACATGACGATCACGG
ACCTGCATGCGGGAGGTTCTGCGATTGTCTAAGAAAGAAGGCTGTGTTCATGACGATCACGG
ACCTGCATGCGGCTCCGTGAAAGTCCTGATGCAGACATTCCGGCCATTGACGATCACGG
ACCTGCATGCGGACTCAAGAGAATAAGTAAGTTATGGAGTTGGAGACCCCAGCTGACGATCACGG
ACCTGCATGCGGATTGCTCTCTCTTTATGTCTGTAATGCAGTTGCTCCTCTCTTGACGATCACGG
ACCTGCATGCGGAAAGTGGCCTGAAGAGAGGCTTCTAGACTATACTCACGTGACGATCACGG
ACCTGCATGCGGAGCGCTCTAGACAGTACTATGGGTAGTATCTGTGGGTGACGATCACGG
ACCTGCATGCGGGAGGTTCTCAGAAACATGTCGCATGCTCTTTTAGAACCTTGACGATCACGG
ACCTGCATGCGGTAAAGTATTCCTAGTCTCCTCTACTTCCAGCCCTTGGCTGACGATCACGG
ACCTGCATGCGGTGAATTCAGTTTCTTTCTGACATTCTGCTCTCTACAAGGGGTGACGATCACGG
ACCTGCATGCGGATAATGAAGTGAGGCCTAATTCCTTACTCTTCAGAGAGCCCTGACGATCACGG
ACCTGCATGCGGACTGTGGAAGTGTCACTGACCTTGTGTATGGGCTGTGACGATCACGG
ACCTGCATGCGGCCCTTCATGGCTCTGGGAGTCATTATAAAGGGCAGTGACGATCACGG
ACCTGCATGCGGCATTTGGCGTGGTGCGTCCTAAGCCAGTGTTTCTCTGACGATCACGG
ACCTGCATGCGGAAAACAGCATTCTGAGGTCAAACATGCTCAGAAAGCTTGACGATCACGG
ACCTGCATGCGGGGGGAAAGCAGGGTACTTGCTGCTTTGCAAGTGTGTGACGATCACGG
ACCTGCATGCGGACGATAGGGCCATTCCAACAAAGCTGTAAAGTGGTTGACGATCACGG
ACCTGCATGCGGACACAGTCATAAAGAAGAGGAGATAGCTCTGGGAGAAAAGGTGACGATCACGG
ACCTGCATGCGGGTCTGCAGGTTCCTGGTCCTGATTGCAGGCCATTTTGACGATCACGG
ACCTGCATGCGGGGGGTATCACCCAGAACACCTAGTACACGAATTTCAGTGACGATCACGG
ACCTGCATGCGGAGAGGACGAAGCATTACTGGAGTATTGTTATGCAGGAAAACTGACGATCACGG
ACCTGCATGCGGTTAAGTGGAAAACATATTGTGTGTACTTGATCTGGCCTGCTTGACGATCACGG
ACCTGCATGCGGTAACCACATTTCATGACACGGCTAGAGAAGAATCATGCATGACGATCACGG
ACCTGCATGCGGTTTCCAACATGTTCGTAAACTGGGAAAGTATTTCACTGGGGTGACGATCACGG
ACCTGCATGCGGAAGAATATCTTTAAATGACACATGCCTTGCTCTGGGACGATGACGATCACGG
ACCTGCATGCGGGGCATCTGCATGGGTGACACATATGTGTTGTGTGTTGACGATCACGG
ACCTGCATGCGGCTCCCAGCATTTCCAGGGCCCTGCTCAGAATGTAGTGACGATCACGG
ACCTGCATGCGGGCCTTACTGATTCTTACAGAGTTACAAGCGCTGGTGATGACGATCACGG
ACCTGCATGCGGGGTTGGCGAAGTTTAGGTAAACACAGCTGGGAATGTGACGATCACGG

ACCTGCATGCGGTTTTACCCTGATAGGGCCTTCCATGATGCTTTCAAAGTTGACGATCACGG
ACCTGCATGCGGGTGTTCATTGCTCTTGTATGGTACTGTCTGTGGCACTGACGATCACGG
ACCTGCATGCGGTGACTCTAAGAATTTAAGGAGCATGTGAGTGTTGATGGCTCTGACGATCACGG
ACCTGCATGCGGTAAAAGGGTAACAGAGCCCAACTAGCTCAGTTCTCAGCTGACGATCACGG
ACCTGCATGCGGATGAAAATAGTCATATGGCACAGACTCAGTGGAGTGGGTGACGATCACGG
ACCTGCATGCGGTGCACTTCAATAACTGGAAGCACAGATGCCCTACATGACGATCACGG
ACCTGCATGCGGATGGAAACGCACATTTAGTATGCATATGACAACACGAAGGATGACGATCACGG
ACCTGCATGCGGCAGAAAGCTCTACCGCCTGTTTATAAGCTCCTGCATGACGATCACGG
ACCTGCATGCGGTGAATATAACTTGAAATGACCGTAATCTCCTGCCCCAACTCTGACGATCACGG
ACCTGCATGCGGATTACCAAACCGCCTCTTTCTTCATTATTTCTCCTGAAGCATGACGATCACGG
ACCTGCATGCGGCAAATCTATAGAGAACTCAGCTGCCAGTCTCTCCCACTTGACGATCACGG
ACCTGCATGCGGGCACTCAGCAGTGAAAGGGTTAGGCCTAGGCTTTTTGACGATCACGG
ACCTGCATGCGGCAAACAGACCAGTGCTTGTATCAGCCCTTAAACATCTCTTGACGATCACGG
ACCTGCATGCGGTTTTGACAGTTGGGGATTAGGTGTTCTGTATCTGGGGTGACGATCACGG
ACCTGCATGCGGACTCAAGCAAAATCTAGACTATGTGTCCCCAAGATCAGAGGTGACGATCACGG
ACCTGCATGCGGGGGGCACATTCTTTAAAATTGGTCCCTGATCTAGACTCTTGACGATCACGG
ACCTGCATGCGGTTTAAGTTCAGAATTTCCACTCATGACCTCACATCTGTGGGTGACGATCACGG
ACCTGCATGCGGAGTCAAGGCAACCTCTTCACTCTTCTGCCAAGCCTTGACGATCACGG
ACCTGCATGCGGGCCAGCCCTGCTCAGTCTTCTTGCTTGTCATCCTTTGACGATCACGG
ACCTGCATGCGGCTAGGCCTTGCATTGCCATAGCCCTCTGCCTCTGTTGACGATCACGG
ACCTGCATGCGGTCACGCTCTCTCATCTTGGAGCATGAGCCTTCCATTGACGATCACGG
ACCTGCATGCGGCTCCTTCTGGAGGGTGGGCACAGCATCCATTCATTTGACGATCACGG
ACCTGCATGCGGCTGGGAATCCTGGTCGGCACCATGCTAGAACTTCTTGACGATCACGG
ACCTGCATGCGGCTTTGGTGCTGGCCCATGGGAGAGCTGTTGGTAAGTGACGATCACGG
ACCTGCATGCGGGCATACTTTTGCAGATTCCAGTTGCTGCTGAGGTTTGACGATCACGG
ACCTGCATGCGGGTTGCTCTTTGCACAAGTTTCTTCTAGTCACCAGTGAAGTGACGATCACGG
ACCTGCATGCGGTCTTTCATCAAGTTAGGGTACCAACCTTTAAAGACTGCCAGTGACGATCACGG
ACCTGCATGCGGATGTAAAAGGCACTGAGCCAGTGCAGCACATCTATTGACGATCACGG
ACCTGCATGCGGGACCAGGAAGGGTCAGCTTCCTGCCTTGCATGAGATGACGATCACGG
ACCTGCATGCGGATATGTATTGAGAAGTGGGTTCTCAGGAGATGATGCATCCATGACGATCACGG
ACCTGCATGCGGCAGCATTGTTTGTATGCCTCTGTCTTTGATGTCCCTGTGACGATCACGG
ACCTGCATGCGGCCTGAGTCGCCCACTTTAGAGCCCTTCTGTTCTTCTGACGATCACGG
ACCTGCATGCGGCAGTGATAATAATCATGAGTGAGCCCCTGCCCGTGTGACGATCACGG
ACCTGCATGCGGCTGGCTGTGTCCTGCTGAAGGCACACTAAGTGCTGTGACGATCACGG
ACCTGCATGCGGTCAGCATCTGTGAAGAGCTGGCCAGGCTTTTGCTCTGACGATCACGG
ACCTGCATGCGGGTTGACACAGTCAAGGAGCAAGGGCCCCGTAGGAGTGACGATCACGG
ACCTGCATGCGGGGGGCTGATGCTGGCTTCTGGAGCACTGTAATGTGTGACGATCACGG
ACCTGCATGCGGAAAAGAAGAAGGCAGGAGCTCGCACAGCTCTTGACTGACGATCACGG
ACCTGCATGCGGTTTTCCTGCTTCATCCAAGCAGGTCGACTCTCTCGTGACGATCACGG
ACCTGCATGCGGTGATCTCAGAGACAGAGTGAAGTCATGAGTGGGAGGTGACGATCACGG
ACCTGCATGCGGGGAGCACAGAAAATAAGACCTTGATTCCCAGCATTGGTGACGATCACGG
ACCTGCATGCGGAGGAGAGATTTGTCTTTCTCAGGTCAGGAAGAAAATGCTGTGACGATCACGG
ACCTGCATGCGGCCCGCTGCACATTCTTCGGGACAGACTCTTTTAATTGACGATCACGG
ACCTGCATGCGGTGTTTTGCTTAGTTAAGGAAAACCCCTGTGGTTTCTTGACGTGACGATCACGG
ACCTGCATGCGGTGCTTCAGTATTCTAACTCACAGCTGATTCAGTTCAGGGTGACGATCACGG
ACCTGCATGCGGCTGGGGAGATGTCCTCGACCTCTGGAAAGGAGGGTTGACGATCACGG
ACCTGCATGCGGTCTCTAGAAATAAGGCTAAGTATGCCACTGACACTGTCTGCTGACGATCACGG
ACCTGCATGCGGATAAACGTGTGTGATCTCAGGTCCAAAGGATGGGGTGACGATCACGG
ACCTGCATGCGGCTGTGGCAACTTGTGAAGACCATTCTGTGACCTTGTGACGATCACGG
ACCTGCATGCGGGTGTCTCTGGGCCTTCTCTTAGATTTTCTAAGTTGGCTTGACGATCACGG
ACCTGCATGCGGCAGTGGAGCTGCCATCCCTCCTTTGCCCATGTTCTATGACGATCACGG

ACCTGCATGCGGCTCCCAGAGTTCCTCCAAGAAATTGCGGAGCAATGTGACGATCACGG
ACCTGCATGCGGCCTGTTTCATGAGAGCTGAGTTTGCTGTGTCTTCCTGACGATCACGG
ACCTGCATGCGGACTTAGAAACAACACTGTGGACCAGGAGGACACACTGACGATCACGG
ACCTGCATGCGGCAGGAACCTCAGCAGGGCAGGGTGTTTCCTAGGAGTGACGATCACGG
ACCTGCATGCGGGACATCCGATTCCCAGCCATTCCTTTCAGTGAATCTGACGATCACGG
ACCTGCATGCGGTTTTGAAGCTCTTAGCCAATCTGCTTCGCGATGAATGACGATCACGG
ACCTGCATGCGGCCAGTTTTGCTTGAAGCAGACAAACCCGATTGTCATGACGATCACGG
ACCTGCATGCGGTCTTCAGTCTCTGAGGAAGAGTTTTCATTTTCCCCAATTCGTGACGATCACGG
ACCTGCATGCGGAATATTTTCCATGTGTGTTAACAATGGCTGAGCGTGGTTGACGATCACGG
ACCTGCATGCGGAGATGCCAGGAATTTCTGTCAACCCTCAAAGAGGATGACGATCACGG
ACCTGCATGCGGCTTATACCTGGCACTGCACGCTTCGCTCCCTCTGATGACGATCACGG
ACCTGCATGCGGTCTCCTGACTGTCATGCCCAGTGTCTCAGCCTATCTGACGATCACGG
ACCTGCATGCGGATTCTACCTCTAACTCGACCTTGAGTGACCTTGAGCATGACGATCACGG
ACCTGCATGCGGAGTTTCTCAGGATTCCACCTCCAAGTCACTCTCCCTGACGATCACGG
ACCTGCATGCGGTTTGGGATATGCAGCACTAAGTTAAGCTTGCCTGGTGACGATCACGG
ACCTGCATGCGGAAAACATCACTTGAAGCTGGAAAACCACTTTTAACACAGCGTGACGATCACGG
ACCTGCATGCGGGGAAAAGCTATTTGTTCAGACAGGAGTGGGGTGGGTGACGATCACGG
ACCTGCATGCGGGAGCTCAGAAAGAGGGGAACGCTCCCTCGTCTCTCTGACGATCACGG
ACCTGCATGCGGAACAGTTGCTCCAGACAGGTCAGCAAACATGGAATTGACGATCACGG
ACCTGCATGCGGTGTTCATTAAACACTGGCTGTGTCTTTTGTGTTCAAAAGCATGACGATCACGG
ACCTGCATGCGGAGACACTCTCTCTGAACCATGGCCCCACAGAGAGTTGACGATCACGG
ACCTGCATGCGGGCAGAATGTGTGAAACCTGCCGGGAAGGTCTGGACTGACGATCACGG
ACCTGCATGCGGCGTTCACACCACTCACATGGCTGTGCCTCTGCTTCTGACGATCACGG
ACCTGCATGCGGCTTCTGGCATGGCTGCTTCTTCCTCAGGTCTCAACTGACGATCACGG
ACCTGCATGCGGTTGCGCACCCATGTTTGCATTTGCAGGCCCTAGAATGACGATCACGG
ACCTGCATGCGGCCTCCTGTTCACAGCGGCACCTGAGTCACACATCTTGACGATCACGG
ACCTGCATGCGGGGGGGTTTCCCTGACTTCATCCATCCTGTCTTTTGGTGACGATCACGG
ACCTGCATGCGGGTCCCCATAATAACTGACATGGGTCGGCCCGTACCTGACGATCACGG
ACCTGCATGCGGCCCCTGCTCATCTTAGAGTCTCTCTATAGTGCTGCTTGACGATCACGG
ACCTGCATGCGGTTCTAACGTGCAGATCTGGTCATGTTACCAGCCTTTGACGATCACGG
ACCTGCATGCGGTTGGCTTTCTCTATCTAAAGTTCAAAACCGAACATGTGCGCTGACGATCACGG
ACCTGCATGCGGGCCCACCAGCAAGATTTATGAACCATTTTCTGCTGTTGACGATCACGG
ACCTGCATGCGGATGAGATGTGAGTCCTTCTACGAACTGACTCACACTGATGACGATCACGG
ACCTGCATGCGGTTGCCCAACTTCCTCTCTCGAGGTCTCATCCTCTTTGACGATCACGG
ACCTGCATGCGGCAGGGTCGATGCCATCTACCCTGGACTTCATCTTGTGACGATCACGG
ACCTGCATGCGGAACTCCTTCGAGTGTGAGTCATTACTCCTTTGTGCATGACGATCACGG
ACCTGCATGCGGCCTCTGCTTTCTCTTCTCAAGATGTTCACCTGCTTGATGACGATCACGG
ACCTGCATGCGGCACCACAGTGCTCAACATGCCTGAATGCATGGATGTGACGATCACGG
ACCTGCATGCGGATTCGTTCCACAACACTTCACCAATTAAAGAGAGGGTACATGACGATCACGG
ACCTGCATGCGGAAAGGTGAACATCCTTGGCTCCCAGCAGATGCTCCTGACGATCACGG
ACCTGCATGCGGTCAGATAGGTGAGGGAAGATTGAATGAAAGGCCTCTTATGATGACGATCACGG
ACCTGCATGCGGTTGAAGTAAAATGGATAGTGAGCCTCTGTGTATGTGAAGGCTGACGATCACGG
ACCTGCATGCGGCCGCATCTGGAACATGAAAGAACCTGTCTGATGTGTGACGATCACGG
ACCTGCATGCGGGAACATGATAGAATTTTACTCATTTGCCAACTGCGGTTCCCTGACGATCACGG
ACCTGCATGCGGATTTCAGGTTCAACCATAGGTCAGAAGTACACGTGTTTTGTTGACGATCACGG
ACCTGCATGCGGTTGTCTCTCAGTCATCTCATGGAAAATTCTCAGCTTTTGGTTGACGATCACGG
ACCTGCATGCGGCTCCTTTCCAATCCATACCCGCAATCACCAGAAACTGACGATCACGG
ACCTGCATGCGGGTGGACTTCCCTGACACTGAGCACCTCTTAATTAAGCTGACGATCACGG
ACCTGCATGCGGATCTCATAAGTGAACAAAACCCAGCCCTTCAAAGAAGTCATGACGATCACGG
ACCTGCATGCGGACTCTTCCACAAACACTTCAAGAAACCTAAGACTTTTGGCCTGACGATCACGG
ACCTGCATGCGGAGAGTAACACCGAGGTTTGAGAGAAAGGATATGTGTGTGATGACGATCACGG

ACCTGCATGCGGATAAAATGGGATTGAGAGGGGAATTTTGAAGTTCCCATGGTTGACGATCACGG
ACCTGCATGCGGCTGTCCTCTACATTCTGACAGCTCATTATCTCTGCGGTGACGATCACGG
ACCTGCATGCGGTATTGTTCTCACATTTAAGTGAGGTTAGCGGAGGCAGAGTGACGATCACGG
ACCTGCATGCGGGCCTCTCAGGCCTGAAGATAGCCTCTGTTTTCAGGTGACGATCACGG
ACCTGCATGCGGACTAGACTGTGAGATCTGTGACACTGAAGCACTAAGTTCATGACGATCACGG
ACCTGCATGCGGTTTTCTGTTGGACCTGGTGTTGGGGTTTCACACTTTGACGATCACGG
ACCTGCATGCGGGTGAATGTGGGCCTCCTGTGAGCACCAGCACAAAATGACGATCACGG
ACCTGCATGCGGAAACACACCAGGAAGCAGATCTTAGATCTCTGTTATCTCCTTGACGATCACGG
ACCTGCATGCGGTGGCACCAGCTGGTATTCATCCTCAATGCTAGCTATGACGATCACGG
ACCTGCATGCGGATCAATCAGTCTTGCAGGATTTCTTATGCGTTACATCGTCCTGACGATCACGG
ACCTGCATGCGGTTATAAGAAAGCAGTCATTCTCACCGAGATGTGCTGAGCTGACGATCACGG
ACCTGCATGCGGAGATACTGGACATGTTCTGACCCAGATAAGGGCTGGTGACGATCACGG
ACCTGCATGCGGCAACTTCCAGGACCTCGGAAGAACTGAAGGCAGAGTGACGATCACGG
ACCTGCATGCGGAACCTACAGACGATTCTGAGAAAGGCTCACAGGGATGACGATCACGG
ACCTGCATGCGGTTATTTCTGTCACTCTGCATTTTCTGACCTTTCCCAAGTGTTGACGATCACGG
ACCTGCATGCGGTCTAGTTTGTTCAGCTGTTCTTACTGGGTGAATTTGTGTGGTGACGATCACGG
ACCTGCATGCGGTCAGTATGGCCAAAGTGAGTTTCTCATTATTTTACCTCCCCTGACGATCACGG
ACCTGCATGCGGTCAGGAAATGACTTTTCATCTTGTGTTTTGGGAGCCATAGATGACGATCACGG
ACCTGCATGCGGTGGTTCTGGGCAGGAAACTGGCTTTGGATAGACCCTGACGATCACGG
ACCTGCATGCGGAGCATGTAGATGGCTATTTGGCCTTGCTCCCAGTATGACGATCACGG
ACCTGCATGCGGTTTGGGGCTCTGGATACCGTGTGGCCTGAAGAGACTGACGATCACGG
ACCTGCATGCGGAAGGGCTCAATGCCAACTCTGCCTGTTTCCAACTGTGACGATCACGG
ACCTGCATGCGGAAAATCATGGACGTGCTCTGGTTAACACTGAGTGGGTGACGATCACGG
ACCTGCATGCGGTGTTACTTGGACATGGCCAAGTTGACTACACTTTATGTTCTTGACGATCACGG
ACCTGCATGCGGGCTACCTGCCAGTCTGAAAGTGACGCCACAGAAGGTGACGATCACGG
ACCTGCATGCGGCAAACACAGCCCATGCGCCTGCTCTTCATGACTGTTGACGATCACGG
ACCTGCATGCGGATCTGTACCAGCAATATTTGTATTGGCAAATCACATGCCCTGACGATCACGG
ACCTGCATGCGGTATGTAAGATATGTGTTGCAATTCAGCTTTCAGGTCCCAGCTGACGATCACGG
ACCTGCATGCGGGCCTTCACAAACTGGCCCACCCACGAGAAGGAAAGTGACGATCACGG
ACCTGCATGCGGCAATTGTCCAAATGTGGGTAGCTTTTCTTCCCACTGTTGACGATCACGG
ACCTGCATGCGGATCCCAGTTTGTGTCTGTATCAGTACAATTCTCCCATGTCATGACGATCACGG
ACCTGCATGCGGATGTGAATTTTAAGCCACAGAGGGAAAGGGGACAGATGACGATCACGG
ACCTGCATGCGGGAATATGCTTTCATTCAGCTCTCCTCGTCTCACACCTTGACGATCACGG
ACCTGCATGCGGCTTGCCCTGCATGCATTTCTTTGCTCTGATTAAACGTGACGATCACGG
ACCTGCATGCGGTTTTGGAACCGAGACAAATGGAACTTAGCTAAAAGATGGGATGACGATCACGG
ACCTGCATGCGGTTTTGCAGCACCCAGAAGCAAGCCTGTTTCAGCACTGACGATCACGG
ACCTGCATGCGGGGCAAAGGCTCAGCTGCTAAGTGGGCAGCATTGTTTGACGATCACGG
ACCTGCATGCGGGGAGGTGAGCAGCTTAGGCTGACTGTTCATCAAAGTGACGATCACGG
ACCTGCATGCGGGGCCATGGAACTCTACTGTCAGTTTCAGGGAAATTTGGTGACGATCACGG
ACCTGCATGCGGGACCCTGGAGCACAAACCAAAACTCCAATTAACCATGACGATCACGG
ACCTGCATGCGGGGAGAGGAACTCGATCCCCAGGAGATAAGTGAAGATGACGATCACGG
ACCTGCATGCGGAGGAGGGTGGAACTGTTCTGGAAGTGGGTCTCAATTGACGATCACGG
ACCTGCATGCGGCTCAGCACCAGCAGCTCTCAAGACTTTCTAGAGAATGACGATCACGG
ACCTGCATGCGGTTAAAATTAGTCTTCAATGACATGGCAGGGATTTCGGCACATGACGATCACGG
ACCTGCATGCGGCTCTCTTGCGTCATAGGCCACTGTGTTGGAGGCAGTGACGATCACGG
ACCTGCATGCGGTAACACGTGAGCACTGAAAAGGTTAAACAGTAGAGACATGGTGACGATCACGG
ACCTGCATGCGGTACCTTAGTGCTTCTGTGAGCATGTTTTCTCTTTCATTGCTTGACGATCACGG
ACCTGCATGCGGTGTGCCATTTGGATCATAACTGATAAGCAACCAAAGAGTCCTGACGATCACGG
ACCTGCATGCGGATATTACTGCACGTTCCCATCGCTATTTTATGTGAAGGTGGTGACGATCACGG
ACCTGCATGCGGCTTGTGGCATCTTTGAAAAGCTACTTAAAATGTGCTGCTGTTGACGATCACGG
ACCTGCATGCGGTTGAAAATGTGCTTTTAATACAAAGTTTGTGCAGCCCTTGCTGACGATCACGG

ACCTGCATGCGGCATTCTCTTCACTGAGGGTCAAATGGAAAGAACACATGGGTGACGATCACGG
ACCTGCATGCGGATAAGAGAAATGAAGAAATATGTGCCCGGAAGCAAGAACGATGACGATCACGG
ACCTGCATGCGGCCGACCTCATTAAACTGGCTCCCTTCACCTCCTCTTGACGATCACGG
ACCTGCATGCGGACTTTGGTCCATAATTTAGATCGAACCACATGTGCTGATGTTGACGATCACGG
ACCTGCATGCGGACGATGTGGAATTCTCTGGACAGAGATAGAATTATGGAGGGTGACGATCACGG
ACCTGCATGCGGAAAAGTTCTCTTCCCATGAGGTTCTCGCAAAGCATTTTTGACGATCACGG
ACCTGCATGCGGTTTTCTCAGTTTTCTGAGATGGGGCCATCTTGAATCCATGACGATCACGG
ACCTGCATGCGGACAGACAACACACAGCATCAGCCAGACTAACACAATGACGATCACGG
ACCTGCATGCGGAGGACGTCATGGGCATGGACGTAAATACTGGTGTCTGACGATCACGG
ACCTGCATGCGGAACACTAGGTCTGCACCTCGAGAGGAGTGGAGCAATGACGATCACGG
ACCTGCATGCGGAAAGGAGGCAGAAATGAAAGGAAGACCATCAGTGCGTGACGATCACGG
ACCTGCATGCGGGTCCTGGAGGGCAAATGCCAGCTGCTCACTTCTGAAAATGACGATCACGG
ACCTGCATGCGGACTTGATACTGTGTATGTCTCTTCTTGGCTTCTCATCTCCATGACGATCACGG
ACCTGCATGCGGATAATTGTGTCGTGTTCTTCCCCACTAGAAGCCAATTATGCTGACGATCACGG
ACCTGCATGCGGTGGAGTACTCATTGCAATTTCACTTATCCAGAATTGGCTGTTGACGATCACGG
ACCTGCATGCGGTGTTCTCAGAGCAGCTTGTGTTGCCTTGTTAAGGATGACGATCACGG
ACCTGCATGCGGATCCAGACATCCAGAAAGGATCCTTTACTGTTTCAGAGTCCTGACGATCACGG
ACCTGCATGCGGATGGTAAGTAGACACAAACCTACTGGCTCCATCATTTCTGTTGACGATCACGG
ACCTGCATGCGGTAGCAAATAACCTGCCACATATACCAATAGCCCAAGAGATGTGACGATCACGG
ACCTGCATGCGGGGCATGTCCCTGCATTTCCTGGTCAAGGTGACAACTGACGATCACGG
ACCTGCATGCGGGCGTCCTCCTGGAAGAGGTCTGCCACTCACCATACTGACGATCACGG
ACCTGCATGCGGAAGCATTCTCGAAGCTTCCAGAAGTCAACTCAAGTTATCTGTGACGATCACGG
ACCTGCATGCGGAAAGTGACACTTTTGATGATTGCTCGCTTAATACTGGGAGATGACGATCACGG
ACCTGCATGCGGGCAACTCTGGAATTTCTTAGTGTTACTGGAGATTCCTGCTTGACGATCACGG
ACCTGCATGCGGACACTGAGATGTTCTAAGGAAATTACTCCCTAGGGAGGAGATGACGATCACGG
ACCTGCATGCGGGGGGTGGACGAGGAGTAAGCTTTGCTGGTGACTCATGACGATCACGG
ACCTGCATGCGGAAAAGGTTCTGGTTGCAGAAGGAATAAGAAATGGCTTGAGATGACGATCACGG
ACCTGCATGCGGTGTTGTAAACATGTTAGCTATGGCTGTGAATTCAACCAGCGTGACGATCACGG
ACCTGCATGCGGGGGGAATTAGGGACATGGCTGTGTTTCCCCAGAGATGACGATCACGG
ACCTGCATGCGGAGTGGCCATTTTACTTTCCCTCTTCACTAACATGCTTTTGATGACGATCACGG
ACCTGCATGCGGCCAAAACTATTCTCTACGGCTGCATTCCTTTCGCTCTGACGATCACGG
ACCTGCATGCGGTTGCCTTCCTTTAGAACCCTGGAGAAGGCCTCCTGTGACGATCACGG
ACCTGCATGCGGTTTTCCTCGCAGTGACTACGTGATGAAAGTACCATGCTGACGATCACGG
ACCTGCATGCGGGAGGAGGTGTCTGACTGAGGCGTTCGTGGTGTGTGTGACGATCACGG
ACCTGCATGCGGGCACTCCCCTCTTGCGTCCTTTCCTCCCATGTTTGTGACGATCACGG
ACCTGCATGCGGGGGGTGTTCTGGCAGTTGACATTTCTGAAAACTACAGTGACGATCACGG
ACCTGCATGCGGACCGTGGTCATAATAGTGTGGCATTTGATAACTAATGAGGCTGACGATCACGG
ACCTGCATGCGGGACAGGGCAAGGGAGCTTGGGTTGTAGCCTCGAATTGACGATCACGG
ACCTGCATGCGGAGGGCTACCGTCAGCCTGCGGCACACAAGTAAATCAAATATTGACGATCACGG
ACCTGCATGCGGTAACTCACTGTAACGAATTATCTTTCGCACATCACAGAGGCTGACGATCACGG
ACCTGCATGCGGAAAAGGGCAGTTCAAAAGCTTTGGTTGCTATTGTGAAAGTTGACGATCACGG
ACCTGCATGCGGGTGGGGTGGGGTTCACGCAGGTTCTCTTTTGTCACTGACGATCACGG
ACCTGCATGCGGCAGGGGTGCTGTGGATTCACAAGTAAGCAAGAGGCTGACGATCACGG
ACCTGCATGCGGCTCCCTGAGGTCAGCTGCCTAGCTTCTCCTCCTCTTGACGATCACGG
ACCTGCATGCGGTCCAAAACGTAAAAGTAAACCATGCATCCACAGAGACATGGTGACGATCACGG
ACCTGCATGCGGAATTACAGAACTGGATGCTGAGCTGGTCACTTGGGTGACGATCACGG
ACCTGCATGCGGGTCCTTGCCATTGGTTTATGCCTCAGCCCCACCATTGACGATCACGG
ACCTGCATGCGGCTCCTCACCTGCCTGGTGGGACAGTGACATCTCTCTGACGATCACGG
ACCTGCATGCGGGTCAGGGTGAGGGAGGTAGGGCGCAGTTTCAGAAATGACGATCACGG
ACCTGCATGCGGCTGCGTGAGCACAGCTGCAGGAGCAAAAGTCAGAATGACGATCACGG
ACCTGCATGCGGAGGTCAGCAAAGGATTTCAGGAGCAAAGGTCAGAATGACGATCACGG

ACCTGCATGCGGCTGTCGTCTCCTGCAATGCTTTCAAGACTATTCAGAAGCTGACGATCACGG
ACCTGCATGCGGCCGGAGCCCATGGGGAAATGACTCCAGAGTGTTCCTGACGATCACGG
ACCTGCATGCGGACGTGTTGGAAGGCATCTGTTGGAAAACGGACATTTGACGATCACGG
ACCTGCATGCGGCAAGCAAATAGTTGCCTGCATAGACAACGCAGAATGATGACGATCACGG
ACCTGCATGCGGCTGGGAAAGCCCCAACAAGTTACCTACTGGTAAATGATGACGATCACGG
ACCTGCATGCGGGGTGAGAAGCTTAAAGTGAGAACCCCATTGCTGCCTGACGATCACGG
ACCTGCATGCGGAGAATACAAAGAGGAGATTTTAAAATGTCAGTGGCAGCCCCTGACGATCACGG
ACCTGCATGCGGTTTCCTTGACTATGGTTATTTCTGGTAGTAGCAGCTCCAGATGACGATCACGG
ACCTGCATGCGGTTAGGCAATGGTTTTCTTCAGAGATAGCTTAGAGTGAGCCCTGACGATCACGG
ACCTGCATGCGGCAGAACAAGGTCAATGCGAAGATTGCTTGTGTCTGCTGACGATCACGG
ACCTGCATGCGGGTGTCCAGGGCACAGTGATCCTCATCACTAGCCGGTGACGATCACGG
ACCTGCATGCGGGGCTCCGTGAGGATCTGCTCCTGGTCGTTTCTGTTTGACGATCACGG
ACCTGCATGCGGCTGTATCTTCTCTGCAGCCCTTACTGAAGCCGTTATGACGATCACGG
ACCTGCATGCGGACAGATGGCTGAAGTATTGAGAACGCTCCAGTGACTGACGATCACGG
ACCTGCATGCGGCGGGAGGCAATAGTCTGTCCACATCTAAGAACACATGACGATCACGG
ACCTGCATGCGGAGGTTATCAAACTCCTATGACTTTTCACAGGAAAAGCCCTCTGACGATCACGG
ACCTGCATGCGGATCCACACCAACTTTAGGAATGTGTAGAAAGAAGGGTCAGTGACGATCACGG
ACCTGCATGCGGATTTGGAGTAGGAGGTCTTGCTTTACTATTGAATTGCAGGGTGACGATCACGG
ACCTGCATGCGGACTCAAGTCTTCTTACAAAGCAGTTTGAGGCTTTGAGTACCTGACGATCACGG
ACCTGCATGCGGAAAAGGAATTTTATCGAGGTGGCCCTTGAGTGCCATGACGATCACGG
ACCTGCATGCGGTGTGCAGCAAGTATCTGCTGAGCTAATAATAAACAGCCTTGACGATCACGG
ACCTGCATGCGGCAGACAGAAAGACAGTGGGCACAAGGTCATGCTTATGACGATCACGG
ACCTGCATGCGGAAAAGACCCCTTGTTCTACTGCATCCCAGCTCCCCTGACGATCACGG
ACCTGCATGCGGCTCAGTCCTGGGAGCCACAGAACCCAGCACGTACTTTATGACGATCACGG
ACCTGCATGCGGCCCCAAGACCAGACTCCAGCTTGGCTTTTGTCCTCTGACGATCACGG
ACCTGCATGCGGCTCTCCAGGATTGGTGACCTCCTAGGTCGTGAAGCTGACGATCACGG
ACCTGCATGCGGTGTGATGAGCAAAGACACACTCCTCTCCATTCTCCTGACGATCACGG
ACCTGCATGCGGTTAAATAGCAGACCACCCACAGCAGGGCTGGTAGATGACGATCACGG
ACCTGCATGCGGTGCAGTGAACTCAGGAAGATGCCTGCATAGACTCTTGACGATCACGG
ACCTGCATGCGGCCCCATAGTTACCTAACTGCTGTCTCCAGTGGTCATGACGATCACGG
ACCTGCATGCGGAAGGAGAACAATGCTATATGCTGTATCCCACCTTTCTCTGATGACGATCACGG
ACCTGCATGCGGATGTTACATTTTCTCCCCTATCCCAGGCTGCATCTTGACGATCACGG
ACCTGCATGCGGATGCTATCTATCTTTTCCGAGCAATGAAAGCTCTGGGTTGACGATCACGG
ACCTGCATGCGGAAAATCACTAACTTTCATGCCTCAGCAAACCTCCACTGTGACGATCACGG
ACCTGCATGCGGAAAATATAGTGAGGTCATTCATCTTCGGACAAATTGCCCCATGACGATCACGG
ACCTGCATGCGGGCAGCAGCAGCCTGTGAAACATTGACAAGACCTGGTGACGATCACGG
ACCTGCATGCGGAGTTGGAAGAGGACTTTGCCATCCTCCAGTCCAACTGACGATCACGG
ACCTGCATGCGGAGTTGCCTGTCACAGATTAGACGACTGGGATGTGCTGACGATCACGG
ACCTGCATGCGGCCCCAGTTTATGTCTTAATTCGCAGCCAGGGCTGATGACGATCACGG
ACCTGCATGCGGAAGCAAATTTGCAAACATGTGCAAGAAGAAATCACACATCCTGACGATCACGG
ACCTGCATGCGGGAACCATTGCTGAGCTGTTAAATTTGTCTCCTTCCCCTGACGATCACGG
ACCTGCATGCGGAAAGCCGGCAGATCCTAGGAGAATCTTATTTAATCCAAGCTGACGATCACGG
ACCTGCATGCGGAGTTTTGCTTTATTCCATGGCAACATGGGTATACACATCCCTGACGATCACGG
ACCTGCATGCGGGGCTGTTTCAGTGGCTCAGAGCAGGTAAGGCCTGTTGACGATCACGG
ACCTGCATGCGGCTAGCAGGAGGAACAACGTGGAGACAGCCCCAGAGTGACGATCACGG
ACCTGCATGCGGCCTTCTGTGGCTCCGGTGTCTCAGGACCTCCCTAAATGACGATCACGG
ACCTGCATGCGGCCCAGCCCTGACACTGAGCAAGTTTCCACCACTGTTATGACGATCACGG
ACCTGCATGCGGCCCCATTCCCATTCCATTAATTGCGTGAGGTAGAGATGACGATCACGG
ACCTGCATGCGGATTTTCACGTGAAGACTTTAGGGGCAAAAGAAAAGAAGCAATGACGATCACGG
ACCTGCATGCGGTTTTGAAAAGAGGACTAAAACAGATGGCCTGGCTGTGTGACGATCACGG
ACCTGCATGCGGTTAAACACAGGGACCAGACCAGCACCCACCTCTCCTGACGATCACGG

ACCTGCATGCGGACCTGCCCTGCCTTCACTGGCAGAATTGTGATCCATTGACGATCACGG
ACCTGCATGCGGCATGTTCTCTGTTCAATGTCATCATCCCTTTCAGAGCATGGTGACGATCACGG
ACCTGCATGCGGGTCTCTTCCTTTCTAGGCAGTCTTACCAGGATGCATGTGACGATCACGG
ACCTGCATGCGGCTCTTTAAAGACAATTAACACAGTTTCTGGGCAAGGCAAGGTGACGATCACGG
ACCTGCATGCGGTCAGTAACTTAGCAACACCAACAGAAGACAGCCAATATTGCTGACGATCACGG
ACCTGCATGCGGTTTTCTCTTGGATTGGGTCAGAGAGTACTGCAGAGAAAATGTGACGATCACGG
ACCTGCATGCGGGAGTAGAGAGACCTGAAATACTTTCGCACACACTGTGGTGACGATCACGG
ACCTGCATGCGGTCAGTGCAGCGTCCACTGTGTGCCACAGTAATACTTGACGATCACGG
ACCTGCATGCGGAGAAACTCCCTGGTTAGGCCTTGGAATCCAGCTCTTGACGATCACGG
ACCTGCATGCGGCATTTCGTATGTGACCTGCAGGGAAGTAAGTTAAATGCACATGACGATCACGG
ACCTGCATGCGGTCAATCATTGCTCTCATGAGTGGTGATGCTGATGCTGACGATCACGG
ACCTGCATGCGGTGATTTTCCCTTACAAAGTTACCAACAGTTTCATGTTGGCCTGACGATCACGG
ACCTGCATGCGGAATTGTTTCCACTAATAGGACCAACAGTGGTAGTCCCATCATGACGATCACGG
ACCTGCATGCGGTTTTATTACTGCTTGTCGTAGCACAAGCAGTTGCTTCATTGTGACGATCACGG
ACCTGCATGCGGAAATATTGACGGCTGCTTTTAACAGTCTGCTGTTTTGTCTCTGACGATCACGG
ACCTGCATGCGGAAAAGATAGTGCAGATGGAAAGATGTCTGGAGTCAGTGAACTGACGATCACGG
ACCTGCATGCGGTAAGGAATTCTCAGGCTTTGCTATATCTGACTGCTCTTGGATGACGATCACGG
ACCTGCATGCGGGGCTGAGCTTTTGGCTAACTACCTGACTACTTTGTCGTGACGATCACGG
ACCTGCATGCGGAAATACTGAAAGACTTCCCTCAATTCAAACTATGTGCCGGATGACGATCACGG
ACCTGCATGCGGTCAAGGAAAGGGCAGTTGGATATTGCAGACAGCATTGACGATCACGG
ACCTGCATGCGGTTCTCAGAAGACTTTCAGGCAGTATGTGGGTGAGGTGACGATCACGG
ACCTGCATGCGGCACATGCTGGAAAGGCTTCTGCAGGTGCAGTGATCTGACGATCACGG
ACCTGCATGCGGTTCTCAGTGTGTACATCCCATAATACAGACACGTTACCAGATGACGATCACGG
ACCTGCATGCGGTGGGCATTTATAGCTATTAATACTTCGTCCCATGTGTCCCATGACGATCACGG
ACCTGCATGCGGTTACATAGAGACATTCACAGCTCTTCTGACCTTATCAGCGTTGACGATCACGG
ACCTGCATGCGGTAAGGAAAACAGAAAACCAGCGTGCTATTTGTTCTGTCCCTGACGATCACGG
ACCTGCATGCGGTTTTCTTCCAAGATTTTGCATGTGCACAGGGATGCCTGACGATCACGG
ACCTGCATGCGGAATGATCATTTCTTAGATTGCATCTCAGACACACCCTTCCCTGACGATCACGG
ACCTGCATGCGGGCCCATGGGTACTGCAAACAGCCTGACGTTGTCAGTGACGATCACGG
ACCTGCATGCGGGTCTCAACGGCTCATTCACCACATCTGCCTCGCGAATGACGATCACGG
ACCTGCATGCGGAGGTACACGCTATTAACCTTGTGAGAAAACAAAGAGGCCTGACGATCACGG
ACCTGCATGCGGCCCACCCTTCCTGCTCACTCTGAGTCACGGTGAAATGACGATCACGG
ACCTGCATGCGGATGTTTCAGGATCTCGGGTTCGACCATGAGTCCTGTGACGATCACGG
ACCTGCATGCGGTCCAGGTCCAGGAGGAAATTCGGAAGGACCACATGTGACGATCACGG
ACCTGCATGCGGTTCACTCTGAGATCCCACTTTCATTTCCCTCCTGGTGACGATCACGG
ACCTGCATGCGGGACAATGTAATCAAAATAGAATGTGGGTTTACAGCTGGCCCTGACGATCACGG
ACCTGCATGCGGAAGTGGTGACATCATATTCTCTCATTCAATGTGAAAGGCCATGACGATCACGG
ACCTGCATGCGGAATGCTTAACCTGTCATTCTAATTACCAAGCACGGTGTTCTTGACGATCACGG
ACCTGCATGCGGAAGAGCATTTGTAAGTAACTGAATAGAAGGCCAAAGGGTGTTGACGATCACGG
ACCTGCATGCGGCAGGGGCCACGGTTGCAGCCTGCATTCTTCTAAAGTGACGATCACGG
ACCTGCATGCGGTGTTTCAAAGAATTCCACCAACCAGAGGATACTACCTAGGATGACGATCACGG
ACCTGCATGCGGCTTCCTCCACACCGGTGGCCACTTTTCATTCACTCTGACGATCACGG
ACCTGCATGCGGAGAATATATTCACTGGGGAATGGACTTAGCAACCACTACCATGACGATCACGG
ACCTGCATGCGGGTACAGCTATGTTCCTCCTGAAACTTGGAGACACATCCTGACGATCACGG
ACCTGCATGCGGTCTTGAGCTGGGTTATAATGGGCCACCCAAAGCTCTGACGATCACGG
ACCTGCATGCGGGAGTTCCTGTAATGGATACACTCAGGCAGCAGAACCTGACGATCACGG
ACCTGCATGCGGCATCACAAGTGCACCCACAGCTGCCTTCTCTGGCACTGACGATCACGG
ACCTGCATGCGGGCTCAGAGCTACACAGTGTACTCTGGGATTGGAACTGACGATCACGG
ACCTGCATGCGGAAATCCATGCACATCAACTCTCCAAATCAGAATTTGCTGAGTGACGATCACGG
ACCTGCATGCGGAAAAGAGCATTAAATTAGATGGGCTGGCTTTCAAGGGGTGACGATCACGG
ACCTGCATGCGGTACAAAGAGACAAGCAAGCACATCGCGTGGAAATTTGACGATCACGG

ACCTGCATGCGGTCCATTCAACTGGAAATGTCCAAGCCTGTTTACCTCATGACGATCACGG
ACCTGCATGCGGATTAATTGTCCTTGTTCACTTGTCCAGCCTAGCAATTGTCCTGACGATCACGG
ACCTGCATGCGGAGCTTGAGATGTATTAATACGAAATCCAAGCTGCATGAGGGTGACGATCACGG
ACCTGCATGCGGAAACCCAAGAAAGCCTTGTAACTTCCTTCAATCTAATGGGATGACGATCACGG
ACCTGCATGCGGACTGATGTCCTTGAGTTTTATTATCAAGACTCAAGGGCACCTGACGATCACGG
ACCTGCATGCGGTAGGACACTCTGTGGCCTAGACTTAGAAACACAACTAATGTTGACGATCACGG
ACCTGCATGCGGAAACAACATACTCACACCCCTTCGCCTATTATCATCTAGGTTGACGATCACGG
ACCTGCATGCGGTTCAATGCTCATTGCAATGAAACCTACTTATTGTGCATGGCTGACGATCACGG
ACCTGCATGCGGCCCCACTGAGGAATACTGTAGTTTCTTTCCCTTTGAACTTTGACGATCACGG
ACCTGCATGCGGTAGTAGAGCACATGGTTCATTCACTCCTGAAGAGTTCTTCGTGACGATCACGG
ACCTGCATGCGGTATACTACAACATAATTTCCATCAGAGCTCTGACCACCCGCTGACGATCACGG
ACCTGCATGCGGTCTATTTTCATAATGCCTGCCACTCCATCATTAGCTGTTGTTGACGATCACGG
ACCTGCATGCGGAATAAAACAGTTAGGGAATGAGGGAAATTGACTAGCAGCCATGACGATCACGG
ACCTGCATGCGGTAGACCCCAAAGATTCCAAAGATGGTATTTCGGTCAGATCTTGACGATCACGG
ACCTGCATGCGGATGGCAGTTTTATTGACTACTATTCTTTGCTGGGTGTGGTTGACGATCACGG
ACCTGCATGCGGTTTTAAACTGAGACATCAGTGTGCCTAGCACAGGGCTGACGATCACGG
ACCTGCATGCGGCCTTTTGCACACCCTGCCTCACACTTGCTGGTCTATGACGATCACGG
ACCTGCATGCGGAATGACAGAAATGTTTACTTCCAAGTGGAACTAAGCCAGGGTGACGATCACGG
ACCTGCATGCGGTAACTCAGGGTAGGGCAGCTGCTTGCACCGAAAGATGACGATCACGG
ACCTGCATGCGGCCAAGACTGCTAGAGAACTAGGAAACAGGCGGTGCTGACGATCACGG
ACCTGCATGCGGTATGGGGCTGCAGAAACTCTTTGGTGCTTGGGGAATGACGATCACGG
ACCTGCATGCGGAAAAGAAACCTGGGAGAGGTAGTTTCCAGATGCAGGTGACGATCACGG
ACCTGCATGCGGCCCGTCTTTTCTTTTAAACAGAGGCAGCTCCGAAGTGACGATCACGG
ACCTGCATGCGGAGCTGGACATTGAACCCTGAGCAGGAACTGGAGGCTGACGATCACGG
ACCTGCATGCGGGCTTTGTTTGGCGAGCGGAGCTTTGCAAGGGTGTATGACGATCACGG
ACCTGCATGCGGGCCCTCTTGGGGTAAGGTACGCCCAGGAACAGTTTTGACGATCACGG
ACCTGCATGCGGCCATCCCCAGGGTACCACGCCACGTAGAGACACTATGACGATCACGG
ACCTGCATGCGGTTTTCACTTCGTGTTTGTCACTCCTAAAGCATGTGTGCTGACGATCACGG
ACCTGCATGCGGCTTCCGCACGTGCAGGAAAATGTCCACGTGCACTTTGACGATCACGG
ACCTGCATGCGGCACGGGCACCACCACTGATCATCTGTGGGATCGAGTGACGATCACGG
ACCTGCATGCGGCAGTCGCTTTGGTGAGAGAGTGGACGCTGTGGTGATGACGATCACGG
ACCTGCATGCGGCTGTGCTCAGGAGGACAGAGAGGGGACATCCTGAGTGACGATCACGG
ACCTGCATGCGGGGATCCTGTGCATGTCCCCAGAGCGTCCACTTTCTTGACGATCACGG
ACCTGCATGCGGGGGCGGGAAAACTTACCACCAGGGGACTCGAGATGTGACGATCACGG
ACCTGCATGCGGGCTCTGTGCGTGCTGAAGGTGTGGTGAGAAGCACTTATGACGATCACGG
ACCTGCATGCGGTGGACTGTGTTAGGATTGCACATTTTACTTTGTTTCTCCCATGACGATCACGG
ACCTGCATGCGGCTGGCTTCATAGCTCTCATCATTCCCACGGAACCTTGACGATCACGG
ACCTGCATGCGGGTAGTCCGTGAAAATCCAGGGTCCCGCCACTCCAGTGACGATCACGG
ACCTGCATGCGGGTGTCTGCAGCAGAGCTGAACACACGTAGGCTCTTTGACGATCACGG
ACCTGCATGCGGAAAATAAATGTTATTCACTTGTCGTGGATGTGGTGGGGACATGACGATCACGG
ACCTGCATGCGGAGGGGTGATCTAGTGACACTAGCCATTTATCAGGACGTGACGATCACGG
ACCTGCATGCGGTATGGGTGCCAGTCAGGATGATAAAGCTTCCTTTTGGTGACGATCACGG
ACCTGCATGCGGCCACTATACTACTTAGAAATGCCCTGCAAAAGGTGCACATGACGATCACGG
ACCTGCATGCGGTTGAAAGCTCAATCCTGGATTTTAAGTGCTTCAAAAGTGCATGACGATCACGG
ACCTGCATGCGGAATGATTAACAGGGCAATTATCAATCATGGACTTTTGGCGGTGACGATCACGG
ACCTGCATGCGGCAGAAGGAACTGTATTGTTTGGTACAGTCTGGGCCATGACGATCACGG
ACCTGCATGCGGGGGCCACACACCGTAACGGAGATACTCTATTCTGTTGACGATCACGG
ACCTGCATGCGGTAGACTGTTCACACTGCTGCCACGAAGGAGTCTTGTGACGATCACGG
ACCTGCATGCGGTTTAGACTGGACCTGGCTTTCTTCTTCGCAATGAGTTGACGATCACGG
ACCTGCATGCGGGTTGCAGACTCCCGACAAAGGCCAGGTGGTAAAGTTGACGATCACGG
ACCTGCATGCGGGTGGTGTCTGTGAGCGAGAGCCTGAGATGCCTGAGTGACGATCACGG

ACCTGCATGCGGCCATCGTGCAGAGGTGAGAGCAGCCCCTGAATTCTTGACGATCACGG
ACCTGCATGCGGGCCCCTCGGTCTCTCCATAGCTAAAGCAAAACCATTGACGATCACGG
ACCTGCATGCGGGCGAGGAGGGGCCATCACTGCACAATTCATCAGTGTGACGATCACGG
ACCTGCATGCGGCTCTCTCCAAAACGTGCTGTCAGTGTGTTCTAATTTACTCTTGACGATCACGG
ACCTGCATGCGGTCCCAATCTAGGGCCTTCCTCTCACTGTACAAAGTTGACGATCACGG
ACCTGCATGCGGAGGTAACTGGGGACATTAGTGGATCAGTGATCAAACCTGACGATCACGG
ACCTGCATGCGGACCAAGTGATATCAGGATGAGAAAGCTGTTAGAGTGTCAGATGACGATCACGG
ACCTGCATGCGGTATGTGAAGGAACTTGGGTCATTCCTGATACCTCAAAGAGATGACGATCACGG
ACCTGCATGCGGAAAAGGTAGTCCTTGAACACCTCCTACTTGTAAAGGATGCATGACGATCACGG
ACCTGCATGCGGTCTCCTACCTGTGACTTTATAGCCTTTTGGAGACTCACATGACGATCACGG
ACCTGCATGCGGGCAATAGTTGTATTTAAACTCAGTGGGTGGCATCCAAGGTGACGATCACGG
ACCTGCATGCGGAAAAGGAGATTGCCTAGACACAAAACCACCCAAGGGTGACGATCACGG
ACCTGCATGCGGTCGCTCCAGTGTAGACATACAATAGACCACTCGTCCTGACGATCACGG
ACCTGCATGCGGGGCAGCAGCCTCATCTGAGACCCTCCTGAGACATCTGACGATCACGG
ACCTGCATGCGGTGCACTCGTTCTTGGAGACGTTGACTTTATTTAGTAGCATTTGACGATCACGG
ACCTGCATGCGGCATGGCAATGTCACCAGCATCGCTGACAGCCTGAATGACGATCACGG
ACCTGCATGCGGTGGGATTTTCTTCTTGACTCACTCCACTGTGGCCTTGACGATCACGG
ACCTGCATGCGGCCCTCATCCAGGACTGTAACAGACGCCTGACGTCATGACGATCACGG
ACCTGCATGCGGGTGGTCTAGACCTCTCTGCTGAATGTCATCTTTGGTGTGACGATCACGG
ACCTGCATGCGGGTCTTATGAGAAAACACATGGTTGGTCACTCTTAGAAGGGCTGACGATCACGG
ACCTGCATGCGGATGAAAGCCTGTCTGCAGTATAACCAAAACAGGCACTGACGATCACGG
ACCTGCATGCGGATGGCGAGGCACACTGTGCGCATGTGTGTACAATTTGACGATCACGG
ACCTGCATGCGGGGTTTTAAATTATTTTCAGGCCAAGGGGAGATCTTTGCTGCTGACGATCACGG
ACCTGCATGCGGAGCAAGATTGGGATCCTAAAATACAGACCCAAATGAATGGATGACGATCACGG
ACCTGCATGCGGTAACAGGGTCAAGGCCAAATCAGTAACAAATATCCTGAGTGTGACGATCACGG
ACCTGCATGCGGAGGTGGTTTAACAAATGCCCCTATGAAAGATAGAGATTGGCTGACGATCACGG
ACCTGCATGCGGTTACCATGATGAGATGTAAGCCCAAGTTATGAGGTTGGCTGACGATCACGG
ACCTGCATGCGGTGGACAAGCTTCTGTTGATTAGAAGAGTTCAGTTACAAGGCTGACGATCACGG
ACCTGCATGCGGTACACTTGGGAGGAATGTTTACAAACTGGAATGGTCAGAGTGACGATCACGG
ACCTGCATGCGGTTTGGCATATTGGATTTTGCCTTCAGAGAAGTGGAGTCCTGACGATCACGG
ACCTGCATGCGGGCTCACCTGCTGTCTTCTGTCCAGGTTCTCGGTAGTGACGATCACGG
ACCTGCATGCGGCAGGGCAGTACAGCCCCATCCGTGATCTTCCATAGTGACGATCACGG
ACCTGCATGCGGTCAGGCATATTGTCACACTCAGTGAGCGGAGAGTCTGACGATCACGG
ACCTGCATGCGGAACCGGGAGGAAGGCACAGTTTCTCTGGAATGACCTGACGATCACGG
ACCTGCATGCGGTACGGAATGGTACGCTCAAATGCAAATTCTCCTTCCCTGACGATCACGG
ACCTGCATGCGGTTCCCCAGTCCTTGTCCTTCAGATGGTAATTTAGGAGCTGACGATCACGG
ACCTGCATGCGGTCACATGAAGGATATTTGCATTGTTTTCCAACTGATGCCAGTGACGATCACGG
ACCTGCATGCGGCAGACACATTGTCAGTGGTATCATGCCTGGGGTATTGACGATCACGG
ACCTGCATGCGGCAGAGTTGACATTGGGTTGCCCCTTCTCTGAGGCATGACGATCACGG
ACCTGCATGCGGTCCTTTTAAGTTTATAAAACCTCCATGTGGCTCCTGCATGCTGACGATCACGG
ACCTGCATGCGGCGCCCTCTTGTAGAGCCCGTTCCGTTGTCCATGAGTGACGATCACGG
ACCTGCATGCGGAAACCTAACAATGTAGAAAGACTAAAGCACATGGGTGGTGCTGACGATCACGG
ACCTGCATGCGGTATTTCTCCCACTTTCTGAAAACTCCTGCTGAAGTAACTGCTGACGATCACGG
ACCTGCATGCGGTGGAATTCTGCTAACGAAAATTGCGAAGGATCAAGGCTGACGATCACGG
ACCTGCATGCGGAGAGTACAGAGCTGGTGTGTAGCGGGTACCTTCTGTGACGATCACGG
ACCTGCATGCGGTCTGCTGGCACTAGGTATTTTACACATTAAATCAGCTCGTTTGACGATCACGG
ACCTGCATGCGGAAAATAAGGAAATGAGGAGCCACAGTGGCCCAACTTGACGATCACGG
ACCTGCATGCGGGATGTGCCTCCGTGTTTTGTCTCCTGAGCATGCTGTGACGATCACGG
ACCTGCATGCGGAGTTTGACAATACCAAGATTTGTACTGGAACATTCCCTCCCTGACGATCACGG
ACCTGCATGCGGAGCCTTCATCCACTACACTGTCAGCTGGTATCTCATGACGATCACGG
ACCTGCATGCGGAAAATGAAGCTGTGGCTTTTCTGTGAGTCAATTCTTTCCTGTGACGATCACGG

ACCTGCATGCGGGCAGCTTCTTGGAATAAGACCAAGTATAGCAGCAGAGTTGACGATCACGG
ACCTGCATGCGGCAGCTGGCAGCCCCATGGGAAAGATGTGTGAAGTGTGACGATCACGG
ACCTGCATGCGGTGGGGTTTGACGACCCATGGGAGAACAGACTTTCTTGACGATCACGG
ACCTGCATGCGGTCCTCTTCTTGTTTTCCCTTCAAAGCCGTGAGTCATGACGATCACGG
ACCTGCATGCGGCTCGTGCCTCTCTCCCTCACGCTCTGCATCTCTCATTGACGATCACGG
ACCTGCATGCGGTTTTCAGGGTCATTCAGGAAGTCATCCAGAGTTATAATGGCTGACGATCACGG
ACCTGCATGCGGCCCCGCGCTGTGAGTGTGCTATTTAGAGCTGACCATGACGATCACGG
ACCTGCATGCGGGGGGGAAGAGAGGATGGCTCGGATGCTGCATTTCCATGACGATCACGG
ACCTGCATGCGGCTGAGAACACAAGGCTGGCAAAGCTTGTCTGCTGCTGACGATCACGG
ACCTGCATGCGGCCAGCAAGCACTTCAGGCTCACACCATTTTAGGTTTGACGATCACGG
ACCTGCATGCGGCATTCTATCCGGCAGTCTTATTCTATGCACCCGAAGGTGACGATCACGG
ACCTGCATGCGGGATAAGAGCCATGCTTTCATGAAACATGGGTTGTGTGTTGACGATCACGG
ACCTGCATGCGGACTTTGAAGGAGAAATAAGAGAGTCGGCATTTTCTTGCACATGACGATCACGG
ACCTGCATGCGGTTCTTTGTGATGTTGTGATGAGTTGGAAACTTCCCGATTGGTGACGATCACGG
ACCTGCATGCGGGTTTATTAGAGCATGAACACCCAGGCACCCAGCTTTGACGATCACGG
ACCTGCATGCGGCTAGCCAGCCCTGTCAGGCAGAGTCTCCTCGAAGATGACGATCACGG
ACCTGCATGCGGTGTGGAAAGGACTGACCAACAGCTGAGGCCTACAGTGACGATCACGG
ACCTGCATGCGGTTGGAGTTGCAGGCCACAGAAAGGAAGTGACATGATGACGATCACGG
ACCTGCATGCGGGTCACTTTGGGCCTTCTTAATTCCTTCATCAAAGGCATGACGATCACGG
ACCTGCATGCGGTTCTCACTTCTTTGGCATCTACCACACCTGCCCCATGACGATCACGG
ACCTGCATGCGGCCCCACAAAGCCTGAGGTACATGGAAAGGAGCAGTTGACGATCACGG
ACCTGCATGCGGGTCTGGCTCCCAGGAGTGTGAGAAGCAGCCATGTTTGACGATCACGG
ACCTGCATGCGGTTCAGAGGCTGTATTCCACTTGGACTTGGCCCTACTGACGATCACGG
ACCTGCATGCGGCCCCTTCGCACAAAGAGCCCCATGAAAGAGTGCACTGACGATCACGG
ACCTGCATGCGGTAAAACAGACGCAGAAAATGTGTGTAGGACTTCTTCCTGAATGACGATCACGG
ACCTGCATGCGGAAAGAGCGTGGTGCGTCCAGTACCTCCATGTTCATTGACGATCACGG
ACCTGCATGCGGACAGGGATAATTCCAGAGTGATTTTAAAGTCAACTGCCAGGTGACGATCACGG
ACCTGCATGCGGCATCCGCACTTGCAAATTAGATGCCTGGCACATGCTGACGATCACGG
ACCTGCATGCGGTTCATTACAATACAAATTACAGGGGAGTTCCTCTGGGCATGTGACGATCACGG
ACCTGCATGCGGCGACCTTTCCCGTCATTTGGCTTTCCCTGTGATTATGACGATCACGG
ACCTGCATGCGGTCAGGGGAGCTTCCATCGTGCTGCTAATGGGACCTTATGACGATCACGG
ACCTGCATGCGGACCATGTGTCAACCCATGGCTGTAATGCTGACACTTGACGATCACGG
ACCTGCATGCGGGTTTTCTTTCTGGAATGAAAGGCCTTCGCAATTGAAACCATGACGATCACGG
ACCTGCATGCGGAACATCAAGTTCTGGAGACTGGCCATCCAGCCTCCTGACGATCACGG
ACCTGCATGCGGTTGAACTGAAGTACGTGTAAGTCAGCCTATGCATGTAATGGTGACGATCACGG
ACCTGCATGCGGAAAAGGGGATTTAGCTTGTTCTGGGCTTTGCAGCCTGACGATCACGG
ACCTGCATGCGGCGTACACTGGGCACTGTTCATACGTACTTCTCTGTTGACGATCACGG
ACCTGCATGCGGGCTAGGGAGGCCTGGCAGAGGGTGCCATTCAAATATGACGATCACGG
ACCTGCATGCGGGGAGCCCATGTTTCTTTCCAGCCAGGCTTGGAGACTGACGATCACGG
ACCTGCATGCGGCCACACGGGATTTGCTTCTTGGGGCCTCCCTCTAATGACGATCACGG
ACCTGCATGCGGAAGCTGCTCAAAGGACTCATTCAGTTCAAATTCACCTATGTTGACGATCACGG
ACCTGCATGCGGTTTTAAAGAACTTCAAATGATCTCCAGGGTACACAGCGCTTGACGATCACGG
ACCTGCATGCGGCCTTCCCAGACCCTGGGTCTTTTGCTGCCACATGAATTGACGATCACGG
ACCTGCATGCGGCTTCTTCAAGGTCCTATGTGTAGATTTTCTTGACTTGGCCATGACGATCACGG
ACCTGCATGCGGACTAAACCCTGATTGAAAATCAGAGCTCAGACATACAGCCTGACGATCACGG
ACCTGCATGCGGAAAATGGCCAGGCTTGTCTGTTGAGAAAGCCATATTGACGATCACGG
ACCTGCATGCGGCTCATCTGACTTCTCAGCTGTCCCTGAGAGGTACCTGACGATCACGG
ACCTGCATGCGGAGTGTGCAAGATCGCTGAGTTGGCAAGTGATAGTGTGACGATCACGG
ACCTGCATGCGGTGTATTGGATTTGCACAATTTCACGTTCTGAAAGAGGATGCTGACGATCACGG
ACCTGCATGCGGCCTCAACTTTGCAAAATGGGCCTTTTGAATGAAAAGGATCATGACGATCACGG
ACCTGCATGCGGTGAAAACTACCATCTTGAAAGGTTCTGGTGATAGTGGTTCCTGACGATCACGG

ACCTGCATGCGGTGCAGTGAAAGATGTGTAAGTCAAAGATTTGTAACCAGCCATGACGATCACGG
ACCTGCATGCGGTACAGGCTTCCTGGTTACCTTCCAGAGAGTAAGGCTGACGATCACGG
ACCTGCATGCGGAAGTCCCTACTACATTCTAAAATCTGGTCCTGACTAGTGGCTGACGATCACGG
ACCTGCATGCGGGGGCCCAGGAGTAGCACTTAAACAATGGCAGGCTTTGACGATCACGG
ACCTGCATGCGGTGTGTGCAGCTGGGGAGACTCACACTCAGAGGATTTGACGATCACGG
ACCTGCATGCGGTCAAAGCAGAGGGCATCTCGTAGAGCAACTTATCCTGACGATCACGG
ACCTGCATGCGGTCAGAGAACTGTTTAATGCGGGCTACCTCTGTTCTTGACGATCACGG
ACCTGCATGCGGCATGCTGCTTGTGGTTCCTGCTCTGCTCAGGACAGTGACGATCACGG
ACCTGCATGCGGAATGGGGAGGAAAACAGGCTCTGCGGCACAATATTTGACGATCACGG
ACCTGCATGCGGATTAGATGCTGAGTCCTACAATGTATTTGAGAAGCCCAGGATGACGATCACGG
ACCTGCATGCGGACATTTGCCTTAATTACTTTGGCAAGATTTGCATCAGTGGTTGACGATCACGG
ACCTGCATGCGGATTAGTCCCTGCCTCACTTGGAGGCCTGCACTTAATGACGATCACGG
ACCTGCATGCGGGTGGCCACATTCAGGCTCCAATTTCCTGGTGATTTTGACGATCACGG
ACCTGCATGCGGCATAGTGTAGGGCACTTGCAATCAAAACTAGGCTTAAAGCTGACGATCACGG
ACCTGCATGCGGTCAACTTCATATATCTTCTCCGCCAGATCAAAACAAGCTCGTGACGATCACGG
ACCTGCATGCGGTAGTATTAGATGTCACCGAGCACCATGACAGGCAGTGACGATCACGG
ACCTGCATGCGGGGGACCGGCACCGTATCTCCAGCAATTCGCAGATATGACGATCACGG
ACCTGCATGCGGTCCCTTTCAAGATTGGATTAGACATTAGGAATTTGGAGGGCTGACGATCACGG
ACCTGCATGCGGTTGATTCTAAAGTCTGAAAGCCACATGGACAGAGTTCATGTTGACGATCACGG
ACCTGCATGCGGAATTGGCTACTTTATGTGCCTCTTCCTAGATTGCCCTTGACGATCACGG
ACCTGCATGCGGTGAGGCTTTGAAAACTCAGCCTATGTTTGTCTTGATTTCCTTGACGATCACGG
ACCTGCATGCGGTAACTGACATCTAGAAGAAAATATGAGCTCAGGGGTCCGCTGACGATCACGG
ACCTGCATGCGGCCTGCTGGAACCAAGCTTTAAATGCACTTGTCAGTTGACGATCACGG
ACCTGCATGCGGTACCTCGTTTTGGAATCACTAATGTGGCCAATTTATAGCCATGACGATCACGG
ACCTGCATGCGGAAAATCAGTATCGTAGAGTGAGCAATGAATGGCATGGTGATGACGATCACGG
ACCTGCATGCGGCTCGCCCCGCGTCTCAGTCCTCAGTGATGGTAAACTGACGATCACGG
ACCTGCATGCGGCGTGATGCGCCTCAGCCCTACTTCCCTTGTCCTTTTGACGATCACGG
ACCTGCATGCGGCTCATTCCAATTTTGATGGAAACTTTTAGCTGGTGGATGGCTGACGATCACGG
ACCTGCATGCGGAATTTAGGGAGATGAAACTAGTCTGTGATCCGAGGTCTCACTGACGATCACGG
ACCTGCATGCGGCCACAGGTGCTTAGCCAGCTGGTCCTGTCGTAACTTGACGATCACGG
ACCTGCATGCGGTGAAATTGGCCTCGTAAAGCATCAGTTATGTTGTGATGGTTGACGATCACGG
ACCTGCATGCGGCAGGCGAGATGGGAAGTGCAGCCACTGAGAACTCATGACGATCACGG
ACCTGCATGCGGAAAAGATGAAACCCATTGTACACAGCTGTGTACTGCCTGACGATCACGG
ACCTGCATGCGGGGGGCTGCTTGACAGGATGACATCTGGTGATATCATGACGATCACGG
ACCTGCATGCGGATTCCTTTGGAAATCAATCTGTGGAAGTGAGTTTCCACTGATGACGATCACGG
ACCTGCATGCGGAAAATGAATTGGCTTCACCCAGCATCCAGCTTCTTTGACGATCACGG
ACCTGCATGCGGCTGGTGCAAGAGCCAAGTTTGTGCAGCCTGCAGAGTGACGATCACGG
ACCTGCATGCGGCTGTCAGGAGACGCAGTTGCCCAGACGTTTTCAAGTGACGATCACGG
ACCTGCATGCGGTTTTGGAGAGTCCTTGAGTTGGAGATTACAGGTGACCTGACGATCACGG
ACCTGCATGCGGTCAGAGGAGGGAGTGAGAACATCTGGGTCATGGGTTGACGATCACGG
ACCTGCATGCGGAGGAGTCCACAGTGAAAACAAGAAGAGGAATTTACGACAAGTGACGATCACGG
ACCTGCATGCGGACAGTCCAGCAACTTCCTTTCTAACTTCTCCTTTCACATGACGATCACGG
ACCTGCATGCGGATGCTGGATACTCCAAGACTTTGCATTTACATGGACATCATGACGATCACGG
ACCTGCATGCGGATTTAACAGAAACATTTTGGACCATTTTGGACCTTGGCAGTTGACGATCACGG
ACCTGCATGCGGCCCAGTGCAGCAGCGGCAACCATAAACCTCTCCATATGACGATCACGG
ACCTGCATGCGGCTGGCACCGGCAGCCTTATGTCAGGGCTCTCTTATTGACGATCACGG
ACCTGCATGCGGTCATCTTTTAAAGAAAGCAAGGACACAGAGGTGGATTCTCCTGACGATCACGG
ACCTGCATGCGGCAAGCCACTCCTGCAGGGCTCTGGTCTAAGTGCAATGACGATCACGG
ACCTGCATGCGGGGAAAAGCTGCAGGTCCCGCAAGACACAGAGCAACTGACGATCACGG
ACCTGCATGCGGACAGTCATTTTACTGAAGCTGCTTCAGTGGGTCCTTGACGATCACGG
ACCTGCATGCGGGAATTAGGCCCTGTCCTTTGGGAGAGACAGTCCTGTGACGATCACGG

ACCTGCATGCGGCAGCTTGGGAGTGTGGGCCAGTTTCGCCTTTAGAATGACGATCACGG
ACCTGCATGCGGAGAAGAGAAAGTGAAGATTCGATTTGAGGTGAGTTCAGCACTGACGATCACGG
ACCTGCATGCGGCCCTCTGGCTCCAACTGCAGCCCTTTCTACCTCATTGACGATCACGG
ACCTGCATGCGGTCCCAGACCCCACCTAAGCCTTTTCTCTTCAAAATTGACGATCACGG
ACCTGCATGCGGTTTTAATTCTCCGGTTGTGTTCACCAGGTGCTTGGTGACGATCACGG
ACCTGCATGCGGCTGAGTTAACTTTGTGGCGGCCTGTGGACTAGACCTGACGATCACGG
ACCTGCATGCGGGCACATGCAATGCAGAACGGCAGGGCCAGATTTGAAATTGACGATCACGG
ACCTGCATGCGGAAAATAACATCAGGCGATGGGGATACGATGCCAGATGACGATCACGG
ACCTGCATGCGGTTATGGCCATTTTACTTCTAGGAAGACAGGAAGTGTCAGGATGACGATCACGG
ACCTGCATGCGGTTTTCCACTCTGAAATAAAGTGACTGACCAGGAGTTCCCTGACGATCACGG
ACCTGCATGCGGCCTGTGGGAACTGCCGCACGGCCACTTTTATGAAGTGACGATCACGG
ACCTGCATGCGGTGGACACGTGTTGGTCCCACTGAAAAGAAACTCCCTGACGATCACGG
ACCTGCATGCGGAAATACAGAGGCTAGCACATGTGCTTGGGGAATGCTGACGATCACGG
ACCTGCATGCGGAGCTACAGTAGTGGAATGAAAGTGCTGTCCGTTCCTGACGATCACGG
ACCTGCATGCGGCAGCTCCTCACCTGTCCTCCACACGCATATCCCTGTGACGATCACGG
ACCTGCATGCGGCTTGCCCGAGGCTGCATACCTGTGCTCTTTCTGAATGACGATCACGG
ACCTGCATGCGGGCCCAAGTCACTGGCTCTAGAATTCTAACCTGTGATGACGATCACGG
ACCTGCATGCGGGTGTCTTTGCACTTCTTCAAGGCAGTACTGCCTGGTGACGATCACGG
ACCTGCATGCGGAACGCAGAATCCCAGCCTCCTCTATCCTCCTTGCCTGACGATCACGG
ACCTGCATGCGGGGAAGGTCACACAGTCATCGCAATAGTAACCTCAGCTGACGATCACGG
ACCTGCATGCGGTACGTGTCACTTTGTACCTGATTTCTATAAGACCCAGGACCTGACGATCACGG
ACCTGCATGCGGGGAGCATGTTGTGCCCAACTAGGGACAGCGCAGATATGACGATCACGG
ACCTGCATGCGGAGAACTGGGCTTTATTGATGTCACCTAAATGCACCTAAACTTGACGATCACGG
ACCTGCATGCGGAAATCCATATTTCCTCTCATACAGACCCCAGAGTTGCTTTGTGACGATCACGG
ACCTGCATGCGGAATCTTTATCCTTAGCCTCCATCTGGTTGCAGACCATGACGATCACGG
ACCTGCATGCGGAAAATGAGGGTGTTGGTAGCTTCGTTAGAAACTGAAAGCTTGACGATCACGG
ACCTGCATGCGGTGTCTTGTCATCAAACAGATAGTAAGAACAGGCAAACCTGGTGACGATCACGG
ACCTGCATGCGGCCCTCCACACTGCCAGCCTTTTGTGATTCAAGGCTTGACGATCACGG
ACCTGCATGCGGAGTATGTGAACCAGTAAACCCTAAAAGGTGTCCAGTGTTGTGACGATCACGG
ACCTGCATGCGGTCTGTGAGCTAATTAAGTGATTTGATTCTGACTCCCCGAGTTGACGATCACGG
ACCTGCATGCGGCTTCTGATTTCGAAGCAGTGGGGAGTCAGACAGGATGACGATCACGG
ACCTGCATGCGGAACCTGGCCTCTGGCAGCTCTTCATAAACGTCCATTGACGATCACGG
ACCTGCATGCGGTGAAAATCAATTTCTAAAACAGCCCTCTGTGTGCTCTCTCGTGACGATCACGG
ACCTGCATGCGGTATCTCTCCTTTTCACACATCGTGGTGGTGGCTTTTGACGATCACGG
ACCTGCATGCGGCTCTGTGTTCCTCTGTTGATTCAGTCTCTGGAATTAACGGTGACGATCACGG
ACCTGCATGCGGATCAGGATTCCATGCCCAGAATGCTACAAAGACTGTTGACGATCACGG
ACCTGCATGCGGGCTTGAGTTCTCCCACATCTCACTCAATTACACAGAAGTTGACGATCACGG
ACCTGCATGCGGTTCAGATTATGTAACAGATGCTGTGCTGGGTTAGGCATGACGATCACGG
ACCTGCATGCGGAAAACCTCCACTTGGCTGTCAGCATTGGAAATCAGTGACGATCACGG
ACCTGCATGCGGATGAAAAGGTGAAATCATTGCTTGAAATCGCTAAGCAGGACTGACGATCACGG
ACCTGCATGCGGGGGGAATTCGGGAGCTGGTTGGAAAAGAGTATTTGTGACGATCACGG
ACCTGCATGCGGGCACTTTGCAGCCTTGAGAGGTGCAGAAGAGACACTGACGATCACGG
ACCTGCATGCGGCGAGGGGTTCACCACCAGAGCCACCATTGTCAGAGTGACGATCACGG
ACCTGCATGCGGCCCCTTCTAGGATCTGATTCCTGAGGAATCACAATGTGGTGACGATCACGG
ACCTGCATGCGGATTTCACAATCACTTCCAGTGTCTTTTGCCAACCTCTGTGACGATCACGG
ACCTGCATGCGGAAAAGAAAGGGGCCCAATTCTCAACACTGTAAGTGGTGACGATCACGG
ACCTGCATGCGGCCAGCAGACTGCCGCCAGTCTTGTCAGAGATGTCATGACGATCACGG
ACCTGCATGCGGGTGGTGGCCACTCTGAATGGAAAGCAGCATCTCTCTGACGATCACGG
ACCTGCATGCGGCATCTCTGAGGCACTGCTCCTCAGCGGAGACTGTGTGACGATCACGG
ACCTGCATGCGGGTGGCTTTGCCTTTCAGCACGCATCCTTTCTACGATGACGATCACGG
ACCTGCATGCGGGAATGGGCAGAGCTGGGAGCTCTGAAGCCCTTTCATGACGATCACGG

ACCTGCATGCGGCCTAAACCACCCTGGGTCACCTGACCTAGTTTTCCTGACGATCACGG
ACCTGCATGCGGTCCCAATTTTAATTATGTCAGGCACTTCACAAAGGCCTCCTGACGATCACGG
ACCTGCATGCGGCTGCCCCAGCAGATGCTGTGTGTAAACAGTTGTATTGACGATCACGG
ACCTGCATGCGGAAAACATGGTTCTTGCCAGATCCTCAGGATTTGGGTGACGATCACGG
ACCTGCATGCGGGGGAGGCCCTCGAGGGAGAAATGTCTGCAGGAAATTGACGATCACGG
ACCTGCATGCGGAAGTTATCTAAGAGCTACTGCAGCTGTTTACTGCAGAGTGATGACGATCACGG
ACCTGCATGCGGCCCTGCTCAAAGCTGTGGTCACCCAAGGCTTTGAATGACGATCACGG
ACCTGCATGCGGCCACCTGAGCCACTCGGAGGCTCTATCCAGAGTCATGACGATCACGG
ACCTGCATGCGGAAAATTGCTGCTGTGAATACCAGTGTCGGCGTCCCTGACGATCACGG
ACCTGCATGCGGTGCCTCACCTTTACCTGGTGCTTTTCCACCACACATGACGATCACGG
ACCTGCATGCGGGTGCTGGCCTTGGGCTTGCAGACAGCTGATTCTTCTGACGATCACGG
ACCTGCATGCGGCTCCTCCGAGCAACCCTCTGACAACTCTGCTCCTTTGACGATCACGG
ACCTGCATGCGGCTGACAACCTCTGCAAGGGCTGCCAGATGTGAACATGACGATCACGG
ACCTGCATGCGGGAGGAAGCCTGCCCTGTCCTGTCCACCAGACTTTATGACGATCACGG
ACCTGCATGCGGCATTTGTTCCTTTTGCAGTTCCTACTGTTCTCAGCACTTCTGACGATCACGG
ACCTGCATGCGGATCGGTACAGACACAGTTCTGCTCCAGCATTTCCCTGACGATCACGG
ACCTGCATGCGGCGTGTTCTTTCTGTCGCTCTATTTACTGAATTACCGTGAGGTGACGATCACGG
ACCTGCATGCGGATGTGGAGCGAGGCTGAGTTCTGTATTTTAACACCATGACGATCACGG
ACCTGCATGCGGATTCTCACCTACTGAGAAATCCATCCTCTTATCACTGTGCTTGACGATCACGG
ACCTGCATGCGGTTTTAACCTGTCACGAATCCATGAAATCCTATCAGCCAGCTGACGATCACGG
ACCTGCATGCGGTGCATACTTCCTTTTAAGGTGCAGTTGAATCAGGAGAAACTTGACGATCACGG
ACCTGCATGCGGTGCCCAGCTATGAAACAGGTTAGCCACACATGACCTGACGATCACGG
ACCTGCATGCGGTTTGACTTAATTTCTTCCCATCCCTTTTCCACGCAGCTGACGATCACGG
ACCTGCATGCGGCCAGAAGCCTTGGATCACGTGGTGAGGGGAAGAGGTGACGATCACGG
ACCTGCATGCGGTTGTGCTATGTCGGGAAACTCTGTATCGAAGCTCGTGACGATCACGG
ACCTGCATGCGGTTGACTAAAACCCTCAGTTTCCATCAAACTTGTCACTCTGGTGACGATCACGG
ACCTGCATGCGGCATTAAAGCCTGTCACTGTGTGGCTCTGAAAACCTTGACGATCACGG
ACCTGCATGCGGCTGCAGGTCCCTGTGCTCCACAGAAGCCCACTTATTGACGATCACGG
ACCTGCATGCGGCACTCATCACCACCTTCCCTCACCCAGAGCCTCAGTGACGATCACGG
ACCTGCATGCGGGATTCCCACCTGCCCCTCAAGATTAATCTCCAAGGTGACGATCACGG
ACCTGCATGCGGACATTTCCAAATTCCTCTCCCCATCTCTCAGCCAGTGACGATCACGG
ACCTGCATGCGGCTTCGACCTCCAGCAGGGCACTCCACTCCACATTCTGACGATCACGG
ACCTGCATGCGGTCCTGGTCTGTCTGGCTCATCTTACCTGAGCCATGTGACGATCACGG
ACCTGCATGCGGCTCTCCAGGTGAAGGACTATGTCTAACTCAACTCTGCTTGACGATCACGG
ACCTGCATGCGGAAAAGCAGCTAACACATTGCTCTTTGCATATTGTTCACTCATGACGATCACGG
ACCTGCATGCGGCTAAGTTGAACTGGACTTGGACATGCACACTGAACTGCTGACGATCACGG
ACCTGCATGCGGACTCCTGATAAACATTCTCAGGAGGCACACTATGTAACTGTTGACGATCACGG
ACCTGCATGCGGATGGTGAAATCAGATTTTGTTTAAAGCATGGAGGAGAGGGATGACGATCACGG
ACCTGCATGCGGTCTTGCAGATTCTGCAGTCCTTAACATCTTTGAAAGAGGAATGACGATCACGG
ACCTGCATGCGGTTTCAGACAATGTAATAAGAAGGCCACGTGCTTTGACTTCTTGACGATCACGG
ACCTGCATGCGGGAACTCTTAGTGCAGTTTCAGTATTTTCTTCCTCAGGAGGGTGACGATCACGG
ACCTGCATGCGGTGAGCTGCTTCCGAATGTCCTCCCCTTCTTTGAGGTGACGATCACGG
ACCTGCATGCGGTGATGACCTTTCCCAGTCTCTGCAAGCCCTTCAGTTGACGATCACGG
ACCTGCATGCGGTGTGTCCTCTCTGAGCAAATCTGAATTGTGTGCTTAATACATGACGATCACGG
ACCTGCATGCGGGATACCCTTACAGAATGTGATCCACCATTTATGAACCTGCATGACGATCACGG
ACCTGCATGCGGGCATTGCATTCAGAGACTAAGTGAAAAGCTGGCAGATGACGATCACGG
ACCTGCATGCGGTTTAAAGCACAAGCTAAGGAAGAAAGCTGGTCTAGAAGGAGTGACGATCACGG
ACCTGCATGCGGCTACAGAAGGGTAATGCTTAGGGAGGGAATGATGTGCTGACGATCACGG
ACCTGCATGCGGTGTGGGTGGTGGTAGTTAAATCTAACCAAAGAATGATGTCGTGACGATCACGG
ACCTGCATGCGGACTTGAACCTATGCCTACCAGACATGACATGTGACTATTCATGACGATCACGG
ACCTGCATGCGGCTCTGGTTTAAGAATAAAGGATGCTGCATAGAAGGCTCACATGACGATCACGG

ACCTGCATGCGGTTTTCCATGCCAGAAGGGTAGAAGTTCCTGTGCTCTGACGATCACGG
ACCTGCATGCGGATTAAGAAACAGCAATGTCAATCGAGGCCCAACTCAAATCTGACGATCACGG
ACCTGCATGCGGATAGGAGTTATAAAGGGCGTGTGCCTGTTTTGTCTAGAAGCTGACGATCACGG
ACCTGCATGCGGAGTGTTGGGCAGCACTGAGTAGGATAGACCACCTGTGACGATCACGG
ACCTGCATGCGGTTGCTACCGATAAAGGAGCAGCTTCTCGAATGCTCTGACGATCACGG
ACCTGCATGCGGCTCATCCACAATCTGTGTGGCAAAAGGCATTGCAGTGACGATCACGG
ACCTGCATGCGGGCAATTCAGTGAGGAGACCGAGGCATGGAGAGCAATGACGATCACGG
ACCTGCATGCGGCTGGGTTGAAACCGTCCTCCGTAGGCTCCCAACTCTGACGATCACGG
ACCTGCATGCGGCCGTCGCTGCTACTGTGCTGGATGATGCCTGGTAGTGACGATCACGG
ACCTGCATGCGGGAGGAGCGATTTATGTTCTTAGTGGAAATATGTCGGCGTTGACGATCACGG
ACCTGCATGCGGGCTAGGCTGCCTCAGTAACTCTGAGGGGCATTGACTGACGATCACGG
ACCTGCATGCGGTTTTCTGTTGTTTTACAACGCAGTAGGAAGTGGGCATGACGATCACGG
ACCTGCATGCGGATGCAGATCAGCTGCACTCAGAAAACTACTCAAGTGATGACGATCACGG
ACCTGCATGCGGTTTTGCCTGATATAGTCACTTTATGCTGCAGGGGTGCTGACGATCACGG
ACCTGCATGCGGAATTATTATTAGAGGAGCTTGCGGTTGTTACATGTCTGCCTTGACGATCACGG
ACCTGCATGCGGTTTTAGCCATCTCCCCTGATGTCAAATGCTCAGGCTGACGATCACGG
ACCTGCATGCGGAATGTGGCTCCTTCAGAAATATACCACATACCTTTTGGTGTTGACGATCACGG
ACCTGCATGCGGGGTTTGTGGCTGAGAAGAGTGGGGAATGCACAAGTTGACGATCACGG
ACCTGCATGCGGACAGAGTCTTTCATGATCAAAGAGAACCAGGCTCTAGTAGTTGACGATCACGG
ACCTGCATGCGGTCCAGTATCCTAACGTGGACACTAATTGTTTCCCTCCTTGACGATCACGG
ACCTGCATGCGGTTCTTCATGAAAACAGCTTCTGCACAAATGATAGCCTTGTGTGACGATCACGG
ACCTGCATGCGGAACTAGCCATGGGCACAACTGGAGAAGCATTTAGGTGACGATCACGG
ACCTGCATGCGGGAGCTTTAGTGCAAATTGAGACCACCTACACATCTGACTTGACGATCACGG
ACCTGCATGCGGCTACAGGGTTTGACAACATCCAGGGTGAATCACAAAACTGACGATCACGG
ACCTGCATGCGGAAGCATAACATGAGAAGGGCTGACACACATAGATTTCTGTTGACGATCACGG
ACCTGCATGCGGCAGGAGACCTAGAGACCCAGGTCTGGGAGCCTAGCTGACGATCACGG
ACCTGCATGCGGCAGCAGCAGCTCAGTGCAGGGCAGTGCTGATAGTATGACGATCACGG
ACCTGCATGCGGCAGTGCCTGCAATCCTGGAGTGGATTTGGATGTGTTGACGATCACGG
ACCTGCATGCGGCCAGCAGTACGTTGAACAGTGTGCGTCCAGGTGTCTGACGATCACGG
ACCTGCATGCGGTGTAGGGCCCCTCGCCCTAACTCACAAAACCATTCTGACGATCACGG
ACCTGCATGCGGTGGGTCAGAAGCCACCAATATTGTCATCATCCTCCTGACGATCACGG
ACCTGCATGCGGCTTTTCTGAGAACCCTAGTAAGTCCCTCCAGTGGGGTGACGATCACGG
ACCTGCATGCGGGCCTTCATTTCCTAATCAGTTTTGCCCTGCTCATTCATGACGATCACGG
ACCTGCATGCGGATGCAAAATGGATCTGCTTTCCTTGGGCACCAATATGACGATCACGG
ACCTGCATGCGGAAAATCAGGCCTGCACTGCCTGTGCACTCCACAATTGACGATCACGG
ACCTGCATGCGGTTGCCAACTTTGTGCCTTCCCAAGCAAAATTAAAATACAGGTGACGATCACGG
ACCTGCATGCGGTACACACTCAGAAAAGATAGAAGCAGCAGCATATTTTGGCATGACGATCACGG
ACCTGCATGCGGGCATCTGGTTTATTGGAACTCAAACGTTCTGATTGTGCATGACGATCACGG
ACCTGCATGCGGTTTTCCTATTTCCATTGGCAAGCCACATGACAAGCATGACGATCACGG
ACCTGCATGCGGATGCTGCAGGGCATGAGACATCTGCACAGAGTTCATGACGATCACGG
ACCTGCATGCGGTCTCTTCCAGCATCTTGCATGTCCCAAGCACTGCCTGACGATCACGG
ACCTGCATGCGGCCAGGCAGAGAATGCTGCAGATCACGGCAGTGAATTGACGATCACGG
ACCTGCATGCGGTCCAGTTGTTCAGAGCACATTTGACTTCCAAATTCTCAAGGTGACGATCACGG
ACCTGCATGCGGTTTTGCACAAGTGCCTATATGCTATATAGAGTTTGCCCACTTGACGATCACGG
ACCTGCATGCGGCCCCGTTATCTGGCACAAGCTATTCAAAAGACACGTGACGATCACGG
ACCTGCATGCGGAAAATGTGAAGAAACACAGAACAGCTCATGAACACCTCCATGACGATCACGG
ACCTGCATGCGGCCCTGAGTTAGTCTATTTGTCATGCAAGAGCCTAGAAAAGCTGACGATCACGG
ACCTGCATGCGGCCTGAGTGATAAGAAATGGCCATAGGCCATTCCCATGACGATCACGG
ACCTGCATGCGGTAGAACTTATAACTCATGTTCGGAACTGTCCATGTTCACGCTGACGATCACGG
ACCTGCATGCGGACATTTTCAGAAGTTGAGAAATAGCAGTAGGCTGAAGGCATGACGATCACGG
ACCTGCATGCGGAGTCGCCATGCCTGGAATTCATGAACACTAGTTGATGACGATCACGG

ACCTGCATGCGGAAGAACTGGCGTGAGTTAGTCATGACAGGAGAGATGTGACGATCACGG
ACCTGCATGCGGTTTAATCTGTAGGACTCCATTCTCAAAGCACAGTCACTCCATGACGATCACGG
ACCTGCATGCGGGAGGACGGTCATTGCTCAGTGAGACTGCAATAGAGTGACGATCACGG
ACCTGCATGCGGGGCTGCAAGGGAGAAGGTGGGAGGGACAGCATTTATGACGATCACGG
ACCTGCATGCGGGCAGAATGAGCAGCACAGTCCCATAGGAAGAAGAATGACGATCACGG
ACCTGCATGCGGATTGCCTCCTTAGGCAAATAAATTCCCAAACCTTGAACATCTGACGATCACGG
ACCTGCATGCGGATTAATGCAACCTAAACAAAAGGGGTCTGGATACCCTCATGACGATCACGG
ACCTGCATGCGGTTTTAAGGACTATCTACAGCAATGGAAGAATCGGGTGTTGGTGACGATCACGG
ACCTGCATGCGGAAAATCAAGGACCTGCTACAAAGCACAAAGCCGACTGACGATCACGG
ACCTGCATGCGGTGCAATGCTTGCTGCTTACTGGTTAGGGCAGCTCCTGACGATCACGG
ACCTGCATGCGGTCTTTGCCAGCGACCAAGCAGAAAGCAAGACAAGATGACGATCACGG
ACCTGCATGCGGCAGGTTCTGAAGCAGTAATTCAAAGCCTTCCTCGCTGACGATCACGG
ACCTGCATGCGGGTGAGTCATTGCTAGTCAGAATATTACCTTTGCAGAGAGGCTGACGATCACGG
ACCTGCATGCGGTCCTCTCCTGGTTTAGGTATAAACTTTTGACTCACAGGACATGACGATCACGG
ACCTGCATGCGGATCATTCCTTTGGGCCTAGGATTGCATTTATTTCCATGACATGACGATCACGG
ACCTGCATGCGGAAAGCCCATCTCCTTTTGAATGAGCTCTAAAACAGTTCTCCTGACGATCACGG
ACCTGCATGCGGGCGACCTCTCCGCGCTGAGAAGGTTATCCGGATAATGACGATCACGG
ACCTGCATGCGGACAGCATAAACACACAGTGTATGTTGTTATGAGGCATCACATGACGATCACGG
ACCTGCATGCGGTGATGAGATACTGCTGGGGAGGGAAGAAGTGAGGATGACGATCACGG
ACCTGCATGCGGAAAAGAGAGAAAAGGGGTCTCTGCAGACATGGATGTTGACGATCACGG
ACCTGCATGCGGACCCAAGAAGATAATAAAGCAGAAGCATGTATCCAGGTTGCTGACGATCACGG
ACCTGCATGCGGCCCCAATTTTCGTTGGCACTGACCCCGATGATTTATGACGATCACGG
ACCTGCATGCGGAGACTGTGTGACTGGGAATTAAGGAGCAAAGCAACTTGACGATCACGG
ACCTGCATGCGGTTCCAGTCTGTGATTTACTGGGTTTCCATTCTGTTTCCTTGACGATCACGG
ACCTGCATGCGGATATTGCTTTTCATGCTTTATTTGGGACCGATTTTAGCCCCTGACGATCACGG
ACCTGCATGCGGCAGATCTTGGGCCCAGGTGTGAAGTCGCTGGAGAATGACGATCACGG
ACCTGCATGCGGGCATTTCAGGGCCAAGATGGGAGTGATTTCATTTTCCTGACGATCACGG
ACCTGCATGCGGATTGACACTATGCAGAAATGAAGGGGATTCAAGTGCCTTGACGATCACGG
ACCTGCATGCGGGGGTTTTCCAGACTTGCAACTGCTTTTATTCTTGGAAGCTGACGATCACGG
ACCTGCATGCGGCCCCTTCCATTTATATCCCAGGAACTGATTCAGAAACCATGACGATCACGG
ACCTGCATGCGGTAGAAATTGGATTTGGAATCGCTGAATGCTAGCAGACAGCTGACGATCACGG
ACCTGCATGCGGTTCCCAAGAAACCCTGCCAGCTGGGTTCGGGTATCTGACGATCACGG
ACCTGCATGCGGTCATTCTTTAGACATTTTGTGGTTTAGGGCTCAATCAGCCATGACGATCACGG
ACCTGCATGCGGGGGTATGATTTGCAATCCACAGTAACCGGTTTCAGAGCTGACGATCACGG
ACCTGCATGCGGCCAGCGAGGCAGGTTTCATCTCGCTTGCTAGACGTTGACGATCACGG
ACCTGCATGCGGCACTGATTAGACAGGCTTGAAGCAGAACCCACCAGTGACGATCACGG
ACCTGCATGCGGAGAAATGAATTCCTGGACTTGATATGTAGCAAGCTGGCATGACGATCACGG
ACCTGCATGCGGTTGGCTCGGGAGTGAGTGGGCTCAGTTAAGTGAGCTGACGATCACGG
ACCTGCATGCGGTAAGATGAGATGGTGCACAGGCGAGCACCCACCTGTGACGATCACGG
ACCTGCATGCGGAGGAGTGTTTGGATGTTATGATAGCCAGCTCCTCTGTGACGATCACGG
ACCTGCATGCGGTAAAGACCTGTCCTTCTATGTCAGCAGCCCAGCAGTGACGATCACGG
ACCTGCATGCGGTGACGTGTAAATACCACATTTAGGAGGGCTTATGATGATGCTGACGATCACGG
ACCTGCATGCGGCACGACGCCCTCACTAAGTGACCCACAGGACTCACTGACGATCACGG
ACCTGCATGCGGCTTAGACAGCAGTTCTGCAGTCGGCGTCTCACCCCTGACGATCACGG
ACCTGCATGCGGTTCGGGTCTCATTGTGACTCACTTTGATAGCCACATGACGATCACGG
ACCTGCATGCGGCGATTTAAGGGTGGTTCAGTAGTGATTTGATGAGTGCTGTTGACGATCACGG
ACCTGCATGCGGGGCTCAGGGTCATTCCCCTGCCCAAGCATTTCAAATGACGATCACGG
ACCTGCATGCGGTTCCAGAAGTTCATGCCCTGCATGGTGGGTGAAAATGACGATCACGG
ACCTGCATGCGGCGGGGTGGCACGTTGCTCAAACCCATCATTTGGAGTGACGATCACGG
ACCTGCATGCGGATCAAGAGTTTTCACATCGCCTAGACATGGCCTCATGACGATCACGG
ACCTGCATGCGGCTGGTGTGGCCAGGGGCTGATCTCACAGTAGACAGTGACGATCACGG

ACCTGCATGCGGCTCAGCAGCCTCCTGTGAAGCGAGGAAGGGTCTTCTGACGATCACGG
ACCTGCATGCGGCTGCCGGCCTCTGGAGATCAGTATGGGAATGCACATGACGATCACGG
ACCTGCATGCGGAGTAGGAAACGCTGGATGGGAATCCCTCTGCCCTGTGACGATCACGG
ACCTGCATGCGGTACAGTCAAGTGCCAGTGCAACACCTGATTCCCAGTGACGATCACGG
ACCTGCATGCGGGAGATGTCTGTACCAGACGCTGGATGACAGGCACCTGACGATCACGG
ACCTGCATGCGGAAAGTCACCTTGGAAAGAGGTTTTGTTCCTTAACCTTCTGTTGACGATCACGG
ACCTGCATGCGGTAAATAGGAAGCTCCGTGAATGAAAACAACTCCCTTCCCTGACGATCACGG
ACCTGCATGCGGTAAACATTCTAGTAATGACCCAACACTGCCAAGCCTGCTGACGATCACGG
ACCTGCATGCGGCAGCTCTGCCTCATGGTCGTGTTGACTGTGTGAGATGACGATCACGG
ACCTGCATGCGGAAAATGTCCTATTCAAATCACCTGGCACCCAGGAAATTTCTGACGATCACGG
ACCTGCATGCGGTTTTCCCAGGTGAAATATACAGTTGAAAACACCTGACAGCATGACGATCACGG
ACCTGCATGCGGTGGTTCCTCCATCTTTGATGTGTACAAGTGTGATGTTTTCCTGACGATCACGG
ACCTGCATGCGGCCCCACAGACAAGTAAACCACATTCTCTTCACATTCCCTGACGATCACGG
ACCTGCATGCGGTTTTGTTCAACACTCTTGCCCTATGAGTGCACAGAACTGACGATCACGG
ACCTGCATGCGGAAACATCCCTGAATGTTTAGCTTTGACAGAGATTCCAAGGTTGACGATCACGG
ACCTGCATGCGGGATTTGATAAGAAGCAGGGCTGTGTTTGGGCTCTGTGACGATCACGG
ACCTGCATGCGGTTTTGATATGGTTTCAAGCCCCATCCAAAACCCACATGACGATCACGG
ACCTGCATGCGGGACCTCTAGAAAGTAGGTGCCTGCCTTCCTGCAGCTGACGATCACGG
ACCTGCATGCGGCTGCTCCACAGAGGTGCCAAAACCAGGTTCCTTCCTGACGATCACGG
ACCTGCATGCGGTTTTATAACAAGGAACCTTTGGCTTTGAGATGACTGGGGATGACGATCACGG
ACCTGCATGCGGACCAAGACTCTGTTCAGACAAAAGTATCAGAAAAGTGAGCCTGACGATCACGG
ACCTGCATGCGGAAAACGCCAAGCACATTGCCTACACAAACTAAATATTCAGTTGACGATCACGG
ACCTGCATGCGGTATAATCAATGGCAGTAAGGGAGAGTAACTCGCAGTTCTCTTGACGATCACGG
ACCTGCATGCGGTATAGTGAAATTTCAGCTTCACATTCCCATCCATGAGCTCCTGACGATCACGG
ACCTGCATGCGGCTGCAAATATCCCGGTCTGCTCTCAGGACCCAGTGTGACGATCACGG
ACCTGCATGCGGGCTTCCTGTGTGCCCTCCTCAAACTCAGCCAATGCTGACGATCACGG
ACCTGCATGCGGCGTCATACAGTAGCAGGCAGGTGTCTTTGCTGGGTTGACGATCACGG
ACCTGCATGCGGAGTTGGACTGGATGTCCCTGGGATTGCAGAACTGGTGACGATCACGG
ACCTGCATGCGGATGGGGAGTGACATCAGGAAACTATAATCATCAGGACAACATGACGATCACGG
ACCTGCATGCGGTGGTTTGCCATAACTTTAAGTTTTAAGCGACCGCAGATTTGACGATCACGG
ACCTGCATGCGGGAGAGAGATGCATGCCCACAGCCATGCTTCCCATGTGACGATCACGG
ACCTGCATGCGGTAACTGGAGAGGGGTCTGAAGTTTGAAACAAGTGTTCCTGACGATCACGG
ACCTGCATGCGGACTCATTTTAATACCTGAAAACATGCTTTCCCCATGCTGGTGACGATCACGG
ACCTGCATGCGGTCGAAAGATTCTCCTAGGAAAAGAAAGGCTTGACAACATCGTGACGATCACGG
ACCTGCATGCGGAAATAACTCTAATGTGCAAGTAGATCCCCTGACCTCAAGCTTGACGATCACGG
ACCTGCATGCGGCAGAAGAGTCCAGGCCTTCACACCTTCTCTGCTTCTGACGATCACGG
ACCTGCATGCGGGGGCCAGCTATTGAGATTCCTGTGCCCACGCAATGTGACGATCACGG
ACCTGCATGCGGCTGGCCGCTGTCCACAAGAAATCCAGTTGCACCAATGACGATCACGG
ACCTGCATGCGGAAAACATGAGTTCCTGACTGGGAATTCGATGCTGCTGACGATCACGG
ACCTGCATGCGGCAGGCAGCTTTGCTCAGAGGGAGCAGCCTTCTAGATGACGATCACGG
ACCTGCATGCGGTAAAGTGGAATAGAGATGTGGGTAAAAGTTGTCTTGCCACCTGACGATCACGG
ACCTGCATGCGGTAATGGTACCCAATGAAAGACATAGCAGTCTACAAGGAGGGTGACGATCACGG
ACCTGCATGCGGGGGGACCCACACTCTGTAACTCCCACATTCAATTATGACGATCACGG
ACCTGCATGCGGGCATGTTATAGGTAAGCTGCAGAAAACGAGGCAGCTGACGATCACGG
ACCTGCATGCGGTTGTCAAAGAGGAACGGCTCTTGGCCATGGTTGCTTGACGATCACGG
ACCTGCATGCGGCAACCCTGGAGGAAACCACCTGGAATGATGGGAGATGACGATCACGG
ACCTGCATGCGGACTCCTCCAGGGAACATGGCCCTTTAATAGATCTCTTGACGATCACGG
ACCTGCATGCGGAAAATAATCCCAAAGCAGCCACCAGGGCATACTGCTGACGATCACGG
ACCTGCATGCGGTGCGATCAAGTCCTAGGCGGTATTCCCTTCTGCGCTGACGATCACGG
ACCTGCATGCGGCATAGACCCTGTGCAGAGTGCCCTCAACGAAGGAGTGACGATCACGG
ACCTGCATGCGGCAAGGAAGACCAAGTCTCCCGAGGGTTTGCATATGTGACGATCACGG

ACCTGCATGCGGTGTATGTGATTCTGCAGTCATGGTGAATGACACAGTCATGACGATCACGG
ACCTGCATGCGGTCAGAAGTTTGAAAAGGCTAGATTTCCTAGGCCAAAACACTTGACGATCACGG
ACCTGCATGCGGGAAAATTTGCAATTAGAACTTCAGTGCTGATGCTGGGAAGATGACGATCACGG
ACCTGCATGCGGCTGGAGTTAGTTTGAGACATGCACCTGTGCAGAACTGACGATCACGG
ACCTGCATGCGGCCCCAGAAAAGGAGAAGGAAGGGAATCCAGACCAGTGACGATCACGG
ACCTGCATGCGGAGTAGGGCCTGACACCACTCAGACTCGGCGTGTCTATAATGACGATCACGG
ACCTGCATGCGGAATGCTTAGATGGAGATTACCCTTTTGAGCATTTTGCCAGTTGACGATCACGG
ACCTGCATGCGGGCTTCTGAAATTAATGGGGACCTCCTGTTGGAGGATGACGATCACGG
ACCTGCATGCGGTTACTAGAGTAAGATAGGCTCAGTAGGTACCTGAAGGCACCTGACGATCACGG
ACCTGCATGCGGATCCCAAAGACCAGAGTGGTAGAAGCAGGTGGACCTGACGATCACGG
ACCTGCATGCGGCTGTGATCTCACAGATCCACAGAGAAGTGTCCCAGTGACGATCACGG
ACCTGCATGCGGATAAGAAGTCAGCCCATCAGCCTGAAATGCTCCCCTGACGATCACGG
ACCTGCATGCGGAAATCTTCCCATTCAGTGTTTTCTCAGTAGCAAACTCGTGGTGACGATCACGG
ACCTGCATGCGGAGAAGTGAACAATGGACCAAGTAAGCCTAAAACTCTGCTTGACGATCACGG
ACCTGCATGCGGCTTTCCCCTGCTCATTTTACAGTTCAAGTGCCATTCATGACGATCACGG
ACCTGCATGCGGTTTTGATCAATAACTACAGGCCTTTTGAGAGAGTGCCCTTGACGATCACGG
ACCTGCATGCGGTCCATACACAGTGTCTATCATGACCTACAAACCCTTTTCCCTGACGATCACGG
ACCTGCATGCGGATGAGGTGTAACAGAGAGAGATTACAGCCTTGGAACTGGTGACGATCACGG
ACCTGCATGCGGAGACTCTCCTGGTTTAAGACAATAAGCCATGACATAGAGCCTGACGATCACGG
ACCTGCATGCGGTGAAACCAACACAATCTTCCGAGTGGTTCCAGAAACTGACGATCACGG
ACCTGCATGCGGATATAGGGGATAATGTTGGCTCTGATGCTGTACATCCCCTGACGATCACGG
ACCTGCATGCGGTGTGGGAAGAGAGTCTCACTGTGTATCTTCTCCTGTTGACGATCACGG
ACCTGCATGCGGAAAAGGTATCCCAAGGATCCCTTTGTAGCTACATCTGTGGTGACGATCACGG
ACCTGCATGCGGGGCCTGGCCCCTCTGCATTTGACAATACGGTCCTATGACGATCACGG
ACCTGCATGCGGCCTGCCCTGGTTCACCTCTCCTCTTGACCCATCCTTATGACGATCACGG
ACCTGCATGCGGTCAGTGTCTTGAAAGGTCCTTCTATTGGAGGACACATTCTTGACGATCACGG
ACCTGCATGCGGGTTGCAGCAGGTGACCACTGCCCCAAATCTGTTTCTGACGATCACGG
ACCTGCATGCGGGTCTCATGAGTTCCTGACATACAAATAGTGCTGAGGCCTGACGATCACGG
ACCTGCATGCGGTTTTCAGTGTTTCTGTTGCATCGCTGTGTCTCTCTGTGACGATCACGG
ACCTGCATGCGGCCTGGCAGGAGGACTGCAGACAGGATGACCAAGAGTGACGATCACGG
ACCTGCATGCGGCCACCCACCCTCGTGGTTCCTGTTCATCTGCCTACTGACGATCACGG
ACCTGCATGCGGACGTGGAGGGCCCAAGAGGGCTAATATGTGACTATTGACGATCACGG
ACCTGCATGCGGCTCCACTTCCTGGTACCCTGTGTGAATAACTTCACTTTGACGATCACGG
ACCTGCATGCGGTTACAACCACAAGGACGTTTACATGGAAATGAACCATCTGCTGACGATCACGG
ACCTGCATGCGGCCCCATTTTCCTTTTAAATCAGCCAATGGGTGGTGGTGACGATCACGG
ACCTGCATGCGGCATAACACTGGTTTGGAGTGGAGGGCATTCATCGGTGACGATCACGG
ACCTGCATGCGGTGCTTATAGGAAGACATTTGATGGAAATGAGTCCAAAGGCATGACGATCACGG
ACCTGCATGCGGAGATAACTGTTTTCCGGTAGAGTGAATTGCCTGTTTGTTGCTGACGATCACGG
ACCTGCATGCGGGTTGGGACTTTGCTGGGCTGGTTTACAGGGCCAAGTGACGATCACGG
ACCTGCATGCGGAAGTGGATCTTCTAGTGAGAGGTCATCTGTTTTGAAAGCCTTGACGATCACGG
ACCTGCATGCGGTCCATGAACTAAATCCAAGTCTTACAACACAGGGAAGTGTGTGACGATCACGG
ACCTGCATGCGGTCATACTGTGCAGGGATGAAGTCTCCAATTTAGCATGAATGACGATCACGG
ACCTGCATGCGGTCTCTCATTGCTGCTGCTCTTTCAGAATCCATTTCAACATGACGATCACGG
ACCTGCATGCGGGAGTCAATGGGAGCCCCGTGCCTCTGGCAGATATCTGACGATCACGG
ACCTGCATGCGGATATGGCGTTTCAGTGGCATTGTGTGTTACCCTTCTGACGATCACGG
ACCTGCATGCGGTTAGGTAACAGCTCAGCCATTAGAAGAATGTCCTACACACCTGACGATCACGG
ACCTGCATGCGGTTCTCATTTTCTGTGATGAGAGGAATGTGAGGTACTGCCTGACGATCACGG
ACCTGCATGCGGAGCCTTCTGCCCAGCCACCACCACTATAGTTTTGTTGACGATCACGG
ACCTGCATGCGGTGGGAATGAAGCTGTCCCATGTGGGGCCTGTTCTTTGACGATCACGG
ACCTGCATGCGGCAGTTCCCTGCTGCCCAGTTCAGTCCAGTGGATCTTGACGATCACGG
ACCTGCATGCGGGGGCATCTCTCTTTAATCCGCATTAGGGGCTCTTTTGACGATCACGG

ACCTGCATGCGGAAAAGACTTGGAGGGGAGACCTGAGCCCACTTCTGTGACGATCACGG
ACCTGCATGCGGATCTTTGTGCAACAAGTCAATTCACTGAAGAGATCTGCTCTTGACGATCACGG
ACCTGCATGCGGCTAGGAGCCTCTGTGACCCCACCATAACTGGGAAGTGACGATCACGG
ACCTGCATGCGGGCTCTACCTCTCCAGTCTTCGGGCCACATTTCTCTTGACGATCACGG
ACCTGCATGCGGGGAGCACTCTAGTCCTCCGTCGACCTTGGCCTTTGTGACGATCACGG
ACCTGCATGCGGTTTTCACCTGGCAGCTCCCATCTGGTCTCACTCCCTGACGATCACGG
ACCTGCATGCGGTTTTGTCCAGTCTGCATGACACAGCCTCACATCGTTGACGATCACGG
ACCTGCATGCGGTAGTGTTCCCTCACTCCCCTCTTACTGCCCAACCTTGACGATCACGG
ACCTGCATGCGGACAGCCACGCTGTGTGCTGCTGAAGAAATGCACAGTGACGATCACGG
ACCTGCATGCGGGCGCATTTCTGCCCTTCTGAGCTCCGCTCACTTTTTGACGATCACGG
ACCTGCATGCGGCAGCAGCCGTTCCAACCTGCATGGGATCTTCACTCTGACGATCACGG
ACCTGCATGCGGTGACTTGCTTTTCTGCCATTCAACTGAGTTCCCTCTGACGATCACGG
ACCTGCATGCGGAGTGCTGGGGCCAGCGTGCAGTGTCTTGTATTCAGTATATGACGATCACGG
ACCTGCATGCGGGGTTCACACTAGCACAGTGCAACCTTCAATGCAAATGACGATCACGG
ACCTGCATGCGGAATTCACATGGTGGGATATTTTAGAAGGATGACCAGGCTTGACGATCACGG
ACCTGCATGCGGACGTTTCAGATTAACTCAAGGTATTGTGGTGATATTGCGGTTGACGATCACGG
ACCTGCATGCGGAGTGTACCTCCTTGTGAAATATCTTCAGAGTCTACTGGTGCTGACGATCACGG
ACCTGCATGCGGACATAAGCAATTGCTGGCAGCAGCTTGAGGGTCTCTGACGATCACGG
ACCTGCATGCGGATCTCATGGATATGAATGTCAATTCAAATCCCAGTGGCAGTTGACGATCACGG
ACCTGCATGCGGGGGGAAAGCCTAGAAGAGAAGAAACCTAGAGGAATCAAGCTGACGATCACGG
ACCTGCATGCGGACTTGATATGTTTCCCAGTGCTATTGTCAGAAAATGTCCCTTGACGATCACGG
ACCTGCATGCGGAAAATGAATAAGCCTGGGCCAAGGCTTGGAGACTTTGACGATCACGG
ACCTGCATGCGGTGAAATCTGCATTATCATCATCTGCAAGTTTAGATGGGGCATGACGATCACGG
ACCTGCATGCGGGGGGCAATTCAACTGGCTGATTCCAGCCAAGATGATGACGATCACGG
ACCTGCATGCGGCAACAGTCAGGATCCGTTCCCTTCTGATCATCCATTGACGATCACGG
ACCTGCATGCGGGACCAGACACTATCTCAGGCTGGCTGCCCCACATGTGACGATCACGG
ACCTGCATGCGGCCTTGCTCCACACCAAATTCACAGTCTATAAACCTGAGCTGACGATCACGG
ACCTGCATGCGGCTCCAGTGCTCCTACTACCATACTCACTCGAACATTCCTGACGATCACGG
ACCTGCATGCGGCGATTCTGACCTGGAGATGTCAACAGCTACTTGATGCTGACGATCACGG
ACCTGCATGCGGCACTCTCTTCTATCTTTCTGTAGCTAAGCCATCCCCAAGTTGACGATCACGG
ACCTGCATGCGGTTGTCGATTCACCCTCTTTAACCCCTGTCGGGGTGTGACGATCACGG
ACCTGCATGCGGATTTATCTCAGTTAGAGAAGAAGAGTCACCTGTCACCAGGGTGACGATCACGG
ACCTGCATGCGGCAGCTTTGGGGTGCCAGCTCCTCCCATAAAGCAAATGACGATCACGG
ACCTGCATGCGGCACATACCTAGGGATGATATTTCTTTGCAAGGGCTCTGCTGACGATCACGG
ACCTGCATGCGGGGGGAGAGAAATGCTCCTGCACTGCCTCTGATTAATGACGATCACGG
ACCTGCATGCGGGTATGAAATAGCCACAAACATTCAGTGAAACCAAACACCCCTGACGATCACGG
ACCTGCATGCGGGGGGCAGTGCAAGTGTATTTCTAATGGTGAAAACCATGACGATCACGG
ACCTGCATGCGGTAATGTACCATCTAGTTAATAAGAGCTCCTCTGACCCACGCTGACGATCACGG
ACCTGCATGCGGAATTCTGTCACAAAGATCTGGGTCTCATTAGGAAGGAGAGGTGACGATCACGG
ACCTGCATGCGGCCCCAAACCAAACATCACGGCCTTCTCAGTTGTTTTGACGATCACGG
ACCTGCATGCGGTTTTCCACGAAAGGAAGTGTTACCACATTTTCCAAGTGGTGACGATCACGG
ACCTGCATGCGGGAGGCATCTATATCCTTACTCCTTCATCCTCTCCTTCCCTGACGATCACGG
ACCTGCATGCGGCCCCACCACCCACACAACATCTGCAATTCTTAAACTGACGATCACGG
ACCTGCATGCGGTTGCACTTTCAAAGGAATGCTTGTATAGAAACTTTCTCGGCTGACGATCACGG
ACCTGCATGCGGTACGCTTTGAATGGCTGTTCAACAGCATAGAAATTAGCTGATGACGATCACGG
ACCTGCATGCGGCCCCTTATAAAGACTTGACAGTGGGCCAGAATAAACAACTTGACGATCACGG
ACCTGCATGCGGTCAGGATGAATTCAGTTGAGACACAAAGTACACACTTCCAGTGACGATCACGG
ACCTGCATGCGGTTTTCCCTTCTCTGGTTACTGGCCTCAATAACCAGGTGACGATCACGG
ACCTGCATGCGGAGATTGCAGACTTCTTATAATATGTCCATTTCACCAGGCCCTGACGATCACGG
ACCTGCATGCGGCCGGGAAAGGCCACTGGAAACCACCTCACATGGTATGACGATCACGG
ACCTGCATGCGGGGGGCCTTGCGGGAGCCAGTAATAACCTTATCTCCGTGACGATCACGG

ACCTGCATGCGGTCAACATGTTCTGTCAGATTGAATGGGGCAGCCAGTGACGATCACGG
ACCTGCATGCGGCCATCCTCCTGGAGTGAGCGGTTGAGCCTGGATTGTGACGATCACGG
ACCTGCATGCGGGTGCTTAGACCTATAATGGGTGCAAGCAGCGTTCATGACGATCACGG
ACCTGCATGCGGTTCATAGTGGCTTTCTAGACCCAGGGACTTGGCCCTGACGATCACGG
ACCTGCATGCGGCAGCCCTGCTGCTCCACTCCTCTTCTTGCTTCATTTGACGATCACGG
ACCTGCATGCGGCACGAGTCTCCTAGACCACCGAACGATGCCTGCATTGACGATCACGG
ACCTGCATGCGGTTGAAAGACACTTCTGCTGATCAAAGCAGCTGATGTGTGACGATCACGG
ACCTGCATGCGGAATATTCTTTGGTCTAAAATTACTGTCATGGCGGAGCAGCTTGACGATCACGG
ACCTGCATGCGGGCTAGTAGGATGTGATAGGAAACTAAAACCTAGGGGAGACCTGACGATCACGG
ACCTGCATGCGGAATACTTCCTCATCCAAACTTGAGAGCAATTTACCGTCAGGTGACGATCACGG
ACCTGCATGCGGAATAATAGAATGCTTTCAGCACTCAAGTGTGAGTGAGTGCTTGACGATCACGG
ACCTGCATGCGGCATACATGAGAGAAAGCCGTGGGGACTACAGAAGCTGACGATCACGG
ACCTGCATGCGGCAAGAAGCAGATCTAGCTGGGGAGGCCTTTGCAGATGACGATCACGG
ACCTGCATGCGGGGATGTAGTTGTGTGGAGAGGCCACACACGTGGAATGACGATCACGG
ACCTGCATGCGGATAACAATCCCCAGCCCCTGCATTTCCCTCTCCGGTGACGATCACGG
ACCTGCATGCGGTCTCTGAGACATGGCAAGCGCTGCTTCAGGCTCATTGACGATCACGG
ACCTGCATGCGGGCTTCACTAGATTCAGGCCTGGGGCAGTAAAGAGCTGACGATCACGG
ACCTGCATGCGGTTTTCAGGCAGGGGAGCCATCTAATGTCAAGTGCCTGACGATCACGG
ACCTGCATGCGGTACGTGCAGGAACTGGTCTGTTAATTAGCAGCTCTTGACGATCACGG
ACCTGCATGCGGTCATGGAAGGGATAATATATTCTAGAAACAGGAGTGCGGCCTGACGATCACGG
ACCTGCATGCGGCCTGAGCCAAAATTAAGATTCTTCTATGGCAGAAACTTGGCTGACGATCACGG
ACCTGCATGCGGTTTTCCTTGTCAACGACCCCAGGAATGAGTTTGGGTGACGATCACGG
ACCTGCATGCGGATTACAGGGTAGCCAGGGAAAGGGAAAGCTTCACGTGACGATCACGG
ACCTGCATGCGGTAAAAGGAGGATTTCAGTTGAGTAGGAAGTGAGAAGAGGGCTGACGATCACGG
ACCTGCATGCGGTTAAAACAAGCGTTAAATGAAAACCCACACACACTCAGAGCTGACGATCACGG
ACCTGCATGCGGACACAAATCCAACCACGCTTACAAAACCATCACAGATGACGATCACGG
ACCTGCATGCGGAAAAGAACAGGGGTGGAGACTGTTCTGAGAGCATGTGACGATCACGG
ACCTGCATGCGGCTGGGTTCCCTGAAGGGAATTCTCAGCTGTATGTGTGACGATCACGG
ACCTGCATGCGGCAGGCCGGCAACCTCTGAGTGCTTAGAGAAAGGGATGACGATCACGG
ACCTGCATGCGGCGGGATTCCTCCTGTGCAACCCCTCTAGTCTCACTTGACGATCACGG
ACCTGCATGCGGCAGACTCAAGTCTGACTAAGGGGCCAGGTGCTTTGTGACGATCACGG
ACCTGCATGCGGACATGTGCCAGTAACAGTGCAGCTGCCTGTGGTTCTGACGATCACGG
ACCTGCATGCGGTTTATGACATGAACATAAACTACTGGCTGCATCGTTCTGCTTGACGATCACGG
ACCTGCATGCGGAAAAGCAAGCAGTGGACTGCTCATTGGTAAAATTGAGTTGACGATCACGG
ACCTGCATGCGGTTTTCTTCCTTTTACTGCTTCTCATCTGTTCCCCGCATGACGATCACGG
ACCTGCATGCGGCCAGCCTGAAGCAGCACCATGCGAGCTTAGACCTTATGACGATCACGG
ACCTGCATGCGGGACACTCCCTCTCGAAGCTGCTTCTCCCCAAGCTATGACGATCACGG
ACCTGCATGCGGCCCCAGCAATTCAATTCACTGGCTGTCCTGCTCCTTGACGATCACGG
ACCTGCATGCGGGTCAGTACTGAGAGTTGCATGTTTGACCCTCGGGGTGACGATCACGG
ACCTGCATGCGGCTGCTGACCCCTTGCGCTGTCAGCATTCAGAGACATTTGACGATCACGG
ACCTGCATGCGGCACAGCCTCCCAGGCTAACAGCTGTCATGGAACAGTGACGATCACGG
ACCTGCATGCGGCACACTGGTGCACTGACCTGCTCCTGGTGTCTTTGTGACGATCACGG
ACCTGCATGCGGAAAAGAACATTTAAACCATCTTGGGCTAGGGGTGGATATGCTGACGATCACGG
ACCTGCATGCGGACCACCCCACGGAAGCCAAACAAAATCTCTCTGCATGACGATCACGG
ACCTGCATGCGGCCCCAATTCGAATCCAGAGGATGGGAAGGCATAAATGACGATCACGG
ACCTGCATGCGGTGGTGGGACTATGTATTTTACGTATGGTGAAGTCACCAAGCTGACGATCACGG
ACCTGCATGCGGCCAACACTTGGCACTTGTAGGCAAGGTAGTCTTCTTGACGATCACGG
ACCTGCATGCGGTTTCATTTGCTTGGTAATGAGGAATATTGGTGCTTTCGTCCTGACGATCACGG
ACCTGCATGCGGGTGTTCCTTTAGCACACCTCTACACTGCATGGCTCTGACGATCACGG
ACCTGCATGCGGTGAGATGTCCTGTGACTGTTTCATGTGTAAAGTTGCCTTGACGATCACGG
ACCTGCATGCGGCCCCAAAGGACTCACATATTCCTTCAGGGCAGTGATGACGATCACGG

ACCTGCATGCGGGTACTTCTGATTCATCCTTAGCAGCTACCTTCGCGCTGACGATCACGG
ACCTGCATGCGGTGAATTAATGAACAAAAGAACAAGCAACTTTGGTGCCTGGTTGACGATCACGG
ACCTGCATGCGGCTTGGTATTCACATGACTTCTAGGGCGACAGCTCATGACGATCACGG
ACCTGCATGCGGCGGGCCTGAACTGAATAGAAGATAGAGGAGAAAAGGTGTGACGATCACGG
ACCTGCATGCGGGGTGCCTTCGGTTGTGAGGGCCTGTCTAGGAAGAGTGACGATCACGG
ACCTGCATGCGGGCAGGCAGCACTGAGCTGTCTCATGCAAAGCTGAGTGACGATCACGG
ACCTGCATGCGGGTAACAAAGAGGGACGTGAAGTCAGAGGGGAAGGGTGACGATCACGG
ACCTGCATGCGGAGGTAGCACAGGGCAGTCTTGACTTTGAACAAAGATGACGATCACGG
ACCTGCATGCGGCTGGCTTCCTGAAGTCAGCTGGCCGGGTTTTGAAGTGACGATCACGG
ACCTGCATGCGGCCGATTTTCCAGCAGTGATCTTTGATGCCAACCCCTGACGATCACGG
ACCTGCATGCGGCCCCTACCTTCTACCAGATGTCTCTGAGCTCACCTTGACGATCACGG
ACCTGCATGCGGTATTCAGTTCAAAAGTATGCCTGTGTGTAAAGAGAGAGCCCTGACGATCACGG
ACCTGCATGCGGCTGAACATGAGTAAGCAAAAGTCTCAGCGCAGAGATTAGATGACGATCACGG
ACCTGCATGCGGCAAGTAGAATGCTGGCCCGAGAGGAGGCGTTTACTTGACGATCACGG
ACCTGCATGCGGGGAAAGCCAGGCCCAGCACGCTCACTGCTATCTATTGACGATCACGG
ACCTGCATGCGGCCTCTCACACAGAGGGATTTTGAATCGAAGCCAGCTGACGATCACGG
ACCTGCATGCGGCCCACCCCAGCCATATCCAGGTTCAGTTCCAGGTATGACGATCACGG
ACCTGCATGCGGTTGAAGGCACATAGGCAGAAATAGATTGCAATAGTTGCTGCTGACGATCACGG
ACCTGCATGCGGAGAAGTTTGATGATGTCACTGGAAGCATATTCCACAGTCCTGACGATCACGG
ACCTGCATGCGGCTTGTGAACTGGCCTTCCTGTGATCAGAAGCATCATGACGATCACGG
ACCTGCATGCGGATCACAAGCTTGGCTTTGGATTCTTTTAATGGCTAGAAGGATGACGATCACGG
ACCTGCATGCGGGAATCATGGGGTTGGAAGTCCACCAGTTTGGGTATTGACGATCACGG
ACCTGCATGCGGTAACTTAATTAGAGATACATGAGTTTGCAGCTGTGCTGGGCTGACGATCACGG
ACCTGCATGCGGATGCCGCAGCTTGGCATGTTTAGTCCAGAAGGCATTGACGATCACGG
ACCTGCATGCGGATTTTATAATGACCAACCAGAAAGTGGCAAGGACTGTCACTTGACGATCACGG
ACCTGCATGCGGCTTCCCATACAGCCCACCTCCTCCTCTATCTCCCTTGACGATCACGG
ACCTGCATGCGGCAGGCACACGGAAACGAGAAAGGCAGAGAAACCCATGACGATCACGG
ACCTGCATGCGGGGACAAGTCATCCAAGACTTTGGTCACATGGCCATTGACGATCACGG
ACCTGCATGCGGTAAATCCAACATGTGTGTGTGCATTTCATACCAGTAGGTCCTGACGATCACGG
ACCTGCATGCGGCTGGGGATGGAAGCAGAGACAACTTGGAGGCCAAGTGACGATCACGG
ACCTGCATGCGGGGGGAGAGACTGACAACAGCCCAGCTCATACCTCAGTGACGATCACGG
ACCTGCATGCGGAATAAGTAATATAGATTTGCCCAGGCTTGTGGGCAGAGTTGACGATCACGG
ACCTGCATGCGGGGTATACACCCCATAATAGCAGAGGAAGGCCACAGTGACGATCACGG
ACCTGCATGCGGAAAATAATAACGAGACCACCGGCCCGCCCTAGATGTGACGATCACGG
ACCTGCATGCGGAATGCACAGATAACTTCGTATATCCACTTTCCTGAACCTGCTGACGATCACGG
ACCTGCATGCGGGTTAGTGCTTTGAAGTCCTAAGTCATAGGGCCTGCTTGACGATCACGG
ACCTGCATGCGGGCTCTTGATGCAGTAGAATTTGTCTTCAGATTTGCAAAGGGTGACGATCACGG
ACCTGCATGCGGAAGGCAAACCACTAGCATTTTGTATGGAACTTGATGCATGACGATCACGG
ACCTGCATGCGGAACTTAAGACAATATAGAGTGGCTGACATCCCCTAACGTGGTGACGATCACGG
ACCTGCATGCGGACCTAGACCAAGTCATTGTGATCCTCATCCTAGATTTCAGTTGACGATCACGG
ACCTGCATGCGGAAAAGTGCTAAATAAGCCCTGAAGGGGCTTTTGGGTGACGATCACGG
ACCTGCATGCGGAAAATTCTTTGGTTCAGCTGAGAGAAACTTGCAAGCTGATGACGATCACGG
ACCTGCATGCGGATTTAAAATGAAAGCAGCTGCCTGTTTTCATCTCTCTGCATGACGATCACGG
ACCTGCATGCGGTCCTGAGGAAACTCTTCTGCAACGTGTTCCAGCCCTGACGATCACGG
ACCTGCATGCGGCCAACTAACACTTGCGGCTACCTGGACGACAGCCATGACGATCACGG
ACCTGCATGCGGATCTAGTTGGAATGAGAGTTAGAGGCCATAGTCTGTCAGCTGACGATCACGG
ACCTGCATGCGGTGGGAAAGCAGCTTTTATTCCAAGGTGTGCCAACCTGACGATCACGG
ACCTGCATGCGGGAAAGGCCACATGTTATTGTCACAACCTGGTACCTACATGACGATCACGG
ACCTGCATGCGGACCTTAGAATTGGGAAATCAGTTTAGCCCTATCTGCATGTGTGACGATCACGG
ACCTGCATGCGGTAGCCGACAACCACACAATTGTTCCAACTTGAGGTTGACGATCACGG
ACCTGCATGCGGCCCCATACTCCTGAGGAAAAGCAGACCAGAGACGCTGACGATCACGG

ACCTGCATGCGGGACTCCATCTTGTCAACTGTCATGGTTCCCATGTGATGACGATCACGG
ACCTGCATGCGGATCACTGAAATTGAATGAGAACACAGAAGGGAAGAACAGGGTGACGATCACGG
ACCTGCATGCGGAAATCCCCACAGAGTTAAAGAGGATGTGAAGATTGCTTCATGACGATCACGG
ACCTGCATGCGGTGTTTAATGTTTGTGTAAGTGCTTTGGGTTGGTTATGTGCTTGACGATCACGG
ACCTGCATGCGGGTCTGAACATGTGCTCATTTCCATGGCTCATTGAGAGTGACGATCACGG
ACCTGCATGCGGGGCAGACAGTCCAATGATACTCTTTAGAATCATTCCCATGGTGACGATCACGG
ACCTGCATGCGGAAAAGCATTGAAGAAAGTGCCAGTGTATTGATTTTGGCCATGACGATCACGG
ACCTGCATGCGGAGAAGTGCCTTTGCAGATGAGGAAGAACCTGGCCATGACGATCACGG
ACCTGCATGCGGAGGCTCAATCCAACATCCAAAGCCAGAGGCCATATTGACGATCACGG
ACCTGCATGCGGTTGGGTTCAAATTATAGTCCCTTTACACACTCTCTGCCTCATGACGATCACGG
ACCTGCATGCGGAAAGGCCCAAGACTCTCTTTTGTTATGCTTGCCTAAACTGACGATCACGG
ACCTGCATGCGGAAAATGGATCACCACGTCCATTTCTATGGTCCAACTTTGTTGACGATCACGG
ACCTGCATGCGGAACATTAGGAACTCAAGCATACAAGATTGTATGCTGGTGGTTGACGATCACGG
ACCTGCATGCGGCCTGTGCTGGGTGGATCAACATGGAGTGTGGACGATGACGATCACGG
ACCTGCATGCGGTTTGTCCCTGATTTTAGGTGTAGTGTTGTGTACACTACTGCTGACGATCACGG
ACCTGCATGCGGATGTTTGTAAACATTCATTGCACGCACAACAATGTGACTCATGACGATCACGG
ACCTGCATGCGGTTTTGCCTACTGGAGCTCTTACAGGTTAATCCTGTCTTCATGACGATCACGG
ACCTGCATGCGGATCAGTCACTAAAGCTCACTAAGTTAGTAAGCTTTGCGCCTGACGATCACGG
ACCTGCATGCGGCAGATGACCTGGGCAGGAATGGGTGAGTCTCTGTGTGACGATCACGG
ACCTGCATGCGGTGGAGAGAGTGAAGAAACTGCTACCCTTAATACCTGGATGACGATCACGG
ACCTGCATGCGGTTTTCTTCCAGTTTGCCAAGGCACGAGTAACAAGCTGACGATCACGG
ACCTGCATGCGGGCCTCCAGAGGATGTTCAATAACTGTGAGGTGGTCTGACGATCACGG
ACCTGCATGCGGTTTTGCTATGGCAATGACAAGTCTTACAGAGCTACAAACGATGACGATCACGG
ACCTGCATGCGGAGAAACAGGCTAATCTGGAGTTCCATGTCATCATAGACACTTGACGATCACGG
ACCTGCATGCGGGACGTTTATCCCTGACCATTACCTCAGTCATGATGTGCTGACGATCACGG
ACCTGCATGCGGGCTAATGACCCCATAGGAGCCAGCTCTGAAGGCTTTGACGATCACGG
ACCTGCATGCGGAGAGTTTCACTGAATTTTGGATGGGGTTTACTTAGCCTTCTTGACGATCACGG
ACCTGCATGCGGGGCATGAGAGCACAGTGTTCCAGAACAATGCTTACTGACGATCACGG
ACCTGCATGCGGTGCTCATTATCACAGGGGTCAAAGGCTAACGTGCATGACGATCACGG
ACCTGCATGCGGGGGATTGTTGCAGATCGTGGACATGCTGCCTCCTGTGACGATCACGG
ACCTGCATGCGGTCCATGACTGCAATCGTCTACCTATTTTACAGTTGTTGAGCTGACGATCACGG
ACCTGCATGCGGACTCGTGTGCATTAGGGTTCAACTGGGCGTCCTAGTGACGATCACGG
ACCTGCATGCGGGGCTCCCTGGACCCATTTTAGACCTTGAGTTCTTGTGACGATCACGG
ACCTGCATGCGGGAGCGAATTCCTTTGGAAAACCTGCAGATCATCAGAGTGACGATCACGG
ACCTGCATGCGGCCATGAGAAATTTACAGGGTGAGAGGCTGGGATGCTGACGATCACGG
ACCTGCATGCGGCCCAGCGAGCTTGTCACTCAATTCCACCTCGGAGATGACGATCACGG
ACCTGCATGCGGTTTTACCCAGTACACGTGCACTGAGTGCCGGCTGTTGACGATCACGG
ACCTGCATGCGGGTGTAAGATACTGCAGGGGAAGTTACTGAGAAGATGGCTGACGATCACGG
ACCTGCATGCGGAGATACTGGAATGGGAAGATTTAAGCGGGGTACCAGTGACGATCACGG
ACCTGCATGCGGATTCTGAGTAAAAGAGAGTATGACCAAACAAGCTGAGCAGGTGACGATCACGG
ACCTGCATGCGGAATCGTGAATCTATGTGTGTAGGCAGTGAATAAACTGCCATGACGATCACGG
ACCTGCATGCGGGCTGCTACACAGAAAGCCCTGTGTAAGTTAAGGATAGAGTGACGATCACGG
ACCTGCATGCGGTTTTGTGAGCAGGACTCGTGAGGTTCCTGTGAGAGTGACGATCACGG
ACCTGCATGCGGCGCCGAGGGTGAAGGCGTCATTAGGTCGGATTTCATGACGATCACGG
ACCTGCATGCGGCATGCAGTTCTGACAGGGAGGCAGTGCTGGTAGGATGACGATCACGG
ACCTGCATGCGGCAAGCCAGCACCTCTGGTCATGGCTCCTCAGTTCATGACGATCACGG
ACCTGCATGCGGCACATGCTTGTGCTGTGCGTGCCTCCTGTGAGATGTGACGATCACGG
ACCTGCATGCGGCTGTAGGAACAGGCCTCAGCAGCTCCTTCCACTTCTGACGATCACGG
ACCTGCATGCGGCTCCTCTGGCAGTCATTGCCTGTGGTGTGCTGAAGTGACGATCACGG
ACCTGCATGCGGCCCCTGCTCTCTCTCAGGTCTTTGGTCTGTTGTCCTGACGATCACGG
ACCTGCATGCGGCCCTGTCCTCTCCTGAGTTCCATCCACCTGTGCATTGACGATCACGG

ACCTGCATGCGGTTTTCATGAGTGTTTTCACGGATGGTTCTGCTGTCATCTGACGATCACGG
ACCTGCATGCGGTCCAACCTGATAAACAAAGCACCACGATTCAGCCCTGACGATCACGG
ACCTGCATGCGGTCTGTGCATCCACTGAAGAAGCCTGTTCCACTGTTTGACGATCACGG
ACCTGCATGCGGCCCTCACTTCCACACTCCTCTAAGATGTGTGCCTGTGACGATCACGG
ACCTGCATGCGGTGCCCTTCTGGGGAAGCTCATTTTCCTAGCAGCCTTGACGATCACGG
ACCTGCATGCGGCCAGGATCTTCAGGGGTGAATCCCTCCTTTCCCACTGACGATCACGG
ACCTGCATGCGGGTTGGTACTCTGTACACACAACATGCCCATTCCCTTGACGATCACGG
ACCTGCATGCGGGCCTGGGGAGCTGGGCATTGCTTCATGAATCAGAGTGACGATCACGG
ACCTGCATGCGGAAAGTCACAGATGCTCATTGCACCATTGTGAGAATGAATGTGACGATCACGG
ACCTGCATGCGGAAGATAGTGCTTATAAATCAGCCAGCAAGGTACCCAGCTGACGATCACGG
ACCTGCATGCGGTAGTTCTTCTTTGGAAACAGCTGTACAGTTTCACCATCCTGTGACGATCACGG
ACCTGCATGCGGTGCATCCCTGGAGTCTACCTGTCTCTGTCATACATTGACGATCACGG
ACCTGCATGCGGCGTGTCACTCTCATATCCTTTTCTCTAATGAAAAGCTCCGCTGACGATCACGG
ACCTGCATGCGGCTCACCTGGCATCCAGGGCTGTAGCACTCAGGAACTGACGATCACGG
ACCTGCATGCGGCCCCTACATCATGCGACCATTCCAGTCCAGCCAATTGACGATCACGG
ACCTGCATGCGGCAGCCCCTTGGGACCCAGCTTACCACATGCATATCTGACGATCACGG
ACCTGCATGCGGGGGGAAGTTCACTGGGCTAATTGCGGGACTCTTGTTTGACGATCACGG
ACCTGCATGCGGAAGAGCACATGCATCCTTCATGGGAATTTAAAGGAGCTGACGATCACGG
ACCTGCATGCGGGAAAGAGTGCTCACCGCAGTTCCATTCTCCCGCAGTGACGATCACGG
ACCTGCATGCGGCAACGTGGAGAGCATCCAGTGGCGGGACATAGTCATGACGATCACGG
ACCTGCATGCGGGCAGTGACTTTCTCAGCAACATGTCGATGGACTTCTGACGATCACGG
ACCTGCATGCGGCAGAACCACCTGGGCAGCTGTAAGTGTCGCATACATGACGATCACGG
ACCTGCATGCGGCTCTACAGCACTGGGGCAGGGGAGAGAAGCCATGTTTATGACGATCACGG
ACCTGCATGCGGCCCTGCTGACACTGGCCAGACTCCTACATGCTTCTTGACGATCACGG
ACCTGCATGCGGCTGGGCACTGCCATGGAGAGGAAGTCTGTCCATGTTGACGATCACGG
ACCTGCATGCGGTTCCTTGAATACTGGTGGCCTGGTCCCTGTCCCATTGACGATCACGG
ACCTGCATGCGGGGACTCTGAGACGCAAGCTCACACTCACCCAGCTCTGACGATCACGG
ACCTGCATGCGGCTGCGCCCGTTCCTGATCGCTTGGACTTTCTGTTCTGACGATCACGG
ACCTGCATGCGGAGTAAGAAGTGATCACCATTTCCTGCTTCTTTGTTTCTCCATGACGATCACGG
ACCTGCATGCGGCAACTGTGCAGTGGATGCCTGTTTGTTTTCTGCCCTGACGATCACGG
ACCTGCATGCGGATAGAGCTGACGTGAATCTTCAAAATCATCAACTACAGGGCTGACGATCACGG
ACCTGCATGCGGTTTTAATGAACCTGGAACGGTTATGGGGCCAGTGTTGACGATCACGG
ACCTGCATGCGGCAGGCAGCCCACAATGGCAGGTCTCCATGTTCTGTATGACGATCACGG
ACCTGCATGCGGCAACAACTGTGGGAAAGACCCACAGAGAAAGTGCTTGACGATCACGG
ACCTGCATGCGGACTACAGGGCATTGAACTATAGGCCAATATAGCATTGCTTTTGACGATCACGG
ACCTGCATGCGGAAAAGGAATTAGGTATGTTGGGTTAGTCGCATTGGAGAGTGTGACGATCACGG
ACCTGCATGCGGAAATTCACCATCGACCTGATACCTGAAATTTCCTCCTTACCTGACGATCACGG
ACCTGCATGCGGGTGGTAAAGGAAGCTCCTCTCTTGACTGGTGCTTTTGACGATCACGG
ACCTGCATGCGGATGGCTACACGTTCCTGCTCAGAATGGATCTCATTTGACGATCACGG
ACCTGCATGCGGTCACAGTGTGCCTAGCTTTTATGTTTATGTTGAAGTTGGGCTGACGATCACGG
ACCTGCATGCGGTAATGGCTACCAGGAAAATATTGTGCAATGTCTGATTGCCATGACGATCACGG
ACCTGCATGCGGAAAAGAAAGCATCACACCTTCCTTTGCAAAGTATTTGGGTTGACGATCACGG
ACCTGCATGCGGAAAAGTTGAAAGAATGGGATTGCACGCTACAGATCTAGCTTGACGATCACGG
ACCTGCATGCGGGCTTTTAGCACGCCTGCGTAGGACCTTGCTTTCTCTGACGATCACGG
ACCTGCATGCGGGTGATGCCAGCCTCCCCGGCCTTCATGTCTCTAAGTGACGATCACGG
ACCTGCATGCGGGCGGCAATTAGCGCCCTTTGCCTTGGTGGTATTCTTGACGATCACGG
ACCTGCATGCGGTTTCTGTTTGAAATCTAAATTCTTCCTGACTGCAGGGGACCTGACGATCACGG
ACCTGCATGCGGTCGGGACCCATGAACACCTCTAGTTTACTATGTCTTCATGACGATCACGG
ACCTGCATGCGGTCTGCATGACTGGACTCTTTAACAAATTTGGTGGTTAACCTTGACGATCACGG
ACCTGCATGCGGAAAAGAAAACACGACAGATGACAGGAAAACCTTTGCAGCTGACGATCACGG
ACCTGCATGCGGAGCTAAGGTTTTACACAGTCATGAAAGTGATCTGCACTGTTGACGATCACGG

ACCTGCATGCGGTTATTTACAGGCTATGACTTAACAATCCCAGAACGGCTGACTGACGATCACGG
ACCTGCATGCGGACATGCAGTCACTCAAGACTGGACACAGCAAGGAATGACGATCACGG
ACCTGCATGCGGGTCCATGCCAAAGGCTCAGCCAGACGAGACACTCTTGACGATCACGG
ACCTGCATGCGGTGGCAGGAGATGCCAGGGAATGCTCCAAGCCTAAGTGACGATCACGG
ACCTGCATGCGGTTCTTTTCTTATCGGGGTCTCAAGTGATTCTACAAACCAGCTGACGATCACGG
ACCTGCATGCGGAGATTGTCTTCATTTATTGAATGTGCTTAACTCAGGCCCGGTGACGATCACGG
ACCTGCATGCGGAAAGGGCGTCATCAGTTTCTCATCATTTCACTGAGATATGCTGACGATCACGG
ACCTGCATGCGGAGTCAGTGAACAGCCTCAGACCCATGTGTGACCGCTGACGATCACGG
ACCTGCATGCGGCCCTCTCTTCCTTCACTTGCTTAGGTGATTGGATTTGTTGACGATCACGG
ACCTGCATGCGGACTTTATTACAGGGTCAGATGTGAACCAGTAGGTGAAGGATGACGATCACGG
ACCTGCATGCGGCAGTCTTGCAAATCTCACCGCATGCAGTTAATCCATGACGATCACGG
ACCTGCATGCGGGGGTGGGCTATTTTGGGAGCTTCAGCCTATCACAATGACGATCACGG
ACCTGCATGCGGCTGAACACAGCACACTTTCCCAGGAATGGAAGACTTGACGATCACGG
ACCTGCATGCGGTCCTCTCCCTCCCTGGCTTTCTTTGGTACCCACCTTGACGATCACGG
ACCTGCATGCGGCACAGAAGCCTGAGCACGGGTTCTCATGGGGACTTTGACGATCACGG
ACCTGCATGCGGTTCCATGTGGACCCTGCTTTACGATGGAGAGGGCCTGACGATCACGG
ACCTGCATGCGGATTCTCCTAGGTATGGTTGTCTGGCTCAGCCTCTCTGACGATCACGG
ACCTGCATGCGGGGGGCTGTTAGGCCCTTCTAAACACTACTTCCCAGTGACGATCACGG
ACCTGCATGCGGTTTTGTTCTTTGTATGTGCTTTCTGCATTGCCCAAGATGCTGACGATCACGG
ACCTGCATGCGGAAAAGAGGCAATCAGAAAAGGGCATGGTTTGACTTAGTTTGTGACGATCACGG
ACCTGCATGCGGATGATCCTACCCTCACTCTTCAGCTCACAGGGAACTGACGATCACGG
ACCTGCATGCGGTTTTCAGTGACCAAAATCATCTGTGCCCAGCAGTGTGACGATCACGG
ACCTGCATGCGGGGGGAGCTGTCAATTAGCATTTGTCATAACAGACAGGATGACGATCACGG
ACCTGCATGCGGCTACATGCAAGTATTTGTCATGAATGTGCCTTGTTGCCATGACGATCACGG
ACCTGCATGCGGGCAGCACACTCTCTTTATTGTTTGACTTCGGCTATACCTTGACGATCACGG
ACCTGCATGCGGCTAGAGACTTGACACTGTGAGGTCCCTAAGAGACCCTGACGATCACGG
ACCTGCATGCGGGAAGCCCTTCCTAAGACTAAATCCAGAGCACTCTGTGTGACGATCACGG
ACCTGCATGCGGCAGGAATTGTGACTTGGTTGGCCTGAGCTGTTTCTTGACGATCACGG
ACCTGCATGCGGAGGGAGAAGACTGTGCGTGGGGTTGGTCCTAGGAAAATGACGATCACGG
ACCTGCATGCGGCCTCGCTGTATTGGGTCTGGCTCCTTTACACGGCATGACGATCACGG
ACCTGCATGCGGACCCTCAATTGTACTTTATAAGGGAAGCCCAATGTCCTTTTGACGATCACGG
ACCTGCATGCGGTTTTAGAGAAGGGTCTTTCTGACTCTGCAGAGGGCTGACGATCACGG
ACCTGCATGCGGGCCAGCTGGGTTTTCCCACACTAGTGGAACACTAGTGACGATCACGG
ACCTGCATGCGGGCTGCAAAGACAGTAACTTGGGCTTTCTGACGGGATGACGATCACGG
ACCTGCATGCGGCGCAAATTCCGAGACGAAGCCACGTGCAAGGACACTGACGATCACGG
ACCTGCATGCGGCCCCACTCATGCTCTACAACCCCACCACGTACCAGTGACGATCACGG
ACCTGCATGCGGATGGATGTGAACCCCGAGGGCAAATACAGCTTTGGTGACGATCACGG
ACCTGCATGCGGCTGCGTGAAGAAGTGTCCCCGTGAGTCCTCCTCTGTGACGATCACGG
ACCTGCATGCGGCCCTCTAACTGGTCAGGCATCCTTGTCCCGCTCTGTGACGATCACGG
ACCTGCATGCGGTCTCCTGCTGAGCCCTGGAGTATCCCATCTTGGAGTGACGATCACGG
ACCTGCATGCGGAGTCTTTGGGTGGATGTGTTTGCCTTGCTTGGAGGTGACGATCACGG
ACCTGCATGCGGTGGGCATGCCACTGTTTAGTGCTTGTGCCAAGCAGTGACGATCACGG
ACCTGCATGCGGAATACACCGAAATCAGAGAGTCTATCAGAAGACCTGCCTTGACGATCACGG
ACCTGCATGCGGACTTACATTTGTACTGGTGGCTCCAGGGTTTAGGGTGACGATCACGG
ACCTGCATGCGGTGAAAGAGTTTTAGAAGCCACTCTTGATCTCTAGAAGGGGATGACGATCACGG
ACCTGCATGCGGAACAGAGCTTTAAAATCCATAGGAAAACACTTCGAGCCTGATGACGATCACGG
ACCTGCATGCGGATACCAAGAGCAGATGGTTCACAGAAGAATCATCAATGTCCTGACGATCACGG
ACCTGCATGCGGCAAATGAATTACTTTAAAGGTGAGAGCAGGTGGAGGAGAGGTGACGATCACGG
ACCTGCATGCGGCAAAACAGCTGGCCCCTCAAGGGACCCAGTGTTTCTGACGATCACGG
ACCTGCATGCGGCTGCCATGATGAAACACCTGTATTGTCCACATTGCGTGACGATCACGG
ACCTGCATGCGGAAACTCTTGAACGGGATTCCTTCTCTATTTGCAACCTTTCATGACGATCACGG

ACCTGCATGCGGTTTGTCCTTAAAGTAAATAAAGCCAAAGGAGGATGGAGCCTTGACGATCACGG
ACCTGCATGCGGGAGCACCTGGTGCCACCGTCATCACCTTCCTTTCATTGACGATCACGG
ACCTGCATGCGGGCTCTCTTCCCCAGGTAATTATGTGGTGACAGATCACTGACGATCACGG
ACCTGCATGCGGACAGAAGATAAAGCTGTAAAGCTAGGTTAGGCAATGGGAAGTGACGATCACGG
ACCTGCATGCGGCAGCTCCAAGCGGCCCATGGGAAATAATGAGGAGATGACGATCACGG
ACCTGCATGCGGACGCAAGGTCAGTGTGAGGTGACAGGGATGGCATCTGACGATCACGG
ACCTGCATGCGGTCCTACACCGCCGTAGCCCCAAAGTGTACTATAGGTGACGATCACGG
ACCTGCATGCGGCTGCACTCTCCCCAGCCCCTTCAGTGTTTGTTGAGTGACGATCACGG
ACCTGCATGCGGGAATGAAGGATGATGTGGCAGTGGCGGTTCCGGTGTGACGATCACGG
ACCTGCATGCGGCCGGAATTCCTTCCTGCTTCCCTCTGCCTGTGGATTGACGATCACGG
ACCTGCATGCGGAAAACTGCACCTCCATCAGTGGCGATCTCCACATCTGACGATCACGG
ACCTGCATGCGGTGGTGATCAATAATCACCCTGTTGTTTGTTTCAGTGACTCCTGACGATCACGG
ACCTGCATGCGGTTCACACATACTCCTCCTCTGGATCCACAGGAACTTGACGATCACGG
ACCTGCATGCGGTTTTCAAAAGTGCAGTTTCTCCTGCAGGCAAAAGGGTGACGATCACGG
ACCTGCATGCGGGACACGTTAAGTCCAGGCTTGGGTCATTCACTGCGTGACGATCACGG
ACCTGCATGCGGGCCTCAGGCTTTTGAAACTGGACGCCAGAGCCTTGTGACGATCACGG
ACCTGCATGCGGGGCAGATAATCGCAGCAGGAGCCTCTTCGGGGTAATGACGATCACGG
ACCTGCATGCGGAAAAGAAACTCCTACGTGGTGTGTGTCTGAAGTCTTTCATGACGATCACGG
ACCTGCATGCGGTTTTGCTGATTCAGGCTTGGCCTGAAAACAGGACGTGACGATCACGG
ACCTGCATGCGGGACCTCCATGCCTTTGAGAACCTAGAAATCATACGCGTGACGATCACGG
ACCTGCATGCGGGCAGGACCAAGCAACAGTAAGTTGACCACAGCCAATGACGATCACGG
ACCTGCATGCGGTTTTCAGCTTTGACAGTGGAATAGACATGCAGGTGCTGACGATCACGG
ACCTGCATGCGGTATGGTAATGGTAATCATCAGTTTAGTGGTGTGGAGGAGGATGACGATCACGG
ACCTGCATGCGGAGCCTCGTTATGAGAACTGCTGCCCCTACTTGACTTGACGATCACGG
ACCTGCATGCGGTTAAAGAGAATAAGAGTGCGTGAAATATGCATTGCCTCCCATGACGATCACGG
ACCTGCATGCGGAACTCCCTTGGCTCTGAATCTCTGATACTAAATATGTGGCTTGACGATCACGG
ACCTGCATGCGGTGCTCTGAATTCTCTGGAATGCTTTCTTTGACCAAGATTCTTGACGATCACGG
ACCTGCATGCGGAAGAGATTTAGAGCAATTTTCTTGGATGGCTGGTATGAGCCTGACGATCACGG
ACCTGCATGCGGTCACATTTAATGAGAAGTCTGGTGATTCCAGCTCCTACTGATGACGATCACGG
ACCTGCATGCGGGGGGAGAATTCCTAATGGTATTGGTTACAGGCTCCTATGCTGACGATCACGG
ACCTGCATGCGGCCCCTGCCATCCAACACCTGAACATCAGTCTCTTCTGACGATCACGG
ACCTGCATGCGGGCAGATCTCTTTCCTGTGGCTCCTTCTCAATTACAGATGACGATCACGG
ACCTGCATGCGGCAGCCTGCATCATCGTGGTCCTGTAGGGGCTAGAGTGACGATCACGG
ACCTGCATGCGGCCATCGTGGTCCGGCAGGGGCTTAGTGGTTCTTATTGACGATCACGG
ACCTGCATGCGGCCCATCGTGGTTCAGGAGGGGCTAGTGGTTCTCACTGACGATCACGG
ACCTGCATGCGGCCCCTTCGGTCTGAATGACTTCCATCCAGTCATCTTGACGATCACGG
ACCTGCATGCGGATATACACATTGGACCACCCAATAGCATCCTAGTGTCATGTTGACGATCACGG
ACCTGCATGCGGTAATGACATTAAGGCATTTTATTCATCCTCCCCATCTGCCATGACGATCACGG
ACCTGCATGCGGAAAACACCTGCAGTTTTCAAAAGGTGCAGTGTGTGTGACGATCACGG
ACCTGCATGCGGCCTCCCACAGCATGACCTACCATCATTGGAAAGCATGACGATCACGG
ACCTGCATGCGGTAGTCAATCAAAGGTGGTCTGGAGAAACAAAGTTTTCAGGGTGACGATCACGG
ACCTGCATGCGGTTTCTTCTCTCCAATGTAGTGGTCAGTTTTCTCTTGCAGTCTGACGATCACGG
ACCTGCATGCGGGTCAGCCTGAACATAACATCCTTGGGATTACGCTCCTGACGATCACGG
ACCTGCATGCGGAAAACTGTTTGGGACCTCCGGTCAGAAAACCAAAATGACGATCACGG
ACCTGCATGCGGCCAGATCATTACCATTCAATGGGATGAATTTACCCTGAGGTTGACGATCACGG
ACCTGCATGCGGTTACCTTTTAGTTGTAGGTCACTCTCTGCTCATTTCAGCCTTGACGATCACGG
ACCTGCATGCGGATGAGCACTTTACATCAACACTTCCTGTTCTGGCTCTTGACGATCACGG
ACCTGCATGCGGCTCAGCTTTTGAAGCTGGTGAGAGCCTGGCCTGTGTGACGATCACGG
ACCTGCATGCGGGGCCCAATGGCTCCTATCAGCTGTGGGATTTGTCATGACGATCACGG
ACCTGCATGCGGCTCTGCTCCAGGCTGCCCTGGGTACCATCAAAACATGACGATCACGG
ACCTGCATGCGGGCCGTGGGAGCACAGAAAACATGGAGTCCCATCTGTGACGATCACGG

ACCTGCATGCGGGGCACATTTCAGTCTTCCCAGCAGCCAGCACAACTTGACGATCACGG
ACCTGCATGCGGCTTTGTCCTTCCAGTCACGGTCGGCCTCTGGGAAGTGACGATCACGG
ACCTGCATGCGGAGTCTGTGTCCTCCTCCTTCAGGGGTAGCCAGCATTGACGATCACGG
ACCTGCATGCGGGTCCCTGCTCTGTCACTGACTGCTGTGACCCACTCTGACGATCACGG
ACCTGCATGCGGGAGGCAGGGAATGCGTGGACAAGTGCAACCTTCTGTGACGATCACGG
ACCTGCATGCGGGAGGGGTAGGAGGTTATTTCTTTAATCCCCTTGCGTTGACGATCACGG
ACCTGCATGCGGTTTTGGGTTTTCTGTGGGGAGACGGGAAGTTGTTTTGACGATCACGG
ACCTGCATGCGGAAGACAAATGAGACAGTCATCAATGGTTCATCTAGCCAACATGACGATCACGG
ACCTGCATGCGGCCGTGGCCATTTGGGCTTTTCTTTGTAGTGCCCGATGACGATCACGG
ACCTGCATGCGGCCCCAGCATGTGTTGAACATCAAACAGTACCAGGGTGACGATCACGG
ACCTGCATGCGGAGTGCAGTTTGTGAATGCTCTGAGGTTCTTGATATTGATGTTGACGATCACGG
ACCTGCATGCGGAAATAGGAAAGTAAAACTGAAGGATAGAGGCCAAGGCCATGTGACGATCACGG
ACCTGCATGCGGAAAAGGGAAATTTGGTGAGCACCATAGGAATTACAGATGGCTGACGATCACGG
ACCTGCATGCGGACAGCTTGGATCCAGAAATATTTAGGAGCAGGATAAGAGGATGACGATCACGG
ACCTGCATGCGGTATTGATGCAATCCTTACAGCTACAAATGGACAGTGGTCTCTGACGATCACGG
ACCTGCATGCGGAGTTTTCAAGTGTTTCTTAAGAGGCAAGGTGATGAAAACGCTGACGATCACGG
ACCTGCATGCGGCCACGTGGGGAGCCCCATGTCCTTCCATTAGTGTATGACGATCACGG
ACCTGCATGCGGCCCTTCAGCAAGGGCAGTGACCCAGAGAAGAAGCATGACGATCACGG
ACCTGCATGCGGAAGGCACACACTTTGGAGAATAAACACTCCTTGTTCGCTGACGATCACGG
ACCTGCATGCGGTGCTGGAGGGTAGAACTATGCTTGACTACTAGGCATGACGATCACGG
ACCTGCATGCGGAGCTTCTTGCTTAACATTGATGTGTACATCTTGAGGGATGTTGACGATCACGG
ACCTGCATGCGGTTTTGATAACCATGAATCTGGTATTAGAGGCAGGGAACACCTGACGATCACGG
ACCTGCATGCGGATGAATGAAGCTCCTGTGTTTACTCAGAGAGAAGATGACCCTGACGATCACGG
ACCTGCATGCGGTACTATATAGTCCTGGAGTCCCAACTCCTTGACCATTACCTTGACGATCACGG
ACCTGCATGCGGTTTGAAGAGGTGATTTGTGTTCCTGCAATAATGTCTCAGGGTGACGATCACGG
ACCTGCATGCGGAGCCAAGGGAGTTTGTGGAGAACTCTGAGTGCATATGACGATCACGG
ACCTGCATGCGGCTCAGGCCATGAACATCACCTGCACAGGACGGGTATGACGATCACGG
ACCTGCATGCGGGAAGGGCCTTCACAGAAGCCGAACAGTGATGATGGTGACGATCACGG
ACCTGCATGCGGCCGGGCAGGGTGTCGGTGACATTAGCACACACATTATGACGATCACGG
ACCTGCATGCGGGCCTGCGATGAACATTCACTCTTTCTGCTGACACCTGACGATCACGG
ACCTGCATGCGGAGCTTATCAAATCCTCACATTTAACGGAGGCTGTTTTCACCTGACGATCACGG
ACCTGCATGCGGCCCCATCCCTGACCTAGTCAGCATTGCTTTATCGCTGACGATCACGG
ACCTGCATGCGGTCTTTTAAAGAATATGAGTGAGAATTCGGGCCCCTCTCACATGACGATCACGG
ACCTGCATGCGGTAGACATTCTCTAAATTGGGCAGAAGATGACAGGACTGTGTTGACGATCACGG
ACCTGCATGCGGTTTGAGGGATAGGCTGCCAGCGTGGCTGCTTACAAATGACGATCACGG
ACCTGCATGCGGTGGCGATATTGACATAGTCATCTAATTATGCTGGCTCTGGGTGACGATCACGG
ACCTGCATGCGGCACACACAGCCCTTGAGTGGACAAAACCAACATGATGACGATCACGG
ACCTGCATGCGGGAGAACTTAGCCAAGGGGAAAGCCTTTCCCTGCTGTGACGATCACGG
ACCTGCATGCGGGTTTTATTTCTGCTACTTCTGAAGTGTGGGGCACACAACTGACGATCACGG
ACCTGCATGCGGACAAATAGAAGGTAGCGTATGACTCAGTCCTTGATATGCCATGACGATCACGG
ACCTGCATGCGGGGAGAAACACTCATGTGGGTTTTCTTAAATTTGCCTCCCTGACGATCACGG
ACCTGCATGCGGTGCAAGAAAGACTATCTGACCATAAATACACATTTGGGCCATGACGATCACGG
ACCTGCATGCGGATCAAGATGGTTTTGCCAAGGAAAGATGCCCACAATGACGATCACGG
ACCTGCATGCGGCAGGGAGAACTTCTAAGCAACTAACAGTGACCATATCAAGCTGACGATCACGG
ACCTGCATGCGGTTTTCTCCACCTTGGTGCAGGGACCAGACAACTGTTGACGATCACGG
ACCTGCATGCGGGTCAAGACCTGCCCGGCAGGAGTCATGGGAGAAAATGACGATCACGG
ACCTGCATGCGGCATGTGTGCCACCTGTGCCATCCAAACTGCACCTATGACGATCACGG
ACCTGCATGCGGTTGTATTTGAGTCAGTTACTTAAGGTGTTTTGGTCCCCACATGACGATCACGG
ACCTGCATGCGGCCATGCCAGTAGCAACTTGCTTGTGAGCAGGCCTCTGACGATCACGG
ACCTGCATGCGGGGGCCTGTGTTGTGCAATGCTGCACATCACAACAGTGACGATCACGG
ACCTGCATGCGGCTGCCACAGTCTCCAGGGGCTTTTGCGTTTCTCTCTGACGATCACGG

ACCTGCATGCGGGCTACTTTGACTGGAATTAGCAGAAGCACTCTGATTGTGTTGACGATCACGG
ACCTGCATGCGGATAAAGCTGTCAATCAAACTGGATCCCACTCAACAATCAGATGACGATCACGG
ACCTGCATGCGGTGTCAGTGTTGATGGTCCGATGGTGATGAGGAGGTTGACGATCACGG
ACCTGCATGCGGGGAGGTGAGAGTCACAATGGTAGTGGCGATGATGGTGACGATCACGG
ACCTGCATGCGGGTGAGGAGGTGGAAGTCACGGTGGTGGCGATGATGTGACGATCACGG
ACCTGCATGCGGAGATTCTTCATATTTTCGCCTTTGGGTATACCAACATGCCATGACGATCACGG
ACCTGCATGCGGGCTCTGTTGGCCACTTTGTGAGCTCGATGAAGCATTGACGATCACGG
ACCTGCATGCGGTATATCAGTATCTCTGTGTCAGTGAGCATATGTTGCTGGGCTGACGATCACGG
ACCTGCATGCGGTTAGGGGAGGTCCAGAAAGTGATTGGGTTTTGGCATGACGATCACGG
ACCTGCATGCGGTTCACTGTCTGACTTTAGTCTCCCACTAAAACTGCATTTCCTGACGATCACGG
ACCTGCATGCGGTTGTCTCTCTCATTTCTCAGATGGGATGTATTGCCTTCTCCTGACGATCACGG
ACCTGCATGCGGAACACTTACAGGGGTTTCTTTAACAACTTGTGAACAGCAGCTGACGATCACGG
ACCTGCATGCGGATCAGAGCCCAGACTACAGCATAAGCAGCTGCTGATGACGATCACGG
ACCTGCATGCGGCTGTTTGGTGTCTGTAGGAGGTAGTGGGGTGGGCATGACGATCACGG
ACCTGCATGCGGGTAACGAGGTCTCCTGTATATTCCACCCCTACGAAGCTGACGATCACGG
ACCTGCATGCGGCTGTGTGTTTGGTTTATGAACTAAGCTCAAAAGCACCACATGACGATCACGG
ACCTGCATGCGGCCCCAAATTAAGAAGAGCAGTGTAGAGAACAGAGACCTGTGACGATCACGG
ACCTGCATGCGGAAACTGTTAGGATCAGATTATAGTGTTACACCAGGGCTCCCTGACGATCACGG
ACCTGCATGCGGCTCACATATTGAAATGTACTTGTCCATCTTTCTCCAGGCCATGACGATCACGG
ACCTGCATGCGGAACTATCATCCTGTAATCAAAGTAATGATGGCAGCGTGTCCTGACGATCACGG
ACCTGCATGCGGGATCCCTTCTCTTCTGCCGTCAGAGTTTCAGCTGGTGACGATCACGG
ACCTGCATGCGGCTGCCACTGAGCCTCATGCCTTCACGTGTCTGTTCTGACGATCACGG
ACCTGCATGCGGGGGCCGCCAGGTTCCCAAGAGTATCCTACCCATTTTGACGATCACGG
ACCTGCATGCGGCCAGGACTGACCTCTTCCTCCTCGCTGCCAGATGATTTGACGATCACGG
ACCTGCATGCGGGTTCAAAGCACAGAATTTGTCAGAAACCTGCAGGGATGACGATCACGG
ACCTGCATGCGGCTCCATGCTGCCAGCCTTCTCCGTAATTAGCATGGTGACGATCACGG
ACCTGCATGCGGCCCTCTGTTTGAAATTCTAGAGCCAGCTGTGGGACTGACGATCACGG
ACCTGCATGCGGAATGTATATTTCTCTTTCACTTCCTACAGATGCACTGGGCCTGACGATCACGG
ACCTGCATGCGGAGGTCTTGAAGGCTGTCCAACGAATGGGTAAGTGTTGACGATCACGG
ACCTGCATGCGGTCACAGCTCTGTGTCACATGGACCTCGTCAAGAATTGACGATCACGG
ACCTGCATGCGGTTTCTGACTGTCCTCTGTCCTGATCAAGTTTCTATGGCTTGACGATCACGG
ACCTGCATGCGGGCTCCACGTGCCAACTTTGTCCTCAGTGGAGGGAAATGACGATCACGG
ACCTGCATGCGGCGTTGCTCGATGCATGGCCTGCCTCTGAATTCCTTTGACGATCACGG
ACCTGCATGCGGTTTTCTGTCCACTTACCCCATTGGTCCCATCACAATGACGATCACGG
ACCTGCATGCGGGCCTGTGTGTGAGTGGCCTTTCTGTTCGATGACAATGACGATCACGG
ACCTGCATGCGGCCTCCAGCATAGGGGAGTGTTTCTCCTTGCTTTCTTGACGATCACGG
ACCTGCATGCGGGACACACTGCCCAGCAAAGGCAAAAGGGCTTCCTTTGACGATCACGG
ACCTGCATGCGGAGGTTGAAAAGTCTTATTCAAACTCACCAGGAAAGAGTGGTTGACGATCACGG
ACCTGCATGCGGGTTACTCTCGATGGCGTCTAGCCAGGAATCATGGATGACGATCACGG
ACCTGCATGCGGATTATACACCGAGCACCTGTTTGCCATTTTGGATGTTTGACGATCACGG
ACCTGCATGCGGTCCAAACATGAACCAAACTTCCAGGCCCCTCTGCCTGACGATCACGG
ACCTGCATGCGGATCTCTGGTAACATTTACAAAGTCCCTTCCTCACCACTGCTGACGATCACGG
ACCTGCATGCGGCCCGAGTTGACAGCCATAGCTCTCTCTCCTGCCACTGACGATCACGG
ACCTGCATGCGGGTGAGCTGCTAGGACACCCAGCAGAACTTCCCCACTGACGATCACGG
ACCTGCATGCGGTCCACACTGCAATCTCAGGGATCTTAGTCACGGGGTGACGATCACGG
ACCTGCATGCGGCTTTCCACCATGTCTCCACCTGGAAACCAGTCATGTGACGATCACGG
ACCTGCATGCGGTTTTCTTCTCCTCCTGTTCACCCGCCCTTACTCTTTGACGATCACGG
ACCTGCATGCGGGTGGCGCCCTATGGATATGCGCTCCATAGCAAATGTGACGATCACGG
ACCTGCATGCGGCTTTATATCTTACGGTATTCTAGTGAGCTGGCACATGTGGCTGACGATCACGG
ACCTGCATGCGGGTGGGAGGGAGGATTAAATGCACCCTACAGTCTGATGACGATCACGG
ACCTGCATGCGGTCCTATTCCTTTATAACCCCTTTCAAGCTCGTTCAGAGAGTTGACGATCACGG

ACCTGCATGCGGATTTCACACAATCCATGTGCTCATCTTAAAAGCCAAGGACTGACGATCACGG
ACCTGCATGCGGGCCAGAGGCAGTGCCTGGTCCATGTGTATGGACTATGACGATCACGG
ACCTGCATGCGGTGGCACTTCAATTGCATGGAAATACTCTTGGAATGAACAAATGACGATCACGG
ACCTGCATGCGGAAAATGAGGAAAAGTGTGCCTGGTAGGGGACTGGGTGACGATCACGG
ACCTGCATGCGGGAGAGCTTGAGAAAGTTGGAAACGTTGCCTTAGAAGCTGACGATCACGG
ACCTGCATGCGGGTGTCTCACTTTCCAAGATCATTCTACAAGATGTCAGTGCATGACGATCACGG
ACCTGCATGCGGCAACCTCACCCTTCCTTGTTCCTCCACCTCATTCCTGACGATCACGG
ACCTGCATGCGGCAGCCCCGAGAACGGGCCATTAGCAGTTGTGTATGTGACGATCACGG
ACCTGCATGCGGATGCAGAAAGAATCTCTGAATGTGCAGTTATACCCAGTTGGTGACGATCACGG
ACCTGCATGCGGTGACATGTTGGTACATCCATCCGAGGAAATGGCAATGACGATCACGG
ACCTGCATGCGGTGTTTCTAGGCTGCACCCTTCAATGTCCACAAAGCTGACGATCACGG
ACCTGCATGCGGCATCTGCTTAGGACCCGGTGCCTGTGTGTGCATAGTGACGATCACGG
ACCTGCATGCGGCACTGTGGCGAAGGCGAGAGATTCCTGCTTTGGAATGACGATCACGG
ACCTGCATGCGGGCCCAGTCCTGTGCTGGAAGCCATGTTCTCTGGAAATGACGATCACGG
ACCTGCATGCGGCAACCCAGCGGCTCATAAGCATAAGCGCGTGTGATTGACGATCACGG
ACCTGCATGCGGCCCAACCAAACGACCGCCATGCACAACTTCCCTACTGACGATCACGG
ACCTGCATGCGGCGGAGTTTTCAATCCAGTTAATAGGCGTGGAAACAGACATGACGATCACGG
ACCTGCATGCGGGAAAGGTAGCTGTTCAGTTAAAGAACACCTGTATCAGAGCCTGACGATCACGG
ACCTGCATGCGGTGTGTTTCTACCAACTTCTGTCAAGCTCTGTAGAGAAGGCTGACGATCACGG
ACCTGCATGCGGGTACATTTGTCCTTCCAAATGAGCTGGCAAGTGCCTGACGATCACGG
ACCTGCATGCGGCCCAGCTTGTGGAGCCTCTTACACCCAGTGGAGAATGACGATCACGG
ACCTGCATGCGGGCTCCCAACCAAGCTCTCTTGAGGATCTTGAAGGATGACGATCACGG
ACCTGCATGCGGCTCCGGTGCGTTCGGCACGGTGTATAAGGTAAGGTTGACGATCACGG
ACCTGCATGCGGGTCTGGTGGGGAGCCCAGAGTCCTTGCAAGCTGTATATTTGACGATCACGG
ACCTGCATGCGGACTTTACTCTTTGTTTCACTGAGTGTTTGGGAAACTCCAGTTGACGATCACGG
ACCTGCATGCGGTATAACCACAGTAATCAGTGGTCCTGTGAGACCAATTCACATGACGATCACGG
ACCTGCATGCGGGGCATCAGAATGCAGCCCAGCTGAAATGGGCTCATTGACGATCACGG
ACCTGCATGCGGATCCTCTTTGCATGAAATCTGATTTCAGTTAGGCCTAGACGTGACGATCACGG
ACCTGCATGCGGAATTCTGGATGAAATGATCCACACGGACTTTATAACAGGCTTGACGATCACGG
ACCTGCATGCGGTTATCTAAATAATCAGTGTGATTCGTGGAGCCCAACAGCTGTGACGATCACGG
ACCTGCATGCGGCTTCGGGGTGCATCGCTGGTAACATCCACCCAGATTGACGATCACGG
ACCTGCATGCGGGGGCAGCATGTGGCACCATCTCACAATTGCCAGTTTGACGATCACGG
ACCTGCATGCGGATCCCAGAAGGTGAGAAAGTTAAAATTCCCGTCGCTTGACGATCACGG
ACCTGCATGCGGAGGAAATCCTCGATGTGAGTTTCTGCTTTGCTGTGTGACGATCACGG
ACCTGCATGCGGGGTCCATGGCTCTGAACCTCAGGCCCACCTTTTCTTGACGATCACGG
ACCTGCATGCGGCATGTCTGGCAGCTGCTCTGCTCTAGACCCTGCTCTGACGATCACGG
ACCTGCATGCGGTGTTCACTTTCTATGTCTTTCCCTTTCTAGCTCTAGTGGGTTGACGATCACGG
ACCTGCATGCGGATAACTCCCTCCCCTTAGAGACAGCACTGGCCTCTTGACGATCACGG
ACCTGCATGCGGCCCATGCTGGTATCCACCCCAAAAGGCTGGAAACATGACGATCACGG
ACCTGCATGCGGAAAGGTGCCTAGAATATGATGGCGTGCAGTCTATAAACTGTGACGATCACGG
ACCTGCATGCGGCCAGTGGTGGGAAGATGCTGGCTGGAGTCTGACAGTGACGATCACGG
ACCTGCATGCGGGCCTCTTCTACACTGGCCTGGGCTTGCTGTGAGTTTGACGATCACGG
ACCTGCATGCGGGGTGGAAACCTTTGCTCTTGTCCCAACACAGAGCATGACGATCACGG
ACCTGCATGCGGCTCAGGCAGCGGACTAGGGAAGCAGAATCGAGGAATGACGATCACGG
ACCTGCATGCGGTTTAGTTGATCTTACAGCCAAGAAGGACAAAAGCCAGAGAGTGACGATCACGG
ACCTGCATGCGGTAATATCCTCCGCCTCATGTCTAACCCACAGAATACATAGCTGACGATCACGG
ACCTGCATGCGGTTAACATAACCACAAGAAATACAGGAAGACGGGCAATCTGTTGACGATCACGG
ACCTGCATGCGGGTGCTGTGTAGGAAAATCTTTGTGCACAATTTGCTTCCTTGACGATCACGG
ACCTGCATGCGGTCAGTGGTATTTATAAATGCAGTGTGAGGAGGGTTTGGATGTGACGATCACGG
ACCTGCATGCGGATTCTAAGACAATAGTTGTGCTTTGGGAAGGAAGCAGTGTTGACGATCACGG
ACCTGCATGCGGCCCCAGGACCTTTTAATTGGAGGAAATATGCTTCTGTGGTGACGATCACGG

ACCTGCATGCGGAGATAAGGTCAAGGCTTAAAAGTTAAGTGCACCCAACATCTTGACGATCACGG
ACCTGCATGCGGATCATTTTATTTCCTTGCCCCATGTTTCTTCACTGCGGTGACGATCACGG
ACCTGCATGCGGGTCAGAGGGCGACAAGCCTGGAGTGTTCCCTGAAGTGACGATCACGG
ACCTGCATGCGGGCAAAAGCCAGCAGTGCCCAGGCCAATTTCAACAGTGACGATCACGG
ACCTGCATGCGGGGAGTTAAATAGCACCTTAATCCTGTGGCAGGACAGCTGACGATCACGG
ACCTGCATGCGGTCATGGGGCCATGTGTGCTCTTAGAAAGACTCACATGACGATCACGG
ACCTGCATGCGGGCACGCATGCACGGCAGCAATGACTCCATACTCACTGACGATCACGG
ACCTGCATGCGGTGGCTTAGTGAATGAGTAAAGTTCTTAAAATGCAGGGGACATGACGATCACGG
ACCTGCATGCGGCCTGCCCTTCATTCATAAGGCTGGACGTACACCTCTGACGATCACGG
ACCTGCATGCGGGAACCAAAAGTCATGTCAGCAGTTAGGACAAAATAACAGGCTGACGATCACGG
ACCTGCATGCGGTTTCAAGGTCACAAAGCCTCAGGGACACTCCTGCGTGACGATCACGG
ACCTGCATGCGGGTGGGTGCTGGGGCTCCACGAAGATTGTTGTGGAATATGACGATCACGG
ACCTGCATGCGGCCAAGCATGCTTGCTGTAGGTCACGGTGCACGTTTTGACGATCACGG
ACCTGCATGCGGACTACTTCCAAGACAAACAGCCGAGAACAAAGCTCGTGACGATCACGG
ACCTGCATGCGGCTTTAGCTTCTGCGTACACCGAACGGGACACACGATGACGATCACGG
ACCTGCATGCGGGTGGGGAGGAAGTGATGGCCCAGTGGGTCTATCAGTGACGATCACGG
ACCTGCATGCGGTCAAGTCTCAACCAGATCTTGAGAAAATAGGAAGAGCCAGATGACGATCACGG
ACCTGCATGCGGGGGTTTCTTTGGTGTTATGGTTGTACAGCTTCCCAGTGACGATCACGG
ACCTGCATGCGGGGGGGAGAGATGTGATTTGTGCTTTCTGGCAATCCCTGACGATCACGG
ACCTGCATGCGGATAGGCTTTCCAGTTTAAATTTAGGGTAGGCAATGGAAGGGTGACGATCACGG
ACCTGCATGCGGAACGCAAAACAGATTTCTAGGTGTACTGTGTGTGTGTCTTGACGATCACGG
ACCTGCATGCGGACACACACTCTCCTTGAAAATAGAGAGCCTAAACACTCTGCTGACGATCACGG
ACCTGCATGCGGTTGACTTGCTTTGAGTCTTTGACCTTCATGACTTCAGTACATGACGATCACGG
ACCTGCATGCGGTGCCCTAGTCCCTGCTGGAATGTTGAAGAAGCAAATGACGATCACGG
ACCTGCATGCGGGCCCTCAGAGCACTTGCCACGTACTTGCCAACAGATATGACGATCACGG
ACCTGCATGCGGCGGGGCGGAGACTTGAGTCAACGTAAGAGCAAGTGTGACGATCACGG
ACCTGCATGCGGGATCCGACACTGCAGAGCGCCAGCTAGACCCTAAGTGACGATCACGG
ACCTGCATGCGGGTGTGCTAGGGGCTGACCAAGCCGTTCTTTCCTCATGACGATCACGG
ACCTGCATGCGGTACCAGGAGAGTTTCATTTAGATTGAAGAGTCGAGGAAGGCTGACGATCACGG
ACCTGCATGCGGTCCTCTGAGAAAGAGTCTGCTAAGGAAGGAGGAGGTGACGATCACGG
ACCTGCATGCGGGGGGCAGGGTTTAGGGCAAGGAAAGCGTGTGAGCATATTTGACGATCACGG
ACCTGCATGCGGCTCGCCCCACCTTGCCTTTGTCTCCTGTCTCCTATTGACGATCACGG
ACCTGCATGCGGGAGCAGCACCCTATCTCAGAGCCTGGCTCAGTGTGTGACGATCACGG
ACCTGCATGCGGTTCACTTCTGCAGAGAAACTAACTTGCCCAAGTCCATGACGATCACGG
ACCTGCATGCGGCACTCAAAACATAGGCATTGCTGAGATGTGAAAAGCAGCTGACGATCACGG
ACCTGCATGCGGTGTGGATGCTTTCTGCTACAGTCTGTGTGTTCTTTTCCTGACGATCACGG
ACCTGCATGCGGCATTCTCAGTCAGACAGAAAAGAGGGCCCCATTGTTGACGATCACGG
ACCTGCATGCGGGTGCCTGATTGCAAATAAATTTAGCTTCCTCAGCCCATGACGATCACGG
ACCTGCATGCGGAAATAAAGTCTGTATTTATGGCTCTGTCAAAGGAAGGCCCCTGACGATCACGG
ACCTGCATGCGGCCGGAATTAGCAGGGCAGCAGATGCCTGACTCAGTTGACGATCACGG
ACCTGCATGCGGGGGGCACAGCACAGAGAGACCACTTCTCTTTAGAATGACGATCACGG
ACCTGCATGCGGCCGGGCAGCCAGGCAGCCTTTAGTCACTGTAGATTTGACGATCACGG
ACCTGCATGCGGGAATGCTCTGTCCATTTCAAAACCTGGGACTGGTCTGACGATCACGG
ACCTGCATGCGGCCCAAATGCCCACCCATTTCCCAGAGCACAGTCAGTGACGATCACGG
ACCTGCATGCGGCACAGCCAGCGTTCCTGATGTGCAGGGTCAGTCATTATGACGATCACGG
ACCTGCATGCGGCCAGGGTGTTCCGGACCCCACACAGATTCCTACAGTGACGATCACGG
ACCTGCATGCGGTCATGATATTTTAAAACACAGCATCCTCAACCTTGAGGCGGTGACGATCACGG
ACCTGCATGCGGAAGATACTATCAGTTCCCAAACTCAGAGATCAGGTGACTCCTGACGATCACGG
ACCTGCATGCGGGACTCCTCCTTTATCCAATGTGCTCCTCATGGCCATGACGATCACGG
ACCTGCATGCGGCCTCTCTGTCATGGGGAATCCCCAGATGCACCCAGTGACGATCACGG
ACCTGCATGCGGCCATGAGTACGTATTTTGAAACTCAAGATCGCATTCATGCGTGACGATCACGG

ACCTGCATGCGGGGCTGCCTCCTGGACTATGTCCGGGAACACAAAGATGACGATCACGG
ACCTGCATGCGGCAATATTGGCTCCCAGTACCTGCTCAACTGGTGTGTGACGATCACGG
ACCTGCATGCGGTGCAGATCGCAAAGGTAATCAGGGAAGGGAGATACTGACGATCACGG
ACCTGCATGCGGGGGGGAGGGGAGATAAGGAGCCAGGATCCTCACATGTGACGATCACGG
ACCTGCATGCGGGGTCTGCGCTCCTGGGATAGCAAGAGTTTGCCATGTGACGATCACGG
ACCTGCATGCGGTGGATTCAATCAAGTTGATCTTCTTGTGCACAAATCAGTGCTGACGATCACGG
ACCTGCATGCGGTTTCTCTTTATTGAGTGCTCAGTGTGGTCTGATGTCTCTGTTGACGATCACGG
ACCTGCATGCGGTACTGTGGTGATTTGTAGTGGAGAAGGAATATTGCTTCCCCTGACGATCACGG
ACCTGCATGCGGCATTCAGGACTTGATAACAAGGTAAGCAAGCCAGGCTGACGATCACGG
ACCTGCATGCGGGGTGCTGGAGACAGGCACAGAACAACAAGCCAGGTATGACGATCACGG
ACCTGCATGCGGCTCACCCAACTCCAGGCAGATGTAAAAGGTGACCTTGACGATCACGG
ACCTGCATGCGGAAAACATCTGGAGCGCTCTTATGCCAGCATCTGCCTGACGATCACGG
ACCTGCATGCGGCTCCAGCATCATCGGGCATTGATATCTCAGCTGCATGACGATCACGG
ACCTGCATGCGGGGAAGTGGGGAGCGCTCATGCTTCTCTGAGCACAAAATGACGATCACGG
ACCTGCATGCGGTTTTGCCATTCTCATGGTCATAACCCGGGCCACAGTGACGATCACGG
ACCTGCATGCGGAGTAGAACACTCCTATCACTGTTGTTAGACAGTGGTCCTGTGACGATCACGG
ACCTGCATGCGGTAGACGTGAAATGTGGAGACACTGTGCAACCCACTTGACGATCACGG
ACCTGCATGCGGGGGCAGATGTTGCTGGGTGTGGAGCTACATCCACTTGACGATCACGG
ACCTGCATGCGGCCAGACCTGCTTCCCCAGCTCTCCTCCTGGTTATCTGACGATCACGG
ACCTGCATGCGGTTTATGGCTTACTTGAGTTTGTGCCTACCTGTCCCATGACGATCACGG
ACCTGCATGCGGCATGTGGGAGCTGCCCTTCCTCCGTTCATCAGAGGTGACGATCACGG
ACCTGCATGCGGGGGCCAACAGTCCACAGCTGTTCTTAATCATCTCCTGACGATCACGG
ACCTGCATGCGGCCCCAGCTCCACAAAGGTGACTCCTTACATGGTGGTGACGATCACGG
ACCTGCATGCGGAGAGGTGGTCGGGCCATCCGTGTGAAATGTGTATGTGACGATCACGG
ACCTGCATGCGGTGACCGTTTTCCTTAAGGGGCACGTAGTCTTGGCATGACGATCACGG
ACCTGCATGCGGTGTTCTGCCCCTTGTCTTTCTGGGAGGTAGGGAGGTGACGATCACGG
ACCTGCATGCGGATACCCAGAAACATCTTCCTAATCTACATCTGTGCCAACCCTGACGATCACGG
ACCTGCATGCGGCAGCATGACCATCACATGCCCGCCATTGTTCCTGATGACGATCACGG
ACCTGCATGCGGCTGCTCATGACCTGCTCTCCAGCGCTCCTTCTCATTGACGATCACGG
ACCTGCATGCGGGCTCACATTCCAGTTGGCCTGACCTAGATAAGTGGTGACGATCACGG
ACCTGCATGCGGAACTTAGTGCGTGTCCCATTTAAGCAGAAGTTACTGAAAGCTGACGATCACGG
ACCTGCATGCGGCTGAGTTTAAGTTTCCAGGGCCTGAAAGTTTTCCATGACTGACGATCACGG
ACCTGCATGCGGTTTGAAGTCCTTTTACATTACGGGCAGTTAACGCTTTGTCTTGACGATCACGG
ACCTGCATGCGGCTGGAGGGAAATGAAAACAGTGATTCCCCAAATTGTGACTGACGATCACGG
ACCTGCATGCGGAACGTTTGAAGTAAAACTACCCAGAAACACTTAGTGGCTGATGACGATCACGG
ACCTGCATGCGGGAAACTAAACTCCTGGCATCCTCAAAATGGGATTTATTGGCTGACGATCACGG
ACCTGCATGCGGAAATGTCCTGTGTTGACTCACAAAGGCACAAACTATCTAGGTGACGATCACGG
ACCTGCATGCGGTAAGTTTTCTTCTAAATGTTGATGGGAGAGCTGGCCACTTGACGATCACGG
ACCTGCATGCGGAAAACAGAGGTGTCTTGGGAGGAATCCATATGAGACCTGACGATCACGG
ACCTGCATGCGGAGTAGACCATGAGAGAGACATCCCTTGCCATCTACATGACGATCACGG
ACCTGCATGCGGTGCGGAGACACCACTTGCTTACTAGCCAGAAAGAGTGACGATCACGG
ACCTGCATGCGGCAGGTGCCTCCTAAATTCCCCACACAGGAGCTCACTGACGATCACGG
ACCTGCATGCGGTGGCTTTCATGCACTGGGATTAAGTTAGACTTAAGAAAGCCTGACGATCACGG
ACCTGCATGCGGTGTCTACTCTTCCTGGGATTTACAAGCCAGCTAGTAAATCCTGACGATCACGG
ACCTGCATGCGGCAGAATAAATCACACGGCACAGTCATCCAAAGATCCCGTGACGATCACGG
ACCTGCATGCGGCATGCCCAGGACTGCAGAGTCCTGCCATTCAGACTTGACGATCACGG
ACCTGCATGCGGAATGCAGATTACAGTCTCACAGAATTGAGTGTTCAGGCATGACGATCACGG
ACCTGCATGCGGCTGTAGTACAGCAAACAATATCAGTTTACAGTCCTGAGGCCTGACGATCACGG
ACCTGCATGCGGCACGGTAGCGGTGGGGCAGGGTTCTCAGAATGAAATGACGATCACGG
ACCTGCATGCGGCTGGCTTACACATGGCACTCTCTGACCACAACTGTTGACGATCACGG
ACCTGCATGCGGAATGCAGATGAACATCAAAGAAAACGTCAAAGGCTCCTTGACGATCACGG

ACCTGCATGCGGTTTTACAAGTACGTGGGCTACTTAATTTGGTCCAAGTCCATGACGATCACGG
ACCTGCATGCGGAAAAGCCCTAGGTGCTTTCACGGCTCTGCTACTGATGACGATCACGG
ACCTGCATGCGGGAAGGAGACTCCAAGCTGGTGGTGCCAGCACATTCTGACGATCACGG
ACCTGCATGCGGCCACTCAGGCCTATTCCTAGGTGCCAGGTCACGAATGACGATCACGG
ACCTGCATGCGGAACCACGCTGACAGATCGTGCTGTGTGCGTGTCATATGACGATCACGG
ACCTGCATGCGGGCACACAAGCAGGACTGTGAGAGAGTGAAAGTGACTGACGATCACGG
ACCTGCATGCGGGGCTGTCATGGAGAAGAGACCCTGGCTTGCTCTACTGACGATCACGG
ACCTGCATGCGGCTTTCCCATCCTCAGATGAGGACTCGGCAGAGCCTTGACGATCACGG
ACCTGCATGCGGCTGATCCCGTCTGCACTGGGCCAGAGAGGATGACTTATGACGATCACGG
ACCTGCATGCGGTTTTATTTGTTCTTGCGTATTCCACAAAGGGTCGCAGCTGACGATCACGG
ACCTGCATGCGGAAAAGACAAAGCTCACAGTGCTGTTGAAGCACATTCATGACGATCACGG
ACCTGCATGCGGATTTCTTATCTCCCCAAGTTTGCCAACAAATACAGAGCAGGTGACGATCACGG
ACCTGCATGCGGCATGCAAGGCACAGGGAGGGTGAGCTCCAAGTTTGTGACGATCACGG
ACCTGCATGCGGCGTGAAGGCACTTGAGGGTGGGGAAGTGACTCTGATGACGATCACGG
ACCTGCATGCGGGCAAGATGGTGGCCAAATTCAAGGTCGCTGCAAAATGACGATCACGG
ACCTGCATGCGGTGGAGAAGAGAGAATAGATTTGGCATTTGGAGGAAATGGTGTGACGATCACGG
ACCTGCATGCGGTGCATGGATGCCCCAGCTCAGAGCATGACTCTCAGTGACGATCACGG
ACCTGCATGCGGGGGTGCCCGTGCTGCACCACTTCTGTCTAGGAATATGACGATCACGG
ACCTGCATGCGGTAGCACTTACTCTATGCCTGCCTGGGAAGGTGGCATGACGATCACGG
ACCTGCATGCGGTTTTAGGGAGCTCAAGGCCACAGATGCTCTGAGATTGACGATCACGG
ACCTGCATGCGGGAGCGGAGGCTTCTCCTGGTGACCACTCTGCTTAATGACGATCACGG
ACCTGCATGCGGCCAGCACCCCAAGGGCTAAAAGGTTGAGGGGAGAATGACGATCACGG
ACCTGCATGCGGCTCAGGACCAGCCCAAAGTGGAGGTGAGAGGTGAGTGACGATCACGG
ACCTGCATGCGGCAGGGGAAGGGTGATGGTGGTCTTGGTCTCAGCATTGACGATCACGG
ACCTGCATGCGGGGTTCTGGTAGAGGTGGGTTATTTTGAAGATGATGAACCTTTGACGATCACGG
ACCTGCATGCGGAGTCACTGCTCCATGAGCTCTTCCACAGGAGATTTTGACGATCACGG
ACCTGCATGCGGACAAAATAGAACACAAACAATCCAGTTCCCGCCTCTCATGACGATCACGG
ACCTGCATGCGGCCTGGGAACACCCAGCCCTTCAAGGTCAAACACAGTGACGATCACGG
ACCTGCATGCGGTTTTGTAAGTGGAAGTTCCAGGCTGGCTTTTCAAATCCTGACGATCACGG
ACCTGCATGCGGCTGAGTTTCCTGAGAGGCCCCGCTGAGGCTTTGTTTGACGATCACGG
ACCTGCATGCGGGATATTTCTTCCTGCGACCTCTGCTCGGACCCTGGTGACGATCACGG
ACCTGCATGCGGGAGCTCACAGGCCGTATCGCAGCTCTTATCTTTGGTGACGATCACGG
ACCTGCATGCGGCAGTGGGACGGAGACCCATGCGTCAAGTCTCCTCTATGACGATCACGG
ACCTGCATGCGGAGTTCACATGGGATTCTCTCCTTGTCCCAAAGCTGTGACGATCACGG
ACCTGCATGCGGAACCCTCCAACTGATTACCTGAATTCCAGAATATGTCCTGTTGACGATCACGG
ACCTGCATGCGGCCTTTCCCACGCCTTTGGTGAAGACCGGTGTTCTGTGACGATCACGG
ACCTGCATGCGGTTTAAAGCCCTGGTGTGAGGAGTGGGGAAGGGCTGTGACGATCACGG
ACCTGCATGCGGGGCCAGAGGTCAGCTGGATTTGTTAGATTGACAGATGACGATCACGG
ACCTGCATGCGGGACCAAGGTGCATGCTCACGCTGTCTCATGCTCTCTGACGATCACGG
ACCTGCATGCGGCAAGCAGCCTCTGGCTTAGCAGGTGCATTTCCCAGTGACGATCACGG
ACCTGCATGCGGCAGGGCAATTAAAGCCATGGTCCCAGTAGTGGTCTTGACGATCACGG
ACCTGCATGCGGTGGGGTCTCAGGGTATTTGGTCTGTGCAGCCACATTGACGATCACGG
ACCTGCATGCGGGCTTCAGTCTCTGGACCCCAGGTCATCTAACGAGGTGACGATCACGG
ACCTGCATGCGGCACATCGCCAGTGCTCAGCACTGTCAGCTGCTATCTGACGATCACGG
ACCTGCATGCGGATGTGCCCAGTATACCAAAATCTCCGCTAAGCATTTAAAGATGACGATCACGG
ACCTGCATGCGGGGCAGAATGAAAGTTAGCAGTGGTGGTGAAACGAATGACGATCACGG
ACCTGCATGCGGGGGAATGTGCTCTGAGGGCCTCCTTGTGGGCTTAATGACGATCACGG
ACCTGCATGCGGCTGGTCCTCTGCATGGGGACTTGTGTGTGGCTCTTTGACGATCACGG
ACCTGCATGCGGCCACAGAGGACGCCTTCAACAAATGTGAAGAACGATGACGATCACGG
ACCTGCATGCGGCTCAGTTGAACCAGCAAGAGGCTGAGCCTAACTCTTGACGATCACGG
ACCTGCATGCGGTCGAATGGGCAGGAGTTCATGATATTTAATAAACAGAGGCCTGACGATCACGG

ACCTGCATGCGGTTGCTCTGTAAGAGACAGGGTACCAGGCAGAGAGCTGACGATCACGG
ACCTGCATGCGGAAGTCAGCATCGCAGGAGTCAAACGAGGCAGACAGTGACGATCACGG
ACCTGCATGCGGCTTGCCTCTGAAGGAGACCCAGGCTGCCAGAGTAGTGACGATCACGG
ACCTGCATGCGGCCAGTCCTCTTTTGGGAAGCGCTTCCTCGGCTTCTTGACGATCACGG
ACCTGCATGCGGCTTCCTAATCTGGAAGTGTTGTCCAGAGAAGAACCTGCTGACGATCACGG
ACCTGCATGCGGGAGACGTGGGTGTCAGGCAAGCGTCTCTTTTCCAATGACGATCACGG
ACCTGCATGCGGAAAACATTAAAGGCCTCACAAACGGCGCCCAAAGATGACGATCACGG
ACCTGCATGCGGCTAATTCTGCATAGCATCTTTGCGAGACCCTAGGTTCTTGACGATCACGG
ACCTGCATGCGGAAAACAATTTGCCAGAAAACTTGGGAATCCATCCACATCTTGACGATCACGG
ACCTGCATGCGGACAGCTTTTCCCTGCAGTCACACTACAGTGGGATCTGACGATCACGG
ACCTGCATGCGGCATACAGGAGCGGCAGAGTGGAGCAGGCTAGAGATTGACGATCACGG
ACCTGCATGCGGATTTCTGTCGTGCCCACTCTGTACTAGAAAGTACATGAACATGACGATCACGG
ACCTGCATGCGGAGAAACAATAATAAGAAGTCTGCAGAACTTGACCCCTCCCTGACGATCACGG
ACCTGCATGCGGCTGGCCCTTGCAGAGAGATGCAGGCTGCCATTCTTATGACGATCACGG
ACCTGCATGCGGGGACAGTTGGGCTCAGCAAGGTAGGCATCCGTCAATGACGATCACGG
ACCTGCATGCGGCAGCAGCCAGCAGGGAGAAAGGTGCATGTGACAAGTGACGATCACGG
ACCTGCATGCGGGTGGGTCAGGATCAGCCAGGGTCAGGGTAGCATTTTGACGATCACGG
ACCTGCATGCGGCTAGGAATTCACTCTGTTGGGCGCTGTGCTGGCTGTGACGATCACGG
ACCTGCATGCGGATTCCTTTCTTACTCTCAGAGCAGAGATTTCAATTGCAGCGTGACGATCACGG
ACCTGCATGCGGAAAACCCTGAACGACATTCCTTTGCACCAGCTTGGTGACGATCACGG
ACCTGCATGCGGCCCCACTTTGCAGATAAACCACATGCAGGAAGGTCTGACGATCACGG
ACCTGCATGCGGAGCCTGGCAAGTCCAGTAAGTTCAAGCCCAGGTCTTGACGATCACGG
ACCTGCATGCGGCAACTGGGCAGCAGAGCTCCTGCTCTTCTTTGTCCTGACGATCACGG
ACCTGCATGCGGTCATATACGAGCACCTCTGGACTTAAAACTTGAGGAACTGGTGACGATCACGG
ACCTGCATGCGGATGGAGAAAAGTTAATGGTCAGCAGCGGGTTACATCTTGACGATCACGG
ACCTGCATGCGGTCTTTCATGCGCCTTTCCATTCTTTGGATCAGTAGTCATGACGATCACGG
ACCTGCATGCGGAGGCTCAGAGCCTGGCATGAACATGACCCTGAATTTGACGATCACGG
ACCTGCATGCGGCGGATGCAGAGCTTCTTCCCATGATGATCTGTCCCTGACGATCACGG
ACCTGCATGCGGTCACAGCAGGGTCTTCTCTGTTTCAGGGCATGAACTGACGATCACGG
ACCTGCATGCGGCGACCTGGCAGCCAGGAACGTACTGGTGAAAACACTGACGATCACGG
ACCTGCATGCGGCGCAGCATGTCAAGATCACAGATTTTGGGCTGGCCTGACGATCACGG
ACCTGCATGCGGAAACTGCTGGGTGCGGAAGAGAAAGAATACCATGCTGACGATCACGG
ACCTGCATGCGGTTTAACACATGCAGGGGAGGATGCTCTCCAGACATTGACGATCACGG
ACCTGCATGCGGCAGGGTCTCCTGGTAGTGTGAGCCAGAGCTGCTTTTGACGATCACGG
ACCTGCATGCGGGGGAACAGTACTTGCTGGGACAGTGAATGAGGATGTGACGATCACGG
ACCTGCATGCGGATCCCCAGGTGATCATTAGCAAATGTTAGGTTTCAGTCTCTGACGATCACGG
ACCTGCATGCGGTCCCTGCAGGATATATAAGTCCCCTTCAATAGCGCATGACGATCACGG
ACCTGCATGCGGATTGGGAAAGGTCACAGCTGCCTTGGTGGTCCACTTGACGATCACGG
ACCTGCATGCGGGCTGTCAAGGACACCTAAGGAACAGGAAAGGCCCCTGACGATCACGG
ACCTGCATGCGGCACTGGCTCGGTTTCTCTTAGGGACCCTCACAGCATGACGATCACGG
ACCTGCATGCGGCCACCAACCCATGATGCTGGGCCCTGAAAACACACTGACGATCACGG
ACCTGCATGCGGCAGAGCAGCCCTGAACTCCGTCAGACTGAAATCCCTGACGATCACGG
ACCTGCATGCGGTTTTCCAACAGAGGGAAACTAATAGTTGTCTCACTGCCTCTGACGATCACGG
ACCTGCATGCGGATCTCTCACCATCCCAAGGTGCCTATCAAGTGGATTGACGATCACGG
ACCTGCATGCGGTTTTACACAGAATCTATACCCACCAGAGTGATGTCTGGAGCTGACGATCACGG
ACCTGCATGCGGGGTGAGTCATAATCCTGATGCTAATGAGTTTGTACTGAGGCTGACGATCACGG
ACCTGCATGCGGAAATCGAGTCCAGCTGCCGTCCAAAAGTCACTGGATGACGATCACGG
ACCTGCATGCGGGGGGTATGGGAGGGAAAGAGCTTAGGAAACGGCTCTGACGATCACGG
ACCTGCATGCGGTCCCTGCAAAGTCCAACCAAACTTTAACGTTAACCAAATGACGATCACGG
ACCTGCATGCGGCCATTAATGTTGCCATGAATTTGAAGTGAACCAGAGGGAGTGACGATCACGG
ACCTGCATGCGGGTGGCAGAAGAAGCTTAATGGGGAATAGTTCCGGTTGACGATCACGG

ACCTGCATGCGGAGAGAAATGAGGCTTAAGATGAACTACCCTGGCCCTGACGATCACGG
ACCTGCATGCGGTTATGTGTCAGAGAGAACGGCTTGACAAACACACACTTGACGATCACGG
ACCTGCATGCGGGTCCTGAGCCTACAGGGGCCTCCCATGTTGAGAAATGACGATCACGG
ACCTGCATGCGGGTGTTGCTGACACATTGTCTCTGACCGCTGTGCCATGACGATCACGG
ACCTGCATGCGGGGCATTTTCTGCTGAATTACCGCACTTGGTCCTTGTGACGATCACGG
ACCTGCATGCGGAATTTCACCCAGCAACTTACTGAAAGGCTGGAACCTGACGATCACGG
ACCTGCATGCGGCATGAACCTACCCCTTCACTGAGGAAAATAAGTTACCCCTGACGATCACGG
ACCTGCATGCGGGACAGGAGCAAGGGAGGAGTCGCCTCACCTCTCTATGACGATCACGG
ACCTGCATGCGGAACTGCAGGGTCCTGCCCCAAGCCATTTATTTTGTTGACGATCACGG
ACCTGCATGCGGCAATGCCCACACTCCATGCACAAGTGGAAGCCCTCTGACGATCACGG
ACCTGCATGCGGTCAAAGTCAGTGGCTTAGTGCCTTGATGTGGTCACTGACGATCACGG
ACCTGCATGCGGACCCATTCTCAGGAAGTCCGTTCCCACTGAAAACATGACGATCACGG
ACCTGCATGCGGTTGTGTGTTTTCAACATCATTGAGGCTGCCACGGCTGACGATCACGG
ACCTGCATGCGGAGATTATAATCACTGGCCTAGGCAGCCCACTGGAATGACGATCACGG
ACCTGCATGCGGAAATTAGGGGCCTAGTGTGTTGTACCAAGGAGCTAAATGACGATCACGG
ACCTGCATGCGGCACCTAACACTGGCTATACTGGCTCTCCTCTCTCTTGACGATCACGG
ACCTGCATGCGGGGGGGCAGGGGATGACACAAGAATCTATTTTCTTGCTGACGATCACGG
ACCTGCATGCGGCCCCAAACCATTGCTTTCCTTCCAATGTTAAGCTTGTTGACGATCACGG
ACCTGCATGCGGATTCTGTGTATTAATTCAGGTGGTTCCGTTTGGGAATGGCTGACGATCACGG
ACCTGCATGCGGAAAATGTGCAGTTTACAGCCCTGTTCAGAATCTGCATGACGATCACGG
ACCTGCATGCGGTCTTCCTCATTCTGCAGATACAGGTCCCTCAGAGCTGACGATCACGG
ACCTGCATGCGGAGGTGACTGAGTGTGTATCCTGTCTGGAGCATAATACTTGACGATCACGG
ACCTGCATGCGGACTGTTTGGGAGTTGATGACCTTTGGATCCAAGCCTGACGATCACGG
ACCTGCATGCGGATATGACGGAATCCCTGCCAGCGAGATCTCCTCCATGACGATCACGG
ACCTGCATGCGGGTACCATCGATGTCTACATGATCATGGTCAAGTGTGAGTGTGACGATCACGG
ACCTGCATGCGGGCCAAACAGCTGAGGCCTTTGCATCCCTGGAGAAATGACGATCACGG
ACCTGCATGCGGTGTCATCACATTACTTAAGGCAGGCACACAAATCCAGATGACGATCACGG
ACCTGCATGCGGTTTAAAGACACTTCTTGACTCATTGGGCAGTATGACCTGACTGACGATCACGG
ACCTGCATGCGGAAGTCTTCCGCAAGCCATTACACCAAAATATTCTATTCGCTTGACGATCACGG
ACCTGCATGCGGTGCATGGCAGTGAGTAACAACATAAGGCTAATAGAGTCAACTGACGATCACGG
ACCTGCATGCGGACCTTTTAAACCAGTGGATTTATGTGAAGTCTCTGCAGTGTTGACGATCACGG
ACCTGCATGCGGGATTGATGCTATTATTCTACCAGATTCACGAGTGCAGTGGGTGACGATCACGG
ACCTGCATGCGGCTCTGGAGGTAGCATTACATGCATGGGATGAGCATTGACGATCACGG
ACCTGCATGCGGAGTCTGGCCTAAAATAGCTTGATATGCTGTGGATGGAATGACGATCACGG
ACCTGCATGCGGAATAAATGTGATCCCTCAAGAGGCATGAGGATTTCCAGGTGACGATCACGG
ACCTGCATGCGGTTAATCTGGATTTAGATTGTTGGGTAGTCACATGCAGCAGCTGACGATCACGG
ACCTGCATGCGGTTGCATGCATGCCCACTGGGTATGTGCTGTACTGGTGACGATCACGG
ACCTGCATGCGGTTTTCTAAGTTGCATCTGGTTTCCTACCTGAAGGAATGCTTGACGATCACGG
ACCTGCATGCGGAAGGGTGGATGTTGAGTGAGGACCTTGGTGCAGGGTGACGATCACGG
ACCTGCATGCGGCACCCTGCAGTCAGGATAGTTCATGGAGAGCAATTTGACGATCACGG
ACCTGCATGCGGTCTGATTCAAATGTACAGTGCTGGCATGGTCTTTAAACAGTTGACGATCACGG
ACCTGCATGCGGAACCAGTACTAGCTGGCCAAGACAGAAAAGTCTACCTGACGATCACGG
ACCTGCATGCGGTTTTCTCATTCCTTCCCCAGGCTGGATGATAGACGTGACGATCACGG
ACCTGCATGCGGCAGATAGTCGCCCAAAGTTCCGTGAGTTGATCATCGTGACGATCACGG
ACCTGCATGCGGCCCCAGCGCTACCTTGTCATTCAGGTACAAATTGCTGACGATCACGG
ACCTGCATGCGGAGTCTGTGCTTCCATTGGGAAGAGTCCCTCTAATGTGACGATCACGG
ACCTGCATGCGGAGCATCTCATGTCACTGTGTTCTGTCACATGCCAGTGACGATCACGG
ACCTGCATGCGGCCTGGCCTCCCTGTGTCCCAGATCGCATTATTAAATGACGATCACGG
ACCTGCATGCGGCATTAGAGCAAGCCTCAGTAAGGCGCAGGCCACATTGACGATCACGG
ACCTGCATGCGGTTTCTGTTTCCTCAGAAGCTATTTGAATCTCATGTAGGGGCTGACGATCACGG
ACCTGCATGCGGAGCATCAAAGGATGGTTCATGTTTTATTTTAAGGCACCCACTGACGATCACGG

ACCTGCATGCGGATCATGTCATGAGGGGAGGCAGCTATAATTTAGAGAACCATGACGATCACGG
ACCTGCATGCGGGGGGATTTCATTATAACAAAATTGGCAAACACACAGGCACCTGACGATCACGG
ACCTGCATGCGGTGCTGGCAATAGACCCCTGCTCCTATAGCCAAGAATGACGATCACGG
ACCTGCATGCGGGTGGAATAGCATCTCTACGGGCCATTCTAATAGCCTCTGACGATCACGG
ACCTGCATGCGGTCCAACTTCTACCGTGCCCTGATGGATGAAGAAGATGACGATCACGG
ACCTGCATGCGGCACGTCACGGACTCCCCTCCTGAGCTCTCTGGTATTGACGATCACGG
ACCTGCATGCGGTTGTTGCATTGCAACAACTTGATTGTAAGCCTTTTAGGTCCTGACGATCACGG
ACCTGCATGCGGAAATTAAACACCTTCACAATATACCCTCCATGAGGCACACCTGACGATCACGG
ACCTGCATGCGGACCTGCATTCAGGAAAAGTGGATGAGATGTGGTACATGACGATCACGG
ACCTGCATGCGGTTTTCAGAGTGCAACCAGCAACAATTCCACCGTGGTGACGATCACGG
ACCTGCATGCGGGAACACCTTATAAGCCAGAATTTACAGCTCTCCACTATGGCTGACGATCACGG
ACCTGCATGCGGGCAAAGCTGTCCCATCAAGGAAGACAGCTTCTTGCTGACGATCACGG
ACCTGCATGCGGTGAGTGGCTTGTCTGGAAACAGTCCTGCTCCTCAATGACGATCACGG
ACCTGCATGCGGGGGGAAACAGTGGCAGATTTGCAGACACAGTGAAGTGACGATCACGG
ACCTGCATGCGGGGCGTAAGGAGCAGATAAACACATGACCGAGCCTGTGACGATCACGG
ACCTGCATGCGGCACAAGCTCTTTGTTGTGTCTGGTTGTTTGCTGTACCTGACGATCACGG
ACCTGCATGCGGACAGATTTGATCCCTGTTCTCTCTGCTGGCTCTATCTTGACGATCACGG
ACCTGCATGCGGATGTCCATTACTTTGAGAAGGACAGGAAAGAACCCACTTTTGACGATCACGG
ACCTGCATGCGGTCCTGCATGGGATGGTGCTTTGCTGATTACTTCACTGACGATCACGG
ACCTGCATGCGGTTTCTTTCCACTTTCAGAATACATAAACCAGTCCGTTCCCATGACGATCACGG
ACCTGCATGCGGAAAGGCCCGCTGGCTCTGTGCAGAATCCTGTCTATTGACGATCACGG
ACCTGCATGCGGCACTGCAGTGGGCAACCCCGAGTATCTCAACACTGTGACGATCACGG
ACCTGCATGCGGGCCCAGAAAGGCAGCCACCAAATTAGCCTGGACAATGACGATCACGG
ACCTGCATGCGGCCCTGACTACCAGCAGGACTTCTTTCCCAAGGAAGTGACGATCACGG
ACCTGCATGCGGCCAAGCCAAATGGCATCTTTAAGGGCTCCACAGCTTGACGATCACGG
ACCTGCATGCGGCAGTGAATTTATTGGAGCATGACCACGGAGGATAGTATGAGTGACGATCACGG
ACCTGCATGCGGAAAATCCAGACTCTTTCGATACCCAGGACCAAGCCTGACGATCACGG
ACCTGCATGCGGCCTCCATCCCAACAGCCATGCCCGCATTAGCTCTTATGACGATCACGG
ACCTGCATGCGGGACCCACAGACTGGTTTTGCAACGTTTACACCGACTGACGATCACGG
ACCTGCATGCGGTAGCCAGGAAGTACTTCCACCTCGGGCACATTTTGTGACGATCACGG
ACCTGCATGCGGTCCTTTGTCTTCAAACTGTGAAGCATTTACAGAAACGCATCTGACGATCACGG
ACCTGCATGCGGTTTTACTTCAATGGGCTCTTCCAACAAGGAAGAAGCTTGCTGACGATCACGG
ACCTGCATGCGGAGAGGATGCTTGATTCCAGTGGTTCTGCTTCAAGGTGACGATCACGG
ACCTGCATGCGGCTTCCACTGCAAAACACTAAAGATCCAAGAAGGCCTTCTGACGATCACGG
ACCTGCATGCGGCCCCAGCAGGCCGGATCGGTACTGTATCAAGTCATTGACGATCACGG
ACCTGCATGCGGGGCAGGTACAGTAGGATAAGCCACTCTGTCCCTTCTGACGATCACGG
ACCTGCATGCGGCTGGGCAAAGAAGAAACGGAGGGGATGGAATTCTTTGACGATCACGG
ACCTGCATGCGGAAAATGTCCCCACGGTACTTACTCCCCACTGATGGTGACGATCACGG
ACCTGCATGCGGACCAGTGGTTTCCAGTCATGAGCGTTAGACTGACTTGACGATCACGG
ACCTGCATGCGGACATCAAATAATAACTCGGATTCCAGCCCACATTGGATTCATGACGATCACGG
ACCTGCATGCGGTCAGCATTTGGACCAATAGCCCACAGCTGAGAATGTGACGATCACGG
ACCTGCATGCGGTGGAATACCTAAGGATAGCACCGCTTTTGTTCTCGCTGACGATCACGG
ACCTGCATGCGGAACGTATCTCCTAATTTGAGGCTCAGATGAAATGCATCAGGTGACGATCACGG
ACCTGCATGCGGAAAATGAAGCTGCTCTGAAATCTCCTTTAGCCATCACCCTGACGATCACGG
ACCTGCATGCGGTTTTACTCAAAGAGTATATGTTCCCTCCAGGTCAGCTGCCTGACGATCACGG
ACCTGCATGCGGAAGTGTCTCTGCCTTGAGTCATCTATTCAAGCACTTACAGCTGACGATCACGG
ACCTGCATGCGGCTACAATTGGAAGATTGGAAGATTCAGCTAGTTAGGAGCCCTGACGATCACGG
ACCTGCATGCGGTTTTCCTAATCTGTGTGTGCCCTGTAACCTGACTGGTGACGATCACGG
ACCTGCATGCGGACAGTGTTTTAAACTCTCCTAGTCAATATCCACCCCATCCATGACGATCACGG
ACCTGCATGCGGTTTTGACTCCCAGATCAGTCAGAGCCCCTACAGCATGACGATCACGG
ACCTGCATGCGGCTCCACTGCTAAAGTCCACATAAGGCTGAGGTCAGTGACGATCACGG

ACCTGCATGCGGTCACCCTAAACAACCTGCTCCCTCTAAGCCAGGGGTGACGATCACGG
ACCTGCATGCGGGGGGATTTTGAGCTATCATCTCTGCACATGCTTAGTGATGACGATCACGG
ACCTGCATGCGGATTCACCTCAGGACATGCAGAAATATTTCAGTCAGAACTGGTGACGATCACGG
ACCTGCATGCGGGAAACAGAAGGACCTACATTCTGCTGTCACTTATGTGTCATGACGATCACGG
ACCTGCATGCGGAGAAGCAGATGATCGATGAGGCAGGTCAGTTGTAATGACGATCACGG
ACCTGCATGCGGAAGATTGTGCTTCCCAAAATAGTTCTCACTTCATCTGTCCATGACGATCACGG
ACCTGCATGCGGCCCTTAGCCTCTTCACGGATCTGGCGACTGTGATGTGACGATCACGG
ACCTGCATGCGGTTTCACCTAACAGCCCTGATCAGTCAGTACTCAAAGCTGACGATCACGG
ACCTGCATGCGGCTGGCCTTAATACCCTACAGAAAGCCTGTCCATTGGTGACGATCACGG
ACCTGCATGCGGCTGTTTCTTCCTCAGTCAGTTCCTGGAAGACCTTACCTGACGATCACGG
ACCTGCATGCGGCCATGACCCCAGCTTCAGATGTGGTCTTTGGAAACTGACGATCACGG
ACCTGCATGCGGAGAGGTCGAAGGAAAGTAAGGAGCTGAGAGCTCACTGACGATCACGG
ACCTGCATGCGGATTCATAGGTGCCGCCAGCCTTCGTGCATCTTCTTTGACGATCACGG
ACCTGCATGCGGGCATCATCTCTAAGGAGCTCCTCTAATTACACCATGCCTGACGATCACGG
ACCTGCATGCGGCGTCACCCCATGAGGGATCAGAGAAGGGATGAGTCTGACGATCACGG
ACCTGCATGCGGGAGCACTGTGGATGCATCCTATTGCACTCCAGCTGTGACGATCACGG
ACCTGCATGCGGATGACACCAAAGCTTAGGTGTTTGCTGAAAGTTCTTGATGTGACGATCACGG
ACCTGCATGCGGTTGTGACTTACCACCCCTGCCTCACAACTGCAGACTGACGATCACGG
ACCTGCATGCGGATAAGGGGACTATGGATTGCTTAGCAGGAAAGGCATGACGATCACGG
ACCTGCATGCGGGCTGCCCTTGGGAATCTTCTGGTCCCAACCAGAAATGACGATCACGG
ACCTGCATGCGGGACTGTGGCTTGATTTTCTCAGGTGCAGCCCAGCCTGACGATCACGG
ACCTGCATGCGGTGCAACATTCATCAAAGTTTCTAGAACCTCTGGCCTAAAGGTGACGATCACGG
ACCTGCATGCGGTTGTGTAGGATGAAGCTGGTGGGTGATGGGAACTCTGACGATCACGG
ACCTGCATGCGGTTATTTCTAGGGTCTTCAGTTGTACAAGACTGTGGGTCTGTTGACGATCACGG
ACCTGCATGCGGCCCCGTCAGAGTAGAATAAAAGGCTGGGTAGGGTATGACGATCACGG
ACCTGCATGCGGGAGATTCCCATGTGCAGTGGAGAGAACAATCTGCATGACGATCACGG
ACCTGCATGCGGGTCACTGATAAGCCTGAGACTTGGCTCATTTCAAAAGCTGACGATCACGG
ACCTGCATGCGGGTTCAATTCATCCTCACCAGCAGTTCAGCTGGAAATGACGATCACGG
ACCTGCATGCGGAAGTTATAGAAACCAGTCTCTTCCCTCCTTTGTGAGTGAGCTGACGATCACGG
ACCTGCATGCGGTGCTATTCCACGTAGGCAACACCTGTTGAAATTGCTGACGATCACGG
ACCTGCATGCGGTTTATTGCAACATAATGATCTGCTCACATTTCCTTGCCTGGTGACGATCACGG
ACCTGCATGCGGGGGCTGTAAAACCTTACAGAACAGAAATCCTTGCCTCTTGACGATCACGG
ACCTGCATGCGGTCTTTGTTTAAAAGGGTTAAGTTGAGGCAAGAGGAAAGCCCTGACGATCACGG
ACCTGCATGCGGAAAAGGCACAACCTCATTGGGGAGCTAAGCTAGGTTGACGATCACGG
ACCTGCATGCGGTTAAAAGATGATGGAAAGCACATTTAGCTTGGTCTGAGGCATGACGATCACGG
ACCTGCATGCGGAAAATCCAGTTGCATGCCATACTCTCATCATCTGCCTGACGATCACGG
ACCTGCATGCGGAACTAGACATGTCTGGAGAGCCTAATAATGTTCAGCACACTTGACGATCACGG
ACCTGCATGCGGTTGGTTAGTTCACCAACAGTCTTACCAAGCCTGGGTGACGATCACGG
ACCTGCATGCGGGCCTAGGGATTAGCCACAAGGACATGGACTTGGAGTGACGATCACGG
ACCTGCATGCGGGCAAATTCTGCAGGTGTATGTGATTCTCAGGCCTAGATGACGATCACGG
ACCTGCATGCGGGAGCTAAGACACAAAGACCTCCACATCTGTCGCTGTGACGATCACGG
ACCTGCATGCGGAGAGTCAAGAACCTGAACAGAGTTTCCATGAAGGTTCTCCTGACGATCACGG
ACCTGCATGCGGTATAATATTCTTGCTGCTTATGCAGCTGACATTGTTGCCCTTGACGATCACGG
ACCTGCATGCGGCCCTAAAGCAACCAAGTAGCCTTTATTTCCCACAGTGATGACGATCACGG
ACCTGCATGCGGAAGAAAACGCTGGCCTATCAGTTACATTACAAAAGGCAGATGACGATCACGG
ACCTGCATGCGGAAAACTATGTAGCAGGCAGTGTGTTTTCCTTCCATGTCTTGACGATCACGG
ACCTGCATGCGGAAGTCTGTCCTATCTGAATTCCCAGCAGAAGCACTAAGATGACGATCACGG
ACCTGCATGCGGAGCTCCACCCTATCACCTAGCAGATAAAACTATGGGGTGACGATCACGG
ACCTGCATGCGGCCTGTCGCTAACCAGCCTCTCCCACCATCACCATCTGACGATCACGG
ACCTGCATGCGGCAGCCCCTCGGGCTCTCAGATGCATTCCAACTCAATGACGATCACGG
ACCTGCATGCGGTTTTCTCCTTCACAGAGCCAAGAGGGAATTCCCTGTGACGATCACGG

ACCTGCATGCGGTTTTCTATGGCAAGCCCAGGTTTTATTTCTTCAGAGTACCATGACGATCACGG
ACCTGCATGCGGAAAATCAGAGGAATTAAGTGGGTCCCAGCATGGGCTGACGATCACGG
ACCTGCATGCGGCATTCAGCTGCCCCACTCACCGCCAACCTCTCTTATGACGATCACGG
ACCTGCATGCGGCTCTCACTGCCGTGTTTAATGTAATGAAGCCACTACCTTGACGATCACGG
ACCTGCATGCGGCAGGGGAGGGTTCTAGGGAGTCCGAGCTCAGACTGTGACGATCACGG
ACCTGCATGCGGAGAGGAGAGAACAGCTTCCCAAAACCCAGCATCAATGACGATCACGG
ACCTGCATGCGGCCCTGGGGCTCCTGGTGGCTCTGACTTCTGTAAAGTGACGATCACGG
ACCTGCATGCGGGTGGCTGCACAGTGCCCTGCCTTGTTTCTTTCACATGACGATCACGG
ACCTGCATGCGGCACACCACCCCTAATGCTCGCAGCGCTGACTCTAATGACGATCACGG
ACCTGCATGCGGTTGTTTCCAGGAAGATCGCAGCTTTGCTTTTGTGCTGACGATCACGG
ACCTGCATGCGGTGTTCCTGGTGAATGCCATCCGTTGTCTATGCACATGACGATCACGG
ACCTGCATGCGGAAGCAGGACTTCAGGATAGCCAGGTCCACGAACAATGACGATCACGG
ACCTGCATGCGGTGAGAAAATCCGGAGCTTCCCAGCATTCTGCTCAATGACGATCACGG
ACCTGCATGCGGGCTGCCTGGGAACACCTCAGGCACATGCAGGTTTAATATGACGATCACGG
ACCTGCATGCGGGGGGTTTAGTCCATATGGAGGAAGTCCAGATTCTGCTGACGATCACGG
ACCTGCATGCGGAGATGAGACAAGCATGGGGAGCATTCTGAGGTGCTTGACGATCACGG
ACCTGCATGCGGCATCCCACTGAACAGCCTTCACCTCAGGGCAGACTTGACGATCACGG
ACCTGCATGCGGCAGTGGCACCACTGATTTGTCACTGACCGTCACGATGACGATCACGG
ACCTGCATGCGGTCCCTTTGGGCCAGTAATCCTATAAAAGTCATCATACCTGTTGACGATCACGG
ACCTGCATGCGGCTGGGGAGATGGGCGATTTCCCTAGAATGGAGAGTTGACGATCACGG
ACCTGCATGCGGCCCTCATAGGGCTCAGGTCCCACTGCTCCAGAAAGTGACGATCACGG
