## Supplemental Table 3 for "BRCA1-A and LIG4 complexes mediate ecDNA biogenesis and cancer drug resistance"

**Table S3: Summary of high-throughput sequencing data**

| File | Genotype | Sequencing type | Sequencing Method | Number of reads | Mean read length | Total bases | Coverage | Mapped reads | Mappability |
|---|---|---|---|---|---|---|---|---|---|
| **CRISPR Screen** | | | | | | | | | |
| no_cut_1.fastq.gz | no CRISPR-C controls | DNA-seq | Amplicon seq | 39,530,878 | 75 | 2,964,815,850 | 1048.0 | 33,340,025 | 84.34% |
| no_cut_2.fastq.gz | no CRISPR-C controls | DNA-seq | Amplicon seq | 60,734,996 | 75 | 4,555,124,700 | 1610.1 | 51,282,406 | 84.44% |
| no_cut_3.fastq.gz | no CRISPR-C controls | DNA-seq | Amplicon seq | 48,033,762 | 75 | 3,602,532,150 | 1273.4 | 40,514,957 | 84.35% |
| GFP_1.fastq.gz | eGFP+ population replicate | DNA-seq | Amplicon seq | 41,119,039 | 75 | 3,083,927,925 | 1090.1 | 34,491,745 | 83.88% |
| GFP_2.fastq.gz | eGFP+ population replicate | DNA-seq | Amplicon seq | 63,371,869 | 75 | 4,752,890,175 | 1680.0 | 53,391,261 | 84.25% |
| GFP_3.fastq.gz | eGFP+ population replicate | DNA-seq | Amplicon seq | 29,569,377 | 75 | 2,217,703,275 | 783.9 | 24,932,246 | 84.32% |
| DN_1.fastq.gz | eGFP- population replicate | DNA-seq | Amplicon seq | 31,492,247 | 75 | 2,361,918,525 | 834.9 | 26,507,906 | 84.17% |
| DN_2.fastq.gz | eGFP- population replicate | DNA-seq | Amplicon seq | 51,934,768 | 75 | 3,895,107,600 | 1376.8 | 43,454,444 | 83.67% |
| DN_3.fastq.gz | eGFP- population replicate | DNA-seq | Amplicon seq | 23,450,279 | 75 | 1,758,770,925 | 621.7 | 19,574,302 | 83.47% |

| File | Genotype | Sequencing type | Sequencing Method | Number of reads | Mean read length | Average reads quality >=3( | PCR primers contained reads |
|---|---|---|---|---|---|---|---|
| **Junction sequencing** | | | | | | | |
| PC9_WT.fastq.gz | wide type PC9 with EGFR CRISPR-C | DNA-seq | Amplicon seq | 394,155 | 150 | 86,355 | 72,641 |
| PC9_UIMC1.fastq.gz | *UIMC1-/-* mutant PC9 with EGFR CRISPR-C | DNA-seq | Amplicon seq | 350,664 | 150 | 94,612 | 64,401 |

| File | Genotype | Sequencing type | Sequencing Method | Number of reads | Mean read length | Total bases | Coverage | Mapped reads | Mappability |
|---|---|---|---|---|---|---|---|---|---|
| **RNA-seq** | | | | | | | | | |
| PC9-Parental-1 | PC9 parental cells sensitive to Osimertinib | RNA-seq | Illumina, PE150 | 264,724,066 | 150 | 39,708,609,900 | 343.6 | 229,053,821 | 86.53% |
| PC9-Parental-2 | PC9 parental cells sensitive to Osimertinib | RNA-seq | Illumina, PE150 | 267,632,832 | 150 | 40,144,924,800 | 347.5 | 231,657,724 | 86.56% |
| PC9-Osi Res-1 | PC9 cells resistant to Osimertinib | RNA-seq | Illumina, PE150 | 251,052,356 | 150 | 37,657,853,400 | 326.3 | 217,510,117 | 86.64% |
| PC9-Osi Res-2 | PC9 cells resistant to Osimertinib | RNA-seq | Illumina, PE150 | 211,206,768 | 150 | 31,681,015,200 | 283.0 | 188,652,330 | 89.32% |
| HeLa-Parental-1 | HeLa parental cells sensitive to MTX | RNA-seq | Illumina, PE150 | 76,788,176 | 150 | 11,518,226,400 | 82.0 | 54,695,278 | 71.23% |
| HeLa-Parental-2 | HeLa parental cells sensitive to MTX | RNA-seq | Illumina, PE150 | 85,062,294 | 150 | 12,759,344,100 | 97.1 | 64,741,990 | 76.11% |
| HeLa-Parental-3 | HeLa parental cells sensitive to MTX | RNA-seq | Illumina, PE150 | 66,127,404 | 150 | 9,919,110,600 | 76.7 | 51,124,445 | 77.31% |
| HeLa-MTX Res-1 | HeLa cells resistant to MTX | RNA-seq | Illumina, PE150 | 70,449,960 | 150 | 10,567,494,000 | 75.7 | 50,497,418 | 71.68% |
| HeLa-MTX Res-2 | HeLa cells resistant to MTX | RNA-seq | Illumina, PE150 | 60,917,090 | 150 | 9,137,563,500 | 72.4 | 48,241,969 | 79.19% |
| HeLa-MTX Res-3 | HeLa cells resistant to MTX | RNA-seq | Illumina, PE150 | 53,211,282 | 150 | 7,981,692,300 | 59.5 | 39,662,392 | 74.54% |

| File | Genotype | Sequencing type | Sequencing Method | Number of reads | Mean read length | Total bases | Coverage | Mapped reads | Mappability |
|---|---|---|---|---|---|---|---|---|---|
| **Drosophila ovary seq** | | | | | | | | | |
| barcode07.fastq | DNA-lig4+/- | ecDNA-seq | SQK-LSK109 | 108,940 | 8,552 | 931,654,880 | 5.2 | 91,482 | 83.97% |
| barcode08.fastq | DNA-lig4+/- | ecDNA-seq | SQK-LSK109 | 103,961 | 11,384 | 1,183,492,024 | 6.6 | 101,237 | 97.38% |
| barcode09.fastq | DNA-lig4+/- | ecDNA-seq | SQK-LSK109 | 105,907 | 10,245 | 1,085,017,215 | 6 | 103,945 | 98.15% |
| barcode10.fastq | DNA-lig4-/- | ecDNA-seq | SQK-LSK109 | 96,536 | 10,688 | 1,031,776,768 | 5.7 | 93,922 | 97.29% |
| barcode11.fastq | DNA-lig4-/- | ecDNA-seq | SQK-LSK109 | 100,353 | 10,752 | 1,078,995,456 | 6 | 97,910 | 97.57% |
| barcode12.fastq | DNA-lig4-/- | ecDNA-seq | SQK-LSK109 | 97,957 | 11,920 | 1,167,647,440 | 6.5 | 96,393 | 98.40% |
| Hetz_1.fastq.gz | DNA-lig4+/- | genomic-seq | FC-131-1024 | 258,044,964 | 150 | 38,706,744,600 | 215 | 254,231,362 | 98.52% |
| Hetz_2.fastq.gz | DNA-lig4+/- | genomic-seq | FC-131-1024 | 131,380,892 | 150 | 19,707,133,800 | 109.5 | 129,396,135 | 98.49% |
| Homo_3.fastq.gz | DNA-lig4-/- | genomic-seq | FC-131-1024 | 280,759,117 | 150 | 42,113,867,550 | 234 | 276,751,931 | 98.57% |
| Homo_4.fastq.gz | DNA-lig4-/- | genomic-seq | FC-131-1024 | 315,107,290 | 150 | 47,266,093,500 | 262.6 | 310,641,374 | 98.58% |

* ecDNA and genomic sequencing coverages are calculated by the effective genome size.

| File | Source | Sequencing type | Number of read pairs | Mean Quality Score | % Bases >= 30 | Total bases | Coverage | Mapped reads | Mappability | Matched Tumor/blood ID |
|---|---|---|---|---|---|---|---|---|---|---|
| **Glioblatoma samples** | | | | | | | | | | |
| A1 | tumor | genomic-seq | 832155368 | 35.22 | 89.98 | 249,646,610,400 | 83.2155368 | 1,311,070,988 | 78.78% | 6 |
| A2 | tumor | genomic-seq | 856522930 | 35.3 | 90.49 | 256,956,879,000 | 85.652293 | 1,418,150,716 | 82.79% | 24 |
| A3 | tumor | genomic-seq | 1295672597 | 35.56 | 91.71 | 388,701,779,100 | 129.5672597 | 2,533,922,180 | 97.78% | 45 |
| A4 | tumor | genomic-seq | 776628355 | 35.63 | 92.1 | 232,988,506,500 | 77.6628355 | 1,519,336,518 | 97.82% | 034-79 |
| A5 | tumor | genomic-seq | 964742110 | 35.55 | 91.65 | 289,422,633,000 | 96.474211 | 1,812,428,472 | 93.93% | 109 |
| B1 | tumor | genomic-seq | 689160597 | 35.27 | 90.21 | 206,748,179,100 | 68.9160597 | 1,155,403,232 | 83.83% | 8 |
| B2 | tumor | genomic-seq | 852030665 | 35.3 | 90.4 | 255,609,199,500 | 85.2030665 | 1,409,570,860 | 82.72% | 028-25 |
| B3 | tumor | genomic-seq | 1022559399 | 35.59 | 91.88 | 306,767,819,700 | 102.2559399 | 1,996,762,114 | 97.64% | 47 |
| B4 | tumor | genomic-seq | 916950854 | 35.6 | 91.99 | 275,085,256,200 | 91.6950854 | 1,780,626,940 | 97.10% | 80 |
| B5 | tumor | genomic-seq | 1037788589 | 35.54 | 91.59 | 311,336,576,700 | 103.7788589 | 2,016,896,194 | 97.17% | 110 |
| C1 | tumor | genomic-seq | 920928533 | 35.31 | 90.46 | 276,278,559,900 | 92.0928533 | 1,583,673,684 | 85.98% | 008-9 |
| C2 | tumor | genomic-seq | 1036298371 | 35.41 | 90.96 | 310,889,511,300 | 103.6298371 | 1,836,317,756 | 88.60% | 26 |
| C3 | tumor | genomic-seq | 937980320 | 35.14 | 89.66 | 281,394,096,000 | 93.798032 | 1,436,043,486 | 76.55% | 61241-049 |
| C4 | tumor | genomic-seq | 735318726 | 35.43 | 91.08 | 220,595,617,800 | 73.5318726 | 1,437,055,104 | 97.72% | 81 |
| C5 | tumor | genomic-seq | 871001721 | 35.34 | 90.61 | 261,300,516,300 | 87.1001721 | 1,697,718,778 | 97.46% | 113 |
| D1 | tumor | genomic-seq | 696342855 | 35.18 | 89.85 | 208,902,856,500 | 69.6342855 | 1,019,692,056 | 73.22% | 11 |
| D2 | tumor | genomic-seq | 800463036 | 35.37 | 90.73 | 240,138,910,800 | 80.0463036 | 1,564,283,008 | 97.71% | 27 |
| D3 | tumor | genomic-seq | 682876980 | 35.34 | 90.64 | 204,863,094,000 | 68.287698 | 1,290,349,080 | 94.48% | 57 |
| D4 | tumor | genomic-seq | 850231075 | 35.6 | 91.93 | 255,069,322,500 | 85.0231075 | 1,664,130,844 | 97.86% | 82 |
| D5 | tumor | genomic-seq | 974620850 | 35.28 | 90.45 | 292,386,255,000 | 97.462085 | 1,807,847,910 | 92.75% | 119 |
| E1 | tumor | genomic-seq | 1148741671 | 35.55 | 91.63 | 344,622,501,300 | 114.8741671 | 2,253,287,760 | 98.08% | 13 |
| E2 | tumor | genomic-seq | 735672804 | 35.46 | 91.32 | 220,701,841,200 | 73.5672804 | 1,427,405,394 | 97.01% | 005-32 |
| E3 | tumor | genomic-seq | 1109230157 | 35.47 | 91.21 | 332,769,047,100 | 110.9230157 | 2,141,833,102 | 96.55% | 61102-002 |
| E4 | tumor | genomic-seq | 917781736 | 35.66 | 92.24 | 275,334,520,800 | 91.7781736 | 1,798,062,990 | 97.96% | 89 |
| F1 | tumor | genomic-seq | 930946935 | 35.13 | 89.73 | 279,284,080,500 | 93.0946935 | 1,598,772,872 | 85.87% | 019-17 |
| F2 | tumor | genomic-seq | 1036395957 | 35.62 | 92.05 | 310,918,787,100 | 103.6395957 | 2,015,579,330 | 97.24% | 40 |
| F3 | tumor | genomic-seq | 903422556 | 35.63 | 92.16 | 271,026,766,800 | 90.3422556 | 1,761,316,572 | 97.48% | 68 |
| F4 | tumor | genomic-seq | 913464997 | 35.5 | 91.44 | 274,039,499,100 | 91.3464997 | 1,777,753,122 | 97.31% | 96 |
| G1 | tumor | genomic-seq | 1105798673 | 35.51 | 91.45 | 331,739,601,900 | 110.5798673 | 2,141,945,110 | 96.85% | 61309-21 |
| G2 | tumor | genomic-seq | 1417702908 | 35.54 | 91.63 | 425,310,872,400 | 141.7702908 | 2,779,015,960 | 98.01% | 41 |
| G3 | tumor | genomic-seq | 922650782 | 35.58 | 91.9 | 276,795,234,600 | 92.2650782 | 1,794,777,166 | 97.26% | 029-73 |
| G4 | tumor | genomic-seq | 852079311 | 35.57 | 91.79 | 255,623,793,300 | 85.2079311 | 1,657,885,874 | 97.28% | 97 |
| H1 | tumor | genomic-seq | 118625649 | 35.33 | 90.83 | 35,587,694,700 | 11.8625649 | 217,688,344 | 91.75% | 006-22 |
| H2 | tumor | genomic-seq | 1181279955 | 35.56 | 91.75 | 354,383,986,500 | 118.1279955 | 2,283,987,734 | 96.67% | 44 |
| H3 | tumor | genomic-seq | 735486922 | 35.67 | 92.32 | 220,646,076,600 | 73.5486922 | 1,431,992,858 | 97.35% | 76 |
| H4 | tumor | genomic-seq | 963305261 | 35.55 | 91.66 | 288,991,578,300 | 96.3305261 | 1,882,195,950 | 97.69% | 102 |
| 6 | blood | genomic-seq | 252471205 | 35.44 | 91.08 | 75,741,361,500 | 25.2471205 | 495,162,852 | 98.06% | A1 |
| 24 | blood | genomic-seq | 315310395 | 35.43 | 91 | 94,593,118,500 | 31.5310395 | 619,071,560 | 98.17% | A2 |
| 45 | blood | genomic-seq | 309190334 | 35.43 | 91 | 92,757,100,200 | 30.9190334 | 607,651,752 | 98.26% | A3 |
| 034-79 | blood | genomic-seq | 317692213 | 35.45 | 91.09 | 95,307,663,900 | 31.7692213 | 623,813,648 | 98.18% | A4 |
| 109 | blood | genomic-seq | 314951770 | 35.6 | 91.89 | 94,485,531,000 | 31.495177 | 618,386,320 | 98.17% | A5 |
| 8 | blood | genomic-seq | 245818514 | 35.43 | 90.96 | 73,745,554,200 | 24.5818514 | 482,582,548 | 98.16% | B1 |
| 028-25 | blood | genomic-seq | 313115695 | 35.41 | 90.92 | 93,934,708,500 | 31.3115695 | 614,361,328 | 98.10% | B2 |
| 47 | blood | genomic-seq | 312832771 | 35.42 | 90.89 | 93,849,831,300 | 31.2832771 | 614,732,342 | 98.25% | B3 |
| 80 | blood | genomic-seq | 314645916 | 35.44 | 91.01 | 94,393,774,800 | 31.4645916 | 617,773,382 | 98.17% | B4 |
| 110 | blood | genomic-seq | 316804785 | 35.53 | 91.53 | 95,041,435,500 | 31.6804785 | 621,808,280 | 98.14% | B5 |
| 008-9 | blood | genomic-seq | 231082993 | 35.45 | 91.07 | 69,324,897,900 | 23.1082993 | 453,654,256 | 98.16% | C1 |
| 26 | blood | genomic-seq | 322761469 | 35.43 | 90.98 | 96,828,440,700 | 32.2761469 | 633,798,292 | 98.18% | C2 |
| 61241-049 | blood | genomic-seq | 322229818 | 35.5 | 91.36 | 96,668,945,400 | 32.2229818 | 633,395,330 | 98.28% | C3 |
| 81 | blood | genomic-seq | 324909833 | 35.5 | 91.35 | 97,472,949,900 | 32.4909833 | 638,368,808 | 98.24% | C4 |
| 113 | blood | genomic-seq | 320118837 | 35.6 | 91.86 | 96,035,651,100 | 32.0118837 | 628,915,168 | 98.23% | C5 |
| 11 | blood | genomic-seq | 239453551 | 35.41 | 90.91 | 71,836,065,300 | 23.9453551 | 469,655,290 | 98.07% | D1 |
| 27 | blood | genomic-seq | 322008852 | 35.39 | 90.77 | 96,602,655,600 | 32.2008852 | 632,493,002 | 98.21% | D2 |
| 57 | blood | genomic-seq | 307008204 | 35.36 | 90.63 | 92,102,461,200 | 30.7008204 | 602,675,004 | 98.15% | D3 |
| 82 | blood | genomic-seq | 316887139 | 35.49 | 91.3 | 95,066,141,700 | 31.6887139 | 623,217,130 | 98.33% | D4 |
| 119 | blood | genomic-seq | 328763101 | 35.55 | 91.63 | 98,628,930,300 | 32.8763101 | 645,549,400 | 98.18% | D5 |
| 13 | blood | genomic-seq | 266459114 | 35.47 | 91.19 | 79,937,734,200 | 26.6459114 | 524,097,792 | 98.34% | E1 |
| 005-32 | blood | genomic-seq | 321202546 | 35.47 | 91.2 | 96,360,763,800 | 32.1202546 | 631,245,506 | 98.26% | E2 |
| 61102-002 | blood | genomic-seq | 310088545 | 35.42 | 90.91 | 93,026,563,500 | 31.0088545 | 609,635,992 | 98.30% | E3 |
| 89 | blood | genomic-seq | 315633534 | 35.58 | 91.73 | 94,690,060,200 | 31.5633534 | 621,817,650 | 98.50% | E4 |
| 019-17 | blood | genomic-seq | 207510132 | 35.5 | 91.38 | 62,253,039,600 | 20.7510132 | 407,265,782 | 98.13% | F1 |
| 40 | blood | genomic-seq | 342044559 | 35.48 | 91.29 | 102,613,367,700 | 34.2044559 | 672,051,686 | 98.24% | F2 |
| 68 | blood | genomic-seq | 317362233 | 35.46 | 91.13 | 95,208,669,900 | 31.7362233 | 623,763,390 | 98.27% | F3 |
| 96 | blood | genomic-seq | 331630786 | 35.54 | 91.57 | 99,489,235,800 | 33.1630786 | 650,959,404 | 98.15% | F4 |
| 61309-21 | blood | genomic-seq | 242028072 | 35.44 | 91.04 | 72,608,421,600 | 24.2028072 | 475,064,776 | 98.14% | G1 |
| 41 | blood | genomic-seq | 317865406 | 35.39 | 90.8 | 95,359,621,800 | 31.7865406 | 624,294,412 | 98.20% | G2 |
| 029-73 | blood | genomic-seq | 314536895 | 35.38 | 90.72 | 94,361,068,500 | 31.4536895 | 616,930,474 | 98.07% | G3 |
| 97 | blood | genomic-seq | 317656311 | 35.44 | 90.99 | 95,296,893,300 | 31.7656311 | 624,333,344 | 98.27% | G4 |
| 006-22 | blood | genomic-seq | 259975670 | 35.38 | 90.75 | 77,992,701,000 | 25.997567 | 510,105,790 | 98.11% | H1 |
| 44 | blood | genomic-seq | 311976313 | 35.36 | 90.64 | 93,592,893,900 | 31.1976313 | 612,117,338 | 98.10% | H2 |
| 76 | blood | genomic-seq | 300777199 | 35.32 | 90.42 | 90,233,159,700 | 30.0777199 | 590,603,646 | 98.18% | H3 |
| 102 | blood | genomic-seq | 307140022 | 35.43 | 90.99 | 92,142,006,600 | 30.7140022 | 603,700,660 | 98.28% | H4 |
